# TEAD4 regulates apical domain homeostasis and cell-positioning to maintain the trophectoderm lineage during preimplantation mouse embryo development

**DOI:** 10.1101/2025.11.18.689016

**Authors:** Rebecca Collier, Martina Bohuslavová (née Stiborová), Michaela Vaškovičová, Aleksandar I. Mihaljović, Andrea Hauserová, Valeriya Zabelina, Monika Fluks, Lenka Gahurová, Dávid Drutovič, Alexander W. Bruce

## Abstract

In mammalian preimplantation embryos, different cell lineages occupy specific niches. For example, the outer trophectoderm (TE) comprises a monolayer of epithelialized cells surrounding the inner-cell mass (ICM) and blastocyst cavity. In mice, TEAD4 is known as a transcription factor that regulates TE-specific genes in a polarity-dependent manner during TE specification. Here we show that it also maintains blastocyst TE integrity, as knocking down (KD) *Tead4* via clonal siRNA causes abnormal morphology of outer-cell apical domains, which correlates with the atypical contribution of *Tead4*-KD cell clones to an enlarged ICM throughout blastocyst maturation; with only minimal feedback on established apical polarity. Light-sheet live-cell embryo imaging reveals these cells either actively migrate into the ICM, sometimes involving apical domain abscission, or are positioned post-division, linking disrupted apical morphology to cell repositioning. RNA-Seq data indicate TEAD4 regulates genes related to the cytoskeleton, particularly actin, and cell adhesion, which we propose are required for the appropriate maintenance of the spatial positioning of specified TE cells in the blastocyst. Indeed, knocking down *Tead4* in combination with two identified target genes, the atypical GTPases *Rnd1* and *Rnd3,* partially rescues aberrant outer-to-inner cell allocations but does not influence the onset of apical domain morphological abnormalities. These findings indicate that *Tead4* and its regulated transcriptome actively contribute to the maintenance of the outer TE lineage until the peri-implantation stage.

## INTRODUCTION

During early mouse embryo development, three distinct lineages form within the blastocyst. The trophectoderm (TE), which later develops into the placenta, is a surface layer of epithelial cells that surrounds a fluid-filled cavity and an inner-cell mass (ICM). The primitive endoderm (PrE) arises within the ICM, forming a monolayer at its interface with the cavity, and eventually contributes to the yolk sac. The epiblast (EPI) consists of pluripotent cells located deep inside the ICM and serves as the source of cells for the future foetus^1–4^.

The first cell fate decision begins at the 8- to 16-cell stage and continues through the 16-to 32-cell stage. During this time, cells that are polarised along their apical-basolateral axis become actively segregated from inner unpolarized cells and ultimately reside on the outer surface of the embryo. This segregation results in the formation of the polarised TE on the outside and an apolar ICM on the inside. Thus, the ICM is composed of primary and secondary cells generated during the subsequent rounds of cell cleavage division^5,6^. The second cell fate decision occurs later within the blastocyst ICM, where cells either specify as the EPI or differentiate into the PrE^7–9^. After cell cleavage, the spatial positioning of 16- and 32-cell stage blastomeres is linked to the unequal inheritance of outer-cell actomyosin-driven apical constriction. Increased contractility causes some cells to internalise^10,11^, in a process closely associated with impaired apical-basolateral polarity^12–16^. Outer-cells lacking polarity proteins or with experimentally depleted polarity factors internalise into the ICM^17–19^, indicating that inherited polarity acts to inhibit contractility and cell internalisation^10,11^. Maintaining outer-cell polarity is central to TE specification, as the Hippo-pathway effector AMOT is sequestered to contactless apical domains enriched in polarity factors, preventing YAP1 phosphorylation. Unphosphorylated YAP1 translocates to the nucleus and forms TEAD4-YAP1 complexes, transcriptionally activating TE-related genes, such as *Cdx2*. In contrast, apolar inner-cells exhibit active Hippo-signalling, leading to cytoplasmic retention of phospho-YAP1, failure to activate TE genes^20,21^, and maintenance of pluripotency^22^. Additionally, asymmetric inheritance of keratin intermediate filaments has been reported to stabilise the outer-cell apical cortex, further promoting Hippo-pathway suppression and TE fate^23^.

Our previous work showed that RNAi knockdown (KD) of *Tead4* not only impairs TE differentiation, aligning with genetic knockout models^24,25^, but leads to increased contribution of *Tead4* KD clones to the late (E4.5) blastocyst ICM. Immunofluorescence revealed that these supernumerary ICM cells predominantly expressed EPI transcription factor profiles rather than those of PrE cells, suggesting that TEAD4-mediated TE-specification cues prime later PrE differentiation within the ICM^26^. Notably, these effects occurred without any overt polarity defects. However, apical-basolateral polarity initiation at the 8-cell stage is reported to involve the partially redundant roles of the *Tead4* and *Tfap2c* genes, and the small GTPase RHOA^15^. Collectively, while TEAD4 promotes TE differentiation by inducing genes such as *Cdx2* through polarity-dependent Hippo-signalling suppression^20,21^, these data also indicate TEAD4 may also actively maintain TE-fated blastomeres in outer positions, supporting the “polarity-dependent cell-positioning model”^27^ and prompting further investigation in this study.

We found that clonal *Tead4* KD in preimplantation mouse embryos leads to an atypical allocation of outer-cell clones to the ICM, which peaks during blastocyst maturation after the 32-cell stage. This is not associated with the formation of apolar outer cells, though the expression of the apical polarity protein PARD6B^28,29^ is reduced and the expression of other Hippo-signalling components is also reduced, indicating a degree of functional TEAD4-mediated feedback. Rather, such cellular misallocations result from a combination of asymmetric cell divisions and apical domain constriction. In addition, we report mechanisms of unconventional cell division associated with the formation of apical domain blebs or the abscission of apical domains that result in cell internalisation independent of cell division. RNA-Seq analysis revealed significant transcriptome differences between *Tead4* KD clones and their non-targeted sister outer-cells, mainly in genes related to the cytoskeleton and actin organisation, plus cell adhesion. Targeting the identified actin-related atypical small GTPases *Rnd1* and *Rnd3* candidate genes partially rescued the *Tead4* KD-induced internalisation phenotypes, although the induced apical domain distortions persisted. These findings show *Tead4* plays a novel role in maintaining apical domain stability and outer cell spatial positioning, ensuring TE-specified cells remain appropriately positioned during blastocyst maturation.

## RESULTS

### Clonal *Tead4* knockdown (KD) induces atypical outer-to-inner cell allocation and distorted apical domain morphology from the 32-cell stage

Previously, we found that microinjecting 2-cell stage mouse embryos with *Tead4* dsRNA reduces the contribution of confirmed *Tead4* KD clones to the PrE lineage and biases them towards EPI in late blastocysts (E4.5). We interpreted this as evidence that TEAD4 primes PrE fate in secondary ICM founders derived from outer 16-cell stage blastomeres, by forming active TEAD4-YAP1 complexes via polarity-dependent Hippo suppression, which are inhibited in the *Tead4* KD clones. Additionally, *Tead4* KD clones (comprising 50% of the embryo) were more frequently located in the E4.5 ICM than the outer TE, resulting in larger ICMs^26^. These findings suggest that TEAD4 not only drives TE specification/differentiation^20,21^ but also maintains proper outer positioning of TE-specified blastomeres; a hypothesis we explore further in this study.

We repeated the 2-cell stage microinjection RNAi strategy, co-injecting a *Tead4*-specific siRNA (si*Tead4*) together with histone H2B-RFP mRNA, producing embryos with both fluorescently marked *Tead4* KD and unmarked control clones. We performed the same approach using a non-targeting control siRNA (siNTC), as a control. Embryos were cultured for 48, 54, 60 or 72 hours (equivalent to E3.5, E3.75, E4.0 & E4.5), fixed, immuno-fluorescently (IF) stained, and imaged by confocal microscopy to assess clonal contributions to outer positions and the ICM (Fig. 1A). At all stages, si*Tead4* injections effectively abolished TEAD4 protein expression in marked clones (Figs. 1B&C and S1A&B). We also performed whole embryo *Tead4* KD, followed by RT-qPCR analysis at E3.5 (Fig. S1D-F), additionally validating the efficiency of the si*Tead4* construct.

**Figure 1:**
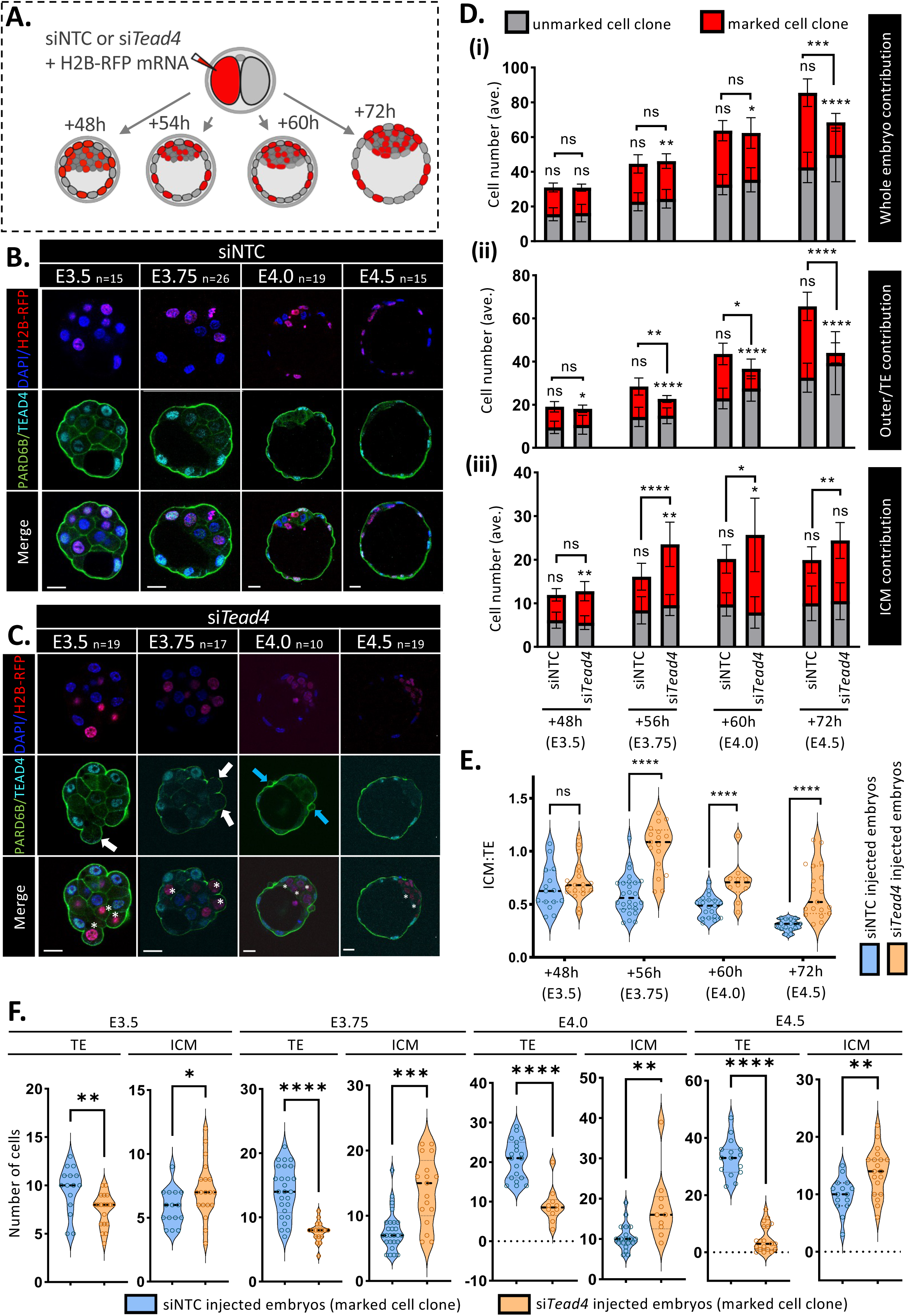
Clonal *Tead4* KD induces atypical cell allocation events during blastocyst maturation. **A)** Schematic of siRNA microinjection procedure generating staged blastocysts containing either control (siNTC) or *Tead4* KD (si*Tead4*) marked clones — comprising 50% of all cells; allowing assessment of clone contributions to the outer/TE or ICM positions. **B)** Example confocal micrographs (single merged z-sections) of IF-stained siNTC-injected blastocysts (targeted clones – red nuclei); PARD6B (green) & TEAD4 (cyan), with DAPI-DNA counterstain (blue). Scale bars = 20 μm; number of embryos per group (n) indicated (see also Fig. S1 for greyscale images). **C)** Similar images as in B), but for si*Tead4*-injected blastocysts. Clonal *Tead4* KD cells are marked with asterisks. Morphological abnormalities such as apical domain blebs and distortions are indicated with white arrows (E3.5 & E3.75). Vesicle-like structures containing PARD6B are marked with blue arrows (E4.0). **D)** Quantification of clone contributions to total, outer/TE, and ICM cell numbers across different blastocyst stages: unmarked (grey) versus marked (red) clones within siNTC and si*Tead4* blastocysts (data are also presented as violin plots and illustrative concentric pie-charts of percentage clone composition in outer/TE and ICM lineages; Fig. S1). Significant differences and standard deviations between control/experimental groups, and between unmarked/marked clones within groups, are indicated (*p<0.05). **E)** ICM:outer/TE cell number ratios in blastocysts containing marked siNTC (blue) and si*Tead4* (orange) clones at specified stages. Significant differences are marked (*p<0.05), with median values shown by dashed lines. **F)** Comparison of outer/TE and ICM cell number contributions of only the marked siNTC (blue) versus si*Tead4* (orange) clones, at the indicated stages. Median contributions shown by dashed lines, with significant inter-group differences indicated (*p<0.05). *All clonal allocation raw data are summarised in S.Tabs. 1a-e. Statistical analyses were performed based on determined data distribution (normal or non-normal) and application of appropriate tests; details in supplementary statistics Excel workbook*.

We then compared the development of embryos microinjected with si*Tead4* versus siNTC (see S.Tabs. 1a-e for detailed cell counts). In siNTC controls, blastocyst total cell numbers increased steadily from E3.5–E4.5, with marked and unmarked clones contributing equally, indicating no adverse developmental impact. In si*Tead4* embryos, total cell numbers were comparable to controls from E3.5–E4.0, but by E3.75, marked *Tead4* KD clone numbers were slightly reduced, with evident compensation from unmarked clones (extending to the E4.0 stage). At E4.5, si*Tead4* blastocysts had significantly fewer total cells on average (68.5±3.28 vs. 85.5±1.7), mainly due to fewer marked clone cells, despite partial compensation from unmarked clones (Figs. 1D and S1A&B). Overall, embryo cell numbers were similar until E4.0, after which si*Tead4* embryos had fewer overall cells, exclusively in the marked clone.

We next assessed marked and unmarked clone contributions to outer and ICM positions (Figs. 1D-F and S1A-C). In siNTC controls, there was no significant difference between clones in either lineage at any stage, indicating unbiased contribution and normal development (see also clonal TE/ICM percentage composition; Fig. S1C). Additionally, the ICM:TE cell ratio decreased appropriately over time, demonstrating normal blastocyst development after siNTC microinjection. In contrast, si*Tead4* clones contributed only 12% of cells to a significantly smaller TE by E4.5, consistent with the characterised role of *Tead4* in TE development^20,21^. Conversely, si*Tead4*-microinjected blastocysts had larger ICMs, with over half (57%) of the cells originating from the marked clone, and significantly increased marked ICM cell numbers compared to marked ICM clones in siNTC controls. These findings confirm a strongly biased ICM contribution from *Tead4* KD clones, even in embryos with significantly fewer marked clones overall, resulting in late (E4.5) blastocysts with predictably reduced TE but atypically expanded ICM.

At E3.75 and E4.0, where overall cell numbers were similar between si*Tead4* and siNTC embryos, si*Tead4* clones made up a larger proportion of the total ICM at 59% and 68%, compared to 48% and 51%, respectively, for the siNTC clone controls. This led to overall significantly smaller TE and larger ICM sizes. Accordingly, the reduced TE contribution was not due to a smaller clone size (as seen at E4.5) but indicated a genuine positional bias caused by *Tead4* KD. Notably, at E3.5, no significant differences were observed in overall TE or ICM cell numbers. However, within just six hours (from E3.5 to E3.75), si*Tead4* embryos exhibited a marked shift in positional allocation, averaging 6.32 fewer TE cells and 6.25 more marked ICM cells. This trend continued at E4.0, with an additional 7.38 marked ICM cells. This suggests a post-cell division mechanism whereby outer blastomeres at the E3.5 stage were atypically redistributed internally after the initial formation of primary and secondary ICM founders. Morphologically, the outer cells of si*Tead4* clones displayed distinctive abnormalities in their apical membranes, which we termed apical blebs (Figs. 1B&C and S1A&B). These abnormalities were particularly evident at E3.5 and E3.75, coinciding with the observed large cell positioning shifts. Additionally, vesicle-like structures containing the apical polarity factor PARD6B were found on outer embryo surfaces (Figs. 1B&C and S1A&B), potentially representing events of apical bleb abscission; events, theoretically resulting from increased contractility, that could facilitate the reallocation of marked *Tead4* KD outer-cells to the ICM.

At E3.5, si*Tead4*-treated blastocysts show no significant difference in total TE or ICM cell numbers compared to controls, but the microinjected clone did contribute less to the TE and more to the ICM, indicating early clonal allocation differences starting before the 32-cell stage; likely during the 16- to 32-cell transition. These data also reveal complete compensatory regulation by the unmarked clone, balancing overall TE and ICM cell allocation in E3.5 stage si*Tead4*-treated embryos. Additionally, at E4.0, unmarked ICM cells in si*Tead4*-treated blastocysts were actually reduced in number compared to those at E3.75 (and the equivalent unmarked cells of siNTC embryos), suggesting further compensatory mechanisms attempting to regulate overall ICM size, following the atypical influx of marked clones. Similar reductions in marked si*Tead4* ICM cells between E4.0 and E4.5 confirm such regulative mechanisms (exemplified by reducing ICM:TE/outer cell ratios from the E3.75 peak; Fig. 1E).

TEAD4-YAP1 complexes in outer-cells activate TE-specific genes like *Cdx2* through polarity-dependent suppression of Hippo-signalling^20,21,26^. We confirmed that global si*Tead4* KD (achieved by microinjecting both 2-cell stage blastomeres) reduces the expression of TE marker mRNAs, such as *Gata3*^30^*, Tfap2c*^31^ and *Cdx2*^32^ (Fig. S1D-F), by the E3.5 blastocyst stage. To determine if such genes also contribute to the atypical outer cell-to-ICM cell allocations, we clonally disrupted *Tfap2c* and *Cdx2* using dsRNA or CRISPR-Cas9 via similar microinjection methods, then examined blastocyst development (Figs. S2 & S3). Despite successful, clone-specific loss of detectable TFAP2C and CDX2 protein expression, we found no evidence of reduced outer cell contribution or increased ICM contribution. Additionally, there were no changes in the morphology of marked outer clone apical domains. These results suggest that, although TEAD4 regulates *Tfap2c* and *Cdx2* expression, neither target gene alone mediates the specific regulation of outer cell positioning attributable to *Tead4*. Instead, TEAD4 likely influences this process through its broader transcriptional network.

Overall, these data show that clonal si*Tead4* KD leads to late-blastocysts (E4.5) with significantly more ICM and fewer TE cells. This occurs because marked clones are preferentially allocated to the ICM starting as early as the 32-cell stage (E3.5), despite compensatory masking in unmarked clones maintaining an appropriate ICM:TE/outer cell ratio. However, between E3.5 and E3.75 marked clones robustly and atypically populate the ICM, likely through direct outer cell-to-ICM re-allocation/internalisation events, supported by morphological apical domain distortions in outer-cells. This abnormal allocation persists into the late (E4.5) blastocyst stage, with varying compensation within unmarked and marked ICM and outer residing clones. However, similar targeted clonal dysregulation of confirmed TEAD4 target TE-specific genes, *Tfap2c* and *Cdx2*, is unable to elicit similar allocation phenotypes. Thus, TEAD4 plays a key role in maintaining relative cell spatial positioning and ensuring TE cells occupy outer lineage-appropriate positions within the maturing blastocyst.

### si*Tead4* KD clones internalise via typical asymmetric cell divisions and apical domain constriction, and unconventional mechanisms associated with apical bleb formation and abscission

Having identified an early blastocyst window (E3.5-E3.75), characterised by atypical outer cell-to-ICM allocation, apical bleb formation, and PARD6B-containing surface vesicles in *siTead4* KD clones within fixed embryos, we employed fluorescent light-sheet microscopy 4D live-cell imaging to monitor the dynamics by which these phenotypes emerge, and with greater spatiotemporal resolution. We co-microinjected single blastomeres from transgenic 2-cell stage embryos expressing a fluorescent plasma membrane-localised Tomato reporter (mT^33^) with either siNTC or si*Tead4*, and mRNA encoding histone H2B-Venus (as an injection/lineage marker). Microinjected embryos were cultured to 89 hours post hCG and then imaged at intervals of 10 minutes for the following 16.5 hours to assess the dynamic behaviour of H2B-Venus-marked control and *Tead4* KD clones, compared with the unmarked clones (marked only by mT) from the same embryos (Fig. 2A).

**Figure 2:**
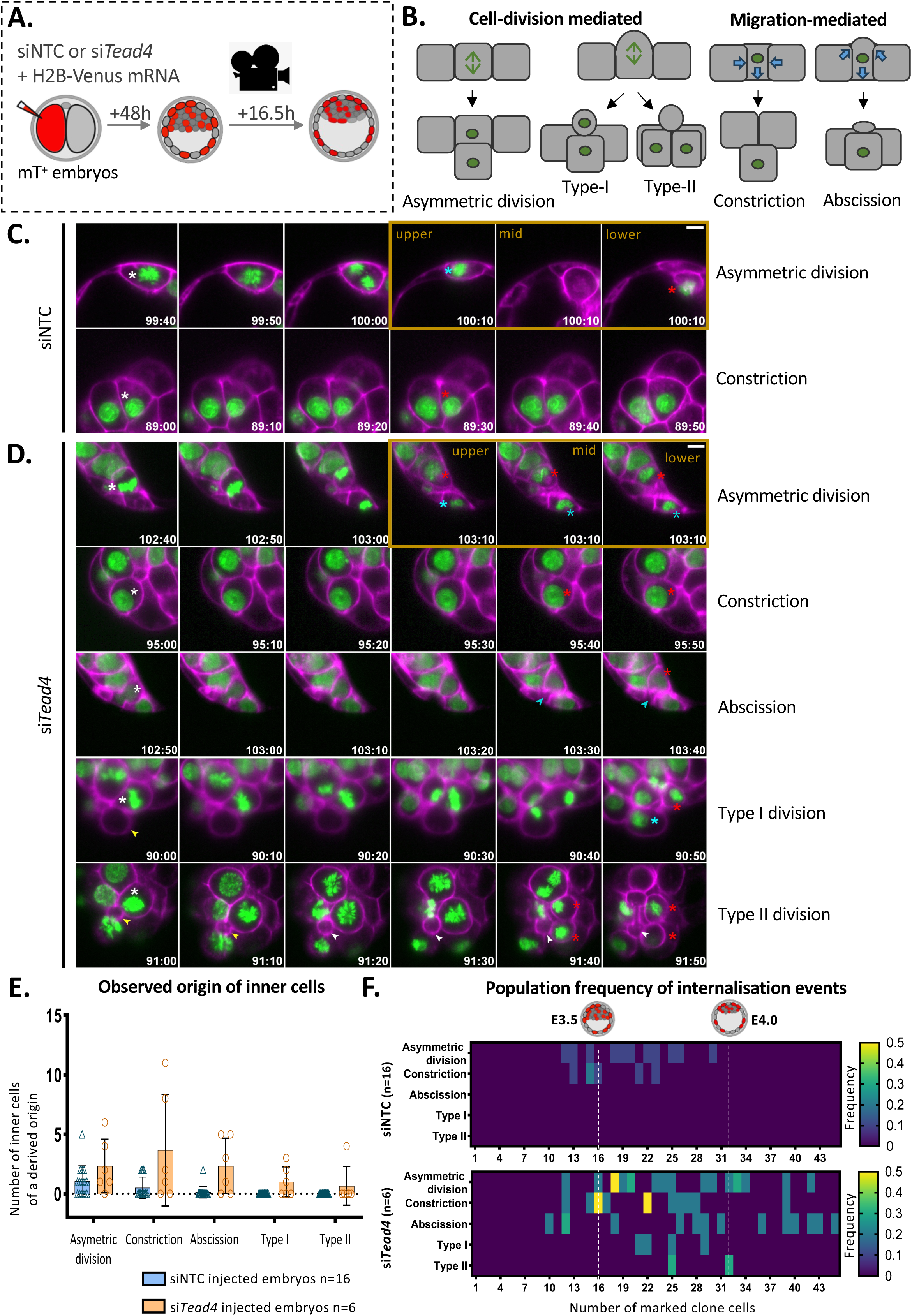
4D light-sheet live-cell embryo imaging of atypical blastocyst internalisation of marked *Tead4* KD cell clones. **A)** Experimental schematic of 2-cell stage microinjections of transgenic membrane-Tomato expressing (mT^+^) 2-cell embryos used to generate early 32-cell stage blastocysts (after 48 hours of culture) containing either histone H2B-Venus marked control (siNTC) or *Tead4* KD (si*Tead4*) clones (comprising 50% of all cells), subject to 16.5 hours of light-sheet live-cell imaging (10 minute intervals); allowing dynamic assessment of individual clone contributions to the outer/TE or ICM positions **B)** Schematic representations of the cell internalisation mechanisms observed for marked (H2B-Venus+) clones during blastocyst maturation. **C)** Examples of cell internalisation events observed in embryos containing marked siTNC clones (i.e. asymmetric cell divisions and apical domain constriction), as successive time-lapse imaging frames (timings reflect hh:mm post-hCG superovulation injection and scale bar 10µm). The cell marked with a white asterisk displays the phenotype described next to the image, cyan and red asterisks denote derived outer and inner cells, respectively. For asymmetric cell divisions, the last three frames represent the same time point in three consecutive z-sections (highlighted by the orange box). Pseudo-colour scheme: magenta – transgenic embryo mT reporter (as a single z-plane), and green – histone H2B-Venus microinjection marker (as a maximum projection from several planes located above and below the selected mT plane). Micrographs taken from the illustrative blastocyst embryo depicted in supplementary movie 1. **D)** Examples of cell internalisation events observed in embryos containing marked si*Tead4* clones (i.e. asymmetric cell divisions, apical domain constriction, plus the additional atypical blastocyst internalisation mechanisms, summarised in panel B), as successive time-lapse imaging frames (timings reflect hh:mm post-hCG superovulation injection and scale bar 10µm). Cells marked with a white asterisk exhibit the phenotype described next to the image, cyan and red asterisks denote outer and inner cells, respectively. Blue arrowheads denote remnants of the apical domain following apical abscission (following cell internalisation independent of cell division), yellow arrowheads indicate formation of apical blebs (associated with type I and type II cell divisions), and white arrowheads highlight the formation of surface vesicles (devoid of segregated chromosomes, after type II cell divisions). For asymmetric cell divisions, the last three frames represent the same time point in three consecutive z-sections (highlighted by the orange box). Pseudo-colour scheme: magenta – transgenic embryo mT reporter (as a single z-plane), and green – histone H2B-Venus microinjection marker (as a maximum projection of several planes located above and below the selected mT plane). Micrographs taken from the illustrative blastocyst embryos depicted in supplementary movie 1. **E)** The summed mechanistic origin of all observed internalised cells, summarised in B), at the end of the imaging period (n=number of embryos in each group) for embryos under siNTC- and si*Tead4*-treatment conditions, as illustrated in panels C) and D). **G)** Heatmap showing the frequency of internalisations at a given developmental stage (delineated by total number of marked clones – representing 50% of the embryo). Internalisations are normalised to the number of blastocysts imaged. *Detailed lineage trees for all analysed embryos are provided in Figs. S4 & S5. Note, the details of the ultimate origin of injected/marked cells at the end of the movie and the observed frequencies of internalisation events are contained in S.Tabs. 5a&5b*.

We found that the marked clones of si*Tead4*-treated blastocysts are internalised through an increased occurrence of classically described internalisation mechanisms, plus the emergence of other undescribed processes. Broadly, these can be divided into division-mediated and migration-mediated events. Internalisations linked to an increase in classical asymmetric divisions, in which an outer cell divides parallel to the apical-basal axis (a perpendicular division plane) to generate one inner and one outer daughter cell (as described for the generation of both primary and secondary ICM founder cells, prior to the 32-cell stage^34^). We also observed atypical division-mediated internalisations in cells with apical blebs that appear to alter the axis of cell division, leading to what we termed either ‘type-I’ or ‘type-II’ divisions. In type-I divisions, cells divide asymmetrically, with one daughter retaining a surface position and the apical bleb membrane while the other cell internalises. In type-II divisions, the division axis is perpendicular to the apical-basal axis, but both daughters lose surface contact and internalise to the ICM, often leaving a surface vesicular apical bleb remnant. Migration-mediated internalisations again include the previously described mechanisms of conventional constriction of a small apical domain, whereby the cell internalises by progressively reducing its surface contact^10,11,17,35^, and unusual apical abscission events. The latter involves the complete resection of the apical domain from non-dividing outer cells, which, post-abscission, yields an inner cell and a vesicular remnant on the blastocyst surface (Fig. 2B&D).

We used light-sheet imaging data to construct annotated lineage trees for each siNTC-and si*Tead4*-treated imaged blastocyst, detailing the incidences of each category of internalisation event and cellular apoptosis (Figs. S4 & S5). Using this information, we determined the developmental origin and timings of all observed internalisation events (Fig. 2E&F, respectively). We found that classical asymmetric cell divisions^34^ and apical constrictions^10,11,17,35^ were the most prevalent origins of marked clone internalisation in si*Tead4*-treated blastocysts, followed by apical abscissions and type-I and type-II division-associated processes (the latter observed in only one embryo, twice). Furthermore, such internalisations peaked between the 32-cell (E3.5) early- and mid- (E4.0) blastocyst stages and waned towards the end of the imaging/tracing period (see heatmap of recorded internalisation events, Fig. 2F); in strong accord with results from fixed embryo samples (Figs. 1 and S1). Moreover, we also observed enhanced frequencies of the classically described internalisation mechanisms before the 32-cell stage (E3.5), plus the emergence of the atypical internalisation mechanism of apical abscission. These observations align with fixed-embryo data showing that the marked *Tead4* KD clone contributes statistically more inner cells than its unmarked counterpart by the early blastocyst stage (E3.5), despite overall compensatory regulation from the unmarked clone in terms of total outer and ICM cell number (Figs. 1 and S1). Interestingly, we did observe unanticipated internalisation of marked siNTC clones during the post-32-cell (E3.5) blastocyst maturation phase (Fig. 2B, E&F). While these data indicate outer-to-inner cell transitions are possible beyond the 32-cell stage under control conditions, their frequency was considerably reduced and almost exclusively restricted to the classically described internalisation mechanisms; (i.e. asymmetric cell division and apical domain constriction). Moreover, post-32-cell-stage internalisation of marked siNTC clones was limited to the earlier phase of blastocyst maturation, whereas similar mechanisms, plus those associated with apical domain abscission and atypical cell divisions, persisted longer and at a significantly higher frequency in marked *Tead4* KD clones (Fig. 2F).

Overall, these data, together with those from fixed blastocyst samples (Figs. 1 and S1), show that marked *Tead4* KD clones become abnormally internalised into the ICM, starting around the 32-cell stage (E3.5) and continuing throughout blastocyst maturation; contributing to an increased number of ICM cells. The mechanisms involve: increased levels of conventional asymmetric cell division and apical domain constriction, and unusual processes related to apical domain abscission and defective cell divisions associated with apical bleb formation. Collectively, these findings support a homeostatic role for *Tead4* in ensuring the correct relative spatial positioning of TE-specified cells within the blastocyst.

### Clonal *Tead4* KD reduces, but does not ablate, outer-cell apical polarity and disrupts YAP1 localisation

From the 16-cell stage, outer-cell TE specification is tightly controlled by intracellular apical-basolateral polarity, which suppresses Hippo-signalling through sequestration of AMOT at the apical domain. This ensures nuclear TEAD4-YAP1 complex formation and TE-related gene expression^20,21,36^. Indeed, chemical inhibition of ROCK1/2 (i.e. Rho-associated coiled-coil containing protein kinases) completely disrupts outer-cell polarity, characterised by uniform plasma membrane distribution of polarity factors, ectopic AMOT at basolateral sites, activation of Hippo-signalling, and loss of CDX2 expression and TE-specification^37,38^. Apical polarity establishment begins at the late 8-cell stage, preceding spatial segregation of blastomeres, in two steps: first, formation of a cortical actomyosin network; second, polarity protein recruitment. This process is asynchronous, dependent on redundant *Tead4* and *Tfap2c* function, plus active RHOA protein, with early polarising blastomeres more likely to contribute to the TE lineage^15,16,39^. However, in addition to specification, maintaining polarity is essential for correct outer TE cell positioning. Time-lapse imaging shows that loss of polarity leads to ICM internalisation of outer-cells ^17^, and clonally targeting apical polarity proteins like PARD3 and PRKCZ causes outer-cells to similarly re-localise^18^. Hence, establishing apical-basolateral polarity promotes TE gene expression but also ensures proper spatial placement of outer TE cells, a concept we termed the “polarity-dependent cell-positioning model”^27^. To investigate this further, we examined whether impaired polarity and ectopic Hippo-signalling occur in blastocysts containing *Tead4* KD clones.

Our analysis of fixed blastocysts revealed that outer-cells in both siNTC and si*Tead4* marked clones displayed apical membranes appropriately associated with PARD6B expression, including within apical blebs or formed surface vesicles associated with outer cell-to-ICM reallocation (Figs. 1, 2 and S1). This shows that apical-basolateral polarity is established and remains intact in *Tead4* KD outer clones. Indeed, similar PARD6B expression was consistently observed in E3.5 blastocysts, also presenting with apical blebs and vesicles, in *Tead4* KD clones independently co-stained for PARD6B and YAP1 (Fig. 3A). However, in such blastocysts, whole-embryo PARD6B levels were significantly lower in those containing marked si*Tead4* clones compared to siNTC controls (Fig. 3B). Analysis of single outer cells showed this reduction was specific to the si*Tead4* clone, which had significantly lower apical PARD6B expression than unmarked outer cells within the same embryo and compared to both siNTC control outer cell clones (Fig. 3C). These findings align with previous data showing *Pard6b* mRNA levels decrease (to ∼80%) in E3.5 blastocysts derived from *Tead4* shRNA-injected zygotes^28^ and to ∼50% from zygotes targeted by efficient *Tfap2c* siRNA KD (associated with reduced apical PARD6B protein, albeit at the earlier morula stage^31,40^). We hypothesise that the si*Tead4*-induced decreases in *Tfap2c* mRNA levels (to ∼40%; Fig. S1) partly explain the significant, but incomplete, reduction in apical PARD6B in marked outer clones. In summary, si*Tead4*-marked outer-cell clones, reallocating to the ICM, show significantly reduced PARD6B levels, indicating diminished but not ablated apical polarity, most likely associated with TEAD4-mediated regulatory gene expression feedback.

**Figure 3.**
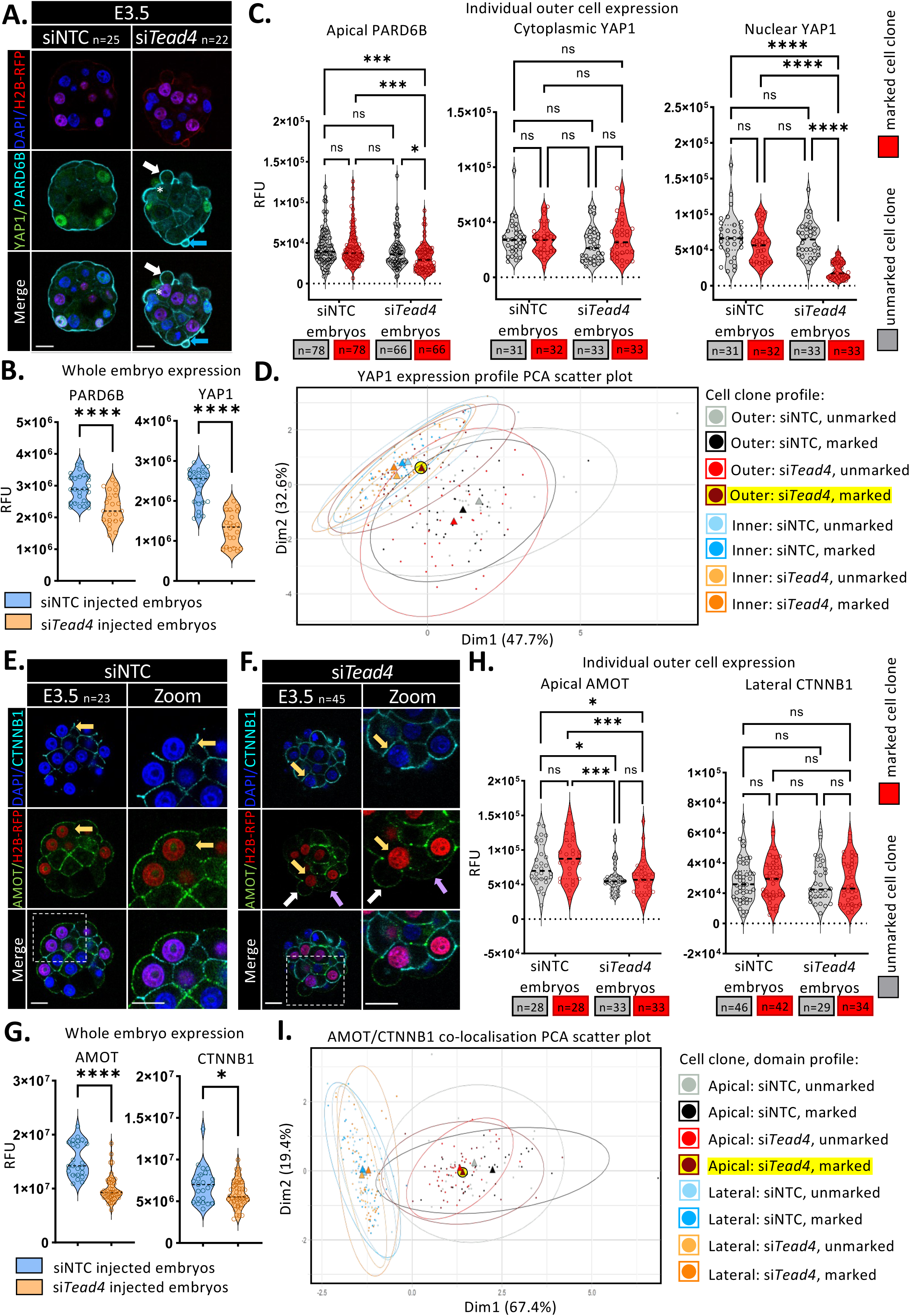
Clonal *Tead4* KD impairs, but does not ablate, outer-cell apical domain polarity and affects Hippo-signalling components. **A)** Example single z-section confocal micrographs of E3.5 blastocysts containing marked (red nuclei) clones injected with siNTC (left) or si*Tead4* (right) constructs. Embryos are IF-stained for PARD6B (cyan; an apical polarity protein) and YAP1 (green; a Hippo-pathway effector and TEAD4 co-factor), plus DAPI DNA stain (blue). Scale bars = 20 μm. The number of embryos per group (n) is indicated. Asterisks highlight atypical cytoplasmic YAP1 localisation in outer-cells, instead of the usual nuclear localisation (associated with Hippo-signalling repression), of si*Tead4* clones; similarly characterised by apical cysts and surface vesicle-like structures, indicated by white and blue arrows (see also Fig. 1). **B)** Comparison of PARD6B and YAP1 expression levels (normalised relative fluorescence units, RFU) across whole embryos at E3.5, including unmarked and marked clones, in control siNTC (blue) and si*Tead4* (orange) embryo groups; as illustrated in A). Median contributions are indicated by dashed lines; statistically significant differences between groups are marked (*p<0.05). **C)** Quantitative analysis of individual outer-cell expression levels (RFU) for apical membrane-associated PARD6B, plus cytoplasmic and nuclear localised YAP1, in siNTC and si*Tead4* blastocysts. Data are separated for unmarked (grey) and marked (red) clone populations. Median levels are denoted by dashed lines, with significant differences indicated (*p<0.05). Sample sizes (n) are noted (see Fig. S6 for similar data on ICM cell YAP1 expression). **D)** Principal Component Analysis (PCA) of subcellular YAP1 expression levels, comparing unmarked and marked clones in siNTC and si*Tead4* blastocysts (expanded in Fig. S6). Each point represents an individual cell, with ellipses denoting 95% confidence intervals. Mean cluster centres are marked with triangles – outer marked si*Tead4* KD clones highlighted in yellow. **E & F)** Similar to panel A), but reporting AMOT (green) and CTNNB1 (cyan) IF-staining in siNTC- (E) and si*Tead4-* (F) treated blastocysts, including magnified insets. White and purple arrows indicate weak but correctly localised apical AMOT in marked and unmarked outer-cells, respectively. Yellow arrows denote the absence of any co-localisation of AMOT and CTNNB1 on marked outer-cell lateral membranes in either embryo group. **G)** Comparison of whole embryo expression levels of AMOT and CTNNB1, derived from the same embryos groups illustrated in panels E and F. **H)** Similar to panel C), quantification of single-outer-cell expression levels (RFU) for apical membrane-associated AMOT and lateral membrane-localised CTNNB1 in siNTC and si*Tead4* blastocysts (see Fig. S7 for similar ICM expression data). **I)** PCA plot similar to panel D), reflecting co-localisation and expression levels of AMOT and CTNNB1 at outer-cell apical and lateral membranes (expanded in Fig. S7) – outer marked si*Tead4* KD clone, apical domain analyses, highlighted in yellow. *All quantified protein expression values (RFU) data are summarised in S.Tabs. 6a & 6b and 7a & 7b. Statistical analyses were performed based on determined data distribution (normal or non-normal) and application of appropriate tests; details in supplementary statistics Excel workbook*.

Alongside PARD6B, we examined the expression of the polarity-sensitive Hippo-signalling effector YAP1. Unlike both outer siNTC control clones and the unmarked outer clones of si*Tead4* embryos, the marked *Tead4* KD outer clones did not display typical nuclear YAP1 localisation (which normally indicates intact apical-basolateral polarity and suppressed Hippo-signalling^20,21^). Instead, YAP1 was predominantly cytoplasmic, resembling all assayed apolar inner-cell populations (Fig. 3A). This implies that, in addition to reduced polarity, such clones may exhibit some mechanistic impairment in Hippo-pathway suppression. Whole-embryo YAP1 expression was also significantly reduced after clonal si*Tead4* treatment, but specifically within the nuclei of marked outer clones, as cytoplasmic levels were unchanged (Fig. 3B&C; see Fig. S6A for inner-cell data). Indeed, the average nuclear:cytoplasmic YAP1 expression ratio robustly and significantly decreased in such marked outer clones, more closely resembling inner cell values (Fig. S6B). Principal Component Analysis (PCA) of YAP1 expression/localisation further confirmed this shift, as marked si*Tead4* outer-clones moved away from the group of control outer-clones and towards the tightly clustered inner-cell profiles (Fig. 3D; expanded in Fig. S6D-F). These findings demonstrate that *Tead4* KD causes a cell-autonomous reduction in nuclear YAP1 in outer-cells, supporting the existence of feedback mechanisms that sustain polarity-dependent suppression of Hippo-signalling and TE differentiation. The altered YAP1 localisation closely mimics inner cell populations, suggesting that *Tead4* KD in clones undergoing outer cell-to-ICM reallocation may lead to aberrant Hippo-pathway activation.

To investigate whether *Tead4* KD induces ectopic outer cell activation of the Hippo-pathway, we performed IF-staining of clonal siNTC and si*Tead4*-injected E3.5 blastocysts for AMOT and the basolaterally localised adherens cell junction (AJ) marker CTNNB1 (β-Catenin^41^). From the 16-cell morula stage, AMOT activates Hippo-signalling by localising to inner cell AJs and promoting LATS1/2-dependent phosphorylation of YAP1, leading to its cytoplasmic retention; an association blocked in outer cells due to AMOT sequestration at the contactless and polarised apical domain^20,21^. Hence, we assayed for AMOT mislocalisation to the lateral domains of outer-cells, marked by CTNNB1 expression, which would indicate aberrant Hippo-pathway activation (Fig. 3E). In control siNTC blastocysts, AMOT was normally confined to the apical domain of outer cells, while CTNNB1 was appropriately basolateral and inner cells showed co-localisation consistent with active Hippo-signalling. In si*Tead4* embryos, despite the presence of marked outer clone-specific apical blebs (marking ICM reallocation), no evidence of abnormal AMOT or CTNNB1 lateral domain co-localisation was observed in either marked or unmarked clones (note, outer cell basal membrane localisation cannot be unambiguously assigned due to neighbouring inner cell plasma-membrane associated signal). Indeed, AMOT was correctly localised to outer cell apical membranes (as was CTNNB1 to all cell-contacted membranes) and the plasma membranes of inner cells. However, AMOT expression levels in general appeared lower than in siNTC controls (Fig. 3E&F). Indeed, whole-embryo quantification analyses confirmed robustly reduced AMOT and modestly reduced CTNNB1 in si*Tead4* blastocysts (Fig. 3G) and on the single-cell level, *Tead4* KD caused significant reductions in apical AMOT in both marked and unmarked outer clones, with no significant changes in lateral CTNNB1 expression (Fig. 3H). Similar clone-independent reductions were also seen in inner cell clones, though CTNNB1 expression was unaffected (Fig. S7A). These analyses demonstrate *Tead4* KD decreases AMOT expression globally, likely through secondary, non-cell-autonomous mechanisms, but has no effect on its sub-cellular localisation (and CTNNB1 expression). Further PCA plots, describing AMOT and CTNNB1 expression levels at outer cell apical and lateral membranes, showed both control siNTC and si*Tead4* KD marked and unmarked clones co-segregated into two main clusters: one with high CTNNB1 and low AMOT levels at lateral membranes (indicating a lack of any *Tead4* KD-induced atypical co-localisation), and a more dispersed apical membrane cluster, associated with reduced and clone-independent AMOT levels in si*Tead4* treated blastocysts; note, tight association of mean-centroids versus equivalent siNTC clones (Fig. 3I; expanded in Fig. S7B-D). Such a membrane-of-origin PCA separation indicates minimal differences in outer-cell AMOT and CTNNB1 (co-) localisation, associated with clonal *Tead4* KD.

Overall, the outer cell-to-ICM reallocations caused by clonal *Tead4* KD are linked to significantly reduced apical polarity and lower clone-independent levels of AMOT across the early-blastocyst (E3.5). Despite these changes, there is no evidence of AMOT mislocalisation to the lateral membranes of outer cells. Therefore, although marked outer si*Tead4* clones display atypical cytoplasmic YAP1 expression, it is highly unlikely that this results from the aberrant activation of the classical-recognised Hippo-signalling pathway^20,21^. This strongly suggests that Hippo-pathway activity is not the primary driver of the observed allocation phenotypes. Instead, TEAD4 is more likely involved in maintaining the proper spatial positioning of TE cells through a broader regulatory mechanism at the transcriptomic level; although this can positively influence apical polarity and appropriate Hippo-signalling feedback, particularly at the level of YAP1 and AMOT protein expression.

### Transcriptomic analysis of internalising *Tead4* KD clones identifies dysregulated cytoskeletal gene expression in early- (E3.5) stage blastocysts

To gain mechanistic insight into outer-cell apical domain homeostasis and *Tead4*-regulated blastomere positioning, we performed RNA-Seq transcriptomic profiling of outer *Tead4* KD clones. Single 2-cell-stage blastomeres were co-injected with dsRNA targeting *Tead4* (ds*Tead4*) and fluorescent rhodamine-conjugated dextran beads (RDBs - as an injection/lineage marker), and embryos were cultured to the 32-cell (E3.5) stage; concurrent with apical bleb formation and the onset of the observed atypical cellular internalisations. We opted for a ds*Tead4*-rather than si*Tead4*-mediated approach to utilise the non-genetically encoded RDB injection marker for the transcriptome assays (instead of H2B-RFP mRNA). Based on our experience, *Tead4* KD efficiency was superior in this context when using the previously reported ds*Tead4* construct^26^. After removal of the *zona pellucida*, blastocysts were briefly cultured with yellow-green-fluorescent microspheres (YGMs) that are readily endocytosed by outer cells. Following embryo disaggregation into single cells, four populations were identifiable: (i) inner cells unmarked by RDBs or YGMs (derived from the non-microinjected 2-cell-stage blastomere); (ii) inner cells marked by RDBs alone (from ds*Tead4*-microinjected clones); (iii) outer cells marked by YGMs alone (from unmarked outer populations); and (iv) outer cells marked by both RDBs and YGMs (from *Tead4* KD outer clones) (Fig. 4A). Therefore, we analysed populations (iii) and (iv) by RNA-Seq to compare outer-cell transcriptome composition between *Tead4* KD and control unmarked clones from the same blastocysts.

**Figure 4:**
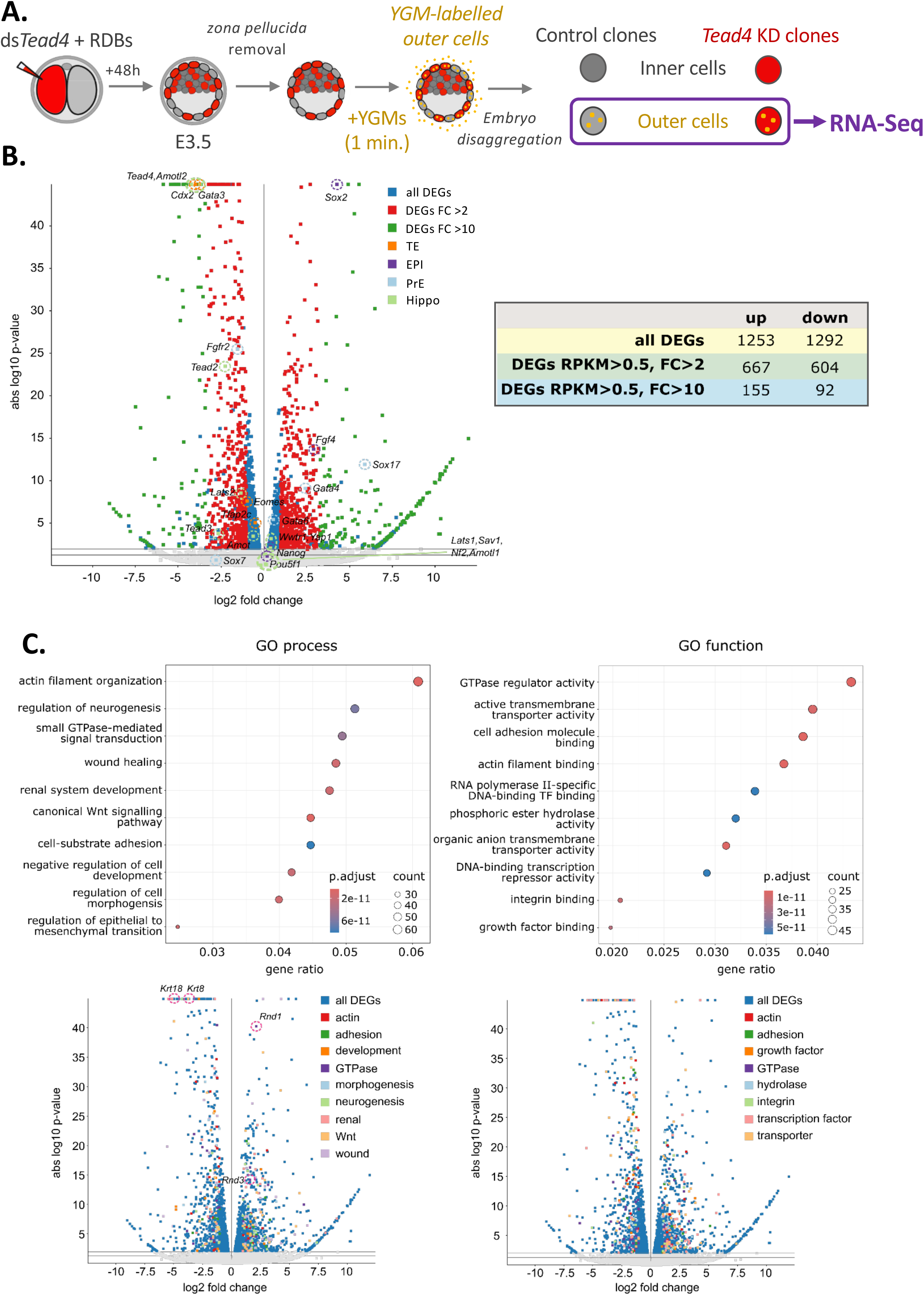
RNA-Seq; *Tead4* KD-mediated dysregulation of cytoskeleton-, actin- and cell adhesion-related gene expression. **A)** Schematic of the dsRNA microinjection protocol used to generate E3.5 blastocysts containing *Tead4* KD (ds*Tead4*) clones, marked by co-injection with rhodamine-conjugated dextran beads (RDBs), representing 50% of all cells. At E3.5, outer cells were labelled via endocytosis of yellow-green microspheres (YGMs), followed by full embryo dissociation into single cells, to enable the identification of inner and outer cell populations from marked and unmarked clones, and preparation of RNA-Seq libraries to compare outer cell transcriptomes. **B)** Left: Volcano plot showing gene expression differences between RNA-Seq transcriptomes of E3.5 *Tead4* KD (marked) and control (unmarked) outer clone cell populations. DEGs are shown in blue, but those with >2-fold and >10-fold changes are highlighted in red and green, respectively. Key marker genes for TE, EPI, and PrE lineages, as well as Hippo-pathway components, are annotated. Right: Summary of up- and downregulated DEGs, plus those with >2-fold and >10-fold expression changes, each with RPKM >0.5 in at least one clone. **C)** Upper panels: Dot plots showing the top ten enriched GO terms, classified by process (left) and function (right) for DEGs (>2-fold change; RPKM >0.5 in at least one clone) between *Tead4* KD and control outer cells. Lower panels: Volcano plots displaying DEG distributions by process-(left; highlighting candidate genes for functional analysis in Fig. 5; *Krt8*, *Krt18*, *Rnd1* & *Rnd3*) and function- (right) related terms. The simplified GO category legends include specific/merged terms: · Process: actin (actin filament organisation), adhesion (cell-substrate adhesion), development (negative regulation of cell development), GTPase (small GTPase-mediated signal transduction, morphogenesis (regulation of cell morphogenesis), neurogenesis (regulation of neurogenesis), renal (renal system development), Wnt (canonical Wnt signalling pathway & regulation of epithelial to mesenchymal transition) and wound (wound healing). · Function: actin (actin filament binding), adhesion (cell adhesion molecule binding), growth factor (growth factor binding), GTPase (GTPase regulator activity), integrin (integrin binding), transcription factor (RNA polymerase II-specific DNA-binding transcription factor binding & DNA-binding transcription factor repressor activity) and transporter (active transmembrane transporter activity & organic anion transmembrane transporter activity). *Lists of all significantly identified DEGs (supplementary data Excel workbook 1), and their association with significantly enriched function- and process-related GO-terms (delineated as those showing >2 fold down- and upregulation or only down- or upregulation alone) are summarised in S.Tabs. 10-15. Comparisons with enriched DEGs identified in E3.25 stage Tead4 genetic knockouts*^49^ *and genes associated with TEAD4 binding in ChIP-Seq analysis of mouse TSCs*^50^ *are provided (S.Tabs. 8 & 9, plus supplementary data Excel workbooks 2 & 3, respectively)*.

Applying a minimal expression threshold (RPKM >0.5 in either derived dataset) and excluding genes located on sex-linked chromosomes, *Tead4* KD triggered extensive transcriptional changes (Fig. 4B). Specifically, 667 mRNAs were significantly upregulated >2-fold (155 >10-fold), while 604 transcripts were downregulated >2-fold (92 >10-fold). A full list of differentially expressed genes (DEGs) is available in the supplementary data Excel workbook 1 (p<0.05). *Tead4* KD caused a marked reduction in *Tead4* transcript levels, with no evidence of compensatory upregulation among other TEAD-family transcription factors; indeed, both *Tead2* and *Tead3* were significantly reduced. Correspondingly, other TE marker mRNAs such as *Cdx2*^32^ and *Gata3*^30^ were downregulated, although *Tfap2c*^31^ showed a significant but less than <2-fold decrease. In contrast, some ICM markers, including early pluripotency genes *Sox2*^22^ and *Fgf4*^42,43^ (but not *Nanog*^7^), were upregulated. The early primitive endoderm (PrE) marker *Fgfr2* was downregulated, consistent with known inner- versus outer-cell expression patterns established at the 16-cell stage^42,43^. Unexpectedly, other PrE genes such as *Gata6*^7^, *Sox17*^44^, and *Gata4*^45,46^ were induced. Genes involved in the Hippo-signalling pathway^20^ (including *Amot*, *Amotl2*, and *Lats1*) were significantly downregulated, consistent with feedback regulation of AMOT protein expression (Figs. 3G&H and S7A). Paradoxically, *Yap1* transcripts were significantly upregulated, despite reduced nuclear YAP1 protein levels in marked *Tead4* KD outer cells (Figs. 3B&C and S6B&C), indicating potential post-transcriptional regulation. Similarly, transcripts for another TEAD4 co-factor *Wwtr1*^47^ (*Taz*) were upregulated, possibly reflecting compensatory responses to targeted *Tead4* mRNA loss. Intriguingly, and relating to the observed *Tead4* KD-induced apical domain distortions/blebs (Figs. 1 & 2 and S1, S4 & S5), we observed that amongst the most significantly reduced DEGs were the cytokeratin genes, *Krt8* and *Krt18* genes (highlighted in Fig. 4C); components of the intermediate filament cytoskeleton governing cell structure and shape^48^. Overall, these findings reveal substantial transcriptomic shifts in the outer cells of 32-cell (E3.5) stage blastocysts following *Tead4* KD. We further compared the DEGs with those from a recent RNA-Seq study of wild-type versus *Tead4* CRISPR-Cas9 knockout embryos at the earlier E3.25 stage^49^. Of the 310 downregulated and 360 upregulated genes reported in that study, 125 (including *Krt8*) and 105 respectively overlapped with our dataset (S.Tab. 8). Additionally, we cross-referenced our DEGs with known TEAD4-genome interactions in mouse trophoblast stem cells (TSCs), identified via chromatin-immunoprecipitation/ChIP-Seq^50^. We found that 110 of the 604 downregulated DEGs and 78 of the 667 upregulated DEGs were associated with TEAD4 binding, with 14 and 8 respectively located in promoter regions (S.Tab. 9 and supplementary data Excel workbooks 2 and 3). While these overlaps support the identification of direct TEAD4 target genes, potentially active from the 16-cell stage onwards, the majority of transcriptional changes observed following *Tead4* KD in E3.5 blastocyst outer cells are likely driven by secondary regulatory mechanisms

We next analysed all DEGs showing >2-fold change (both upregulated and downregulated) for enrichment in gene ontology (GO) terms, using process- and function-related classifications (Fig. 4C and S.Tabs. 10 & 11). Consistent with the dynamic cell behaviours observed during live-embryo (Figs. 2 and S4 & S5), the most significantly enriched GO categories were associated with actin, small GTPase activity (including association with known actin regulator genes such as *Rnd1* and *Rnd3* - atypical, constitutively active small GTPases^51^ that negatively regulate cell adhesion and actomyosin stress-fibre formation^52–55^, that were transcriptionally induced after *Tead4* KD), cell adhesion, and morphogenesis. After narrowing the GO analyses to either upregulated or downregulated DEGs (Figs. S8 & S9 and S.Tabs. 12 & 13 for downregulated; Figs. S10 & S11 and S.Tabs. 14 & 15 for upregulated), we found that these terms were predominantly enriched among the downregulated genes. This was particularly evident for actin-related terms (e.g., ‘actin filament binding’, ‘actin filament organisation’, ‘regulation of actin filament-based process’, and ‘microfilament motor activity’) and GTPase-related labels (e.g., ‘GTPase regulator activity’, ‘GTPase binding’, and ‘small GTPase-mediated signal transduction’). Cell adhesion-related terms (e.g., ‘cell adhesion molecule binding’) were enriched in both up- and downregulated gene datasets. We also identified significant GO terms linked to epithelial development. For instance, ‘cell-cell junction organisation’ and ‘epithelial cell development’ labels were enriched in downregulated DEGs, while ‘regulation of epithelial to mesenchymal transition’ was associated with upregulated DEGs.

These findings are consistent with our observations of apical bleb formation, apical domain abscission, and apparent increased intrinsic contractility driving outer cell-to-ICM reallocation in both fixed (Figs. 1 & 3 and S1) and live-imaged embryos (Figs. 2 and S4 & S5). They also suggest that *Tead4* KD induces substantial transcriptomic changes that remodel the cytoskeleton (particularly the actin network, but also intermediate filament formation) and alter the adhesive properties of early- (E3.5) blastocysts stage outer cells. Thus, facilitating their atypical internalisation and implicating *Tead4* as a key homeostatic regulator of spatial positioning under normal conditions, ensuring specified TE cells remain correctly positioned on the embryo surface.

### Clonal *Tead4*-KD blocks outer-cell intermediate filament formation, yet co-expression of recombinant *Krt8* & *Krt18* mRNAs fails to rescue blastocyst outer cell-to-ICM allocations

Cytoskeletal intermediate filaments, predominantly comprising of KRT18 (type-I) and KRT8 (type-II)^56^, show marked inter-blastomere variability in expression at their late 8-cell onset^57,58^ and are subsequently restricted to TE/outer blastocyst cells^59^, where their asymmetric inheritance is proposed to support TE specification by stabilising the polarised apical domain and repressing Hippo-signalling^23^. Our RNA-Seq revealed robust *Krt8*/*Krt18* suppression (Fig. 4C), and we confirmed the absence of endogenous KRT8-containing intermediate filaments in marked outer cells of *siTead4*-treated E3.5 embryos (Fig. 5A–C). However, although outer cell-to-ICM allocation is enhanced in marked *siTead4* KD clones at E3.5 (Figs. 1 and S1), siRNA targeted knockdown of *Krt8* alone, or together with *Krt18*, did not elicit any similar allocation phenotypes (Fig. S12). This result indicates that loss of cytokeratin intermediate filament cytoskeletal networks alone is unlikely to be the main driver of the observed si*Tead4* KD-mediated allocations. Therefore, to test whether experimentally inducing replacement cytokeratin expression could counteract *Tead4* KD, we co-microinjected single 2-cell stage blastomeres with recombinant KRT8 and KRT18 encoding mRNAs (including HA-tagged KRT8 variants) and confirmed precocious and marked clone-specific intermediate filament formation in early 8-cell embryos (Fig. S13A–D). Repeating clonal *Tead4* KD (plus siNTC controls) with co-injected recombinant *Krt8/Krt18* mRNAs showed marked *Tead4* KD clones still contributed more cells to the E3.5 stage ICM (and fewer cells to outer positions) than controls, despite recombinant intermediate filament expression (Figs. 5D–F and S13E&F). This demonstrates that induced recombinant intermediate filament network replacement cannot rescue outer cell-to-ICM allocations caused by *Tead4* KD. Indeed, apical domain distortions/blebs persisted in such marked si*Tead4*-containing outer clones, and recombinant KRT8-containing intermediate filament networks were unusually observed in marked inner cells of both treatment conditions (Figs. 5E and S13F).

**Figure 5.**
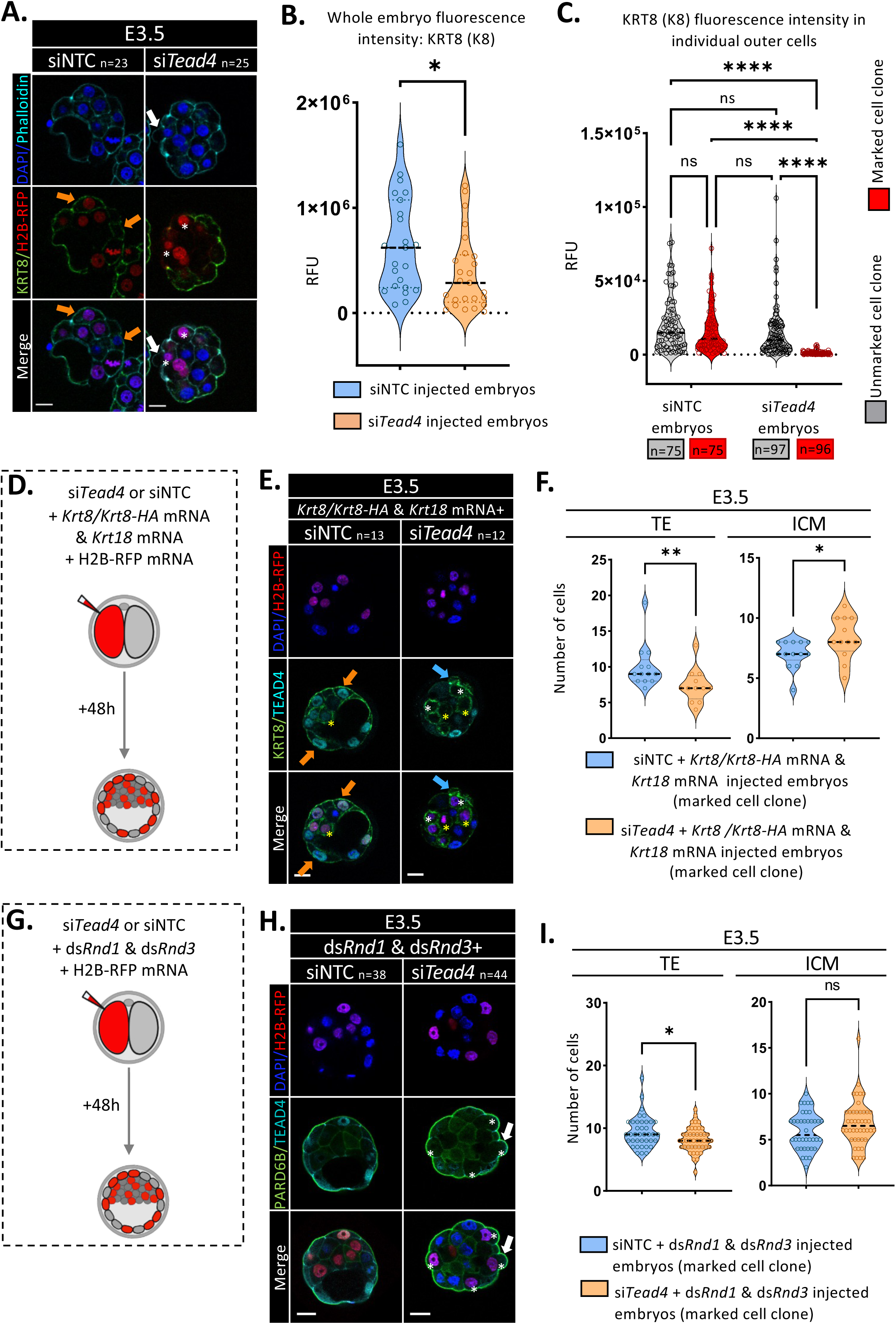
Clonal *Tead4* KD and attempted phenotypic rescue by *Krt8*/*Krt18* overexpression or *Rnd1*/*Rnd3* KD. **A)** Example single-z-section confocal images of E3.5 blastocysts containing H2B-RFP marked clones (red) generated by 2-cell microinjections with either siNTC (left) or si*Tead4* (right). Embryos are IF-stained for KRT8 (green; intermediate filaments) and counterstained with phalloidin (cyan; actin) and DAPI (blue; DNA). Scale bar, 20μm. The number of embryos per group (n) is indicated. Orange arrows denote typical outer-cell KRT8 expression in siNTC embryos, irrespective of clonal origin. White asterisks indicate the absence of detectable KRT8 in outer si*Tead4* clones, which can show abscising apical blebs (white arrows). **B)** KRT8 expression across whole embryos at E3.5 (RFU, normalised), including unmarked and marked clones, in siNTC control (blue) versus *Tead4* KD (orange, si*Tead4*) groups. Median values are shown by dashed lines. Significant differences: *p<0.05. **C)** KRT8 expression in single outer cells (RFU) of siNTC control and si*Tead4* blastocysts; data separated for unmarked (grey) and marked (red) clones. Medians (dashed lines) and significant differences (*p<0.05) are indicated. Sample sizes (n) are shown. **D)** Schematic of the microinjection procedure to generate E3.5 early-blastocysts with control (siNTC) or *Tead4* KD (si*Tead4*) marked clones, co-expressing recombinant *Krt8* (with or without an N-terminal HA tag) and *Krt18* mRNA (50% of cells) to test for phenotypic rescue of *Tead4* KD-induced clonal outer/TE-to-ICM cell allocation phenotypes. **E)** As in A) but for rescue experiments described in D); phalloidin actin staining is replaced by TEAD4 IF (cyan) to confirm specific KD (white asterisks). Yellow asterisks indicate enhanced recombinant KRT8 expression and intermediate filament formation in marked inner-cell clones (of both treatment groups, atypical of endogenous expression). Blue arrows highlight abscised apical bleb-derived surface vesicles from outer *Tead4* KD clones (recombinant KRT8 expression data expanded in Fig. S13). **F)** Outer/TE and ICM cell-number contributions of marked siNTC versus si*Tead4* clones expressing recombinant *Krt8* and *Krt18* mRNAs. Medians (dashed lines); significant differences: *p<0.05. Note that *Krt8*/*Krt18* overexpression did not rescue *Tead4* KD-induced outer/TE-to-ICM cell allocations (expanded in Fig. S13E–F). **G)** Schematic of parallel microinjections to generate E3.5 early-blastocysts with control (siNTC) or *Tead4* KD (si*Tead4*) marked clones (50% of cells), co-injecting either control dsEGFP or ds*Rnd1* plus ds*Rnd3* constructs to attempt phenotypic rescue. **H)** As in E) but for the rescue experiments described in G); TEAD4 IF (cyan) replaces KRT8 staining. White asterisks denote outer-marked si*Tead4* clones, some with disrupted apical domains (white arrows). **I)** Outer/TE and ICM cell-number contributions for marked siNTC/dsEGFP (blue) versus si*Tead4*/ds*Rnd1* & ds*Rnd3* (orange) clones. Medians (dashed lines); significant differences: *p<0.05. Previously significant *Tead4* KD–induced increases in ICM contribution are rescued by concomitant *Rnd1*/*Rnd3* KD, but apical-domain distortions persist, indicating partial phenotypic rescue (expanded in Fig. S14A). *Quantified KRT8 expression values (RFU) for panels B) and C) are summarised in S.Tabs. 17; raw clonal-allocation data are in S.Tabs. 19 (Krt8 & Krt18 over-expression; as in panel F) and S.Tabs. 20 (Rnd1 & Rnd3 KD; as in panel I). Statistical analyses were performed based on determined data distribution (normal or non-normal) and application of appropriate tests; details in supplementary statistics Excel workbook*.

Collectively, the data indicate TE-specific cytokeratin intermediate filaments arise as a consequence of the TEAD4-regulome, potentially via direct transcriptional regulation, but their establishment or maintenance is not essential for correct apical domain morphology or preventing outer cell-to-ICM misallocation in the blastocyst, at least under the context of clonal *Tead4* KD. Consequently, TEAD4-mediated control of TE spatial allocation likely operates through additional mechanisms, beyond KRT8 and KRT18 and intermediate filament formation.

### Dual *Rnd1* and *Rnd3* KD partially rescues clonal *Tead4* KD–induced blastocyst outer cell-to-ICM allocation

Our RNA-Seq analysis showed clonal *Tead4* KD in outer E3.5 blastocyst cells drives robust gene expression changes enriched for GTPase- and actin cytoskeleton-related ontologies (Figs. 4C and S8-11; plus S.Tabs. 10-15). Among these were the *Rnd1* and *Rnd3* genes. As atypical small GTPases that bind GTP but lack intrinsic GTPase activity^51^, *Rnd1* and *Rnd3* can act as negative regulators of actomyosin stress-fibre formation and cell adhesion by inhibiting RHOA and downstream signalling (e.g. ROCK1/2)^52–55^, and in the case of human RND3 are associated with expanding plasma-membrane blebs^52^, reminiscent of marked outer si*Tead4* blastocyst clones (Figs. 1-3 & 5 and S1 & S13). We hypothesised that the outer cell-to-ICM reallocations observed after clonal *Tead4* KD (Figs. 1 & 2 and S1) would be sensitive to *Rnd1*/*Rnd3* expression levels. Therefore, as both the functionally related *Rnd1* and *Rnd3* transcripts were robustly induced in the outer si*Tead4* clone-derived RNA-Seq data (Fig. 4C and supplementary data Excel workbook 1), and were also elevated after global *Tead4* KD by the E3.5 stage (see RT-qPCR data; Fig. S14B–D – note, these data are derived from global rather than clonal *Tead4* KD, as derived in the RNA-Seq data), we targeted both for dsRNA microinjection-mediated KD (note, co-microinjected ds*Rnd1*/ds*Rnd3* global KD efficacy, versus dsEGFP negative control, was validated at E3.5 by RT-qPCR; Fig. S14E-G). We found that in the context of global si*Tead4*-mediated KD, ds*Rnd1*/ds*Rnd3* co-microinjection reduced the enhanced expression of *Rnd1* and *Rnd3* transcripts to levels significantly lower or equivalent to those observed in control siNTC/dsEGFP-treated E3.5 blastocysts (Fig. S14H–J). Hence, we assayed the effect of this reduction on the relative-spatial allocation of si*Tead4* marked clones (again, versus equivalent siNTC/dsEGFP microinjection controls; Fig. 5G&I). We found E3.5 blastocysts containing marked si*Tead4* clones co-injected with ds*Rnd1*/ds*Rnd3,* exhibited a rescue of ICM cell contribution to levels comparable to control clones; reversing the enhanced ICM allocation associated with *Tead4* KD alone (Figs 1 and S1). However, the marked clone TE contribution was significantly reduced, suggesting only a partial rescue, as supported by the presence of apical distortions/blebs in some marked outer clones (Fig. 5H). The direct unmarked versus si*Tead4*/ds*Rnd1*/ds*Rnd3* marked clone comparisons, within the same embryos, showed the same trend (Fig. S14A; i.e. were comparable to the equivalent and unbiased ICM contribution in siNTC/dsEGFP-treated controls).

Overall, these data indicate that clonal *Tead4* KD disrupts the expression of the atypical *Rnd1* and *Rnd3* small GTPases, which have established links to actomyosin stress-fibre formation and plasma membrane deformation^51–55^. Moreover, this contributes to abnormal blastocyst outer cell-to-ICM allocations. They also imply that *Tead4*-driven regulation of actomyosin contractility plays a role in the relevant spatial positioning of specifying TE cells (consistent with the dynamic cell behaviours observed in 4D live-embryo imaging; Figs. 2 and S4 & S5). Consistently, we discovered that treating late morulae/early blastocysts (E3.25) for 12 hours (i.e. spanning the period during which clonal *Tead4* KD is associated with the greatest extent of atypical outer cell-to-ICM cell allocations; Figs. 1 and S1) with Jasplakinolide, a potent inducer and stabiliser of actin filaments^60^, resulted in the formation of outer-cell polarised apical-domain distortion/blebs (containing PARD6B) and a failure to accumulate nuclear YAP1, with reduced levels equal to those observed in outer-residing marked *Tead4* KD clones at the equivalent developmental stage (Fig. S15). These phenotypes are highly reminiscent of those elicited by clonal *Tead4* KD and support the existence of a broad link between actomyosin organisation, apical-domain homeostasis and the correct spatial positioning of TE-specified cells under the regulation of *Tead4*; although the precise role of *Rnd1* and *Rnd3* in regulating apical domain morphology, within the spectrum of other *Tead4*-regulated genes, as opposed to directly contributing to regulating spatial positioning (Figs. 5G-I and S14), is less clear.

### Conclusions

We show that *Tead4* plays a critical, cell-autonomous role in maintaining the correct outer spatial positioning of TE-specified cells in the developing mouse blastocyst, extending beyond its overlapping functions with *Tfap2c* in establishing apical-basolateral polarity^15^ and its involvement in regulated Hippo-signalling and TE cell fate specification^20,21^. Clonal KD of *Tead4* disrupts apical-domain morphology, causing apical distortions that are associated with abnormal cell divisions and apical bleb abscissions; these defects, together with classical asymmetric cell division and apical domain constriction, result in the ectopic formation of blastocyst ICM cells. Although apical polarity is reduced, overall apical-basolateral polarity remains intact, and lateral AMOT mislocalisation is not observed, suggesting that classical Hippo-signalling activation^20,21^ is not the principal driver of such outer cell-to-ICM transitions, despite reductions in AMOT and nuclear YAP1 levels. RNA sequencing reveals that *Tead4* broadly regulates cytoskeletal processes, particularly actomyosin-related factors such as *Rnd1* and *Rnd3*, and cell adhesion, indicating that *Tead4* ensures proper TE cell positioning by maintaining polarised apical-domain homeostasis and appropriate actomyosin dynamics.

## DISCUSSION

The first cell fate decision in mouse preimplantation development involves the spatial segregation of outer TE progenitors from the ICM by the early blastocyst stage (E3.5), typically associated with the 8- to 16- and 16-to 32-cell transitions, though infrequently observed later^34^. Our 4D live-embryo imaging confirms rare internalisations during blastocyst maturation in control siNTC-injected clones via classical mechanisms such as asymmetric cell division or apical domain constriction. In contrast, *Tead4* KD clones show a significantly and atypically increased frequency of such outer-to-inner cell internalisations, peaking post-32-cell stage (E3.5-E3.75) and continuing throughout the whole of blastocyst maturation, that are associated with additional mechanisms including apical blebbing, apical domain abscission, and anomalous type-I or type-II divisions (Figs. 1 & 2 and S1, S4 & S5). Our findings reveal an expanded role for *Tead4* in maintaining cell fate-appropriate spatial positioning and regulating apical domain morphology to ensure that TE cells remain in an outer position. Prior to the 32-cell (E3.5) blastocyst stage, cell internalisations coincide with Hippo-dependent upregulation of pluripotency genes like *Sox2* and downregulation of TE markers such as *Cdx2*^20–22^. Although derived ICM cells retain plasticity, TE fate becomes fixed around the late E3.5 blastocyst stage, concomitant with the peak of atypical internalisations we observe in *Tead4* KD clones. Chimaera studies show that individual 32-cell stage blastomeres with low CDX2-EGFP reporter expression almost exclusively contribute to the ICM^61^. Hence, since *Cdx2* is a transcriptional target gene for TEAD4-mediated activation^24,25^, this supports a model in which TEAD4 ordinarily governs TE versus ICM cell spatial segregation at this stage. Therefore, *Tead4* KD appears to disrupt this process, leading to increased and, importantly, prolonged classical internalisation events, and those involving apical blebbing, abscissions and abnormal divisions, throughout blastocyst maturation. Clonal disruption of other TE-related/TEAD4 target gene expression, such as *Tfap2c*^31^ or *Cdx2*^24,25^, does not replicate this phenotype (Figs. S2 & S3). Such data indicate that spatial regulation of TE cells is *Tead4*-specific, and whilst low CDX2 expression levels may be a good marker of eventual TE versus ICM cell fate, it also suggests an alternative mechanism. Additionally, whilst *Tfap2c* is reported to have overlapping roles with *Tead4* in the 8-cell stage establishment of apical polarisation, in conjunction with RHOA signalling^15^, marked *Tead4* KD clones retain intact apical polarity (evidenced by PARD6B localisation), despite reduced PARD6B expression (Figs 1C & 3A-C and S1B; note, transcript levels were also significantly reduced after clonal *Tead4* KD in the outer cell clonal RNA-Seq data - supplementary data Excel workbook 1); indicating initiation of polarity can form independently of *Tead4*. Although *Tead4* KD significantly reduces *Tfap2c* mRNA levels in early (E3.5) blastocysts (Fig. S1D-F), the degree of functional redundancy with *Tead4* in regulating homeostatic spatial allocation during TE-maintenance remains uncertain, especially as clonal *Tfap2c* KD alone does not produce comparable atypical blastocyst cell internalisations (Fig. S2). Therefore, although reports indicate PARD6B levels are positively regulated by TFAP2C itself^31,40^, potentially explaining the clonal *Tead4* KD-associated reductions we present here, these reductions alone would be insufficient to explain the overall observed allocation phenotypes.

*Tead4* plays a well-established role in specifying the outer TE cell lineage through polarity-dependent regulation of Hippo-signalling and localisation of its transcriptional co-factor YAP1, in both mice^20–22,24,25,61^ and other species such as cattle^49^. The acquisition and maintenance of polarity are closely linked to the spatial positioning of outer TE progenitors and nascent ICM cells by the 32-cell stage, with an absence or loss of polarity triggering internalisation of outer cells via actomyosin-driven contractility^10,11,17–19,35–38^; supporting the “polarity-dependent cell-positioning model”^27^. Our findings both align with and have clear distinctions from such previous reports. For example, while *Tead4* KD clones retain apically restricted PARD6B expression (likely established at the 8-cell stage via *Tfap2c* and RHOA activity^15^) they nevertheless show significantly reduced PARD6B expression by the 32-cell stage (Fig. 3A-C), often accompanied by apical bleb formation (Figs. 1–3 & 5 and S1, S4, S5, S13 & S14). Such data suggest that *Tead4* KD clones may fail to maintain polarity at sufficient levels to preserve outer positioning, leading to increased internalisation via classical mechanisms such as asymmetric division or actomyosin-driven apical constriction, as reported for completely apolar outer cells in the 16-cell stage morula^17^. Supporting this, our 4D live-embryo imaging reveals elevated frequencies of these internalisation events, plus apical domain abscission events, in *Tead4* KD clones compared to controls before the 32-cell (E3.5) stage (Figs. 2 and S4 & S5). Additionally, recent findings show that inter-blastomere differences in the timing of apical-basolateral polarity initiation at the 8-cell stage influence subsequent asymmetric divisions and spatial cell fate allocation, with earlier polarising cells biased towards forming outer TE cells at the blastocyst stage (and *vice-versa*)^39^. Thus, it is possible that *Tead4*-regulated feedback mechanisms help maintain polarity thresholds that prevent internalisation of TE-fated cells, either by limiting asymmetric divisions^39^ or suppressing apical constriction^10,11,17^. Indeed, those early- (E3.5) blastocyst cells with low CDX2-EGFP reporter expression, that is presumably *Tead4*-dependent^24,25^, preferentially contribute to the ICM in morula chimeras^61^. Further evidence for a broader *Tead4*-linked polarity–Hippo-signalling feedback loop comes from observed transcriptional dysregulation of other pathway components (e.g., *Amot*, *Amotl2*, *Lats2*, *Yap1* and *Wwtr1* - Figs. 4A and supplementary data Excel workbook 1). Indeed, we observed a marked reduction in nuclear YAP1 protein expression within outer *Tead4* KD clones (Figs. 3C–D and S6B&C); correlating with not just our clonal RNA-Seq data (Figs. 4 and supplementary data Excel workbook 1), but also similar transcriptomic analyses after whole embryo *Tead4* gene knockout at the earlier E3.25 stage^49^. This failure to accumulate nuclear YAP1 may result from one or more factors: impaired TEAD4-dependent transcriptional feedback, inability to form sequestering TEAD4–YAP1 nuclear complexes, mechanosensory-related mechanisms associated with reduced actomyosin-driven contractility affecting YAP1 localisation^10^, or aberrant Hippo-pathway activation leading to cytoplasmic retention. However, the latter is unlikely, as cytoplasmic YAP1 levels in *Tead4* KD clones match those of control outer cells (Fig. 3C), and no ectopic lateral AMOT localisation, typically linked to Hippo-activation during the preimplantation stage^20,38,62^, was observed in marked outer clones (Figs. 3E–H & I and S7); distinguishing them from phenotypes associated with similar cytoplasmic YAP1 enrichment, without evident reductions in expression, in naturally occurring apolar outer cells^17^, or after experimental disruption of apical-basolateral polarity (e.g. via chemical inhibition of ROCK1/2^37,38^). Notwithstanding, these findings suggest a broader *Tead4*-mediated feedback mechanism that may reinforce polarity-dependent Hippo-signalling, contributing to TE cell fate maintenance and spatial positioning, in conjunction with other invoked mechanisms, during normal development.

Beyond established links with apical-basolateral polarity and cell-positioning prior to the 32-cell (E3.5) stage, our data reveal a distinct and robust increase in atypical internalisation of marked *Tead4* KD clones during blastocyst maturation itself, contributing to an enlarged ICM population (Figs. 1 & 2, and S1, S4 & S5). 4D live-embryo imaging (Figs. 2 and S4 & S5) shows that while internalisation can occur via conventionally recognised asymmetric cell division or apical constriction^11,17,34^, *Tead4* KD clones also exhibit novel mechanisms associated with apical domain distortions and bleb formation: type-I divisions (one daughter inherits the bleb and remains outside, the other internalises), type-II divisions (both daughters internalise, leaving a vesicular bleb remnant on the outside), and bleb abscission events followed by cell internalisation independent of division (also atypically observed, versus controls, prior to the 32-cell stage). Fixed blastocyst analyses show that apical blebs and surface vesicles/bleb remnants contain PARD6B (Figs. 1C, 3A & 5A, E&H and S1B, S13F & S14A), indicating that these atypical events involve resection of polarised apical domains. The persistence and higher frequency of asymmetric cell division and apical domain constriction beyond the 32-cell (E3.5) stage, alongside the emergence of other atypical internalisation events, demonstrates that prolonged *Tead4* KD must disrupt the homeostatic regulation of TE cell positioning, most likely on the transcriptome level. Consistently, RNA-Seq analysis (Figs. 4 and S4 & S5); plus supplementary data Excel workbook 1) identified significant cohorts of *Tead4*-regulated DEGs enriched for cytoskeletal and actin-related functions, including downregulated expression of the cytokeratin genes *Krt8* and *Krt18*. KRT8 and KRT18 form intermediate filaments in 8-cell stage blastomeres, which stabilise the outer cortex, promote apical polarity to support CDX2 expression, and become TE-restricted by the blastocyst stage^23,56^. However, whilst these represent appealing candidates, and *Tead4* KD clones show loss of KRT8-containing intermediate filaments (Fig. 5A–C) and reduced apical PARD6B expression (Fig. 3B & C – indicative of the above mentioned feedback), clonal KD of *Krt8* or *Krt8/Krt18* (Fig. S12) did not lead to increased cell internalisation, nor loss of detectable CDX2 protein expression (possibly reflecting differences in mouse strains between our study and previous reports^23^). Furthermore, reconstitution of recombinant KRT8/KRT18 intermediate filament networks failed to rescue *Tead4* KD-induced cell internalisation phenotypes (Figs. 5D–F and S13), and they were persistent in *Tead4* KD-induced apical blebs and within internalised cell clones, albeit at reduced levels; consistent with their reported asymmetric inheritance from the 8-cell stage (Fig. S13F). Moreover, inner marked siNTC control clones also atypically expressed recombinant KRT8/KRT18 intermediate filaments (Figs. 5E and S13E&F). Together, these results indicate that, whilst these cytokeratin genes are part of the *Tead4* regulome, they are not key regulators of TE spatial cell positioning and expression of KRT8/18 intermediate filaments alone is insufficient to sequester cells to an outer position. However, the RNA-Seq data also revealed strong enrichment of actin-related (and associated GTPase) gene ontologies (Figs. 4C and S8-S11; plus supplementary data Excel workbook 1), implicating actin dynamics in the observed atypical internalisation phenotypes. This is supported by the dynamic cell behaviours seen in 4D live-embryo imaging and the induction of apical blebs containing PARD6B, plus reduced nuclear YAP1 levels, following Jasplakinolide-mediated actin stabilisation (Fig. S15). Together, these findings align well with the established role of *Tead4* (and *Tfap2c*) in the two-step process of apical-basolateral polarity establishment at the late 8-cell stage (E2.5), involving transcriptional activation of actin regulators essential for cortical actomyosin remodelling and polarity factor recruitment^15,16^. Therefore, whilst our findings indicate that *Tead4* alone is not essential for polarity establishment (although there may be some role as part of a reinforcing feedback loop), its downstream transcriptional regulome, enriched in candidate actin-related genes (Figs. 4C and S8-S11), represents a strong source of candidates playing a key role in maintaining lineage-appropriate blastocyst spatial positioning. Indeed, small GTPases represent a major class of dysregulated genes following clonal *Tead4* KD, with *Rnd1* and *Rnd3* notably upregulated in outer *Tead4* KD clones (Fig. 4C and S14B-D). These constitutively active GTPases are known to inhibit actomyosin stress fibre formation and are linked to membrane blebbing^15,51–55^. Consistently, the co-clonal KD of *Rnd1* and *Rnd3*, within the context of *Tead4* KD, is able to rescue enhanced inner cell allocation defects at the 32-cell (E3.5) stage (Figs. 5 G-I and S14A), although apical blebs containing PARD6B persisted, indicating an incomplete phenotypic rescue. These findings suggest that *Tead4* KD-induced upregulation of *Rnd1* and *Rnd3* contributes to the misallocation phenotype by disrupting actomyosin dynamics, but their role is limited within broader transcriptome dysregulation, including the coordinated involvement of other actin-related genes, as well as those associated with other significantly enriched GO-terms. For example, ‘cell adhesion/integrins’ and ‘morphogenesis’ (Fig. 4 and S8-S11). Thus, whilst targeting *Rnd1/Rnd3* alone is insufficient for full *Tead4* KD-mediated phenotypic correction, the data highlight an exemplative role for actin-related genes, under the wider transcriptional control of *Tead4*, in maintaining homeostatic regulation of correct spatial positioning of TE specified cells. Moreover, these functional studies help disentangle apical domain deformation from other *Tead4* KD-associated internalisation mechanisms, such as apical domain constriction and unconventional cell divisions, also active during this developmental window (Figs. 2 and S4 & S5). Lastly, the formation of *Tead4* KD-induced apical blebs containing PARD6B, observed in internalising cells both before the 32-cell (E3.5) stage and throughout blastocyst maturation (see fixed embryo data - Figs. 1, 3 & 5 and S1, S13 & S14, and 4D live-cell embryo data - Figs. 2, and S4 & S5), suggests a conflicted state between increases in actomyosin contractility and the maintenance of apical polarity. As such, internalisation may only occur when contractility surpasses a threshold that any accumulated, or indeed reduced, apical polarity can no longer counteract. Therefore, we suggest that the emergence of the additional unconventional internalisation mechanisms, such as apical domain abscission and atypical type-I and type-II divisions, which become more prominent during the blastocyst maturation period, results from progressive imbalance between polarity and contractility. Hence, under clonal *Tead4* KD conditions, there is a functional uncoupling of the presence of apical polarity from the ability of an outer cell to contribute to the ICM, as would ordinarily ensure maintenance of the appropriate spatial allocation of specified TE-cells.

Our findings broaden the known role of *Tead4* in preimplantation mouse development (summarised in Fig. 6). Beyond initiating apical-basolateral polarity at the 8-cell stage^16^ and promoting outer cell TE fate specification via polarity-dependent Hippo-signalling suppression in the morula and early blastocyst (E3.5)^20,21,61^, we identify *Tead4* as a key regulator of the spatial positioning of TE-specified cells during blastocyst maturation through to the peri-implantation stage (E4.5). Transcriptomic analysis of *Tead4* KD clones highlights a strong contribution from a significant cohort of actin-related genes within the *Tead4* regulome, likely supporting apical domain integrity and preventing atypical internalisation of TE cells; exemplified, in part, by functional studies targeting *Rnd1* and *Rnd3*, although the overall process is most likely highly multi-factorial. Additionally, while feedback on apical polarity appears limited, it may still be functionally relevant, acting to reinforce TE-appropriate gene expression networks contributing to this homeostatic maintenance. Future work should further identify and functionally characterise the key *Tead4*-regulated genes that are collectively responsible for maintaining the appropriately segregated TE lineage, especially during the blastocyst maturation period.

**Figure 6:**
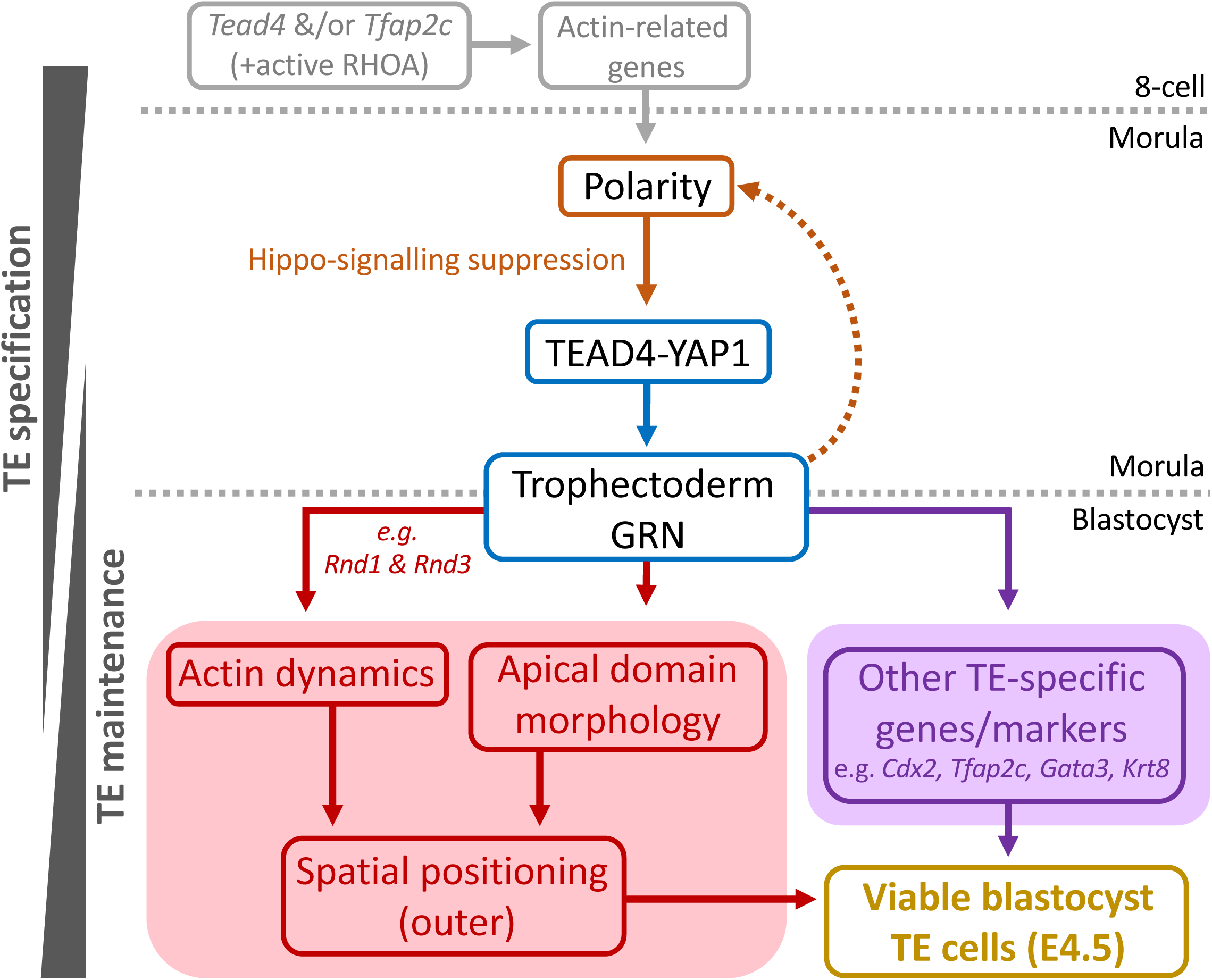
A proposed model of *Tead4*-mediated apical-domain homeostasis and regulation of relative cellular positioning within the emerging mouse blastocyst. At the 8-cell stage, *Tead4* (in partial redundancy with *Tfap2c*) activates genes involved in actin remodelling, and with active RHOA signalling initiates apical-basolateral polarity^15^. Inherited polarity in outer cells at the morula stage contributes to suppression of the Hippo-signalling pathway, leading to the formation of nuclear TEAD4–YAP1 transcriptional complexes^20,21^ that initiate the TE gene regulatory network (GRN) during TE specification; e.g. activating the expression of recognised TE-specific marker genes by the early- (E3.5) blastocyst stage (highlighted in purple). Simultaneously, TEAD4–YAP1 complexes promote appropriate expression of genes that maintain outer-cell apical domain morphology and actin dynamics (e.g. *Rnd1* and *Rnd3*), collectively ensuring that TE-specified cells remain on the surface during blastocyst maturation (highlighted in red), with only limited feedback to apical-basolateral polarity *per se*. Thus, *Tead4* plays a sequential role in TE specification and then TE maintenance during the transition from the morula to the eventual peri-implantation (E4.5) blastocyst stage.

## MATERIALS & METHODS

### Superovulation, embryo isolation & culture (including Jasplakinolide treatment)

All animal work complied with Czech Act No. 246/1992 under the Central Commission for Animal Welfare (approval ID 51/2015). Preimplantation stage embryos were derived as previously described^26^. Eight-week-old F1 hybrid females (C57BL/6 × CBA/W) intraperitoneally received 7.5 IU PMSG, followed 48 hours later by 7.5 IU of hCG, and were mated overnight with F1 stud males. Embryos expressing membrane-associated fluorescent Tomato-reporter were derived in similar crosses using heterozygous F1 mT/mG stud males (Jackson Laboratory, stock 00757631^33^ – with fluorescence-microscopy confirmed mT expression in tail biopsies), although 5 IU of PMSG and hCG were used in the superovulations. From oviducts, superovulated 2-cell-stage embryos (E1.5; 46 hours post-hCG) were micro-dissected and washed through five 20μl drops of prewarmed M2 medium (containing 4mg/ml BSA). Unless proceeding to immediate microinjection (see below), embryos were transferred through fifteen KSOM+AA media drops (Embryo-Max; Millipore) and cultured (at 37°C in 5% CO_2_ in the terminal drop) to the desired developmental stage; note, all 35mm culture plates were pre-equilibrated for 4 hours and were overlaid with 1.5ml of mineral oil (Irivine Scientific). Some embryos cultured to the E3.25 stage (i.e. 42 hours post-2-cell stage recovery) were exposed to similar culture in 15nM Jasplakinolide (Abcam), or an equivalent volume of DMSO solvent vehicle control, containing KSOM+AA drops for 12 hours (i.e. from E3.25-E3.75). For light-sheet live-cell imaging, embryos were isolated in M2 medium and cultured in global® (LGGG-020, Origio) supplied with 1mg/ml BSA in 37°C in 5% CO_2_ and 5% O_2_.

### 2-cell stage RNAi (siRNA & dsRNA) and CRISPR-Cas9 embryo microinjections

To induce gene-specific dysregulated expression, preimplantation mouse embryo microinjections were performed with in-house prepared filamented borosilicate needles (Harvard Apparatus & Narishige) as previously described^26^. Specific details of the injected constructs are in the supplementary information (S.Tab. 22). Briefly, recovered superovulated 2-cell embryos were co-microinjected with indicated combinations of control or gene-specific siRNA, dsRNA, sgRNA, and, as appropriate, recombinant mRNAs (e.g. encoding fluorescent Histone H2B markers, Cas9 endonuclease, KRT8 & KRT18; or non-genetically encoded rhodamine-conjugated dextran beads/RDBs, as an alternative lineage marker for RNA-Seq). To generate clones comprising 50% of the embryo (e.g. cell lineage studies), a single blastomere was injected; to assess specific gene knockdown/KD (knockout/KO) efficacy by RT-qPCR (and assay TE-associated gene expression), both blastomeres were injected (without lineage-marker constructs). Injections were performed in M2+BSA media drops under mineral oil on a 37°C heated stage using an Olympus IX71 microscope, manual Leica micromanipulators, and an Eppendorf FemtoJet injector. Regarding CRISPR-Cas9 targeting of the *Cdx2* gene, minor modifications to a published protocol^63^ were employed. After successful microinjection, embryos were cultured in KSOM+AA drops to the desired stage; non-injected embryos (1–3 per experiment) served as culture control sentinels. For light-sheet live-cell imaging, single blastomeres were microinjected in M2 medium with approximately 10pl of siNTC or si*Tead4* (10µM) and H2B-Venus mRNA (40ng/µl) using a PM2000B microinjector (MicroData Instrument).

### Derivation of microinjected dsRNA and recombinant mRNA constructs

dsRNA constructs targeting murine *Tfap2c*, *Rnd1*, and *Rnd3*, or *EGFP* (enhanced green fluorescent protein) as a negative/non-targeting control (or *Tead4* for RNA-Seq), were designed with the ERNAi design tool^64^ under default settings and produced in-house from T7 promoter-linked PCR products derived from E3.5 blastocyst cDNA templates, using the MEGAscript T7 kit (ThermoFisher). dsRNA integrity was checked by agarose gel electrophoresis; primer sequences are provided in the supplementary information (S.Tab. 22).

Recombinant *Krt8* and *Krt18* (plus HA-epitope tagged *Krt8-HA*) mRNAs were generated via T3 RNA polymerase-mediated IVT from in-house cloned E3.5 blastocyst-derived cDNAs, inserted into *SfiI*-linearised pRN3P^65^ plasmid vectors (which incorporate frog beta-globin UTRs for stability^66^); according to provided IVT and polyadenylation protocols (ThermoFisher; mMESSAGE mMACHINE T3 and poly-A-tailing kit). PCR cloning primer sequences, used to derive cDNA inserts, are provided in S.Tab. 22. mRNAs encoding Histone H2B-RFP/YFP were similarly produced from published *SfiI*-linearised pRN3 templates^67^. For CRISPR-Cas9 targeting of *Cdx2*, recombinant poly-adenylated *Cas9* mRNA (in this case, *Cas9* fused to monomeric streptavidin; *Cas9-mSA*) was synthesised from *NotI*-linearised pCS2+Cas9-mSA plasmid^63^ (Addgene #103882) template, using the mMESSAGE mMACHINE SP6 kit (ThermoFisher). The integrity of all recombinant mRNAs was confirmed by denaturing agarose gel electrophoresis.

### RNA-Seq of marked ds*Tead4* and unmarked E3.5 stage blastocyst outer-cell clones

Per biological replicate, ten 2-cell stage mouse blastomeres were co-microinjected in single blastomeres with *Tead4*-specific dsRNA (ds*Tead4*, prepared in-house) and RDBs, and cultured to the 32-cell stage (E3.5 - apical bleb formation was visually confirmed on a stereo-dissecting microscope). They were then transferred to 20μl drops of prewarmed, BSA-free M2 media containing 1:100 diluted yellow-green fluorescent microspheres (0.20μm YGMs; Fluoresbrite YG, Polysciences), incubated 1 minute at 37°C/5% CO₂, washed through 10 drops of BSA-free M2 media lacking YGMs, and outer-cell endocytic YGM labelling confirmed on an IX71 inverted fluorescence microscope. The *zona pellucidae* were removed in 20μl drops of 37°C Acid Tyrode’s solution (ThermoFisher), and embryos were transferred to prewarmed Ca²⁺/Mg²⁺-free M2 drops and then disaggregated into single blastomeres by repeated pipetting; after which two groups of blastomeres were collected (YGM labelled outer cells from the RDB marked/labelled *Tead4* KD clone, and outer cells only labelled with YGMs representing the unmarked control outer cell clones). The pooled cell populations were placed into 2μl RNA isolation buffer (PicoPure RNA Isolation Kit; Arctus Biosciences), flash-frozen in liquid N_2_, and stored at −80°C (note, the inner-cell populations were not collected). An additional 9 biological replicates were prepared for RDB-marked outer *Tead4* KD and unmarked control outer clones (total n = 10); per replicate, outer-residing *Tead4* KD and control samples yielded pair-wise combinations of, 54, 65, 85, 78, 97, 89, 63, 94, 52, 47 and 89, 76, 118, 114, 77, 98, 91, 124, 75, 75 cells respectively. Next, the marked *Tead4* KD and unmarked outer clone-derived isolation buffer lots were pooled (each 20 μl in total) and RNA purified and eluted in 9.5 μl HPLC-grade water, as per manufacturer’s instructions (PicoPure RNA Isolation Kit) before preparing duplicate RNA-Seq libraries, as previously described^68^ using the NEBNext Ultra II RNA Library Prep Kit from Illumina, using the NEBNext poly(A) mRNA Magnetic Isolation Module, according to manufacturer’s instructions. Libraries were sequenced with 50bp single-end reads on the HiSeq4000 platform, yielding 38 to 57 million reads per library. Data were then trimmed using TrimGalore! v0.6.1 and mapped to the mouse GRCm38 genome using Hisat2 v2.1.0. Gene expression (RPKM) was measured in Seqmonk v1.47.2. Differentially expressed genes (DEGs) were identified using DESeq2 (genes on X and Y chromosomes were excluded and a minimum per gene RPKM expression threshold of >0.5 was applied in relation to gene expression within at least one of the clone datasets). Gene ontology (GO) analysis was performed using the clusterProfile package in R. Datasets are available in GEO database under accession number [*PENDING*]. The list of identified significant DEGs (>2 fold) was compared with similar RNA-Seq-derived DEG lists from the report of Wu et al.^49^, detailing transcript changes in whole embryo *Tead4* genetic knockouts, after CRISPR-Cas9 gene editing, at the E3.25 stage. For DEG comparison with TEAD4 ChIP-seq data, from mouse TSCs derived by Home et al.,^50^ raw data were trimmed using TrimGalore! v0.6.1 and mapped to mouse GRCm38 genome using Bowtie2 v2.3.5. Peaks were then called using MACS2 within Seqmonk v1.47.2. and filtered against identified DEG inputs.

### Immuno-fluorescent (IF) staining and confocal microscopy

Prior to embryo fixation, *zona pellucidae* were removed in acid Tyrode’s solution (Sigma-Aldrich, T1788) followed by sequential M2 washes. Fixation and IF stainings were performed in 96-well plates (Greiner, 655101) at room temperature (unless otherwise stated). Embryos were fixed in 4% paraformaldehyde (ThermoScientific; J61899.AK) for 20 minutes, followed by several brief 0.15% PBST washes and a terminal 20-minute incubation (0.15% Tween-20; Sigma-Aldrich, P1379, in 1X PBS; Gibco, 70011-036). For cell membrane permeabilisation, embryos were transferred to 0.5% TritonX-100 (Sigma-Aldrich, T8787) in 1X PBS for 15 minutes and washed in 0.15% PBST, as above. For non-specific epitope blocking, embryos were incubated in 3% BSA (Sigma-Aldrich, A7906, in 0.15% PBST) for 30 minutes at 4°C. Embryos were then incubated in primary antibodies diluted in BSA blocking buffer for approximately 16 hours at 4°C. Prior to secondary antibody exposure, embryos were washed in 0.15% PBST and transferred to BSA blocking buffer for 30 minutes. Embryos were incubated with fluorescently-conjugated secondary antibodies diluted in BSA blocking buffer for 1 hour at room temperature before final washing in 0.15% PBST. In some cases, phalloidin staining was performed before mounting. Embryos were incubated in phalloidin (50X in 1X PBS) for 30 minutes at room temperature followed by washing in 0.15% PBST for 20 minutes. Details and dilutions of the primary and secondary antibodies used are outlined in the supplements S.Tabs. 23. Embryos were mounted in DAPI-containing media (Vetashield; Vector Laboratories, H-1200, diluted 1:9 in 1X PBS) in glass-bottom 35mm dishes (Mattek, P35G-1.5-14-C) on replacement cover slides (VWR International, 631-0124) overlaid with mineral oil (Sigma-Aldrich, M5904). Complete embryo fluorescent z-section series were obtained by confocal imaging using FV10i confocal laser scanning microscope and FV10i-SW image acquisition software (Olympus), using identical acquisition settings for all control and experimental conditions for a single dataset. All images were captured at 520 x 520 image resolution. For embryonic stages up to and including E3.75, 2μm z-stacks were obtained. For embryos E4.0 and E4.5, 1.5μm z-stacks were obtained.

### Image analyses of fixed samples

Fluoview FV10-ASW 4.2 Viewer software (Olympus) was used to classify inner and outer cell populations in fixed embryo samples. Outer-cell populations were distinguished from inner cell populations by the presence of a contactless apical domain, indicated by PARD6B accumulation. Protein expression levels of defined regions of interest (ROI) were quantified manually in Image J (Fiji^69^) using the Polygon tool or the Line tool, depending on the nature of the analyses. The relative fluorescence units (RFU) were calculated by subtracting the background calculation (from a region outside the imaged embryo) from the total fluorescence (integrated density) of each ROI: RFU = IntDen – (A*X), where IntDen is the integrated density of the ROI, A is the area of the ROI, and X is the mean grey area of the background. Whole embryo quantification was acquired by projecting the stacks using the Stk function and manually tracing around the embryo using the Polygon tool (in the case of KRT8 outer cells quantifications, the central. Apical/lateral domain protein quantifications, such as for PARD6B, AMOT, CTNNB1 and KRT8, were obtained from the centre stack of a given analysed cell (or in the case of KRT8 quantification from the central amd two z-sections above and below) using the Polygon tool to trace around the apical/lateral domains. YAP1 quantification was obtained using the Polygon tool (tracing around the given cell/nucleus) and was calculated by averaging RFU from the central three stacks of a given cell/nucleus. Cytoplasmic YAP1 RFU was quantified by subtracting nuclear YAP1 RFU from the whole cell YAP1 RFU. Principle component analyses (PCA) were performed in R (R 4.4.1, R Core Team^70^, https://www.R-project.org/) using RStudio (Posit team^71^, https://posit.co/), FactoMineR^72^ and factoextra^73^ (https://CRAN.R-project.org/package=factoextra) packages, providing PCA functions and extraction of function data, respectively. The gglot2 function was utilised to create customised graphs, the RColorBrewer was utilised for colour palette visualisation of data points, and the dplyr function for the visualisation of 3D plots. Input data were imported as a CSV files, the categorical variables, outlined in Figs. S6D & S7B, were converted into factors and further grouped into interactive terms to represent experimental classification of each data point. The interactive terms were assigned a factor level to ensure visual representation of clusters on the final scatter plot. Prior to scatterplot generation, the eigenvalues, cumulative variance, and correlation matrices were examined to determine which dimensions explain the majority of variance within the data set and which variables correlate strongly to each dimension. PCA, eigenvalue, and centroid data were extracted from the data and mapped onto the final PCA scatter plot. Clusters are visualised by differential colouring of the experimental classification and 95% confidence ellipses. The axes represent the amount of explained variance in each dimension (>70%). The full code for the AMOT/CTNNB1 and YAP localisation/expression PCAs can be found at: https://github.com/Collier0123/AMOT_CNTTB1_PCA, and https://github.com/Collier0123/YAP1_PCA, respectively.

### Live embryo time-lapse imaging and analyses

Transgenic mT^+33^ 2-cell stage embryos were recovered into M2 media and single blastomeres microinjected with siNTC or si*Tead4* (10µM) and H2B-Venus mRNA (40ng/µl) and cultured in global® (LGGG-020, Origio) supplied with 1mg/ml BSA at 37°C in 5% CO_2_ and 5% O_2_ for 48 hours; as described above. Time-lapse imaging was performed using Viventis LS1 Live inverted light-sheet microscope system (Viventis Microscopy Sarl, Switzerland), as previously described^74^. Complete embryo imaging was performed between 89-105.5 hours post-hCG (relative to super-ovulation). Embryos were placed in a 125μL culture medium covered with 125μL oil (OVOIL, Vitrolife) in a sample-holder multi-well (Viventis Microscopy). Fifty-one 3μm optical sections were taken with a 1024×1024-pixel image resolution using 10-minute time intervals. Venus and mT fluorescence were excited by 515 and 561nm laser lines, respectively. To detect Venus emissions, a band-pass filter with a 539/30 bandwidth was used. mT was detected using a triple-band-pass filter at 488/561/640. Marked clones were tracked using Imaris software (version 9.9.1, Oxford Instruments), up to 100 frames (16.5 hours). Light-sheet time-lapse imaging files are available in the BioStudies database under accession number S-BSST2271 (https://www.ebi.ac.uk/biostudies/studies/S-BSST2271). Other data are available from the corresponding author upon reasonable request. Lineage trees were generated automatically using the spots tool and edited manually. Each track was assigned a label based on its localisation and/or internalisation mechanism (or apoptotic events). At frame 100, inner cells originating from an internalisation were recorded manually for each video (Fig. 2E), and each internalisation event was placed into a frequency matrix (heat map) corresponding to cell stage, normalised to the sample size (n) (Fig. 2F). Full-length example time-lapse movies are provided (supplementary movie 1).

### Quantitative reverse-transcription PCR (RT-qPCR)

To assess dsRNA/siRNA KD, total RNA was prepared from 30 cultured 32-cell (E3.5) blastocysts (typically microinjected in both blastomeres at the 2-cell stage) using the PicoPure RNA Isolation kit (Arcturus). Eluted RNA (10μl) was DNase I–treated (Ambion DNA-free kit) and reverse-transcribed to cDNA (30μl) with oligo-dT priming (Superscript III, Invitrogen). Per quantitative PCR run, specific gene transcripts were measured as technical triplicates; 0.5μl of diluted cDNA (1:3 in nuclease-free water) was used as template in 10μl SYBR-Green-based reactions (Mastermix - Qiagen), with gene-specific oligonucleotide primer pairs (final concentration 400 nM; sequences provided in S.Tabs. 24), on a Bio-Rad CFX96 cycler system (adopting default automatic threshold detection settings). Within a control/experimental sample, transcript levels were internally normalised to those for *H2afz* and fold expression changes (including standard deviations) between the described groups calculated (according to the ΔΔCt method^75^); raw data summarised in S.Tabs. 2 & 16.

## Supporting information

Supplementary figures (1-15 & SM1), supplementary tables (1-24), supplementary movie 1 legend, and supplemenarty Excel workbook legends (1-3, & stats)

Supplementary Excel data workbook 1

Supplementary Excel data workbook 2

Supplementary Excel data workbook 3

Supplementary Excel statistics summary workbook 1

## ACKNOWLEDGEMENTS

We thank the Institute of Parasitology (Biology Centre of the Czech Academy of Sciences, České Budějovice) for housing mice. We are grateful to Marta Gajewska (Institute of Oncology, Warsaw, Poland) and Anna Piliszek (Institute of Genetics and Animal Breeding, Polish Academy of Sciences, Jastrzębiec, Poland) for providing founder CBA/W mice. Special thanks go to Hiroshi Sasaki (Osaka University, Japan) for the anti-AMOT antibody.

