## Supplementary figures (1-15 & SM1), supplementary tables (1-24), supplementary movie 1 legend, and supplemenarty Excel workbook legends (1-3, & stats) for "TEAD4 regulates apical domain homeostasis and cell-positioning to maintain the trophectoderm lineage during preimplantation mouse embryo development"

**Supplementary Figures and legends (Figs. S1-S15)**

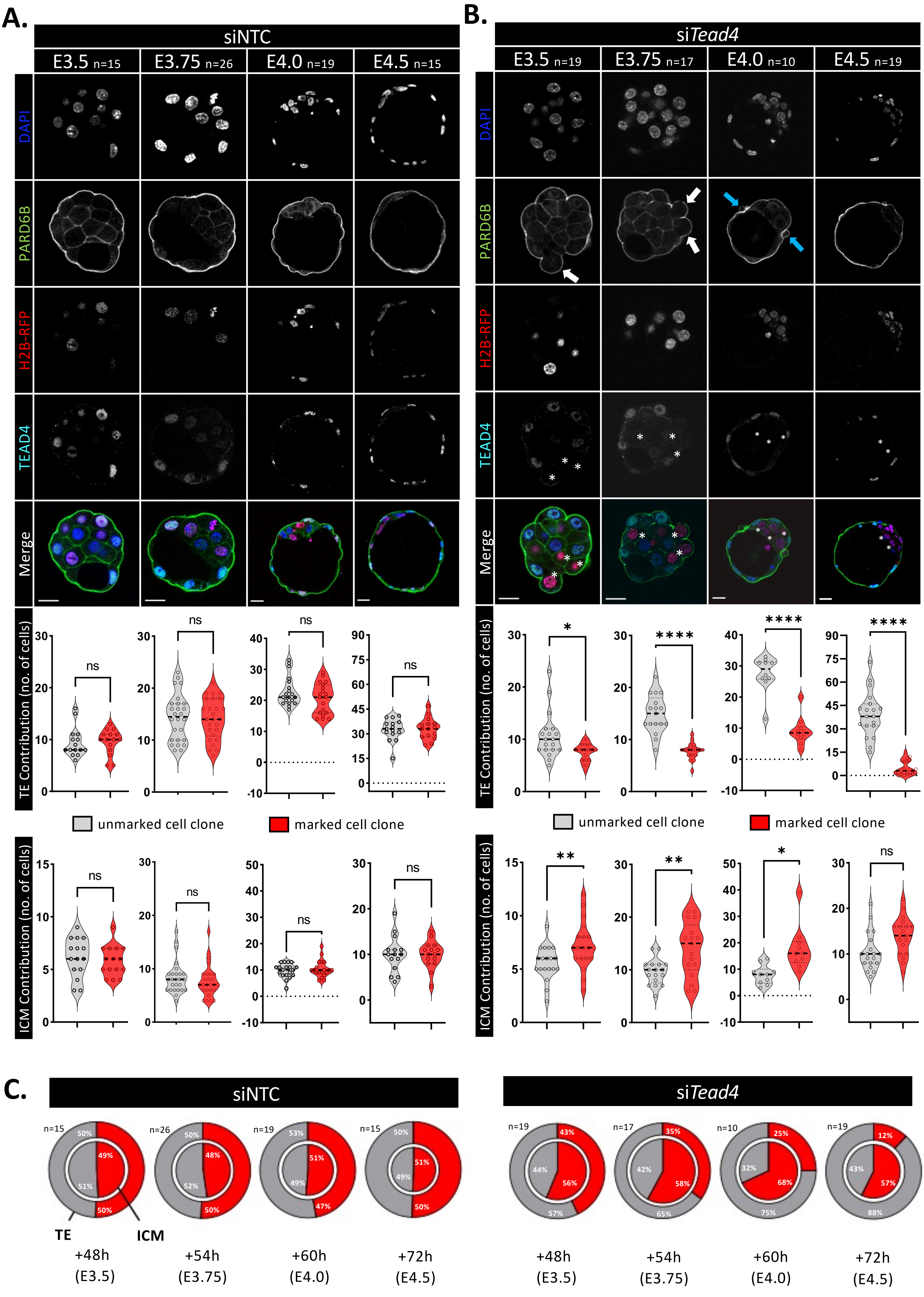

unmarked cell clone marked cell clone

Fig. S1 (part one).

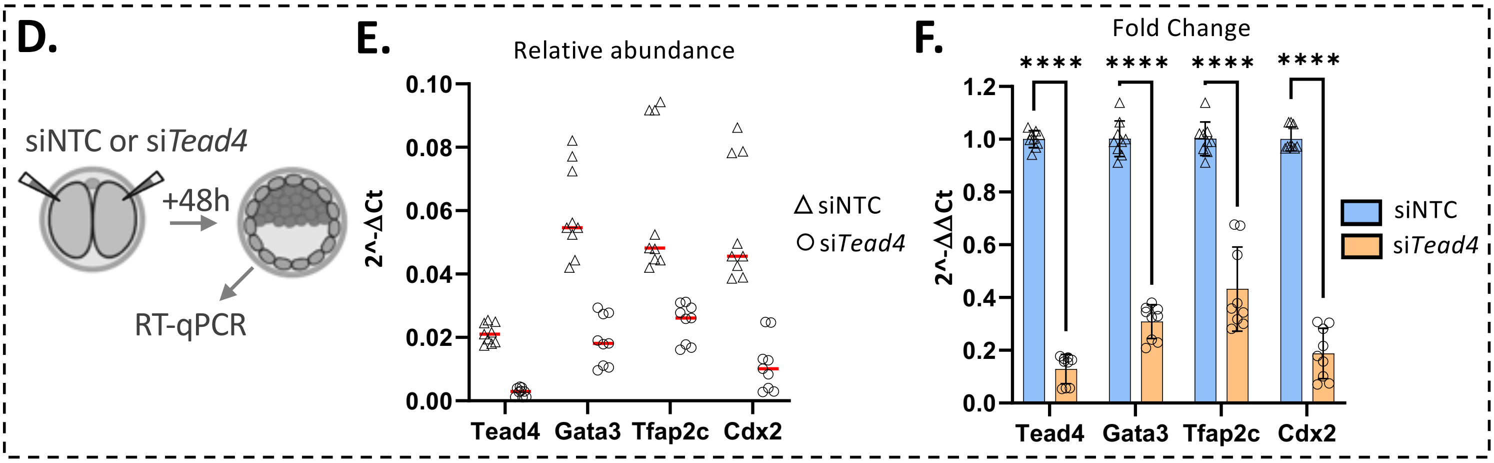

**Fig. S1 (part two).**

**Supplementary Figure S1: Clonal *Tead4* KD induces atypical cell allocation events during blastocyst maturation**

**A)** Upper panels: Example single z-section IF-stained greyscale confocal micrographs (+pseudo-coloured merge) of siNTC-injected clone containing staged blastocysts (same embryos as in Fig. 1). The number of embryos per group (n) is indicated. Scale bar = 20 µm. Lower charts: Violin plots of unmarked (grey) and H2B-RFP marked (red) cells to outer TE (top) and ICM (bottom) populations in siNTC-injected staged blastocysts. Dashed lines indicate median values. No significant differences (ns) in clone contribution between unmarked /marked cells were observed at any stage (*$p<0.05$).

**B)** Upper panels: Similar images as in A), but for si*Tead4*-injected staged blastocysts. Asterisks indicate confirmed TEAD4 protein KD/absence in H2B-RFP marked clones (red). Arrows highlight morphologically abnormal outer-cell membrane structures: apical domain blebs at E3.5 & E3.75 (white arrows) and surface vesicle-like structures, containing PARD6B, at E4.0 (blue arrows). Lower charts: Violin plots illustrating unmarked/marked clone number contributions to outer TE and ICM in staged si*Tead4* blastocysts, as in panel A).

**C)** Concentric pie-charts detailing average percentage contribution of unmarked (grey) and marked (red) cells to outer TE (outer ring segment) and ICM (inner segment) populations in siNTC (left) and si*Tead4* (right) staged blastocysts. Numbers of embryos analysed per group (n) indicated; as in A) & B).

**D)** RT-qPCR experimental schema: Both blastomeres of 2-cell stage embryos were microinjected with either control siNTC or si*Tead4* constructs, to induce global *Tead4* KD, and cultured for 48 hours to the E3.5 blastocyst stage. Normalised mRNA expression levels of *Tead4* (to confirm KD) and TE markers *Gata3*, *Tfap2c*, & *Cdx2* were measured by RT-qPCR (see S.Tabs. 2).

**E)** Normalised relative expression of *Tead4* and indicated TE marker transcripts in siNTC- (triangles) and si*Tead4*- (circles) treated blastocysts. The median values are indicated by red bars; data are from two biological replicates with at least three technical replicates each.

**F)** Median gene expression fold changes of the indicated transcripts in si*Tead4* versus siNTC embryos (orange vs. blue); based on data in panel E). Statistical significance (*$p<0.05$) and standard deviations (error bars) are shown.

*All raw clonal allocation data are summarised in S.Tabs. 1a-e. Statistical analyses were performed based on determined data distribution (normal or non-normal) and application of appropriate tests; details in supplementary statistics Excel workbook.*

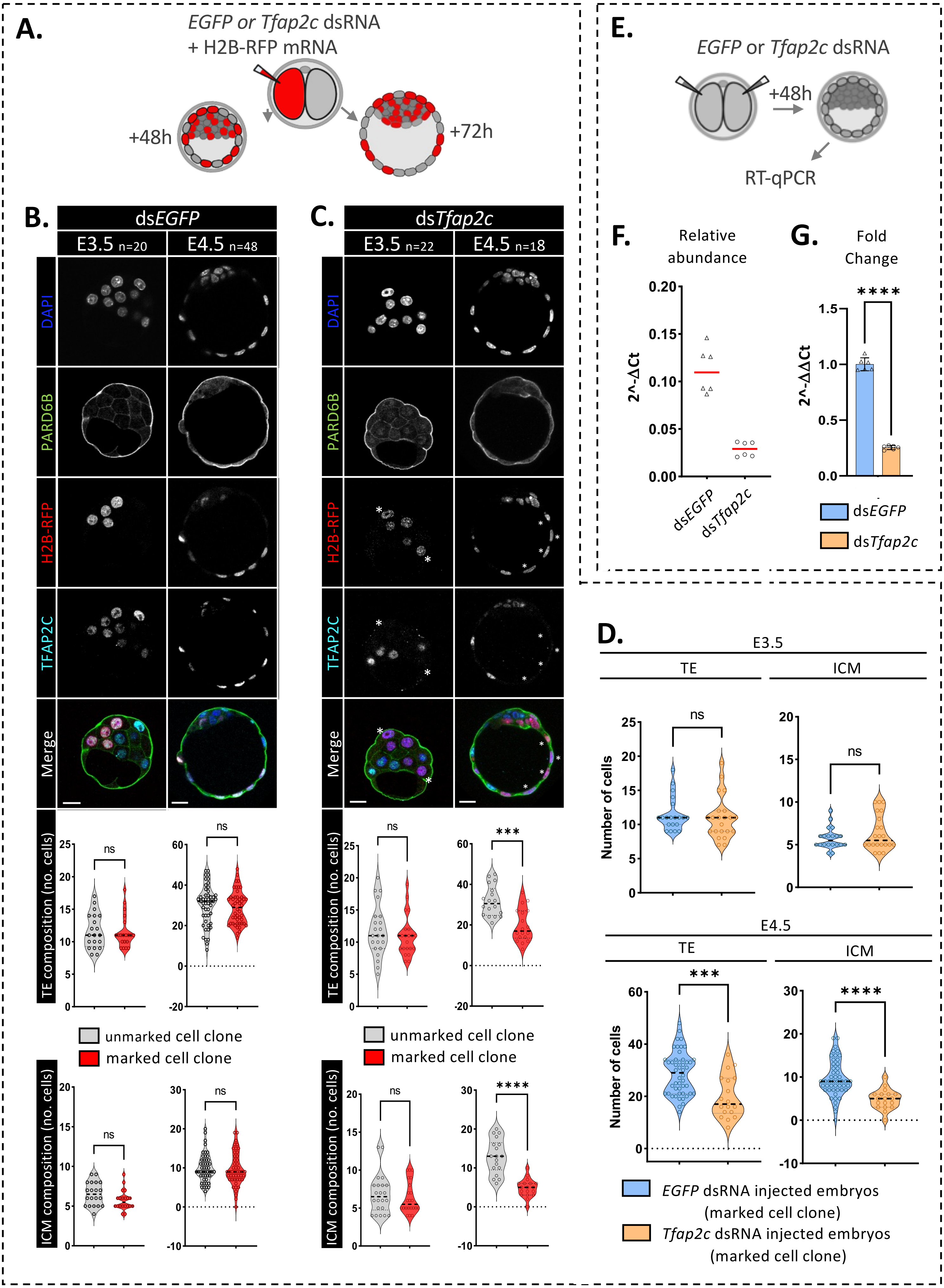

A.
EGFP or Tfap2c dsRNA
+ H2B-RFP mRNA
+48h
+72h
B.
dsEGFP
E3.5 n=20
E4.5 n=48
C.
dsTfap2c
E3.5 n=22
E4.5 n=18
DAPI
PARD6B
H2B-RFP
TFAP2C
Merge
TE composition (no. cells)
ICM composition (no. cells)
unmarked cell clone
marked cell clone
ns
***
****
E.
EGFP or Tfap2c dsRNA
+48h
RT-qPCR
F.
Relative abundance
2^-ΔCt
dsEGFP
dsTfap2c
G.
Fold Change
2^-ΔΔCt
dsEGFP
dsTfap2c
D.
E3.5
TE
ICM
E4.5
Number of cells
EGFP dsRNA injected embryos
(marked cell clone)
Tfap2c dsRNA injected embryos
(marked cell clone)

Fig. S2

**Supplementary Figure S2: Clonal *Tfap2c* KD does not cause abnormal outer/TE to ICM allocation during blastocyst maturation, unlike clonal *Tead4* KD**

**A)** Schematic of dsRNA microinjection procedure generating staged blastocysts (E3.5 & E4.5) containing either control/non-targeting (dsEGFP) or *Tfap2c* KD (ds*Tfap2c*) marked clones—comprising 50% of all cells; allowing assessment of clone contributions to the outer TE or ICM positions.

**B)** Upper panels: Example single z-section IF-stained greyscale confocal micrographs (+pseudo-coloured merge) of control (non-targeting) dsEGFP-injected clone containing staged blastocysts. The number of embryos per group (n) is indicated. Scale bar = 20 µm. Lower charts: Violin plots of unmarked (grey) and H2B-RFP marked (red) cells to outer TE (top) and ICM (bottom) populations in dsEGFP-injected staged blastocysts. Dashed lines indicate median values. No significant differences (ns) in clone contribution between unmarked /marked cells were observed at either stage (E3.5 & E4.5, $^*p<0.05$).

**C)** Upper panels: Similar images as in B), but for ds*Tfap2c*-injected staged blastocysts. Asterisks indicate confirmed TFAP2C protein KD/absence in H2B-RFP marked clones (red). Lower charts: Violin plots illustrating unmarked/marked clone number contributions to outer TE and ICM in staged ds*Tfap2c* staged blastocysts, as in panel B).

**D)** Comparison of TE and ICM cell number contributions of only the marked dsEGFP (blue) versus ds*Tfa2p2c* (orange) clones, at the indicated stages. Median contributions shown by dashed lines, with significant inter-group differences indicated ($^*p<0.05$). As in C) only significant differences in reduced contribution of marked ds*Tfap2c* clones to both TE and ICM at E4.5 were observed (reflecting reduced general viability of the clone, *c.f.* TE-to-ICM allocations seen in si*Tead4* embryos).

**E)** RT-qPCR experimental schema: Both blastomeres of 2-cell stage embryos were microinjected with either control dsEGFP or ds*Tfap2c* constructs, to induce global *Tfap2c* KD, and cultured for 48 hours to the E3.5 blastocyst stage. Normalised mRNA expression levels of *Tfp2c* (to confirm KD) were measured by RT-qPCR (see S.Tabs. 2).

**F)** Normalised relative expression of *Tfap2c* transcripts in dsEGFP (triangles) and ds*Tfap2c* (circles) blastocysts. The median values are indicated by red bars; data are from two triplicate technical replicates.

**G)** Median *Tfap2c* gene expression fold changes in ds*Tfap2c* versus dsEGFP embryos (orange vs. blue); based on data in panel F). Statistical significance ($^*p<0.05$) and standard deviations (error bars) are shown.

*All raw clonal allocation data are summarised in S.Tabs. 3a & 3b. Statistical analyses were performed based on determined data distribution (normal or non-normal) and application of appropriate tests; details in supplementary statistics Excel workbook.*

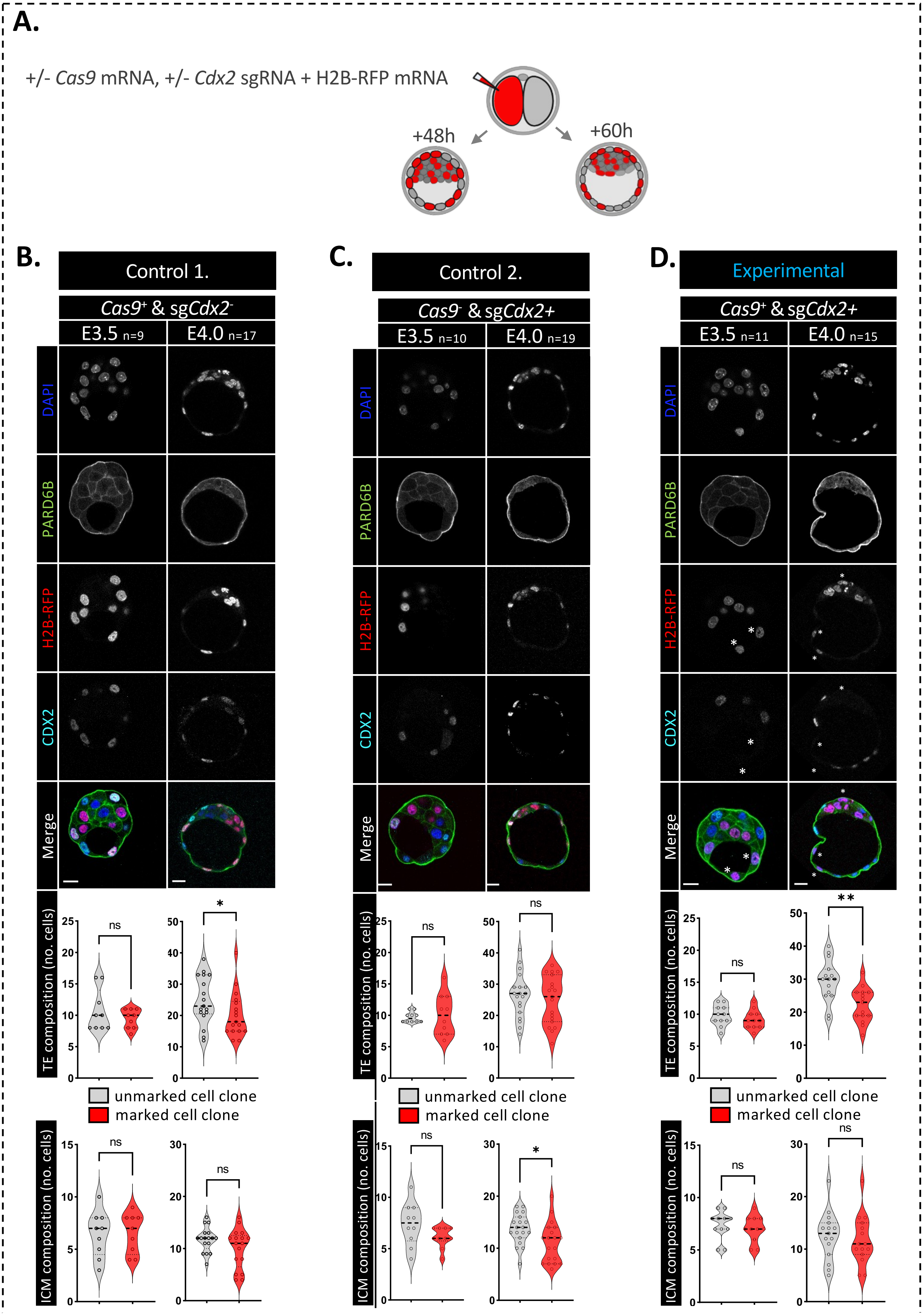

Fig. S3 (part one).

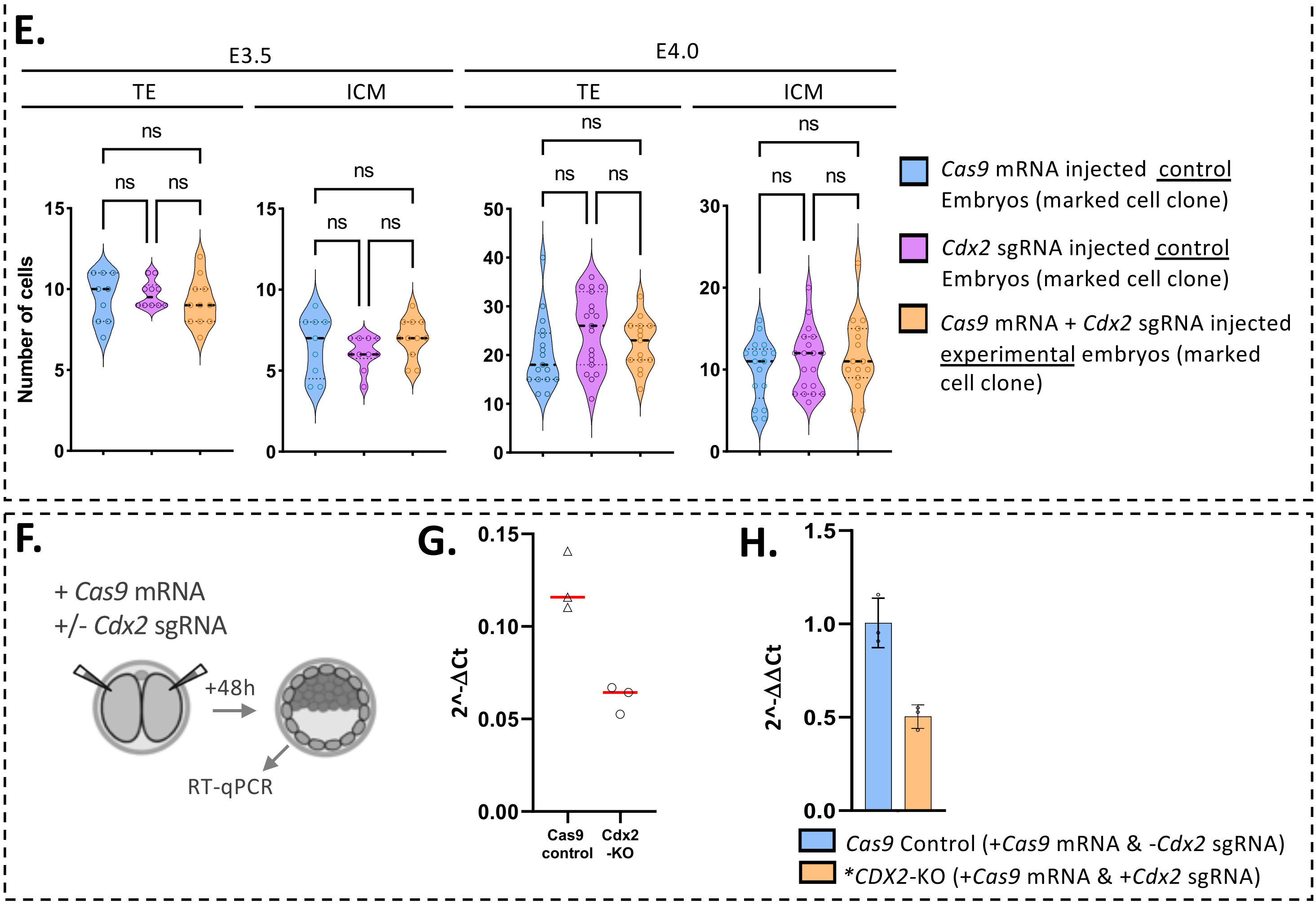

**Fig. S3 (part two).**

**Supplementary Figure S3: Clonal CRISPR-Cas9-mediated *Cdx2* knockout (KO) does not cause abnormal TE to ICM allocation during blastocyst maturation, unlike clonal *Tead4* KD**

**A)** Schematic of CRISPR-Cas9 (±*Cas9* mRNA & *Cdx2* sgRNA/sg*Cdx2*) microinjection procedure generating staged blastocysts (E3.5 & E4.0) containing either control (*Cas9*+ & sg*Cdx2*- or *Cas9*- & sg*Cdx2*+) or *Cdx2* KO (*Cas9*+ & sg*Cdx2*+) marked clones—comprising 50% of all cells; allowing assessment of clone contributions to the outer TE or ICM positions.

**B)** Upper panels: Example single z-section IF-stained greyscale confocal micrographs (+pseudo-coloured merge) of control (non-targeting) *Cas9*+ & sg*Cdx2*- injected clone containing staged blastocysts. The number of embryos per group (n) is indicated. Scale bar = 20 µm. Lower charts: Violin plots of unmarked (grey) and H2B-RFP marked (red) cells to outer TE (top) and ICM (bottom) populations in *Cas9*+ & sg*Cdx2*- injected staged blastocysts. Dashed lines indicate median values. Significant differences in clone contribution between unmarked /marked cells are highlighted (E3.5 & E4.0, *$p<0.05$).

**C)** Upper panels: Similar images as in B), but for control (lacking endonuclease activity) *Cas9*- & sg*Cdx2+* injected staged blastocysts. Lower charts: Violin plots illustrating unmarked/marked clone number contributions to outer TE and ICM in staged *Cas9*- & sg*Cdx2+* injected blastocysts, as in panel B).

**D)** Upper panels: Similar images as in B) & C), but for experimental (*Cdx2* KO) *Cas9*+ & sg*Cdx2+* injected staged blastocysts. Asterisks indicate confirmed outer-cell CDX2 protein KD/absence in H2B-RFP marked clones (red). Lower charts: Violin plots illustrating unmarked/marked clone number contributions to outer TE and ICM in staged *Cas9*+ & sg*Cdx2+* (*Cdx2* KO) injected blastocysts, as in panels B) & C).

**E)** Comparison of TE and ICM cell number contributions of only the marked controls (blue; *Cas9*+ & sg*Cdx2*- & pink; *Cas9*- & sg*Cdx2*+) versus experimental *Cdx2* KO (orange *Cas9*+ & sg*Cdx2*+) clones, at the indicated stages. Median contributions shown by dashed lines, with significant inter-group differences indicated (*$p<0.05$). Note, all marked clones contribute equally (ns) to both E3.5 and E4.5 TE/ICM.

**F)** RT-qPCR experimental schema: Both blastomeres of 2-cell stage embryos were microinjected with either *Cas9* mRNA minus *Cdx2* sgRNA (control) or both constructs to induce global *Cdx2* KO, and cultured for 48 hours to the E3.5 blastocyst stage. Normalised mRNA expression levels of *Cdx2* were measured by RT-qPCR (see S.Tabs. 2).

**G)** Normalised relative expression of *Cdx2* transcripts in *Cas9*+ & sg*Cdx2*- control (triangles) and *Cdx2* KO (circles) blastocysts. The median values are indicated by red bars.

**H)** Median *Cdx2* gene expression fold changes in *Cdx2* KO versus *Cas9*+ & sg*Cdx2*- control embryos (orange vs. blue); based on data in panel G). Statistical significance (*$p<0.05$) and standard deviations (error bars) are shown (*note, *Cdx2* KO strategy generates genomic indels producing, reduced and nonsense frame-shift mutant transcripts, detectable by RT-qPCR; but do not yield detectable CDX2 proteins – see panel D).

*All raw clonal allocation data are summarised in S.Tabs. 4a & 4b. Statistical analyses were performed based on determined data distribution (normal or non-normal) and application of appropriate tests; details in supplementary statistics Excel workbook.*

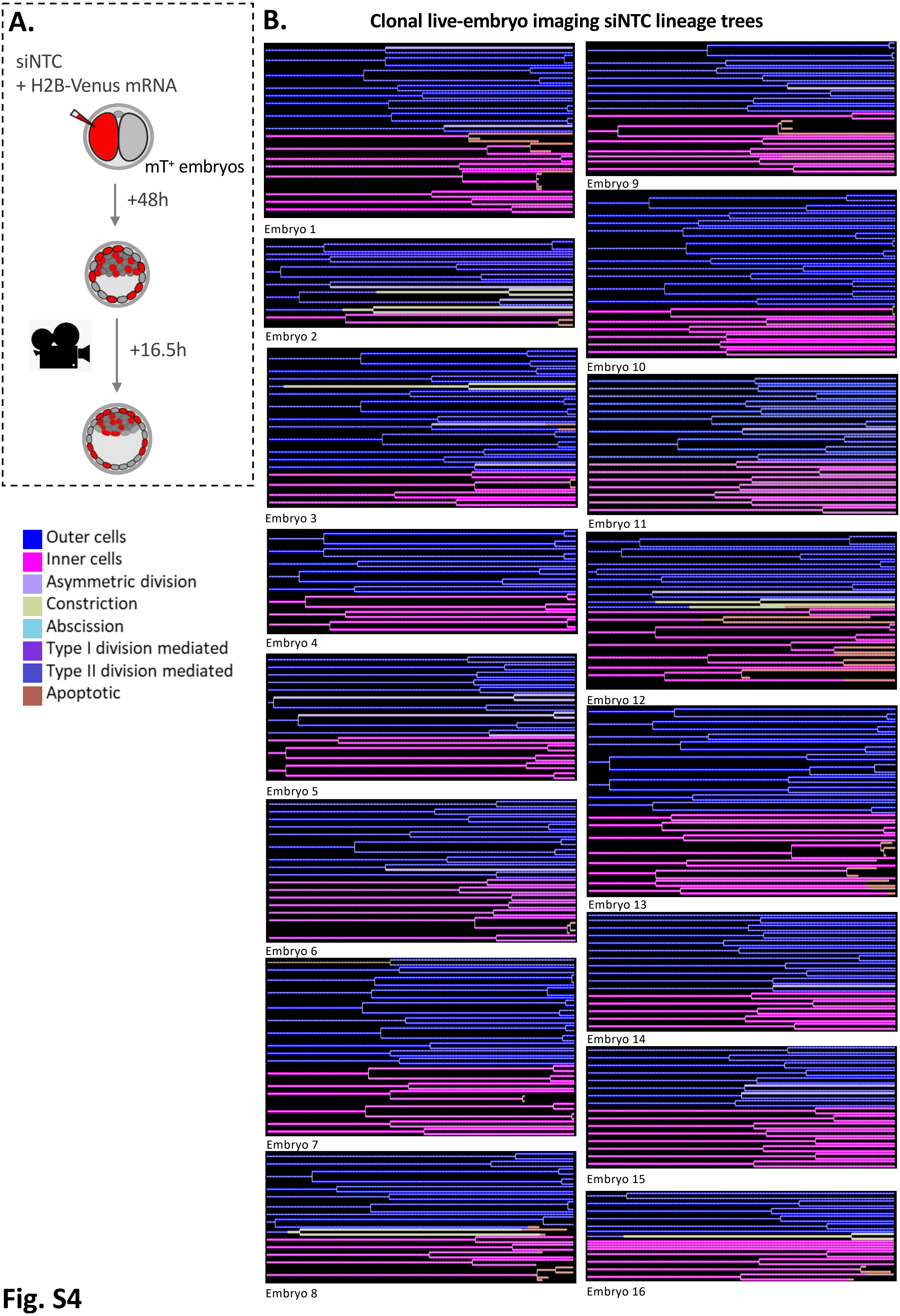
A.
siNTC
+ H2B-Venus mRNA
mT+ embryos
+48h
+16.5h
Outer cells
Inner cells
Asymmetric division
Constriction
Abscission
Type I division mediated
Type II division mediated
Apoptotic
B.
Clonal live-embryo imaging siNTC lineage trees
Embryo 1
Embryo 2
Embryo 3
Embryo 4
Embryo 5
Embryo 6
Embryo 7
Embryo 8
Embryo 9
Embryo 10
Embryo 11
Embryo 12
Embryo 13
Embryo 14
Embryo 15
Embryo 16

Fig. S4

**Supplementary Figure S4: Lineage-tracing of marked control siNTC clones in embryos subject to light-sheet microscopy live-embryo imaging (*related to* *Fig. 2*)**

**A)** Experimental schematic of 2-cell stage microinjections of transgenic membrane-Tomato expressing ($mT^{+}$) 2-cell embryos used to generate early 32-cell stage blastocysts (after 48 hours of culture) containing H2B-Venus marked control (siNTC) clones (comprising 50% of all cells), subject to 16.5 hours of light-sheet microscopy live-embryo imaging (10 minute intervals); allowing dynamic assessment of individual clone contributions to the outer TE or ICM positions.

**B)** Individual embryo developmental cell lineage trees of the fate of marked siNTC clones (originating in either outer or inner positions at the start of recording) during blastocyst maturation. Cell internalisation events, via various mechanisms, or apoptotic events are colour-coded as indicated. n=16.

*A summary of the origin of all observed internalisation events and their frequencies in provided in S.Tabs. 5a & 5b.*

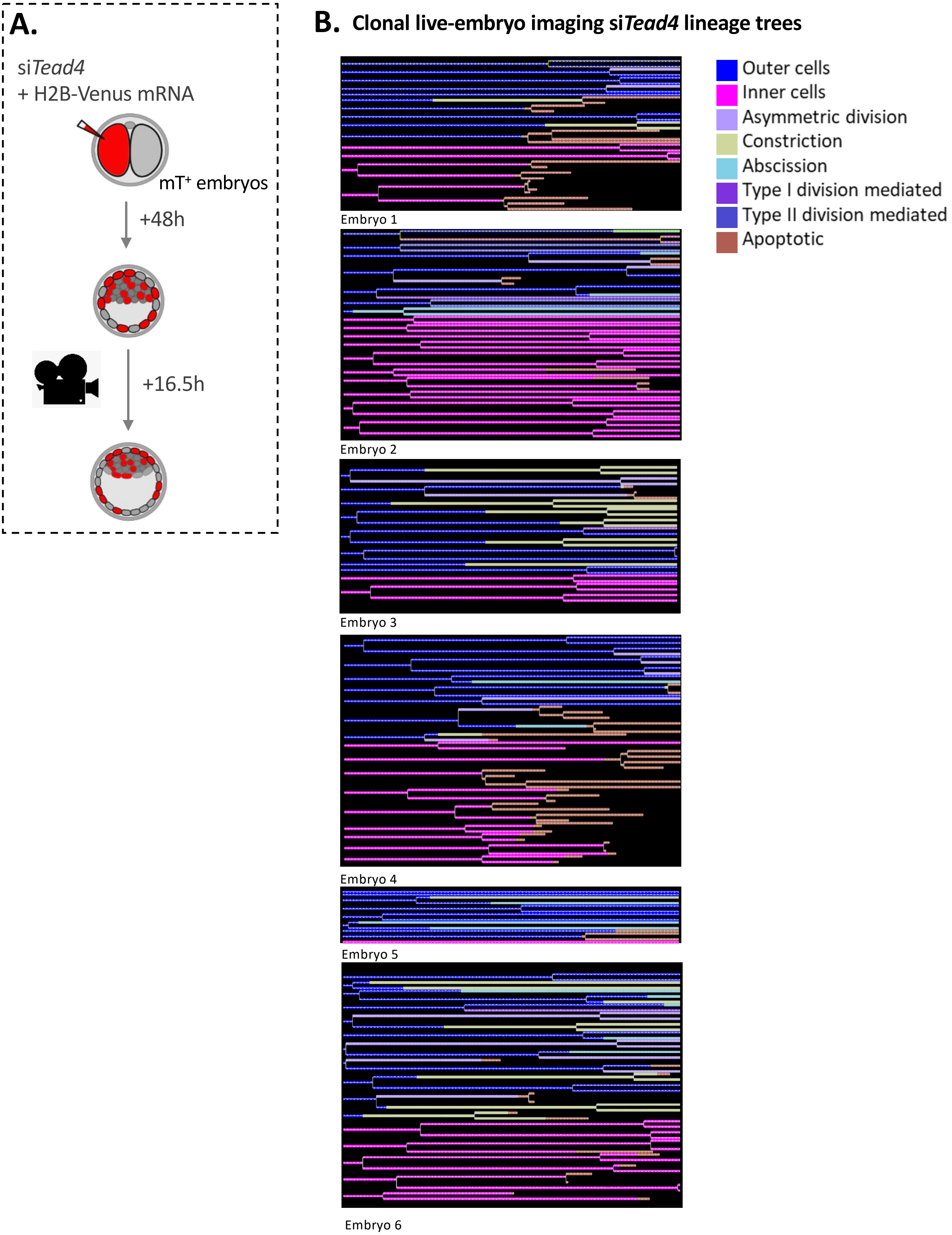
A.
siTead4
+ H2B-Venus mRNA
mT+ embryos
+48h
+16.5h
B. Clonal live-embryo imaging siTead4 lineage trees
Outer cells
Inner cells
Asymmetric division
Constriction
Abscission
Type I division mediated
Type II division mediated
Apoptotic
Embryo 1
Embryo 2
Embryo 3
Embryo 4
Embryo 5
Embryo 6

**Fig. S5**

**Supplementary Figure S5: Lineage-tracing of marked si*Tead4* clones in embryos subject to light-sheet microscopy live-embryo imaging (*related to* Fig. 2).**

**A)** Experimental schematic of 2-cell stage microinjections of transgenic membrane-Tomato expressing ($mT^{+}$) 2-cell embryos used to generate early 32-cell stage blastocysts (after 48 hours of culture) containing H2B-Venus marked si*Tead4* clones (comprising 50% of all cells), subject to 16.5 hours of light-sheet microscopy live-cell imaging (10 minute intervals); allowing dynamic assessment of individual clone contributions to the outer TE or ICM positions.

**B)** Individual embryo developmental cell lineage trees of the fate of marked si*Tead4* clones (originating in either outer or inner positions at the start of recording) during blastocyst maturation. Cell internalisation events, via various mechanisms, or apoptotic events are colour-coded as indicated. n=6.

*A summary of the origin of all observed internalisation events and their frequencies in provided in S.Tabs. 5a & 5b.*

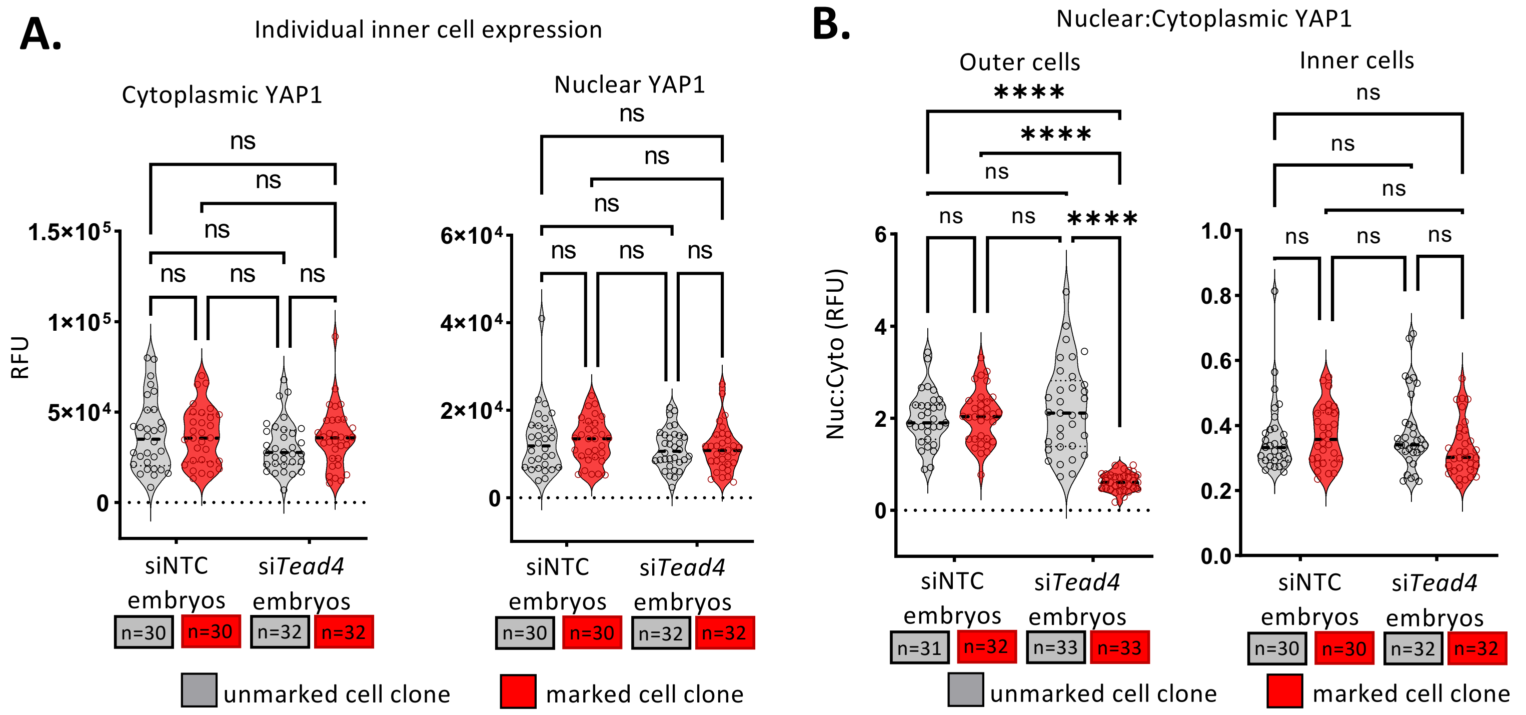

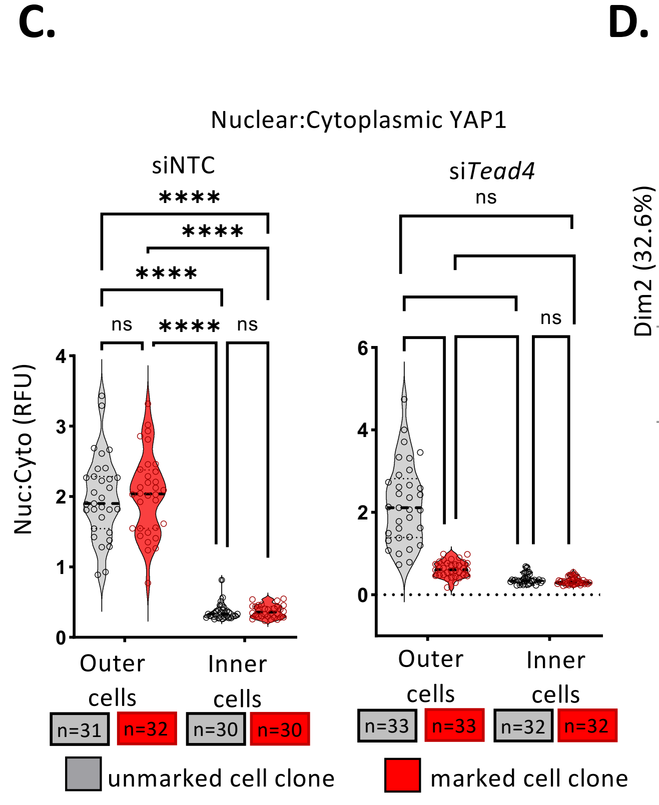

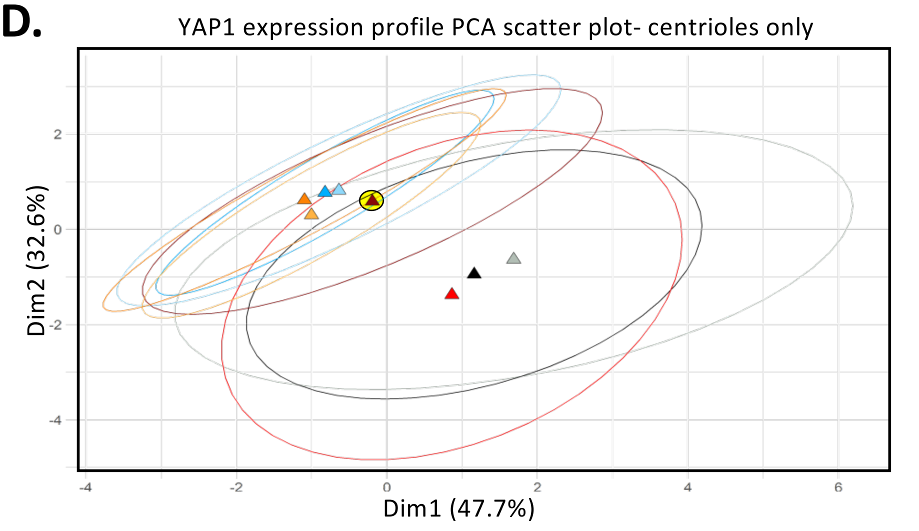

| YAP1 localisation PCA variables (per cell assayed outer/inner cell) | |
|---|---|
| 1 | Cytoplasmic area (µm^2) |
| 2 | Nuclear area (µm^2) |
| 3 | Nuclear YAP1 RFU |
| 4 | Cytoplasmic YAP1 RFU |
| 5 | Nuc:Cyto YAP1 Ratio (RFU) |

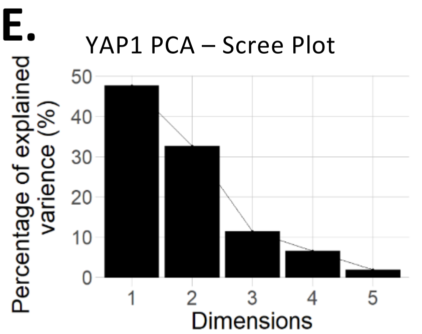

| | Eigen-values | % of variance | Cumulative % of var. |
|---|---|---|---|
| Dim1 | 2.383 | 47.652 | 47.652 |
| Dim2 | 1.629 | 32.586 | 80.238 |
| Dim3 | 0.571 | 11.429 | 91.667 |
| Dim4 | 0.327 | 6.541 | 98.208 |
| Dim5 | 0.090 | 1.792 | 100.000 |

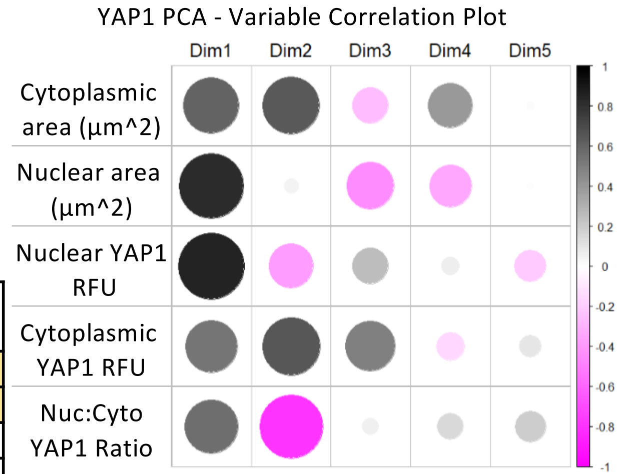

| Vai able | Correlation Coefficient | | | | |
|---|---|---|---|---|---|
| | Dim1 | Dim2 | Dim3 | Dim4 | Dim5 |
| 1 | 0.602 | 0.646 | -0.253 | 0.396 | 0.012 |
| 2 | 0.826 | 0.043 | -0.440 | -0.349 | 0.008 |
| 3 | 0.858 | -0.390 | 0.258 | 0.065 | -0.205 |
| 4 | 0.536 | 0.659 | 0.494 | -0.158 | 0.099 |
| 5 | 0.561 | -0.790 | 0.053 | 0.141 | 0.194 |

**Fig. S6**

**Supplementary Figure S6: Effects of clonal *Tead4* KD on YAP1 expression and subcellular localisation**

**A)** Quantitative analysis of single inner-cell YAP1 expression levels (RFU), separated into cytoplasmic and nuclear compartments, in siNTC and si*Tead4* blastocysts. Data are further divided into unmarked (grey) and marked (red) clone populations. Median levels are indicated by dashed lines; significant differences are marked (*p<0.05). Sample sizes (n) are noted. Quantified protein expression value (RFU) data are summarised in Supplementary Tables S.Tabs. 6a.

**B)** Determined single cell nuclear:cytoplasmic YAP1 expression ratios (RFU) for indicated outer- and inner-cell populations, in siNTC and si*Tead4* blastocysts. Data are further divided into unmarked (grey) and marked (red) clone populations. Median ratios are indicated by dashed lines; significant differences are marked (*p<0.05). Sample sizes (n) are noted. Quantified protein expression values (RFU) used to generate displayed ratios are summarised in S.Tabs. 6a.

**C)** As in B) but comparing nuclear:cytoplasmic YAP1 expression ratios within, rather than between, siNTC and si*Tead4* groups. Sample sizes (n) are noted. Quantified protein expression values (RFU) used to generate displayed ratios are summarised in S.Tabs. 6a.

**D)** Principal Component Analysis (PCA) of subcellular YAP1 expression levels (as in Fig. 3D), comparing outer and inner cells within unmarked and marked clones of siNTC and si*Tead4* blastocysts. Ellipses depict 95% confidence intervals, and mean cluster centres (triangles) are shown without displaying all individual cell data points to facilitate interpretation – outer marked si*Tead4* KD clones highlighted in yellow. PCA variables used are listed in the accompanying table and described in the Materials & Methods section.

**E)** Scree plot showing the percentage of total variance explained by each PCA dimension (Dim), with expanded details in the accompanying table. The first two dimensions (Dim1 and Dim2) together explain over 70% of the variance (specifically 80.2%), and are used in the PCA plot in panel D) (Dim1 = 47.7%, Dim2 = 32.6%).

**F)** Variable correlation plot, illustrating the relationships between the employed PCA variables within each dimension of explained variance. Circle area indicates the strength of the correlation, while graduated shading (black or pink) denotes positive or negative correlations, respectively. The accompanying table provides the numerical correlation coefficients.

*Statistical analyses for panels A), B) & C) were performed based on determined data distribution (normal or non-normal) and application of appropriate tests; details in supplementary statistics Excel workbook.*

**A.**

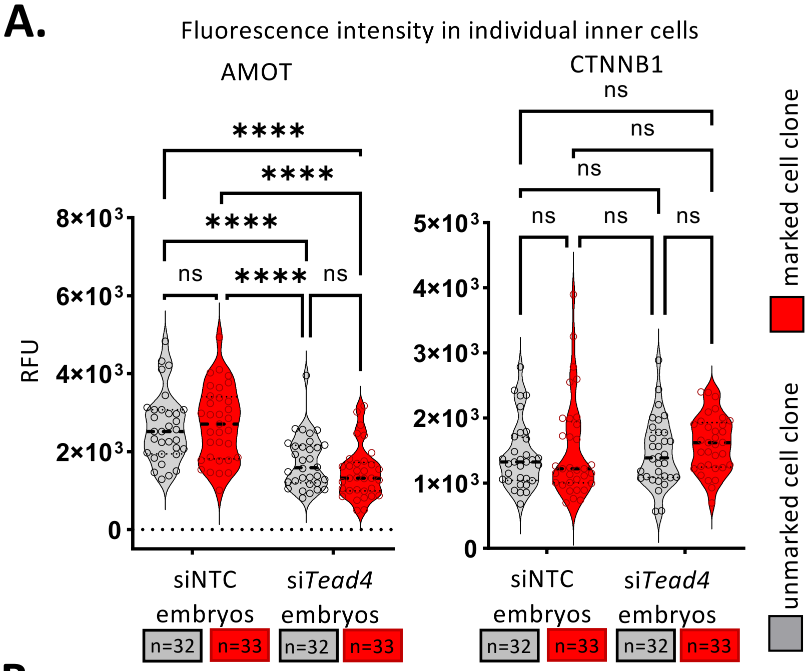

**B.**

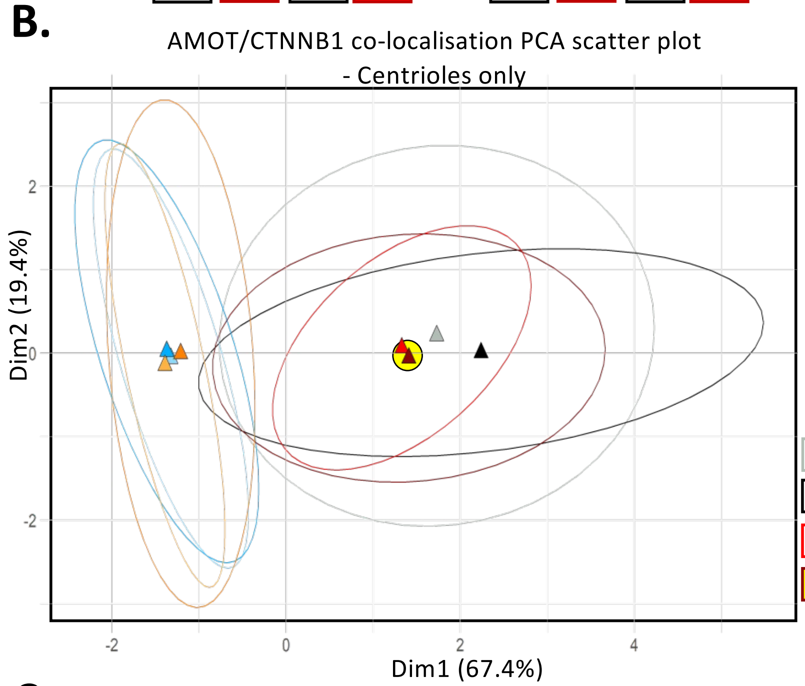

| AMOT/CTNNB1 co-localisation PCA variables (per assayed outer cell) | |
|---|---|
| 1 | Length of assayed domain (apical or lateral - µM) |
| 2 | AMOT RFU (at given apical or lateral domain) |
| 3 | CTNNB1 RFU (at given apical or lateral domain) |
| 4 | Median RFU ratio (AMOT/CTNNB1 - at given apical or lateral domain) |

Apical: siNTC, unmarked.
Apical: siNTC, marked.
Apical: si*Tead4*, unmarked
Apical: si*Tead4*, marked
Lateral: siNTC, unmarked
Lateral: siNTC, marked
Lateral : si*Tead4*, unmarked
Lateral : si*Tead4*, marked

**C.**

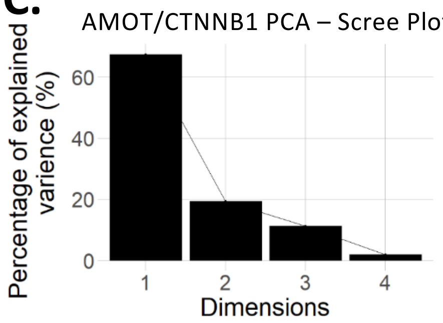

| | Eigen-values | % of variance | Cumulative % of var. |
|---|---|---|---|
| Dim1 | 2.695 | 67.373 | 67.373 |
| Dim2 | 0.778 | 19.446 | 86.819 |
| Dim3 | 0.448 | 11.208 | 98.027 |
| Dim4 | 0.079 | 1.973 | 100.000 |

**D.**

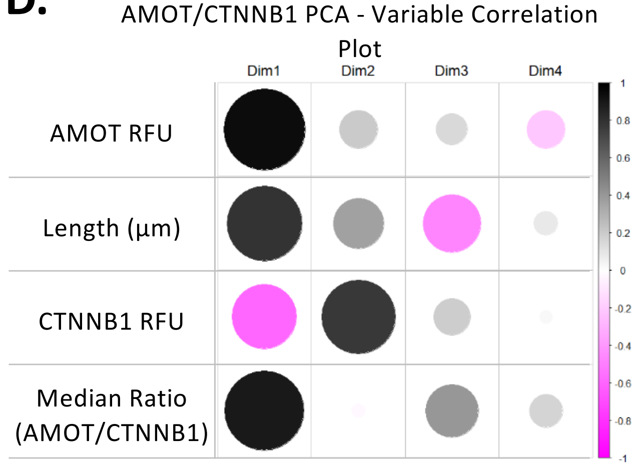

| Variable | Correlation Value | | | |
|---|---|---|---|---|
| | Dim1 | Dim2 | Dim3 | Dim4 |
| 1 | 0.944 | 0.209 | 0.142 | -0.212 |
| 2 | 0.797 | 0.364 | -0.475 | 0.084 |
| 3 | -0.599 | 0.775 | 0.198 | 0.023 |
| 4 | 0.900 | -0.025 | 0.404 | 0.163 |

**Fig. S7**

**Supplementary Figure S7: Impact of clonal *Tead4* KD on AMOT and CTNNB1 expression and membrane co-localisation**

**A)** Quantitative analysis of single inner-cell AMOT (left) & CTNNB1 (right) membrane-associated expression levels (RFU) in siNTC and si*Tead4* blastocysts. Data are further divided into unmarked (grey) and marked (red) clone populations. Median levels are indicated by dashed lines; significant differences are marked (*p<0.05). Sample sizes (n) are noted. Quantified protein expression value (RFU) data are summarised in S.Tabs. 7a & 7b.

**B)** Principal Component Analysis (PCA) of outer cell membrane-associated AMOT and CTNNB1 expression levels and protein co-localisation (as in Fig. 3I), comparing apical and lateral domains within unmarked and marked clones of siNTC and si*Tead4* blastocysts. Ellipses depict 95% confidence intervals, and mean cluster centres (triangles) are shown without displaying all individual cell data points to facilitate interpretation – outer marked si*Tead4* KD clone, apical domain analyses, highlighted in yellow. PCA variables used are listed in the accompanying table and described in the Materials & Methods section.

**C)** Scree plot showing the percentage of total variance explained by each PCA dimension (Dim), with expanded details in the accompanying table. The first two dimensions (Dim1 and Dim2) together explain over 70% of the variance (specifically 86.8%), and are used in the PCA plot in panel B (Dim1 = 67.4%, Dim2 = 19.4%).

**D)** Variable correlation plot, illustrating the relationships between the employed PCA variables within each dimension of explained variance. Circle area indicates the strength of the correlation, while graduated shading (black or pink) denotes positive or negative correlations, respectively. The accompanying table provides the numerical correlation coefficients.

*Statistical analyses for panel A) were performed based on determined data distribution (normal or non-normal) and application of appropriate tests; details in supplementary statistics Excel workbook.*

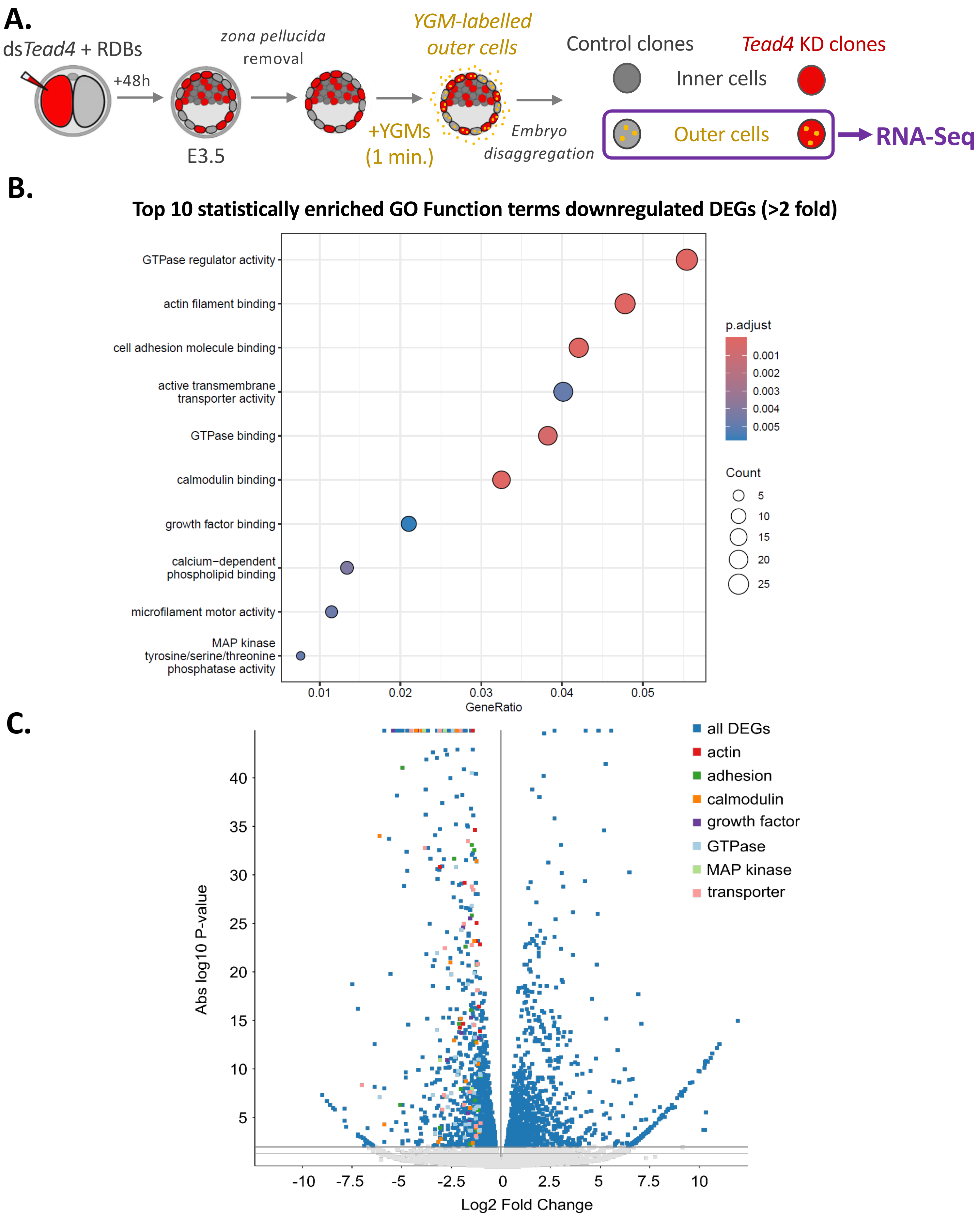

**Legend (amalgamated Function GO terms):**

actin – actin filament binding, microfilament motor activity

adhesion – cell adhesion molecule binding

calmodulin – calmodulin binding, calcium-dependent phospholipid binding

growth factor – growth factor binding

GTPase – GTPase regulator activity, GTPase binding

MAP kinase – MAP kinase tyrosine/serine/threonine phosphatase activity

transporter – active transmembrane transporter activity

**Fig. S8**

**Supplementary Figure S8: Clonal *Tead4* KD-induced downregulated (>2 fold) DEGs enriched GO functional terms**

**A)** Schematic of the dsRNA microinjection protocol used to generate E3.5 blastocysts containing *Tead4* KD (ds*Tead4*) clones, marked by co-injection with rhodamine-conjugated dextran beads (RDBs), representing 50% of all cells. At E3.5, outer cells were labelled via endocytosis of yellow-green microspheres (YGMs), followed by full embryo dissociation into single cells, to enable the identification of inner and outer cell populations from marked and unmarked clones, and preparation of RNA-Seq libraries to compare outer cell transcriptomes.

**B)** Dot plots describing the top ten enriched gene ontology (GO) terms, classified by function, for DEGs significantly downregulated (>2 fold and RPKM >0.5 in at least one of the clones) between marked *Tead4* KD versus unmarked control outer clones at the E3.5 stage after RNA-Seq.

**C)** Volcano plot comparing gene expression changes between RNA-Seq derived transcriptomes of E3.5 stage marked *Tead4* KD and unmarked control outer clone cell populations (all DEGs), highlighting significantly enriched GO function terms associated with genes >2 fold downregulated expression and RPKM >0.5 in at least one of the clones (note, related GO function terms were merged, according to the provided lower legend, to aid interpretation).

*S.Tab. 12 provides a summary of all the significantly enriched GO function terms and associated downregulated DEGs.*

.

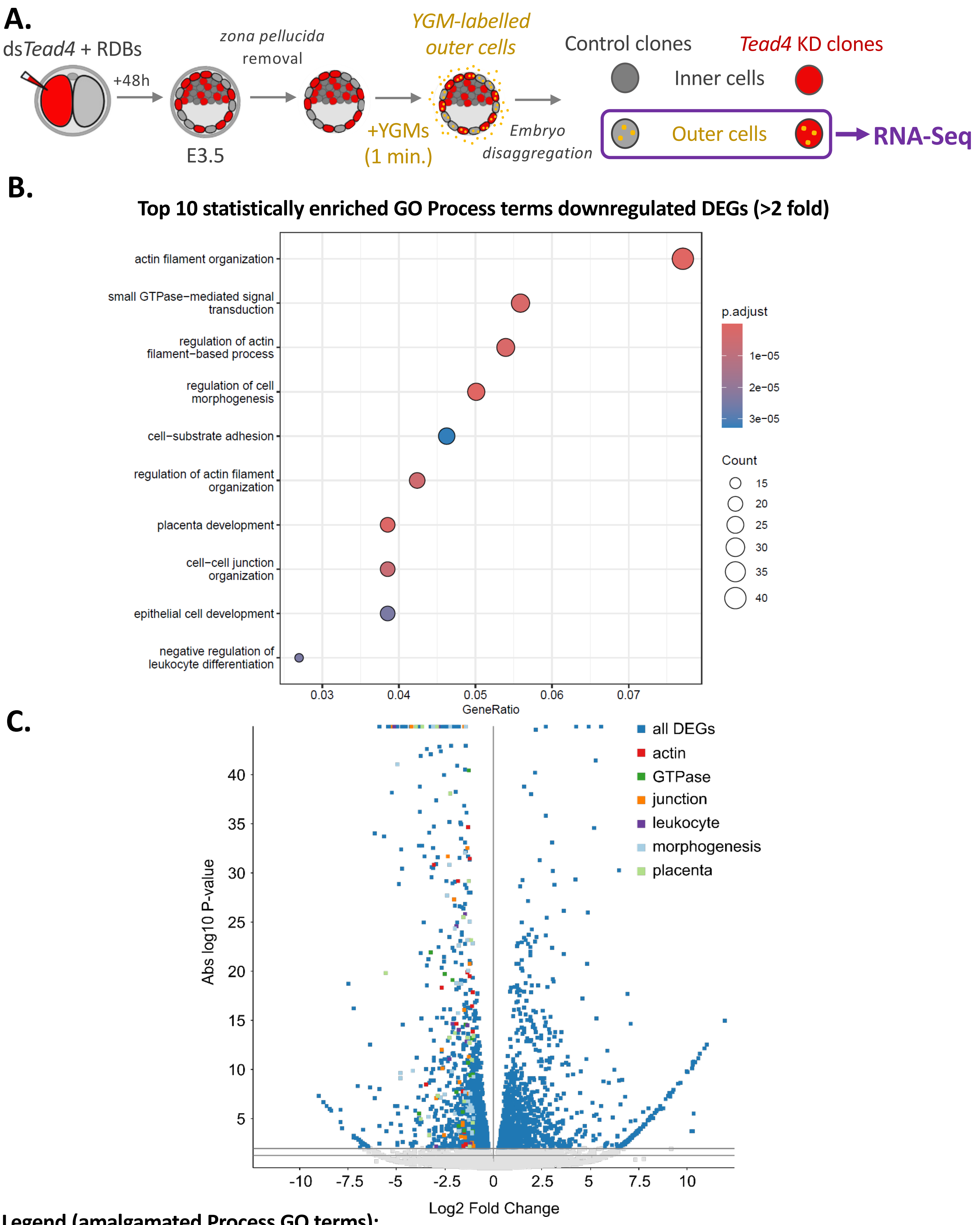

**Legend (amalgamated Process GO terms):**

actin – actin filament organization, regulation of actin filament-based process, regulation of actin filament organization

GTPase – small GTPase-mediated signal transduction

junction – cell-cell juction organization, epithelial cell development

leukocyte – negative regulation of leukocyte differentiation

morphogenesis – regulation of cell morphogenesis, cell-substrate adhesion

placenta – placenta development

**Fig. S9**

**Supplementary Figure S9: Clonal *Tead4* KD-induced downregulated (>2 fold) DEGs enriched GO process terms**

**A)** Schematic of the dsRNA microinjection protocol used to generate E3.5 blastocysts containing *Tead4* KD (ds*Tead4*) clones, marked by co-injection with rhodamine-conjugated dextran beads (RDBs), representing 50% of all cells. At E3.5, outer cells were labelled via endocytosis of yellow-green microspheres (YGMs), followed by full embryo dissociation into single cells, to enable the identification of inner and outer cell populations from marked and unmarked clones, and preparation of RNA-Seq libraries to compare outer cell transcriptomes.

**B)** Dot plots describing the top ten enriched gene ontology (GO) terms, classified by process, for DEGs significantly downregulated (>2 fold and RPKM >0.5 in at least one of the clones) between marked *Tead4* KD versus unmarked control outer clones at the E3.5 stage after RNA-Seq.

**C)** Volcano plot comparing gene expression changes between RNA-Seq derived transcriptomes of E3.5 stage marked *Tead4* KD and unmarked control outer clone cell populations (all DEGs), highlighting significantly enriched GO process terms associated with genes >2 fold downregulated expression and RPKM >0.5 in at least one of the clones (note, related GO process terms were merged, according to the provided lower legend, to aid interpretation).

*S.Tab. 13 provides a summary of all the significantly enriched GO process terms and associated downregulated DEGs.*

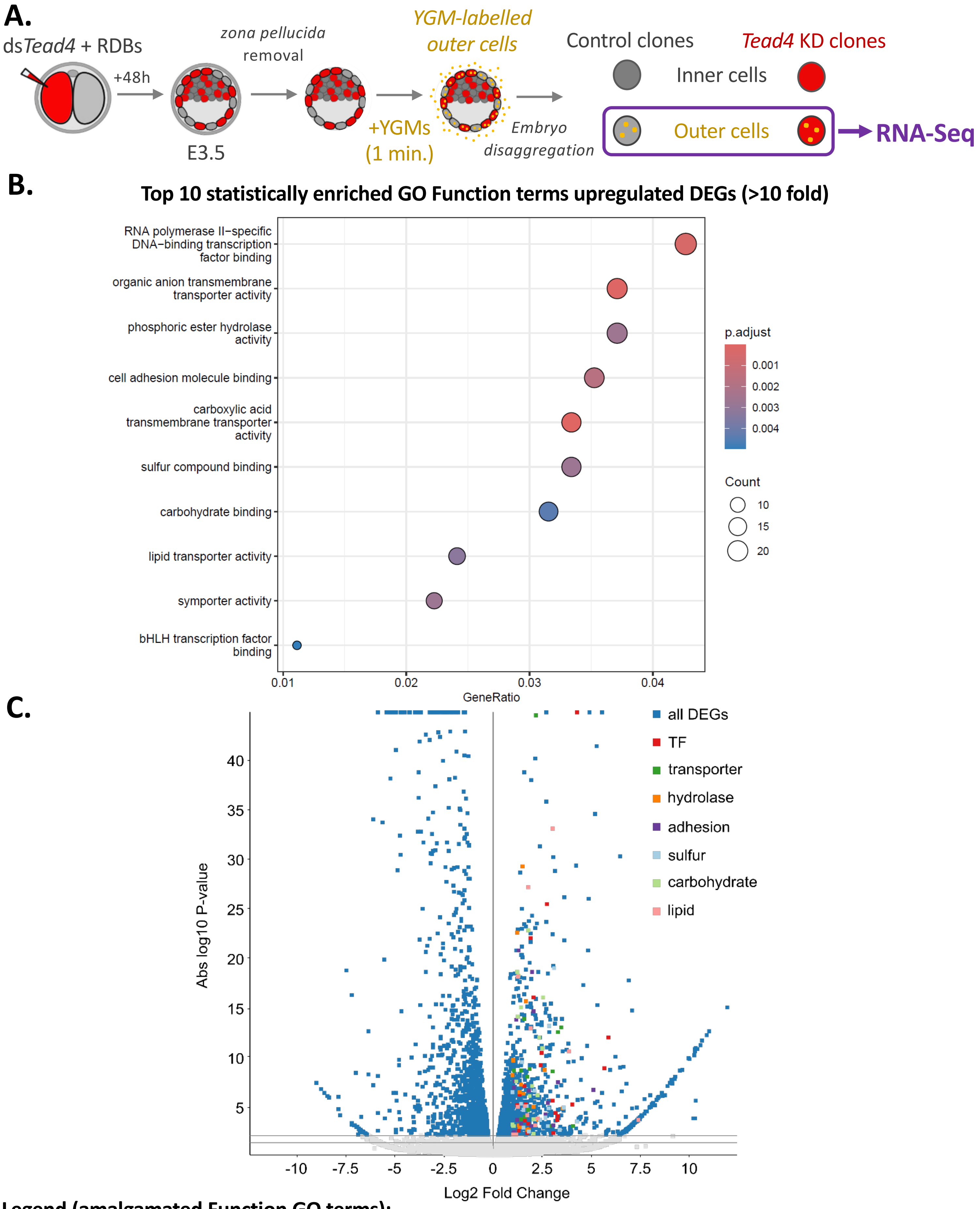

**Legend (amalgamated Function GO terms):**

TF – RNA polymerase II-specific DNA-binding transcription factor binding, bHLH transcription factor binding

transporter – organic anion transmembrane transporter activity, carboxylic acid transmembrane transporter activity, symporter activity

hydrolase – phosphoric ester hydrolase activity

adhesion – cell adhesion molecule binding

sulfur – sulfur compound binding

carbohydrate – carbohydrate binding

lipid – lipid transporter activity

**Fig. S10**

**Supplementary Figure S10: Clonal *Tead4* KD-induced upregulated (>2 fold) DEGs enriched GO functional terms**

**A)** Schematic of the dsRNA microinjection protocol used to generate E3.5 blastocysts containing *Tead4* KD (ds*Tead4*) clones, marked by co-injection with rhodamine-conjugated dextran beads (RDBs), representing 50% of all cells. At E3.5, outer cells were labelled via endocytosis of yellow-green microspheres (YGMs), followed by full embryo dissociation into single cells, to enable the identification of inner and outer cell populations from marked and unmarked clones, and preparation of RNA-Seq libraries to compare outer cell transcriptomes.

**B)** Dot plots describing the top ten enriched gene ontology (GO) terms, classified by function, for DEGs significantly upregulated (>2 fold and RPKM >0.5 in at least one of the clones) between marked *Tead4* KD versus unmarked control outer clones at the E3.5 stage after RNA-Seq.

**C)** Volcano plot comparing gene expression changes between RNA-Seq derived transcriptomes of E3.5 stage marked *Tead4* KD and unmarked control outer clone cell populations (all DEGs), highlighting significantly enriched GO function terms associated with genes >2 fold upregulated expression and RPKM >0.5 in at least one of the clones (note, related GO function terms were merged, according to the provided lower legend, to aid interpretation).

*S.Tab. 14 provides a summary of all the significantly enriched GO function terms and associated upregulated DEGs.*

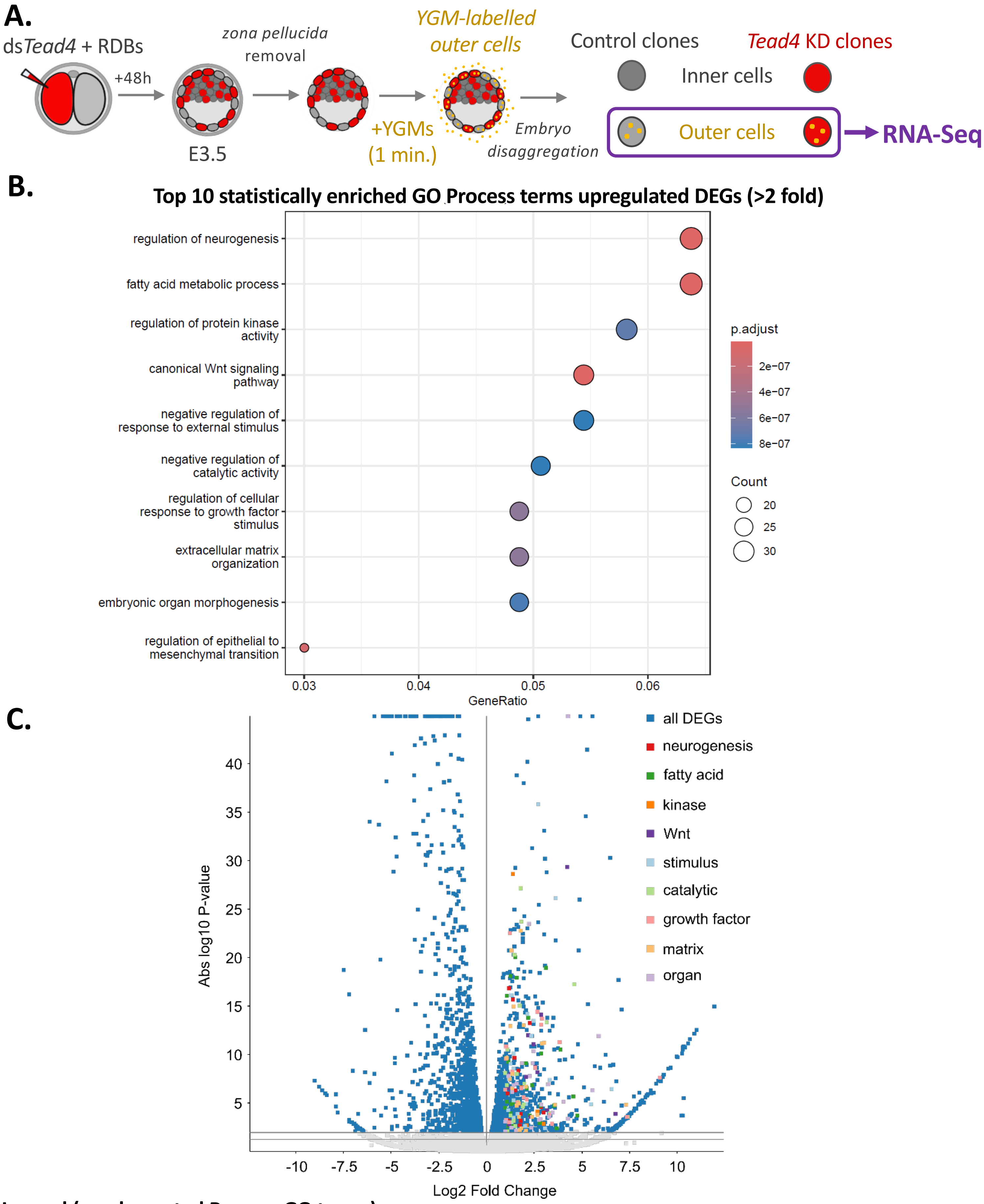

**Legend (amalgamated Process GO terms):**

neurogenesis – regulation of neurogenesis
fatty acid – fatty acid metabolic process
kinase – regulation of protein kinase activity
Wnt – canonical Wnt signaling pathway, regulation of epithelial to mesenchymal transition
stimulus – negative regulation of response to external stimulus
catalytic – negative regulation of catalytic activity
growth factor – regulation of cellular response to growth factor stimulus
matrix – extracellular matrix organization
organ – embryonic organ morphogenesis

**Fig. S11**

**Supplementary Figure S11: Clonal *Tead4* KD-induced upregulated (>2 fold) DEGs enriched GO process terms**

**A)** Schematic of the dsRNA microinjection protocol used to generate E3.5 blastocysts containing *Tead4* KD (ds*Tead4*) clones, marked by co-injection with rhodamine-conjugated dextran beads (RDBs), representing 50% of all cells. At E3.5, outer cells were labelled via endocytosis of yellow-green microspheres (YGMs), followed by full embryo dissociation into single cells, to enable the identification of inner and outer cell populations from marked and unmarked clones, and preparation of RNA-Seq libraries to compare outer cell transcriptomes.

**B)** Dot plots describing the top ten enriched gene ontology (GO) terms, classified by process, for DEGs significantly upregulated (>2 fold and RPKM >0.5 in at least one of the clones) between marked *Tead4* KD versus unmarked control outer clones at the E3.5 stage after RNA-Seq.

**C)** Volcano plot comparing gene expression changes between RNA-Seq derived transcriptomes of E3.5 stage marked *Tead4* KD and unmarked control outer clone cell populations (all DEGs), highlighting significantly enriched GO process terms associated with genes >2 fold upregulated expression and RPKM >0.5 in at least one of the clones (note, related GO process terms were merged, according to the provided lower legend, to aid interpretation).

*S.Tab. 15 provides a summary of all the significantly enriched GO process terms and associated upregulated DEGs.*

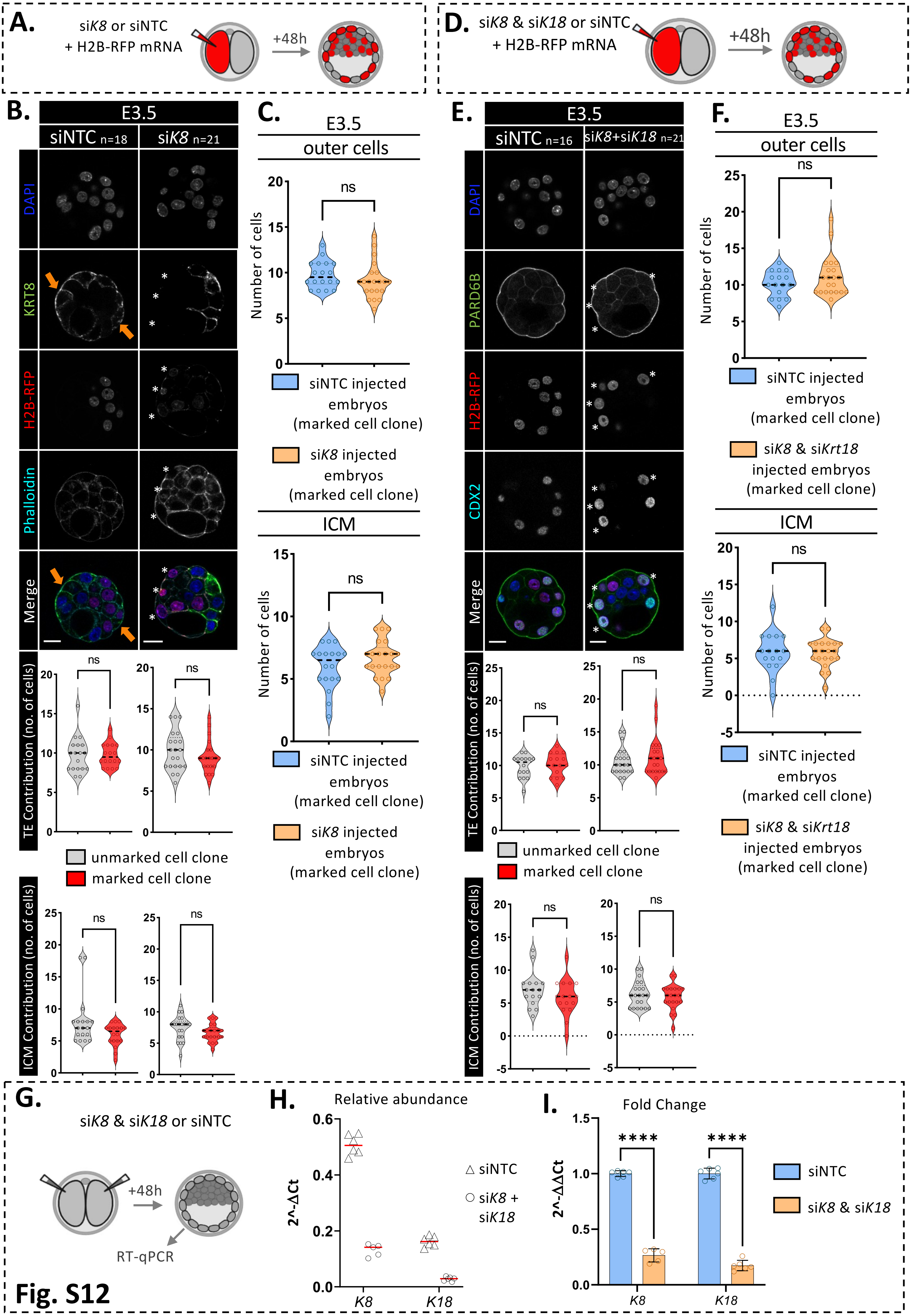
A.
siK8 or siNTC
+ H2B-RFP mRNA
+48h
D.
siK8 & siK18 or siNTC
+ H2B-RFP mRNA
+48h
B.
E3.5
siNTC n=18
siK8 n=21
DAPI
KRT8
H2B-RFP
Phalloidin
Merge
TE Contribution (no. of cells)
ns
unmarked cell clone
marked cell clone
ICM Contribution (no. of cells)
C.
E3.5
outer cells
Number of cells
siNTC injected embryos (marked cell clone)
siK8 injected embryos (marked cell clone)
ICM
E.
E3.5
siNTC n=16
siK8+siK18 n=21
PARD6B
CDX2
F.
siK8 & siKrt18 injected embryos (marked cell clone)
G.
siK8 & siK18 or siNTC
RT-qPCR
H.
Relative abundance
2^-ΔCt
siNTC
siK8 + siK18
K8
K18
I.
Fold Change
2^-ΔΔCt
****
siK8 & siK18
Fig. S12

**Supplementary Figure S12: Clonal KD of *Krt8* or combined *Krt8* and *Krt18* does not mimic *Tead4* KD-induced outer/TE-to-ICM cell allocations in blastocysts**

**A)** Schematic of siRNA microinjection procedure generating E3.5 stage early-blastocysts containing either control (siNTC) or *Krt8* KD (si*K8*) marked clones-comprising 50% of all cells; allowing assessment of clone contributions to the outer TE or ICM positions, in the absence of intermediate filament formation (as also impaired by clonal *Tead4* KD; Fig. 5A-C).

**B)** Confocal micrographs (single channel greyscale z-sections, plus merge – upper panels) of IF-stained control siNTC- and si*K8*-injected blastocysts. Scale bars = 20 µm; number of embryos per group (n) indicated. Orange arrows mark KRT8-positive intermediate filaments restricted to outer cell populations in siNTC embryos (irrespective of clone origin; also seen in unmarked outer clones of the experimental group). White asterisks indicate successful KRT8 KD in marked outer siK8 clones (which still express phalloidin-stained actin). Lower panels show violin plots of unmarked (grey) and H2B-RFP–marked (red) cells allocated to outer TE (top) and ICM (bottom) in siNTC (left) and siK8 (right) embryos. Dashed lines show medians. No significant differences (ns) in clone contribution between unmarked and marked cells in either group was observed; significance marked as *$p<0.05$.

**C)** Comparison of TE (top) and ICM (bottom) cell contributions for only the marked siNTC (blue) versus si*K8* (orange) clones in E3.5 blastocysts. Medians (dashed lines); no significant inter-group differences (ns) observed; *$p<0.05$.

**D)** Schematic of a similar siRNA-microinjection procedure used in panel A) to co-target *Krt8* and *Krt18* (si*K8* & si*K18*) for clonal KD, to assess lineage contribution at E3.5.

**E)** As in B) but for combined *Krt8* and *Krt18* KD versus siNTC controls. Here, KRT8 IF is replaced by the polarity marker PARD6B (green) and CDX2 IF (cyan) as a TE marker, with H2B-RFP marked outer si*K8*+*K18* clones indicated with white asterisks (exhibiting intact apical polarity/PARD6B and CDX2 expression). Lower violin plots show no significant differences (ns) in marked versus unmarked contributions to outer/TE or ICM cells in either group.

**F)** As in C) but for combined *Krt8* and *Krt18* KD; no significant differences (ns) in marked clone contributions to outer/TE or ICM cell between groups were observed (*$p<0.05$).

**G)** RT-qPCR schema: Both blastomeres of 2-cell embryos were microinjected with either control siNTC or si*K8* & si*K18* to induce global *Krt8* and *Krt18* KD, and cultured to the E3.5 blastocyst stage. Normalised *Krt8* and *Krt18* mRNA levels were measured by RT-qPCR to confirm KD efficiency (see S.Tabs. 16).

**H)** Normalised relative expression of *Krt8* (*K8*) and *Krt18* (*K18*) mRNA in siNTC- (triangles) and combined si*K8* plus si*K18*- (circles) treated E3.5 blastocysts. Medians shown by red bars; two biological replicates with three technical replicates each.

**I)** Median fold changes of *Krt8* (*K8*) and *Krt18* (*K18*) mRNA levels for combined si*K8* plus si*Krt18*-versus siNTC-treated embryos (orange vs. blue); based on data in panel H). Statistical significance (*$p<0.05$) and standard deviations (error bars) are shown.

*All raw clonal allocation data are summarised in S.Tabs. 18a & 18b. Statistical analyses were performed based on determined data distribution (normal or non-normal) and application of appropriate tests; details in supplementary statistics Excel workbook.*

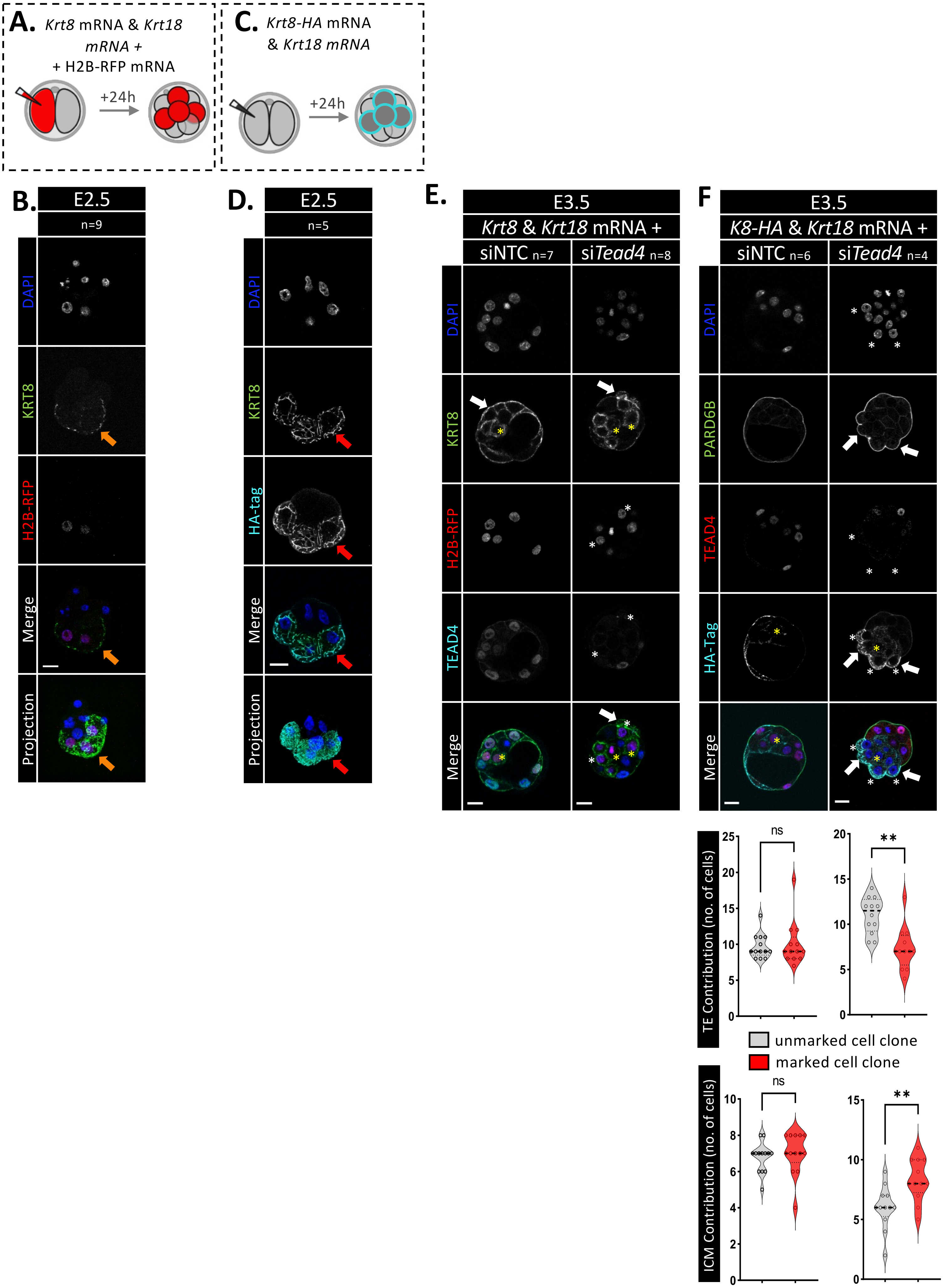
A. Krt8 mRNA & Krt18 mRNA + + H2B-RFP mRNA
+24h
C. Krt8-HA mRNA & Krt18 mRNA
+24h
B. E2.5
n=9
DAPI
KRT8
H2B-RFP
Merge
Projection
D. E2.5
n=5
DAPI
KRT8
HA-tag
Merge
Projection
E. E3.5
Krt8 & Krt18 mRNA +
siNTC n=7
siTead4 n=8
DAPI
KRT8
H2B-RFP
TEAD4
Merge
F E3.5
K8-HA & Krt18 mRNA +
siNTC n=6
siTead4 n=4
DAPI
PARD6B
TEAD4
HA-Tag
Merge
TE Contribution (no. of cells)
ns
**
unmarked cell clone
marked cell clone
ICM Contribution (no. of cells)
ns
**

Fig. S13

**Supplementary Figure S13: Expression of recombinant *Krt8* (including HA-tagged variant) and *Krt18* mRNAs in preimplantation mouse embryos**

**A)** Schematic of the microinjection strategy to express recombinant mouse *Krt8* and *Krt18* mRNAs in marked clones (50% of all cells) by single-blastomere microinjections at the 2-cell stage, followed by 24 h culture to the 8-cell (E2.5) stage; enabling assessment of precocious, clone-specific intermediate filament expression and potential phenotypic rescue of clonal *Tead4* KD–induced outer/TE-to-ICM cell allocations (Fig. 5D–F; plus the accompanying panels E) and F)).

**B)** Example confocal micrographs (single channel-greyscale z-sections, plus merges and projections) of IF-stained 8-cell (E2.5) stage embryos microinjected as in A). Scale bars, 20µm; n indicated. Orange arrows KRT8-positive intermediate filaments restricted to the H2B-RFP–marked (red) clone, confirming precocious recombinant intermediate-filament expression versus the unmarked clone.

**C)** Parallel schematic to A), substituting recombinant *Krt8* mRNA with an N-terminal HA-tagged version (*Krt8*-HA) to unequivocally identify recombinant intermediate filaments by HA-specific IF (cyan).

**D)** As in B), but showing marked clone-specific recombinant KRT8 expression (red arrows) in 8-cell (E2.5) embryos treated as in C), detected by both anti-KRT8 (green) and anti-HA IF (cyan; used to identify the microinjected clones, in the absence of H2B-RFP).

**E)** Confocal micrographs (single channel greyscale z-sections) of E3.5 blastocysts from Fig. 5E containing H2B-RFP marked (red) siNTC (left) or si*Tead4* (right) clones that also over-express co-microinjected recombinant *Krt8* and *Krt18* mRNA. Embryos are IF stained for KRT8 (green) and TEAD4 (cyan) with DNA-DAPI counterstain (blue). Scale bar, 20µm. n indicated. KRT8 expression is observed in both outer-residing clones of siNTC- and si*Tead4*-treated embryos (but stronger in marked clones; indicated with white arrows). Yellow asterisks indicate recombinant KRT8 expression and intermediate filament formation in marked inner-cell clones from both groups (not usual for endogenous KRT8). White asterisks denote confirmed TEAD4 KD in marked si*Tead4* clones associated with recombinant KRT8 expression.

**F)** Upper panels; as in E) showing E3.5 embryos containing siNTC or si*Tead4* clones but co-injected with recombinant HA-tagged *Krt8* mRNA (*Krt8*-HA), rather than untagged *Krt8* transcripts (but still in combination with *Krt18* mRNA). Embryos are IF-stained for PARD6B (green), TEAD4 (red) and the HA tag (cyan; to identify the microinjected clones, in the absence of a H2B-RFP marker). Asterisks denote the progeny of the 2-cell injections, marked by KRT8-HA expression (white for outer cells, yellow for inner cells), which are also TEAD4-negative in si*Tead4*-treated (KD) embryos. Despite KRT8-HA expression, outer clones in the si*Tead4*-treted group still form apical blebs (white arrows) and injected clones from both groups populate inner positions (yellow asterisks). Lower panels: composite violin plots showing the distribution of unmarked (grey) versus H2B-RFP/KRT8-HA-marked (red) clones in siNTC- (left) or si*Tead4*- (right) treated groups (also co-injected with *Krt8* (or Krt18-HA) and *Krt18* mRNA), allocated to outer TE (top) and ICM (bottom) positions. Dashed lines indicate medians, significant differences are indicated (*$p<0.05$). There were no significant differences (ns) in outer-TE or ICM clone contribution between unmarked and marked cells in siNTC embryos, and recombinant intermediate filament expression did not alter the *Tead4* KD-driven outer/TE-to-ICM cell allocations (i.e. phenotypic rescue was not achieved).

*Raw clonal allocation data for panel F) are summarised in S.Tabs. 19. Statistical analyses were performed based on determined data distribution (normal or non-normal) and application of appropriate tests; details in supplementary statistics Excel workbook.*

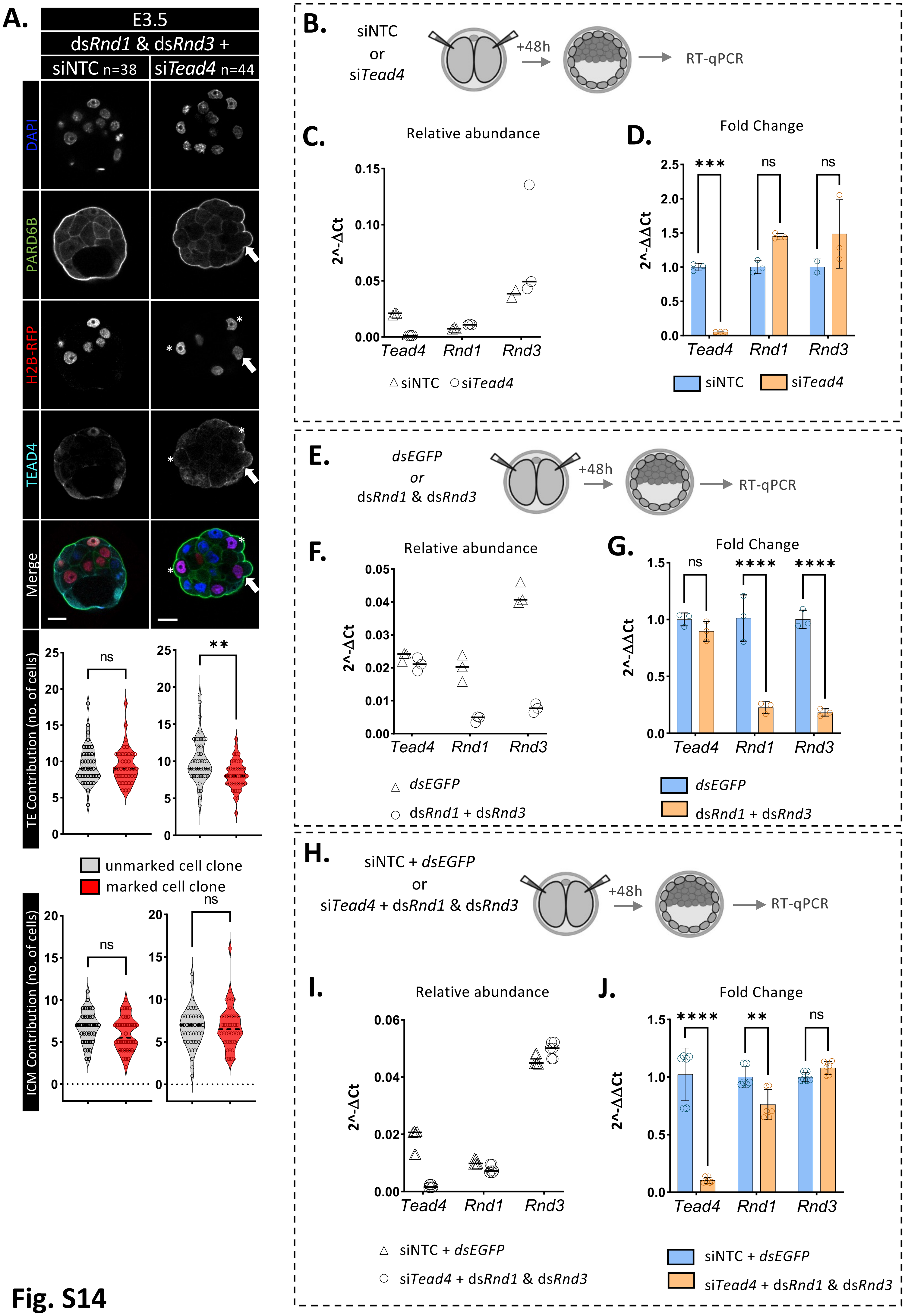

A.
E3.5
dsRnd1 & dsRnd3 +
siNTC n=38
siTead4 n=44
DAPI
PARD6B
H2B-RFP
TEAD4
Merge
TE Contribution (no. of cells)
ICM Contribution (no. of cells)
unmarked cell clone
marked cell clone
ns
**
B.
siNTC
or
siTead4
+48h
RT-qPCR
C.
Relative abundance
2^-ΔCt
Tead4
Rnd1
Rnd3
△ siNTC ○ siTead4
D.
Fold Change
2^-ΔΔCt
***
siNTC
siTead4
E.
dsEGFP
or
dsRnd1 & dsRnd3
F.
G.
****
dsRnd1 + dsRnd3
H.
siNTC + dsEGFP
siTead4 + dsRnd1 & dsRnd3
I.
J.

Fig. S14

**Supplementary Figure S14: Confirmed clonal KD of *Rnd1* and *Rnd3* mRNA partially rescues *Tead4* KD–induced outer/TE-to-ICM cell allocations at the E3.5 blastocyst stage**

**A)** Upper panels: confocal micrographs (single-channel greyscale z-sections with merges) of E3.5 blastocysts from Fig. 5H, showing H2B-RFP–marked clones (red) in siNTC- (left) or si*Tead4*- (right) treatment groups that were co-microinjected with control dsEGFP or ds*Rnd1* & ds*Rnd3* constructs, respectively. Embryos stained for PARD6B (green) and TEAD4 (cyan) with DNA-DAPI counterstain (blue). Scale bar, 20μm. The number (n) of embryos per group is indicated. White asterisks indicate outer si*Tead4* clones with example persistent apical blebs (white arrows). Lower panels: violin plots of unmarked (grey) vs H2B-RFP–marked (red) cells for control siNTC/dsEGFP (left) or si*Tead4*/ds*Rnd1* & ds*Rnd3* (right) conditions, allocated to outer TE (top) and ICM (bottom) positions. Dashed lines indicate medians; *p<0.05. There was no significant difference (ns) in TE/ICM contributions between unmarked and marked cells in control embryos. However, *Tead4* KD-driven increases in ICM contribution (via the marked clone, see Figs. 1F and S1B) are rescued by *Rnd1*/*Rnd3* KD (as no significant difference from the unmarked clone is observed), though apical distortions persist, indicating partial rescue.

**B)** RT-qPCR schematic: both blastomeres of 2-cell embryos are microinjected with either control siNTC or si*Tead4* to induce global *Tead4* KD and cultured to the E3.5stage; normalised *Tead4*, *Rnd1* and *Rnd3* mRNA levels were measured to confirm *Tead4*-KD-associated *Rnd1* and *Rnd3* induction observed in RNA-Seq (Fig. 4) - data summarised in S.Tabs.16).

**C)** Normalised expression of *Tead4*, *Rnd1* and *Rnd3* in siNTC (triangles) and *siTead4* (circles) treated E3.5 blastocysts. Medians shown; three technical replicates.

**D)** Median fold changes for si*Tead4* versus s*i*NTC treatment groups, based on panel C) data. Statistical significance (*p<0.05) and standard deviations (error bars) shown.

**E)–G)** As in panels B)–D), but comparing mRNA expression levels in control dsEGFP and ds*Rnd1* plus ds*Rnd3* microinjected embryos; confirming efficient/significant *Rnd1* and *Rnd3* gene KD (without significantly affecting *Tead4* derived transcript levels).

**H)–J)** As in panels B)–D) and E–G), but comparing mRNA expression levels in control siNTC/dsEGFP and experimental si*Tead4*/ds*Rnd1* plus ds*Rnd3* microinjected embryos; i.e. relating to attempted clonal si*Tead4*-associated outer/TE-to-ICM blastocyst cell allocation phenotypic rescue – Fig. 5G-I and panel A), above). Note, *Rnd1* and *Rnd3* mRNA levels, normally induced after *Tead4* KD (Fig. 4 and panels B)–D), above), are reduced to levels more equivalent to those observed in controls (when targeting each gene alone; panels E)–G), above), but the KD is sufficient to mediate a partial phenotypic rescue (Fig 5I and panel A), above).

*Raw clonal allocation data, panel F, are summarised in S.Tabs. 20. Statistical analyses were performed based on determined data distribution (normal or non-normal) and application of appropriate tests; details in supplementary statistics Excel workbook.*

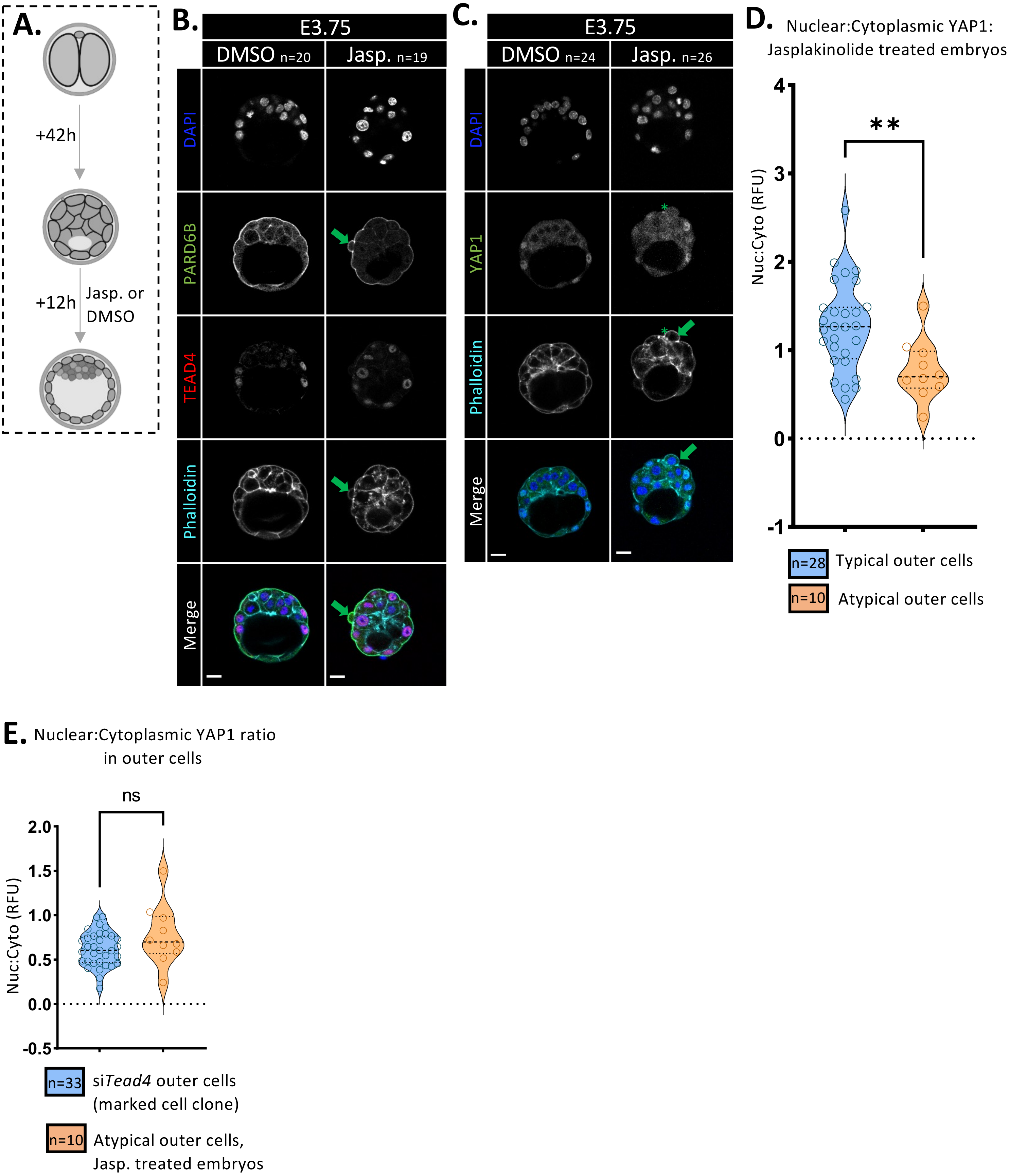
A.
+42h
+12h
Jasp. or
DMSO
B.
E3.75
DMSO n=20
Jasp. n=19
DAPI
PARD6B
TEAD4
Phalloidin
Merge
C.
E3.75
DMSO n=24
Jasp. n=26
DAPI
YAP1
Phalloidin
Merge
D.
Nuclear:Cytoplasmic YAP1:
Jasplakinolide treated embryos
Nuc:Cyto (RFU)
**
4
3
2
1
0
-1
n=28 Typical outer cells
n=10 Atypical outer cells
E.
Nuclear:Cytoplasmic YAP1 ratio
in outer cells
ns
Nuc:Cyto (RFU)
2.0
1.5
1.0
0.5
0.0
-0.5
n=33 siTead4 outer cells
(marked cell clone)
n=10 Atypical outer cells,
Jasp. treated embryos

**Fig. S15**

**Supplementary Figure S15: Jasplakinolide stabilisation of early-blastocyst actin filaments phenocopies apical-domain distortions and reduced YAP1 nuclear localisation associated with clonal *Tead4* KD**

**A)** Schematic of embryo drug treatment: 2-cell (E1.5) embryos cultured for 42 hours to the early (E3.5) blastocyst stage were exposed to a further 12 hour culture in Jasplakinolide (Jasp., an actin filament inducer and stabiliser) or DMSO vehicle control containing media; embryos, at the equivalent early/mid-blastocyst (E3.75) stage, were then fixed for confocal IF-based morphological examination.

**B)** Example confocal micrographs (single greyscale z-sections with merges) of DMSO- and Jasp.-treated E3.75 blastocysts, IF-stained for PARD6B (green), TEAD4 (red), actin (Phalloidin; cyan), and DNA (DAPI; blue). Scale bar = 20 µm. The number of embryos (n) per treatment group is indicated. Green arrows denote apical-domain distortions reminiscent of clonal *Tead4* KD–induced apical blebs (as in Fig. 1C), specifically in Jasp.-treated embryos.

**C)** As in B), but IF-stained for YAP1 (green), omitting TEAD4 staining, to assess expression and subcellular YAP1 localisation. Green arrow indicates a Jasp.-treatment–specific apical bleb structure, associated with an outer cell showing atypical cytoplasmic and depleted nuclear YAP1 expression (green asterisk), reminiscent of marked outer *siTead4* clones (Fig. 3A).

**D)** Quantified single outer-cell nuclear:cytoplasmic YAP1 expression ratios (RFU) in Jasplakinolide-treated E3.75 blastocysts, categorised by presence or absence of apical-domain blebs (i.e. typical versus atypical outer cells). Medians indicated with dashed lines; significant differences (*p<0.05) indicated. n denotes cell sample sizes. Note the reduction in nuclear YAP1 accumulation in outer cells with Jasplakinolide-induced apical blebs.

**E)** Comparison of outer-cell nuclear:cytoplasmic YAP1 RFU ratios in atypical outer cells (with apical blebs) induced by Jasplakinolide- (Jasp.) treatment versus outer-residing marked si*Tead4* clones (as described for Fig. 1A) at the E3.75 stage. n denotes sample size. Medians indicated with dashed lines; no significant difference (ns) observed between groups (*p<0.05), indicating equivalence in the induced phenotypes. Raw data for YAP1 expression (RFU) siTead4 outer cells is seen in *S.Tabs S6a.*

*Quantified RFU values used to generate panels D) and E) are summarised in S.Tabs. 21a-21d. Statistical analyses were performed according to normality and experimental design. Parametric tests were applied to normally distributed data. Unpaired t-tests were applied to between group analysis. Details of t-tests and corresponding p-values are provided in the supplementary statistics Excel workbook.*

### Supplementary Movie 1 legend

**Supplementary movie 1: 4D time-lapse light-sheet live embryo imaging (*related to Fig. 2*)**
Projected illustrative examples of transgenic membrane-Tomato/mT+ blastocysts, containing either histone H2B-Venus marked control siNTC- (left) or si*Tead4*-microinjected (*Tead4* KD – middle corresponding to si*Tead4* #1 and right relating to si*Tead4*#2 embryo examples) clones (after single blastomere microinjection at the 2-cell stage), during 16.5 hours of time-lapse light-sheet microscopy imaging (10-minute frame intervals). Merged fluorescent channels are shown in the upper panels and brightfield (of the equatorial z-section) in the lower panels. Developmental timings (hh:mm) relative to post-hCG superovulation microinjections and scale bar (10μm) are indicated. The number of TE and ICM-marked (Histone H2B-Venus+) cells at the end of the imaging period are indicated. Pseudo-colour scheme: magenta – transgenic embryo mT+ reporter (standard deviation projection), and green – histone H2B-Venus microinjection/clone marker (maximum projection). In the central si*Tead4*-related movie, incidences of atypical blastocyst cell internalisation events are highlighted by yellow dots. Supplementary Movie Figure SM1, provides extracted frames from the start, mid, and end-points of the imaging period (for the merged fluorescent channels). Relates to data in Figs. 2 and S4 & S5.

**Supplementary Movie Figure SM1 and legend**

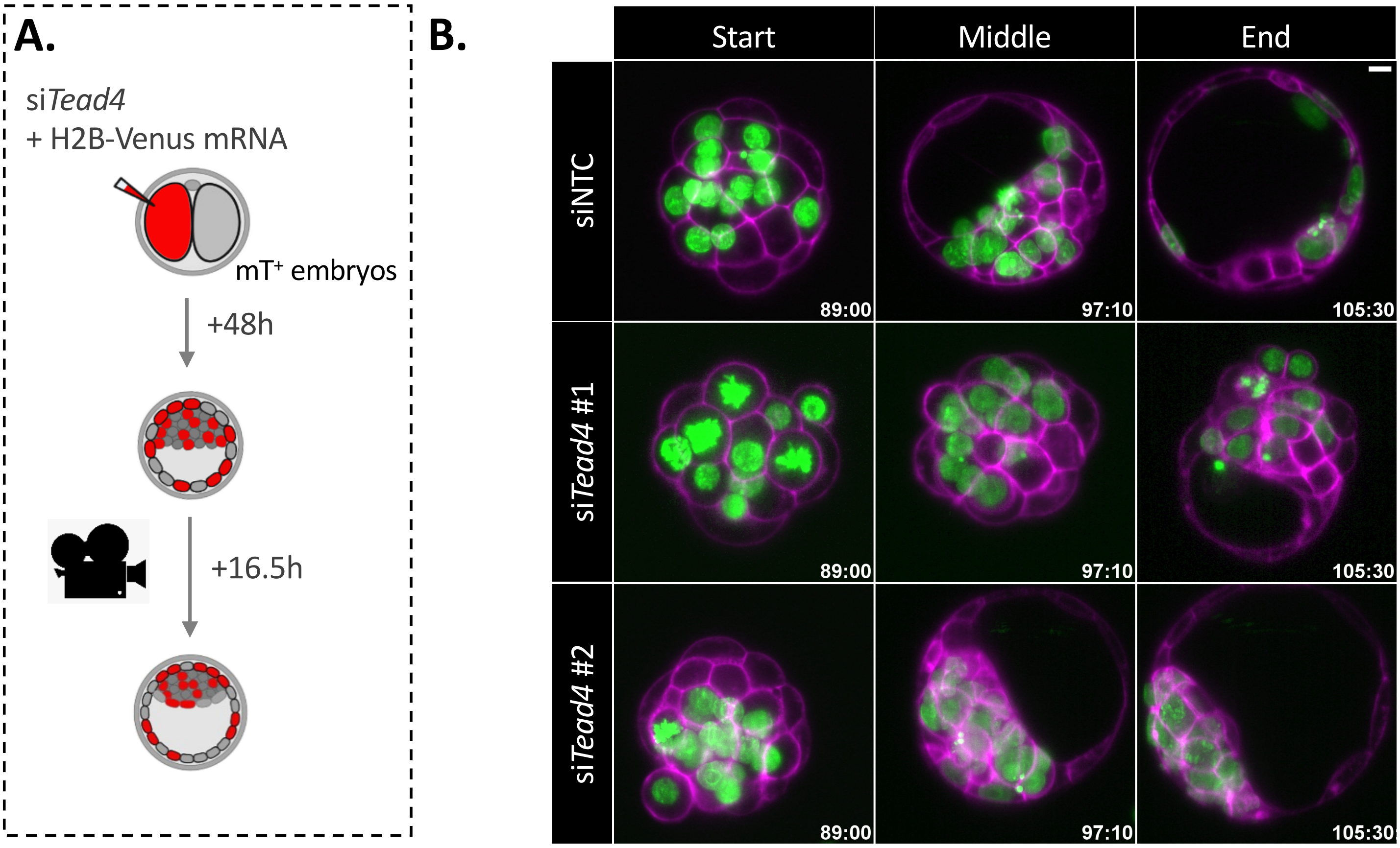

**Supplementary Movie Fig. SM1**

**Supplementary Movie Figure SM1. Exemplar micrographs of embryos subject to light-sheet microscopy live-embryo imaging (*related to supplementary movie 1*)**

**A)** Experimental schematic of 2-cell stage microinjections of transgenic membrane-Tomato expressing ($mT^+$) 2-cell embryos used to generate early 32-cell stage blastocysts (after 48 hours of culture) containing either histone H2B-Venus marked control (siNTC) or *Tead4* KD (si*Tead4*) clones (comprising 50% of all cells), subject to 16.5 hours of light-sheet live-cell embryo imaging (10 minute intervals); allowing dynamic assessment of individual clone contributions to the outer/TE or ICM positions.

**B)** Exemplar projections of time-lapse imaged control (siNTC) or *Tead4* KD (si*Tead*4 – x2) microinjected embryos (as depicted in supplementary movie 1) at the start, mid, and end-points of imaging (time post-hCG treatment indicated as hh:mm). Pseudo-colour scheme: magenta – transgenic embryo mT reporter (comprising a single equatorial z-section), and green – histone H2B-Venus microinjection/clone marker (as a maximum projection from several planes located above and below the selected mT plane); scale bar – 10μm.

### Supplementary Tables and legends (S.Tabs 1-24)

**Supplementary Tables 1a: Lineage Contribution Overview and Statistics (Clonal *Tead4* KD)**

| | Whole embryo (combined lineage) contribution statistics | | | | | | | | | | | | | | | |
|---|---|---|---|---|---|---|---|---|---|---|---|---|---|---|---|---|
| | +48h (E3.5) | | | | +54h (E3.75) | | | | +60h (E4.0) | | | | +72h (E4.5) | | | |
| | siNTC | | siTead4 | | siNTC | | siTead4 | | siNTC | | siTead4 | | siNTC | | siTead4 | |
| **n** | 15 | | 19 | | 26 | | 17 | | 19 | | 10 | | 15 | | 19 | |
| **Av. Total cell no.** | 31.00 | | 30.89 | | 44.50 | | 46.18 | | 63.68 | | 62.30 | | 85.53 | | 68.47 | |
| **Inj. status** | Unmarked | Marked | Unmarked | Marked | Unmarked | Marked | Unmarked | Marked | Unmarked | Marked | Unmarked | Marked | Unmarked | Marked | Unmarked | Marked |
| **Mean** | 15.60 | 15.40 | 16.21 | 14.68 | 22.73 | 21.77 | 24.47 | 21.71 | 32.63 | 31.05 | 35.40 | 26.90 | 42.53 | 43.00 | 49.74 | 18.74 |
| **Mean % of total** | 0.50 | 0.50 | 0.52 | 0.48 | 0.51 | 0.49 | 0.53 | 0.47 | 0.51 | 0.49 | 0.57 | 0.43 | 0.50 | 0.50 | 0.71 | 0.29 |
| **Sd** | 3.76 | 2.59 | 5.04 | 2.06 | 5.17 | 5.30 | 5.41 | 4.18 | 5.85 | 5.83 | 6.98 | 8.81 | 8.78 | 7.93 | 15.47 | 5.14 |
| **Median** | 15.00 | 16.00 | 16.00 | 16.00 | 23.00 | 20.50 | 24.00 | 23.00 | 31.00 | 31.00 | 36.50 | 24.50 | 42.00 | 44.00 | 51.00 | 18.00 |
| **Median % of total cells** | 0.50 | 0.50 | 0.52 | 0.48 | 0.50 | 0.50 | 0.53 | 0.47 | 0.52 | 0.48 | 0.59 | 0.41 | 0.51 | 0.49 | 0.74 | 0.26 |
| **Q1** | 14.00 | 15.00 | 14.00 | 14.00 | 17.75 | 17.00 | 21.50 | 17.50 | 28.00 | 27.00 | 30.50 | 20.75 | 38.00 | 36.00 | 37.00 | 14.00 |
| **Q3** | 16.00 | 17.00 | 17.00 | 16.00 | 27.25 | 26.50 | 29.00 | 24.00 | 37.00 | 37.00 | 40.00 | 30.00 | 48.00 | 46.00 | 56.00 | 25.00 |

| | TE contribution statistics | | | | | | | | | | | | | | | |
|---|---|---|---|---|---|---|---|---|---|---|---|---|---|---|---|---|
| | +48h (E3.5) | | | | +54h (E3.75) | | | | +60h (E4.0) | | | | +72h (E4.5) | | | |
| | siNTC | | si*Tead4* | | siNTC | | si*Tead4* | | siNTC | | si*Tead4* | | siNTC | | si*Tead4* | |
| **n** | 15 | | 19 | | 26 | | 17 | | 19 | | 10 | | 15 | | 19 | |
| **Av. no.TE cells** | 19.07 | | 18.11 | | 28.42 | | 22.65 | | 43.53 | | 36.60 | | 65.60 | | 44.05 | |
| **Inj. status** | Unmarked | Marked | Unmarked | Marked | Unmarked | Marked | Unmarked | Marked | Unmarked | Marked | Unmarked | Marked | Unmarked | Marked | Unmarked | Marked |
| **Mean** | 9.47 | 9.60 | 10.63 | 7.47 | 14.35 | 14.08 | 14.88 | 7.76 | 22.89 | 20.63 | 27.50 | 9.10 | 32.53 | 33.07 | 39.26 | 4.79 |
| **Mean % of total** | 0.50 | 0.50 | 0.57 | 0.43 | 0.50 | 0.50 | 0.65 | 0.35 | 0.53 | 0.47 | 0.75 | 0.25 | 0.50 | 0.50 | 0.88 | 0.12 |
| **Sd** | 2.85 | 2.41 | 4.39 | 1.71 | 4.47 | 3.93 | 3.66 | 1.60 | 4.78 | 4.97 | 5.85 | 4.58 | 6.73 | 6.56 | 14.62 | 4.67 |
| **Median** | 8.00 | 10.00 | 10.00 | 8.00 | 14.50 | 14.00 | 15.00 | 8.00 | 21.00 | 21.00 | 29.00 | 8.50 | 33.00 | 33.00 | 38.00 | 3.00 |
| **Median % of total TE** | 0.48 | 0.52 | 0.56 | 0.44 | 0.50 | 0.50 | 0.68 | 0.32 | 0.53 | 0.47 | 0.76 | 0.24 | 0.49 | 0.51 | 0.92 | 0.08 |
| **Q1** | 8.00 | 8.00 | 8.00 | 6.00 | 10.00 | 10.75 | 13.00 | 7.00 | 19.00 | 16.00 | 25.75 | 6.50 | 30.00 | 28.00 | 32.00 | 1.00 |
| **Q3** | 11.00 | 11.00 | 12.00 | 9.00 | 17.00 | 18.00 | 18.00 | 8.50 | 26.00 | 25.00 | 31.25 | 10.50 | 37.00 | 36.00 | 46.00 | 9.00 |

| | ICM contribution statistics | | | | | | | | | | | | | | | |
|---|---|---|---|---|---|---|---|---|---|---|---|---|---|---|---|---|
| | +48h (E3.5) | | | | +54hr(E3.75) | | | | +60h (E4.0) | | | | +72h (E4.5) | | | |
| | siNTC | | si*Tead4* | | siNTC | | si*Tead4* | | siNTC | | si*Tead4* | | siNTC | | si*Tead4* | |
| **n** | 15 | | 19 | | 26 | | 17 | | 19 | | 10 | | 15 | | 19 | |
| **Av. no. ICM cells** | 11.93 | | 12.79 | | 16.08 | | 23.53 | | 20.16 | | 25.70 | | 19.93 | | 24.42 | |
| **Inj. status** | Unmarked | Marked | Unmarked | Marked | Unmarked | Marked | Unmarked | Marked | Unmarked | Marked | Unmarked | Marked | Unmarked | Marked | Unmarked | Marked |
| **Mean** | 6.13 | 5.80 | 5.58 | 7.21 | 8.38 | 7.69 | 9.59 | 13.94 | 9.74 | 10.42 | 7.90 | 17.80 | 10.00 | 9.93 | 10.47 | 13.95 |
| **Mean % of total** | 0.51 | 0.49 | 0.44 | 0.56 | 0.52 | 0.48 | 0.42 | 0.58 | 0.49 | 0.51 | 0.32 | 0.68 | 0.49 | 0.51 | 0.43 | 0.57 |
| **Sd** | 1.85 | 1.42 | 1.57 | 2.23 | 3.05 | 3.02 | 2.40 | 5.10 | 2.66 | 3.25 | 3.60 | 8.43 | 3.98 | 3.01 | 4.22 | 4.12 |
| **Median** | 6.00 | 6.00 | 6.00 | 7.00 | 8.00 | 7.00 | 10.00 | 15.00 | 10.00 | 10.00 | 8.00 | 16.00 | 10.00 | 10.00 | 10.00 | 14.00 |
| **Median % of total ICM** | 0.53 | 0.47 | 0.42 | 0.58 | 0.53 | 0.47 | 0.41 | 0.59 | 0.50 | 0.50 | 0.31 | 0.69 | 0.48 | 0.52 | 0.38 | 0.62 |
| **Q1** | 5.00 | 5.00 | 5.00 | 6.00 | 6.00 | 6.00 | 7.50 | 10.00 | 8.00 | 9.00 | 4.75 | 12.50 | 7.00 | 8.00 | 8.00 | 10.00 |
| **Q3** | 8.00 | 7.00 | 7.00 | 8.00 | 9.00 | 9.00 | 11.00 | 18.50 | 11.00 | 13.00 | 10.00 | 20.50 | 11.00 | 12.00 | 13.00 | 16.00 |

**Supplementary Tables S1a:** Summary of blastocyst lineage/positional contributions from injected/marked control siNTC and si*Tead4* cell clones, compared with equivalent non-injected/unmarked clones, at +48 h (E3.5), +54 h (E3.75), +60 h (E4.0), and +72 h (E4.5) post-single 2-cell stage blastomere microinjection. Relates to data in Figs. 1 and S1.

**Supplementary Tables 1b: +48h (Clonal *Tead4* KD - E3.5)**

**siNTC**

| Embryo Number | Raw values | | | | | Marked | | Unmarked | | Contribution to lineage (%) | | | | ICM:TE |
|---|---|---|---|---|---|---|---|---|---|---|---|---|---|---|
| | | | | | | | | | | Outer | | Inner | | |
| | Total | Outer | Inner | Marked | Unmarked | Outer | Inner | Outer | Inner | Marked | Unmarked | Marked | Unmarked | ICM:TE |
| _001 | 31 | 19 | 12 | 16 | 15 | 11 | 5 | 8 | 7 | 57.89% | 42.11% | 41.67% | 58.33% | 0.63 |
| _002 | 20 | 13 | 7 | 9 | 11 | 5 | 4 | 8 | 3 | 38.46% | 61.54% | 57.14% | 42.86% | 0.54 |
| _003 | 28 | 14 | 14 | 15 | 13 | 8 | 7 | 6 | 7 | 57.14% | 42.86% | 50.00% | 50.00% | 1.00 |
| _004 | 30 | 18 | 12 | 15 | 15 | 10 | 5 | 8 | 7 | 55.56% | 44.44% | 41.67% | 58.33% | 0.67 |
| _005 | 29 | 14 | 15 | 14 | 15 | 5 | 9 | 9 | 6 | 35.71% | 64.29% | 60.00% | 40.00% | 1.07 |
| _006 | 31 | 18 | 13 | 17 | 14 | 10 | 7 | 8 | 6 | 55.56% | 44.44% | 53.85% | 46.15% | 0.72 |
| _007 | 33 | 18 | 15 | 18 | 15 | 11 | 7 | 7 | 8 | 61.11% | 38.89% | 46.67% | 53.33% | 0.83 |
| _008 | 29 | 21 | 8 | 18 | 11 | 13 | 5 | 8 | 3 | 61.90% | 38.10% | 62.50% | 37.50% | 0.38 |
| _009 | 32 | 21 | 11 | 16 | 16 | 11 | 5 | 10 | 6 | 52.38% | 47.62% | 45.45% | 54.55% | 0.52 |
| _010 | 31 | 17 | 14 | 16 | 15 | 10 | 6 | 7 | 8 | 58.82% | 41.18% | 42.86% | 57.14% | 0.82 |
| _011 | 25 | 17 | 8 | 11 | 14 | 7 | 4 | 10 | 4 | 41.18% | 58.82% | 50.00% | 50.00% | 0.47 |
| _012 | 43 | 28 | 15 | 19 | 24 | 12 | 7 | 16 | 8 | 42.86% | 57.14% | 46.67% | 53.33% | 0.54 |
| _013 | 32 | 23 | 9 | 16 | 16 | 12 | 4 | 11 | 5 | 52.17% | 47.83% | 44.44% | 55.56% | 0.39 |
| _014 | 32 | 21 | 11 | 16 | 16 | 10 | 6 | 11 | 5 | 47.62% | 52.38% | 54.55% | 45.45% | 0.52 |
| _015 | 39 | 24 | 15 | 15 | 24 | 9 | 6 | 15 | 9 | 37.50% | 62.50% | 40.00% | 60.00% | 0.63 |
| Mean | 31.00 | 19.07 | 11.93 | 15.40 | 15.60 | 9.60 | 5.80 | 9.47 | 6.13 | 50.39% | 49.61% | 49.16% | 50.84% | 0.65 |
| Median | 31.00 | 18.00 | 12.00 | 16.00 | 15.00 | 10.00 | 6.00 | 8.00 | 6.00 | 52.38% | 47.62% | 46.67% | 53.33% | 0.63 |
| Sd | 5.28 | 4.06 | 2.84 | 2.59 | 3.76 | 2.41 | 1.42 | 2.85 | 1.85 | 9.08% | 9.08% | 7.05% | 7.05% | 0.21 |
| SEM | 1.36 | 1.05 | 0.73 | 0.67 | 0.97 | 0.62 | 0.37 | 0.74 | 0.48 | 2.34% | 2.34% | 1.82% | 1.82% | 0.05 |

**si*Tead4***

| Embryo Number | Raw values | | | | | Marked | | Unmarked | | Contribution to lineage (%) | | | | ICM:TE |
|---|---|---|---|---|---|---|---|---|---|---|---|---|---|---|
| | | | | | | | | | | Outer | | Inner | | |
| | Total | Outer | Inner | Marked | Unmarked | Outer | Inner | Outer | Inner | Marked | Unmarked | Marked | Unmarked | ICM:TE |
| _001 | 30 | 15 | 15 | 14 | 16 | 5 | 9 | 10 | 6 | 33.33% | 66.67% | 60.00% | 40.00% | 1.00 |
| _002 | 42 | 25 | 17 | 16 | 26 | 6 | 10 | 19 | 7 | 24.00% | 76.00% | 58.82% | 41.18% | 0.68 |
| _003 | 30 | 19 | 11 | 14 | 16 | 8 | 6 | 11 | 5 | 42.11% | 57.89% | 54.55% | 45.45% | 0.58 |
| _004 | 33 | 16 | 17 | 16 | 17 | 4 | 12 | 12 | 5 | 25.00% | 75.00% | 70.59% | 29.41% | 1.06 |
| _005 | 31 | 19 | 12 | 15 | 16 | 8 | 7 | 11 | 5 | 42.11% | 57.89% | 58.33% | 41.67% | 0.63 |
| _006 | 32 | 20 | 12 | 16 | 16 | 9 | 7 | 11 | 5 | 45.00% | 55.00% | 58.33% | 41.67% | 0.60 |
| _007 | 30 | 18 | 12 | 16 | 14 | 10 | 6 | 8 | 6 | 55.56% | 44.44% | 50.00% | 50.00% | 0.67 |
| _008 | 26 | 14 | 12 | 12 | 14 | 6 | 6 | 8 | 6 | 42.86% | 57.14% | 50.00% | 50.00% | 0.86 |
| _009 | 29 | 16 | 13 | 16 | 13 | 8 | 8 | 8 | 5 | 50.00% | 50.00% | 61.54% | 38.46% | 0.81 |
| _010 | 21 | 13 | 8 | 13 | 8 | 8 | 5 | 5 | 3 | 61.54% | 38.46% | 62.50% | 37.50% | 0.62 |
| _011 | 35 | 24 | 11 | 16 | 19 | 9 | 7 | 15 | 4 | 37.50% | 62.50% | 63.64% | 36.36% | 0.46 |
| _012 | 34 | 20 | 14 | 16 | 18 | 8 | 8 | 12 | 6 | 40.00% | 60.00% | 57.14% | 42.86% | 0.70 |
| _013 | 30 | 17 | 13 | 15 | 15 | 8 | 7 | 9 | 6 | 47.06% | 52.94% | 53.85% | 46.15% | 0.76 |
| _014 | 46 | 28 | 18 | 16 | 30 | 5 | 11 | 23 | 7 | 17.86% | 82.14% | 61.11% | 38.89% | 0.64 |
| _015 | 31 | 18 | 13 | 16 | 15 | 10 | 6 | 8 | 7 | 55.56% | 44.44% | 46.15% | 53.85% | 0.72 |
| _016 | 31 | 19 | 12 | 16 | 15 | 9 | 7 | 10 | 5 | 47.37% | 52.63% | 58.33% | 41.67% | 0.63 |
| _017 | 32 | 15 | 17 | 16 | 16 | 8 | 8 | 7 | 9 | 53.33% | 46.67% | 47.06% | 52.94% | 1.13 |
| _018 | 17 | 12 | 5 | 9 | 8 | 6 | 3 | 6 | 2 | 50.00% | 50.00% | 60.00% | 40.00% | 0.42 |
| _019 | 27 | 16 | 11 | 11 | 16 | 7 | 4 | 9 | 7 | 43.75% | 56.25% | 36.36% | 63.64% | 0.69 |
| Mean | 30.89 | 18.11 | 12.79 | 14.68 | 16.21 | 7.47 | 7.21 | 10.63 | 5.58 | 42.84% | 57.16% | 56.23% | 43.77% | 0.72 |
| Median | 31.00 | 18.00 | 12.00 | 16.00 | 16.00 | 8.00 | 7.00 | 10.00 | 6.00 | 43.75% | 56.25% | 58.33% | 41.67% | 0.68 |
| Sd | 6.33 | 4.11 | 3.19 | 2.06 | 5.04 | 1.71 | 2.23 | 4.39 | 1.57 | 11.44% | 11.44% | 7.70% | 7.70% | 0.19 |
| SEM | 1.45 | 0.94 | 0.73 | 0.47 | 1.16 | 0.39 | 0.51 | 1.01 | 0.36 | 2.62% | 2.62% | 1.77% | 1.77% | 0.04 |

**Supplementary Tables S1b:** Blastocyst lineage/positional contributions from injected/marked control siNTC and si*Tead4* cell clones, versus equivalent non-injected/unmarked clones, specifically at +48 h (E3.5) post-single 2-cell stage microinjection. Data include raw cell counts, clonal composition percentages for outer/TE and ICM populations, and the outer cell/TE:ICM ratio for the whole embryo. Relates to data in Figs. 1 and S1.

**Supplementary Tables 1c: +54h (Clonal *Tead4* KD - E3.75)**

**siNTC**

| Embryo Number | Raw values | | | | | | | | | Contribution to lineage (%) | | | | |
|---|---|---|---|---|---|---|---|---|---|---|---|---|---|---|
| | | | | | | Marked | | Unmarked | | Outer | | Inner | | |
| | Total | Outer | Inner | Marked | Unmarked | Outer | Inner | Outer | Inner | Marked | Unmarked | Marked | Unmarked | ICM:TE |
| _001 | 37 | 21 | 16 | 18 | 19 | 9 | 9 | 12 | 7 | 42.86% | 57.14% | 56.25% | 43.75% | 0.76 |
| _002 | 34 | 20 | 14 | 17 | 17 | 10 | 7 | 10 | 7 | 50.00% | 50.00% | 50.00% | 50.00% | 0.70 |
| _003 | 48 | 34 | 14 | 26 | 22 | 18 | 8 | 16 | 6 | 52.94% | 47.06% | 57.14% | 42.86% | 0.41 |
| _004 | 35 | 23 | 12 | 20 | 15 | 14 | 6 | 9 | 6 | 60.87% | 39.13% | 50.00% | 50.00% | 0.52 |
| _005 | 39 | 27 | 12 | 16 | 23 | 10 | 6 | 17 | 6 | 37.04% | 62.96% | 50.00% | 50.00% | 0.44 |
| _006 | 31 | 21 | 10 | 16 | 15 | 11 | 5 | 10 | 5 | 52.38% | 47.62% | 50.00% | 50.00% | 0.48 |
| _007 | 32 | 20 | 12 | 16 | 16 | 12 | 4 | 8 | 8 | 60.00% | 40.00% | 33.33% | 66.67% | 0.60 |
| _008 | 59 | 32 | 27 | 30 | 29 | 18 | 12 | 14 | 15 | 56.25% | 43.75% | 44.44% | 55.56% | 0.84 |
| _009 | 54 | 37 | 17 | 29 | 25 | 16 | 13 | 21 | 4 | 43.24% | 56.76% | 76.47% | 23.53% | 0.46 |
| _010 | 60 | 40 | 20 | 30 | 30 | 19 | 11 | 21 | 9 | 47.50% | 52.50% | 55.00% | 45.00% | 0.50 |
| _011 | 57 | 43 | 14 | 29 | 28 | 21 | 8 | 22 | 6 | 48.84% | 51.16% | 57.14% | 42.86% | 0.33 |
| _012 | 51 | 30 | 21 | 22 | 29 | 18 | 4 | 12 | 17 | 60.00% | 40.00% | 19.05% | 80.95% | 0.70 |
| _013 | 50 | 33 | 17 | 23 | 27 | 16 | 7 | 17 | 10 | 48.48% | 51.52% | 41.18% | 58.82% | 0.52 |
| _014 | 32 | 17 | 15 | 15 | 17 | 8 | 7 | 9 | 8 | 47.06% | 52.94% | 46.67% | 53.33% | 0.88 |
| _015 | 63 | 37 | 26 | 31 | 32 | 14 | 17 | 23 | 9 | 37.84% | 62.16% | 65.38% | 34.62% | 0.70 |
| _016 | 51 | 36 | 15 | 28 | 23 | 19 | 9 | 17 | 6 | 52.78% | 47.22% | 60.00% | 40.00% | 0.42 |
| _017 | 42 | 28 | 14 | 19 | 23 | 13 | 6 | 15 | 8 | 46.43% | 53.57% | 42.86% | 57.14% | 0.50 |
| _018 | 44 | 29 | 15 | 20 | 24 | 13 | 7 | 16 | 8 | 44.83% | 55.17% | 46.67% | 53.33% | 0.52 |
| _019 | 44 | 33 | 11 | 23 | 21 | 19 | 4 | 14 | 7 | 57.58% | 42.42% | 36.36% | 63.64% | 0.33 |
| _020 | 34 | 16 | 18 | 17 | 17 | 8 | 9 | 8 | 9 | 50.00% | 50.00% | 50.00% | 50.00% | 1.13 |
| _021 | 50 | 31 | 19 | 24 | 26 | 16 | 8 | 15 | 11 | 51.61% | 48.39% | 42.11% | 57.89% | 0.61 |
| _022 | 40 | 24 | 16 | 21 | 19 | 14 | 7 | 10 | 9 | 58.33% | 41.67% | 43.75% | 56.25% | 0.67 |
| _023 | 36 | 21 | 15 | 18 | 18 | 12 | 6 | 9 | 9 | 57.14% | 42.86% | 40.00% | 60.00% | 0.71 |
| _024 | 55 | 32 | 23 | 25 | 30 | 16 | 9 | 16 | 14 | 50.00% | 50.00% | 39.13% | 60.87% | 0.72 |
| _025 | 40 | 25 | 15 | 14 | 26 | 7 | 7 | 18 | 8 | 28.00% | 72.00% | 46.67% | 53.33% | 0.60 |
| _026 | 39 | 29 | 10 | 19 | 20 | 15 | 4 | 14 | 6 | 51.72% | 48.28% | 40.00% | 60.00% | 0.34 |
| Mean | 44.50 | 28.42 | 16.08 | 21.77 | 22.73 | 14.08 | 7.69 | 14.35 | 8.38 | 49.76% | 50.24% | 47.68% | 52.32% | 0.59 |
| Median | 43.00 | 29.00 | 15.00 | 20.50 | 23.00 | 14.00 | 7.00 | 14.50 | 8.00 | 50.00% | 50.00% | 46.67% | 53.33% | 0.56 |
| Sd | 9.69 | 7.25 | 4.44 | 5.30 | 5.17 | 3.93 | 3.02 | 4.47 | 3.05 | 7.79% | 7.79% | 11.13% | 11.13% | 0.19 |
| SEM | 1.90 | 1.42 | 0.87 | 1.04 | 1.01 | 0.77 | 0.59 | 0.88 | 0.60 | 1.53% | 1.53% | 2.18% | 2.18% | 0.04 |

**si*Tead4***

| Embryo Number | Raw values | | | | | | | | | Contribution to lineage (%) | | | | |
|---|---|---|---|---|---|---|---|---|---|---|---|---|---|---|
| | | | | | | Marked | | Unmarked | | Outer | | Inner | | |
| | Total | Outer | Inner | Marked | Unmarked | Outer | Inner | Outer | Inner | Marked | Unmarked | Marked | Unmarked | ICM:TE |
| _001 | 34 | 21 | 13 | 16 | 18 | 10 | 6 | 11 | 7 | 47.62% | 52.38% | 46.15% | 53.85% | 0.62 |
| _002 | 45 | 21 | 24 | 23 | 22 | 8 | 15 | 13 | 9 | 38.10% | 61.90% | 62.50% | 37.50% | 1.14 |
| _003 | 53 | 23 | 30 | 24 | 29 | 4 | 20 | 19 | 10 | 17.39% | 82.61% | 66.67% | 33.33% | 1.30 |
| _004 | 49 | 23 | 26 | 23 | 26 | 7 | 16 | 16 | 10 | 30.43% | 69.57% | 61.54% | 38.46% | 1.13 |
| _005 | 33 | 16 | 17 | 20 | 13 | 8 | 12 | 8 | 5 | 50.00% | 50.00% | 70.59% | 29.41% | 1.06 |
| _006 | 41 | 19 | 22 | 17 | 24 | 6 | 11 | 13 | 11 | 31.58% | 68.42% | 50.00% | 50.00% | 1.16 |
| _007 | 39 | 24 | 15 | 17 | 22 | 11 | 6 | 13 | 9 | 45.83% | 54.17% | 40.00% | 60.00% | 0.63 |
| _008 | 39 | 22 | 17 | 18 | 21 | 9 | 9 | 13 | 8 | 40.91% | 59.09% | 52.94% | 47.06% | 0.77 |
| _009 | 61 | 30 | 31 | 29 | 32 | 8 | 21 | 22 | 10 | 26.67% | 73.33% | 67.74% | 32.26% | 1.03 |
| _010 | 47 | 25 | 22 | 21 | 26 | 8 | 13 | 17 | 9 | 32.00% | 68.00% | 59.09% | 40.91% | 0.88 |
| _011 | 48 | 23 | 25 | 19 | 29 | 8 | 11 | 15 | 14 | 34.78% | 65.22% | 44.00% | 56.00% | 1.09 |
| _012 | 46 | 23 | 23 | 23 | 23 | 7 | 16 | 16 | 7 | 30.43% | 69.57% | 69.57% | 30.43% | 1.00 |
| _013 | 59 | 27 | 32 | 29 | 30 | 8 | 21 | 19 | 11 | 29.63% | 70.37% | 65.63% | 34.38% | 1.19 |
| _014 | 59 | 25 | 34 | 26 | 33 | 6 | 20 | 19 | 14 | 24.00% | 76.00% | 58.82% | 41.18% | 1.36 |
| _015 | 51 | 23 | 28 | 24 | 27 | 7 | 17 | 16 | 11 | 30.43% | 69.57% | 60.71% | 39.29% | 1.22 |
| _016 | 48 | 21 | 27 | 24 | 24 | 8 | 16 | 13 | 11 | 38.10% | 61.90% | 59.26% | 40.74% | 1.29 |
| _017 | 33 | 19 | 14 | 16 | 17 | 9 | 7 | 10 | 7 | 47.37% | 52.63% | 50.00% | 50.00% | 0.74 |
| Mean | 46.18 | 22.65 | 23.53 | 21.71 | 24.47 | 7.76 | 13.94 | 14.88 | 9.59 | 35.02% | 64.98% | 57.95% | 42.05% | 1.04 |
| Median | 47.00 | 23.00 | 24.00 | 23.00 | 24.00 | 8.00 | 15.00 | 15.00 | 10.00 | 32.00% | 68.00% | 59.26% | 40.74% | 1.09 |
| Sd | 8.88 | 3.22 | 6.54 | 4.18 | 5.41 | 1.60 | 5.10 | 3.66 | 2.40 | 9.09% | 9.09% | 9.28% | 9.28% | 0.23 |
| SEM | 2.15 | 0.78 | 1.59 | 1.01 | 1.31 | 0.39 | 1.24 | 0.89 | 0.58 | 2.20% | 2.20% | 2.25% | 2.25% | 0.06 |

**Supplementary Tables S1c:** Blastocyst lineage/positional contributions from injected/marked control siNTC and si*Tead4* cell clones, versus equivalent non-injected/unmarked clones, specifically at +54 h (E3.75) post-single 2-cell stage microinjection. Data include raw cell counts, clonal composition percentages for outer/TE and ICM populations, and the outer cell/TE:ICM ratio for the whole embryo. Relates to data in Figs. 1 and S1.

### Supplementary Tables 1d: +60h (Clonal *Tead4* KD - E4.0)

#### siNTC

| Embryo Number | Raw values | | | | | Marked | | Unmarked | | Contribution to lineage (%) Outer | | Inner | | ICM:TE |
|---|---|---|---|---|---|---|---|---|---|---|---|---|---|---|
| | Total | Outer | Inner | Marked | Unmarked | Outer | Inner | Outer | Inner | Marked | Unmarked | Marked | Unmarked | |
| _001 | 68 | 49 | 19 | 23 | 45 | 17 | 6 | 32 | 13 | 34.69% | 65.31% | 31.58% | 68.42% | 0.39 |
| _002 | 53 | 34 | 19 | 25 | 28 | 15 | 10 | 19 | 9 | 44.12% | 55.88% | 52.63% | 47.37% | 0.56 |
| _003 | 56 | 37 | 19 | 27 | 29 | 16 | 11 | 21 | 8 | 43.24% | 56.76% | 57.89% | 42.11% | 0.51 |
| _004 | 66 | 49 | 17 | 34 | 32 | 25 | 9 | 24 | 8 | 51.02% | 48.98% | 52.94% | 47.06% | 0.35 |
| _005 | 60 | 42 | 18 | 29 | 31 | 19 | 10 | 23 | 8 | 45.24% | 54.76% | 55.56% | 44.44% | 0.43 |
| _006 | 57 | 37 | 20 | 29 | 28 | 16 | 13 | 21 | 7 | 43.24% | 56.76% | 65.00% | 35.00% | 0.54 |
| _007 | 53 | 31 | 22 | 23 | 30 | 14 | 9 | 17 | 13 | 45.16% | 54.84% | 40.91% | 59.09% | 0.71 |
| _008 | 63 | 44 | 19 | 37 | 26 | 24 | 13 | 20 | 6 | 54.55% | 45.45% | 68.42% | 31.58% | 0.43 |
| _009 | 50 | 33 | 17 | 22 | 28 | 14 | 8 | 19 | 9 | 42.42% | 57.58% | 47.06% | 52.94% | 0.52 |
| _010 | 61 | 41 | 20 | 28 | 33 | 21 | 7 | 20 | 13 | 51.22% | 48.78% | 35.00% | 65.00% | 0.49 |
| _011 | 60 | 44 | 16 | 28 | 32 | 22 | 6 | 22 | 10 | 50.00% | 50.00% | 37.50% | 62.50% | 0.36 |
| _012 | 65 | 44 | 21 | 32 | 33 | 22 | 10 | 22 | 11 | 50.00% | 50.00% | 47.62% | 52.38% | 0.48 |
| _013 | 62 | 39 | 23 | 31 | 31 | 21 | 10 | 18 | 13 | 53.85% | 46.15% | 43.48% | 56.52% | 0.59 |
| _014 | 69 | 45 | 24 | 39 | 30 | 26 | 13 | 19 | 11 | 57.78% | 42.22% | 54.17% | 45.83% | 0.53 |
| _015 | 74 | 54 | 20 | 33 | 41 | 23 | 10 | 31 | 10 | 42.59% | 57.41% | 50.00% | 50.00% | 0.37 |
| _016 | 82 | 62 | 20 | 38 | 44 | 29 | 9 | 33 | 11 | 46.77% | 53.23% | 45.00% | 55.00% | 0.32 |
| _017 | 74 | 54 | 20 | 37 | 37 | 28 | 9 | 26 | 11 | 51.85% | 48.15% | 45.00% | 55.00% | 0.37 |
| _018 | 57 | 35 | 22 | 33 | 24 | 14 | 19 | 21 | 3 | 40.00% | 60.00% | 86.36% | 13.64% | 0.63 |
| _019 | 80 | 53 | 27 | 42 | 38 | 26 | 16 | 27 | 11 | 49.06% | 50.94% | 59.26% | 40.74% | 0.51 |
| Mean | 63.68 | 43.53 | 20.16 | 31.05 | 32.63 | 20.63 | 10.42 | 22.89 | 9.74 | 47.20% | 52.80% | 51.34% | 48.66% | 0.48 |
| Median | 62.00 | 44.00 | 20.00 | 31.00 | 31.00 | 21.00 | 10.00 | 21.00 | 10.00 | 46.77% | 53.23% | 50.00% | 50.00% | 0.49 |
| Sd | 9.04 | 8.36 | 2.63 | 5.83 | 5.85 | 4.97 | 3.25 | 4.78 | 2.66 | 5.69% | 5.69% | 12.86% | 12.86% | 0.10 |
| SEM | 2.07 | 1.92 | 0.60 | 1.34 | 1.34 | 1.14 | 0.75 | 1.10 | 0.61 | 1.31% | 1.31% | 2.95% | 2.95% | 0.02 |

#### si*Tead4*

| Embryo Number | Raw values | | | | | Marked | | Unmarked | | Contribution to lineage (%) Outer | | Inner | | ICM:TE |
|---|---|---|---|---|---|---|---|---|---|---|---|---|---|---|
| | Total | Outer | Inner | Marked | Unmarked | Outer | Inner | Outer | Inner | Marked | Unmarked | Marked | Unmarked | |
| _001 | 66 | 40 | 26 | 20 | 46 | 7 | 13 | 33 | 13 | 17.50% | 82.50% | 50.00% | 50.00% | 0.65 |
| _002 | 54 | 31 | 23 | 25 | 29 | 5 | 20 | 26 | 3 | 16.13% | 83.87% | 86.96% | 13.04% | 0.74 |
| _003 | 67 | 47 | 20 | 36 | 31 | 20 | 16 | 27 | 4 | 42.55% | 57.45% | 80.00% | 20.00% | 0.43 |
| _004 | 59 | 39 | 20 | 22 | 37 | 8 | 14 | 31 | 6 | 20.51% | 79.49% | 70.00% | 30.00% | 0.51 |
| _005 | 59 | 34 | 25 | 20 | 39 | 9 | 11 | 25 | 14 | 26.47% | 73.53% | 44.00% | 56.00% | 0.74 |
| _006 | 42 | 25 | 17 | 21 | 21 | 12 | 9 | 13 | 8 | 48.00% | 52.00% | 52.94% | 47.06% | 0.68 |
| _007 | 61 | 34 | 27 | 25 | 36 | 3 | 22 | 31 | 5 | 8.82% | 91.18% | 81.48% | 18.52% | 0.79 |
| _008 | 68 | 41 | 27 | 28 | 40 | 10 | 18 | 31 | 9 | 24.39% | 75.61% | 66.67% | 33.33% | 0.66 |
| _009 | 59 | 34 | 25 | 24 | 35 | 8 | 16 | 26 | 9 | 23.53% | 76.47% | 64.00% | 36.00% | 0.74 |
| _010 | 88 | 41 | 47 | 48 | 40 | 9 | 39 | 32 | 8 | 21.95% | 78.05% | 82.98% | 17.02% | 1.15 |
| Mean | 62.30 | 36.60 | 25.70 | 26.90 | 35.40 | 9.10 | 17.80 | 27.50 | 7.90 | 24.99% | 75.01% | 67.90% | 32.10% | 0.71 |
| Median | 60.00 | 36.50 | 25.00 | 24.50 | 36.50 | 8.50 | 16.00 | 29.00 | 8.00 | 22.74% | 77.26% | 68.33% | 31.67% | 0.71 |
| Sd | 11.76 | 6.24 | 8.21 | 8.81 | 6.98 | 4.58 | 8.43 | 5.85 | 3.60 | 11.87% | 11.87% | 15.12% | 15.12% | 0.19 |
| SEM | 3.72 | 1.97 | 2.60 | 2.79 | 2.21 | 1.45 | 2.67 | 1.85 | 1.14 | 3.75% | 3.75% | 4.78% | 4.78% | 0.06 |

**Supplementary Tables S1d:** Blastocyst lineage/positional contributions from injected/marked control siNTC and si*Tead4* cell clones, versus equivalent non-injected/unmarked clones, specifically at +60 h (E4.0) post-single 2-cell stage microinjection. Data include raw cell counts, clonal composition percentages for outer/TE and ICM, and the outer cell/TE:ICM ratio for the whole embryo. Relates to data in Figs. 1 and S1.

### Supplementary Tables 1e: +72h (Clonal *Tead4* KD - E4.5)

#### siNTC

| Embryo Number | Raw values | | | | | Marked | | Unmarked | | Contribution to lineage (%) | | | | ICM:TE |
|---|---|---|---|---|---|---|---|---|---|---|---|---|---|---|
| | | | | | | | | | | Outer | | Inner | | |
| | Total | Outer | Inner | Marked | Unmarked | Outer | Inner | Outer | Inner | Marked | Unmarked | Marked | Unmarked | ICM:TE |
| _001 | 79 | 65 | 14 | 49 | 30 | 39 | 10 | 26 | 4 | 60.00% | 40.00% | 71.43% | 28.57% | 0.22 |
| _002 | 79 | 60 | 19 | 33 | 46 | 23 | 10 | 37 | 9 | 38.33% | 61.67% | 52.63% | 47.37% | 0.32 |
| _003 | 83 | 65 | 18 | 44 | 39 | 33 | 11 | 32 | 7 | 50.77% | 49.23% | 61.11% | 38.89% | 0.28 |
| _004 | 73 | 60 | 13 | 32 | 41 | 24 | 8 | 36 | 5 | 40.00% | 60.00% | 61.54% | 38.46% | 0.22 |
| _005 | 85 | 62 | 23 | 61 | 24 | 47 | 14 | 15 | 9 | 75.81% | 24.19% | 60.87% | 39.13% | 0.37 |
| _006 | 94 | 69 | 25 | 46 | 48 | 35 | 11 | 34 | 14 | 50.72% | 49.28% | 44.00% | 56.00% | 0.36 |
| _007 | 86 | 67 | 19 | 36 | 50 | 27 | 9 | 40 | 10 | 40.30% | 59.70% | 47.37% | 52.63% | 0.28 |
| _008 | 95 | 72 | 23 | 54 | 41 | 42 | 12 | 30 | 11 | 58.33% | 41.67% | 52.17% | 47.83% | 0.32 |
| _009 | 85 | 67 | 18 | 39 | 46 | 36 | 3 | 31 | 15 | 53.73% | 46.27% | 16.67% | 83.33% | 0.27 |
| _010 | 81 | 59 | 22 | 45 | 36 | 33 | 12 | 26 | 10 | 55.93% | 44.07% | 54.55% | 45.45% | 0.37 |
| _011 | 94 | 69 | 25 | 34 | 60 | 28 | 6 | 41 | 19 | 40.58% | 59.42% | 24.00% | 76.00% | 0.36 |
| _012 | 91 | 72 | 19 | 45 | 46 | 36 | 9 | 36 | 10 | 50.00% | 50.00% | 47.37% | 52.63% | 0.26 |
| _013 | 91 | 69 | 22 | 40 | 51 | 29 | 11 | 40 | 11 | 42.03% | 57.97% | 50.00% | 50.00% | 0.32 |
| _014 | 83 | 63 | 20 | 45 | 38 | 30 | 15 | 33 | 5 | 47.62% | 52.38% | 75.00% | 25.00% | 0.32 |
| _015 | 84 | 65 | 19 | 42 | 42 | 34 | 8 | 31 | 11 | 52.31% | 47.69% | 42.11% | 57.89% | 0.29 |
| Mean | 85.53 | 65.60 | 19.93 | 43.00 | 42.53 | 33.07 | 9.93 | 32.53 | 10.00 | 50.43% | 49.57% | 50.72% | 49.28% | 0.30 |
| Median | 85.00 | 65.00 | 19.00 | 44.00 | 42.00 | 33.00 | 10.00 | 33.00 | 10.00 | 50.72% | 49.28% | 52.17% | 47.83% | 0.32 |
| Sd | 6.39 | 4.21 | 3.51 | 7.93 | 8.78 | 6.56 | 3.01 | 6.73 | 3.98 | 9.92% | 9.92% | 15.54% | 15.54% | 0.05 |
| SEM | 1.65 | 1.09 | 0.91 | 2.05 | 2.27 | 1.69 | 0.78 | 1.74 | 1.03 | 2.56% | 2.56% | 4.01% | 4.01% | 0.01 |

#### si*Tead4*

| Embryo Number | Raw values | | | | | Marked | | Unmarked | | Contribution to lineage (%) | | | | ICM:TE |
|---|---|---|---|---|---|---|---|---|---|---|---|---|---|---|
| | | | | | | | | | | Outer | | Inner | | |
| | Total | Outer | Inner | Marked | Unmarked | Outer | Inner | Outer | Inner | Marked | Unmarked | Marked | Unmarked | ICM:TE |
| _001 | 50 | 24 | 26 | 25 | 25 | 9 | 16 | 15 | 10 | 37.50% | 62.50% | 61.54% | 38.46% | 1.08 |
| _002 | 59 | 28 | 31 | 26 | 33 | 4 | 22 | 24 | 9 | 14.29% | 85.71% | 70.97% | 29.03% | 1.11 |
| _003 | 47 | 25 | 22 | 17 | 30 | 1 | 16 | 24 | 6 | 4.00% | 96.00% | 72.73% | 27.27% | 0.88 |
| _004 | 55 | 32 | 23 | 18 | 37 | 0 | 18 | 32 | 5 | 0.00% | 100.00% | 78.26% | 21.74% | 0.72 |
| _005 | 67 | 36 | 31 | 13 | 54 | 3 | 10 | 33 | 21 | 8.33% | 91.67% | 32.26% | 67.74% | 0.86 |
| _006 | 70 | 40 | 30 | 14 | 56 | 2 | 12 | 38 | 18 | 5.00% | 95.00% | 40.00% | 60.00% | 0.75 |
| _007 | 68 | 42 | 26 | 16 | 52 | 0 | 16 | 42 | 10 | 0.00% | 100.00% | 61.54% | 38.46% | 0.62 |
| _008 | 68 | 47 | 21 | 21 | 47 | 15 | 6 | 32 | 15 | 31.91% | 68.09% | 28.57% | 71.43% | 0.45 |
| _009 | 67 | 46 | 21 | 20 | 47 | 12 | 8 | 34 | 13 | 26.09% | 73.91% | 38.10% | 61.90% | 0.46 |
| _010 | 51 | 27 | 24 | 25 | 26 | 9 | 16 | 18 | 8 | 33.33% | 66.67% | 66.67% | 33.33% | 0.89 |
| _011 | 80 | 61 | 19 | 13 | 67 | 3 | 10 | 58 | 9 | 4.92% | 95.08% | 52.63% | 47.37% | 0.31 |
| _012 | 106 | 75 | 31 | 22 | 84 | 2 | 20 | 73 | 11 | 2.67% | 97.33% | 64.52% | 35.48% | 0.41 |
| _013 | 81 | 61 | 20 | 25 | 56 | 11 | 14 | 50 | 6 | 18.03% | 81.97% | 70.00% | 30.00% | 0.33 |
| _014 | 61 | 44 | 17 | 10 | 51 | 1 | 9 | 43 | 8 | 2.27% | 97.73% | 52.94% | 47.06% | 0.39 |
| _015 | 64 | 39 | 25 | 16 | 48 | 1 | 15 | 38 | 10 | 2.56% | 97.44% | 60.00% | 40.00% | 0.64 |
| _016 | 70 | 46 | 24 | 18 | 52 | 4 | 14 | 42 | 10 | 8.70% | 91.30% | 58.33% | 41.67% | 0.52 |
| _017 | 78 | 53 | 25 | 27 | 51 | 10 | 17 | 43 | 8 | 18.87% | 81.13% | 68.00% | 32.00% | 0.47 |
| _018 | 89 | 61 | 28 | 13 | 76 | 0 | 13 | 61 | 15 | 0.00% | 100.00% | 46.43% | 53.57% | 0.46 |
| _019 | 70 | 50 | 20 | 17 | 53 | 4 | 13 | 46 | 7 | 8.00% | 92.00% | 65.00% | 35.00% | 0.40 |
| Mean | 68.47 | 44.05 | 24.42 | 18.74 | 49.74 | 4.79 | 13.95 | 39.26 | 10.47 | 11.92% | 88.08% | 57.29% | 42.71% | 0.62 |
| Median | 68.00 | 44.00 | 24.00 | 18.00 | 51.00 | 3.00 | 14.00 | 38.00 | 10.00 | 8.00% | 92.00% | 61.54% | 38.46% | 0.52 |
| Sd | 14.29 | 13.99 | 4.31 | 5.14 | 15.47 | 4.67 | 4.12 | 14.62 | 4.22 | 12.24% | 12.24% | 14.27% | 14.27% | 0.25 |
| SEM | 3.28 | 3.21 | 0.99 | 1.18 | 3.55 | 1.07 | 0.94 | 3.35 | 0.97 | 2.81% | 2.81% | 3.27% | 3.27% | 0.06 |

**Supplementary Tables S1e:** Blastocyst lineage/positional contributions from injected/marked control siNTC and si*Tead4* cell clones, versus equivalent non-injected/unmarked clones, specifically at +72 h (E4.5) post-single 2-cell stage microinjection. Data include raw cell counts, clonal composition percentages for outer/TE and ICM, and the outer cell/TE:ICM ratio for the whole embryo. Relates to data in Figs. 1 and S1.

**Supplementary Tables 2: RT-qPCR (*Tead4* KD, *Tfap2c* KD and *Cdx2* KO)**

| si*Tead4* | | | | | | | | | | | | | | | |
|---|---|---|---|---|---|---|---|---|---|---|---|---|---|---|---|
| | | | *H2afz* | *Tead4* | | | *Gata3* | | | *Tfap2c* | | | *Cdx2* | | |
| | | | Ct | Ct | 2^-ΔCt | 2^-ΔΔCt | Ct | 2^-ΔCt | 2^-ΔΔCt | Ct | 2^-ΔCt | 2^-ΔΔCt | Ct | 2^-ΔCt | 2^-ΔΔCt |
| BioRep1 | Tech1 | si*Tead4* | 22.36 | 30.12 | 0.004 | 0.168 | 27.39 | 0.028 | 0.360 | 27.48 | 0.026 | 0.282 | 27.55 | 0.025 | 0.307 |
| | | | 22.15 | 30.04 | 0.004 | 0.177 | 27.31 | 0.029 | 0.381 | 27.31 | 0.029 | 0.317 | 28.51 | 0.013 | 0.158 |
| | | | 22.15 | 30.21 | 0.004 | 0.157 | 27.41 | 0.027 | 0.355 | 27.39 | 0.028 | 0.300 | 27.56 | 0.025 | 0.305 |
| | | siNTC | 21.62 | 27 | 0.025 | 0.984 | 25.44 | 0.072 | 0.940 | 25.10 | 0.092 | 0.991 | 25.32 | 0.079 | 0.973 |
| | | | 21.53 | 26.95 | 0.025 | 1.019 | 25.35 | 0.077 | 1.000 | 25.06 | 0.094 | 1.019 | 25.19 | 0.086 | 1.064 |
| | | | 21.81 | 26.98 | 0.025 | 0.998 | 25.26 | 0.082 | 1.064 | 25.10 | 0.092 | 0.991 | 25.33 | 0.078 | 0.966 |
| | Tech2 | si*Tead4* | 22.19 | 30.61 | 0.003 | 0.163 | 28.00 | 0.018 | 0.325 | 28.01 | 0.018 | 0.379 | 28.44 | 0.013 | 0.281 |
| | | | 22.17 | 30.53 | 0.003 | 0.172 | 27.98 | 0.018 | 0.329 | 28.15 | 0.016 | 0.344 | 29.10 | 0.008 | 0.178 |
| | | | 22.23 | 30.69 | 0.003 | 0.154 | 27.91 | 0.019 | 0.345 | 28.09 | 0.017 | 0.358 | 28.82 | 0.010 | 0.216 |
| | | siNTC | 21.48 | 27.33 | 0.018 | 1.002 | 25.73 | 0.055 | 0.991 | 25.91 | 0.048 | 1.028 | 25.99 | 0.046 | 0.973 |
| | | | 21.51 | 27.29 | 0.019 | 1.030 | 25.73 | 0.055 | 0.991 | 25.92 | 0.048 | 1.021 | 25.99 | 0.046 | 0.973 |
| | | | 21.62 | 27.38 | 0.017 | 0.968 | 25.69 | 0.056 | 1.019 | 26.02 | 0.045 | 0.953 | 25.87 | 0.050 | 1.057 |
| BioRep2 | Tech1 | si*Tead4* | 24.12 | 33.85 | 0.001 | 0.057 | 30.61 | 0.011 | 0.241 | 29.13 | 0.031 | 0.672 | 32.58 | 0.003 | 0.071 |
| | | | 24.11 | 33.94 | 0.001 | 0.053 | 30.68 | 0.011 | 0.230 | 29.39 | 0.026 | 0.561 | 32.51 | 0.003 | 0.074 |
| | | | 24.12 | 33.86 | 0.001 | 0.056 | 30.82 | 0.010 | 0.208 | 29.12 | 0.031 | 0.677 | 32.06 | 0.004 | 0.102 |
| | | siNTC | 22.51 | 28.18 | 0.019 | 0.942 | 26.99 | 0.044 | 0.964 | 26.99 | 0.044 | 0.964 | 27.19 | 0.039 | 0.966 |
| | | | 22.43 | 28.03 | 0.022 | 1.045 | 26.75 | 0.052 | 1.138 | 26.75 | 0.052 | 1.138 | 27.05 | 0.043 | 1.064 |
| | | | 22.55 | 28.07 | 0.021 | 1.016 | 27.07 | 0.042 | 0.912 | 27.07 | 0.042 | 0.912 | 27.18 | 0.039 | 0.973 |

| ds*Tfap2c* | | | | | | | | | | | | | | | |
|---|---|---|---|---|---|---|---|---|---|---|---|---|---|---|---|
| | | | *H2afz* | *Tead4* | | | *Gata3* | | | *Tfap2c* | | | *Cdx2* | | |
| | | | Ct | Ct | 2^-ΔCt | 2^-ΔΔCt | Ct | 2^-ΔCt | 2^-ΔΔCt | Ct | 2^-ΔCt | 2^-ΔΔCt | Ct | 2^-ΔCt | 2^-ΔΔCt |
| BioRep1 | Tech1 | ds*Tfap2c* | 22.13 | 26.79 | 0.039 | 0.891 | 24.15 | 0.241 | 0.689 | 27.53 | 0.023 | 0.255 | 25.67 | 0.084 | 0.893 |
| | | | 22.07 | 26.56 | 0.045 | 1.045 | 24.20 | 0.233 | 0.666 | 27.63 | 0.022 | 0.238 | 26.17 | 0.059 | 0.631 |
| | | | 22.09 | 26.52 | 0.047 | 1.074 | 24.27 | 0.222 | 0.634 | 27.70 | 0.021 | 0.227 | 26.01 | 0.066 | 0.705 |
| | | ds*EGFP* | 21.32 | 25.11 | 0.073 | 1.033 | 22.88 | 0.345 | 0.986 | 24.78 | 0.092 | 1.019 | 24.76 | 0.094 | 0.995 |
| | | | 21.44 | 25.26 | 0.066 | 0.931 | 22.81 | 0.362 | 1.035 | 24.87 | 0.087 | 0.957 | 24.77 | 0.093 | 0.989 |
| | | | 21.27 | 25.1 | 0.074 | 1.040 | 22.89 | 0.342 | 0.979 | 24.77 | 0.093 | 1.026 | 24.73 | 0.096 | 1.016 |
| | Tech2 | ds*Tfap2c* | 23.28 | 27.65 | 0.047 | 0.702 | 25.31 | 0.238 | 0.555 | 28.02 | 0.036 | 0.072 | 27.03 | 0.274 | 0.897 |
| | | | 23.21 | 27.66 | 0.047 | 0.697 | 25.21 | 0.255 | 0.595 | 28.07 | 0.035 | 0.077 | 26.94 | 0.265 | 0.955 |
| | | | 23.23 | 27.43 | 0.055 | 0.818 | 25.23 | 0.252 | 0.586 | 28.05 | 0.036 | 0.085 | 26.80 | 0.269 | 1.052 |
| | | ds*EGFP* | 22.48 | 26.16 | 0.084 | 1.260 | 23.83 | 0.424 | 0.989 | 25.58 | 0.126 | 0.950 | 25.76 | 0.950 | 1.382 |
| | | | 22.58 | 26.99 | 0.047 | 0.709 | 23.82 | 0.427 | 0.995 | 25.37 | 0.146 | 1.099 | 26.65 | 1.099 | 0.746 |
| | | | 22.72 | 26.33 | 0.075 | 1.120 | 23.79 | 0.436 | 1.016 | 25.57 | 0.127 | 0.957 | 26.27 | 0.957 | 0.970 |

| *Cdx2* CRISPR-Cas9 | | | | | | | | | | | | | | | |
|---|---|---|---|---|---|---|---|---|---|---|---|---|---|---|---|
| | | | H2afz | *Tead4* | | | *Gata3* | | | *Tfap2c* | | | *Cdx2* | | |
| | | | Ct | Ct | 2^-ΔCt | 2^-ΔΔCt | Ct | 2^-ΔCt | 2^-ΔΔCt | Ct | 2^-ΔCt | 2^-ΔΔCt | Ct | 2^-ΔCt | 2^-ΔΔCt |
| BioRep1 | Tech1 | Cdx2-KO | 27.83 | 30.96 | 0.103 | 1.030 | 29.65 | 0.255 | 0.572 | 30.79 | 0.116 | 0.984 | 31.64 | 0.064 | 0.529 |
| | | | 27.54 | 31.02 | 0.099 | 0.989 | 29.56 | 0.272 | 0.609 | 31.71 | 0.061 | 0.520 | 31.58 | 0.067 | 0.551 |
| | | | 27.67 | 31.03 | 0.098 | 0.982 | 29.67 | 0.252 | 0.564 | 31.04 | 0.097 | 0.827 | 31.93 | 0.053 | 0.432 |
| | | Cas9 Control | 26.11 | 28.92 | 0.132 | 1.087 | 27.25 | 0.420 | 0.942 | 29.20 | 0.109 | 0.924 | 29.18 | 0.110 | 0.908 |
| | | | 25.93 | 28.9 | 0.134 | 1.102 | 27.15 | 0.451 | 1.009 | 29.05 | 0.121 | 1.026 | 29.11 | 0.116 | 0.953 |
| | | | 25.96 | 29.02 | 0.123 | 0.950 | 27.09 | 0.470 | 1.052 | 29.01 | 0.124 | 1.055 | 28.83 | 0.141 | 1.157 |

**Supplementary Tables S2:** Summary of confirmatory RT-qPCR data showing the effects of global gene knockdown (KD) by si*Tead4* and ds*Tfap2c*, and CRISPR-Cas9–directed *Cdx2* knockout (KO), on TE-specific mRNA expression (*Tead4*, *Gata3*, *Tfap2c*, and *Cdx2*) after microinjection into both 2-cell blastomeres and culture to the E3.5 early-blastocyst stage (all normalised to *H2afz* expression levels). Relates to data in Figs. S1, S2, & S3.

**Supplemenatry Tables 3a: +48h (Clonal *Tfap2c* KD - E3.5)**

**ds*EGFP***

| Embryo Number | Total | Outer | Inner | Marked | Unmarked | Marked | | Unmarked | |
|---|---|---|---|---|---|---|---|---|---|
| | | | | | | Outer | Inner | Outer | Inner |
| _001 | 37 | 25 | 12 | 16 | 21 | 11 | 5 | 14 | 7 |
| _002 | 32 | 20 | 12 | 16 | 16 | 10 | 6 | 10 | 6 |
| _003 | 32 | 18 | 14 | 15 | 17 | 10 | 5 | 8 | 9 |
| _004 | 36 | 25 | 11 | 18 | 18 | 14 | 4 | 11 | 7 |
| _005 | 33 | 19 | 14 | 15 | 18 | 9 | 6 | 10 | 8 |
| _006 | 46 | 33 | 13 | 22 | 24 | 16 | 6 | 17 | 7 |
| _007 | 33 | 23 | 10 | 16 | 17 | 11 | 5 | 12 | 5 |
| _008 | 32 | 20 | 12 | 17 | 15 | 10 | 7 | 10 | 5 |
| _009 | 52 | 34 | 18 | 27 | 25 | 18 | 9 | 16 | 9 |
| _010 | 30 | 20 | 10 | 16 | 14 | 11 | 5 | 9 | 5 |
| _011 | 37 | 27 | 10 | 21 | 16 | 15 | 6 | 12 | 4 |
| _012 | 31 | 19 | 12 | 15 | 16 | 11 | 4 | 8 | 8 |
| _013 | 32 | 18 | 14 | 17 | 15 | 9 | 8 | 9 | 6 |
| _014 | 38 | 25 | 13 | 16 | 22 | 11 | 5 | 14 | 8 |
| _015 | 35 | 22 | 13 | 18 | 17 | 13 | 5 | 9 | 8 |
| _016 | 35 | 24 | 11 | 17 | 18 | 11 | 6 | 13 | 5 |
| _017 | 32 | 20 | 12 | 15 | 17 | 9 | 6 | 11 | 6 |
| _018 | 33 | 22 | 11 | 16 | 17 | 11 | 5 | 11 | 6 |
| _019 | 38 | 26 | 12 | 16 | 22 | 11 | 5 | 15 | 7 |
| _020 | 35 | 23 | 12 | 16 | 19 | 9 | 7 | 14 | 5 |
| Mean | 35.45 | 23.15 | 12.30 | 17.25 | 18.20 | 11.50 | 5.75 | 11.65 | 6.55 |
| Median | 34.00 | 22.50 | 12.00 | 16.00 | 17.00 | 11.00 | 5.50 | 11.00 | 6.50 |
| Sd | 5.29 | 4.46 | 1.84 | 2.95 | 3.05 | 2.48 | 1.25 | 2.66 | 1.47 |
| SEM | 1.18 | 1.00 | 0.41 | 0.66 | 0.68 | 0.55 | 0.28 | 0.60 | 0.33 |

**ds*Tfap2c***

| Embryo Number | Total | Outer | Inner | Marked | Unmarked | Marked | | Unmarked | |
|---|---|---|---|---|---|---|---|---|---|
| | | | | | | Outer | Inner | Outer | Inner |
| _001 | 45 | 26 | 19 | 22 | 23 | 12 | 10 | 14 | 9 |
| _002 | 37 | 21 | 16 | 15 | 22 | 8 | 7 | 13 | 9 |
| _003 | 31 | 19 | 12 | 16 | 15 | 9 | 7 | 10 | 5 |
| _004 | 39 | 27 | 12 | 21 | 18 | 17 | 4 | 10 | 8 |
| _005 | 58 | 36 | 22 | 28 | 30 | 19 | 9 | 17 | 13 |
| _006 | 36 | 24 | 12 | 16 | 20 | 12 | 4 | 12 | 8 |
| _007 | 51 | 33 | 18 | 25 | 26 | 15 | 10 | 18 | 8 |
| _008 | 37 | 20 | 17 | 19 | 18 | 15 | 4 | 5 | 13 |
| _009 | 31 | 20 | 11 | 14 | 17 | 9 | 5 | 11 | 6 |
| _010 | 24 | 15 | 9 | 13 | 11 | 9 | 4 | 6 | 5 |
| _011 | 31 | 22 | 9 | 16 | 15 | 11 | 5 | 11 | 4 |
| _012 | 31 | 20 | 11 | 14 | 17 | 9 | 5 | 11 | 6 |
| _013 | 35 | 25 | 10 | 17 | 18 | 11 | 6 | 14 | 4 |
| _014 | 31 | 16 | 15 | 16 | 15 | 7 | 9 | 9 | 6 |
| _015 | 24 | 15 | 9 | 14 | 10 | 9 | 5 | 6 | 4 |
| _016 | 28 | 16 | 12 | 14 | 14 | 8 | 6 | 8 | 6 |
| _017 | 48 | 31 | 17 | 21 | 27 | 11 | 10 | 20 | 7 |
| _018 | 32 | 20 | 12 | 15 | 17 | 10 | 5 | 10 | 7 |
| _019 | 32 | 20 | 12 | 16 | 16 | 11 | 5 | 9 | 7 |
| _020 | 38 | 27 | 11 | 15 | 23 | 9 | 6 | 18 | 5 |
| _021 | 31 | 18 | 13 | 16 | 15 | 11 | 5 | 7 | 8 |
| _022 | 32 | 20 | 12 | 15 | 17 | 7 | 8 | 13 | 4 |
| Mean | 35.55 | 22.32 | 13.23 | 17.18 | 18.36 | 10.86 | 6.32 | 11.45 | 6.91 |
| Median | 32.00 | 20.00 | 12.00 | 16.00 | 17.00 | 10.50 | 5.50 | 11.00 | 6.50 |
| Sd | 8.44 | 5.75 | 3.53 | 3.90 | 5.02 | 3.15 | 2.10 | 4.14 | 2.54 |
| SEM | 1.80 | 1.23 | 0.75 | 0.83 | 1.07 | 0.67 | 0.45 | 0.88 | 0.54 |

**Supplementary Tables S3a:** Blastocyst lineage/positional contributions from injected/marked control dsEGFP and ds*Tfap2c* cell clones, versus equivalent non-injected/unmarked clones, specifically at +48 h (E3.5) post-single 2-cell stage microinjection. Data include raw cell counts. Relates to data in Fig. S2.

**Supplemenatry Tables 3b: +72h (Clonal *Tfap2c* KD - E4.5)**

**ds*EGFP***

| Embryo Number | Total | Outer | Inner | Marked | Unmarked | Marked Outer | Marked Inner | Unmarked Outer | Unmarked Inner |
|---|---|---|---|---|---|---|---|---|---|
| _001 | 46 | 30 | 16 | 24 | 22 | 17 | 7 | 13 | 9 |
| _002 | 53 | 34 | 19 | 24 | 29 | 16 | 8 | 18 | 11 |
| _003 | 83 | 62 | 21 | 34 | 49 | 27 | 7 | 35 | 14 |
| _004 | 62 | 44 | 18 | 35 | 27 | 26 | 9 | 18 | 9 |
| _005 | 69 | 49 | 20 | 34 | 35 | 24 | 10 | 25 | 10 |
| _006 | 81 | 64 | 17 | 33 | 48 | 25 | 8 | 39 | 9 |
| _007 | 82 | 57 | 25 | 51 | 31 | 32 | 19 | 25 | 6 |
| _008 | 76 | 55 | 21 | 36 | 40 | 23 | 13 | 32 | 8 |
| _009 | 69 | 45 | 24 | 36 | 33 | 20 | 16 | 25 | 8 |
| _010 | 100 | 78 | 22 | 38 | 62 | 33 | 5 | 45 | 17 |
| _011 | 83 | 66 | 17 | 35 | 48 | 25 | 10 | 41 | 7 |
| _012 | 67 | 51 | 16 | 31 | 36 | 23 | 8 | 28 | 8 |
| _013 | 97 | 69 | 28 | 50 | 47 | 31 | 19 | 38 | 9 |
| _014 | 65 | 34 | 31 | 37 | 28 | 20 | 17 | 14 | 14 |
| _015 | 82 | 64 | 18 | 37 | 45 | 30 | 7 | 34 | 11 |
| _016 | 72 | 53 | 19 | 32 | 40 | 21 | 11 | 32 | 8 |
| _017 | 89 | 78 | 11 | 52 | 37 | 45 | 7 | 33 | 4 |
| _018 | 101 | 85 | 16 | 52 | 49 | 48 | 4 | 37 | 12 |
| _019 | 81 | 58 | 23 | 46 | 35 | 37 | 9 | 21 | 14 |
| _020 | 49 | 27 | 22 | 23 | 26 | 14 | 9 | 13 | 13 |
| _021 | 87 | 52 | 35 | 34 | 53 | 19 | 15 | 33 | 20 |
| _022 | 91 | 72 | 19 | 49 | 42 | 39 | 10 | 33 | 9 |
| _023 | 69 | 50 | 19 | 35 | 34 | 21 | 14 | 29 | 5 |
| _024 | 83 | 63 | 20 | 40 | 43 | 31 | 9 | 32 | 11 |
| _025 | 58 | 41 | 17 | 45 | 13 | 33 | 12 | 8 | 5 |
| _026 | 57 | 41 | 16 | 28 | 29 | 21 | 7 | 20 | 9 |
| _027 | 61 | 40 | 21 | 30 | 31 | 21 | 9 | 19 | 12 |
| _028 | 78 | 59 | 19 | 34 | 44 | 28 | 6 | 31 | 13 |
| _029 | 96 | 81 | 15 | 37 | 59 | 34 | 3 | 47 | 12 |
| _030 | 93 | 76 | 17 | 49 | 44 | 42 | 7 | 34 | 10 |
| _031 | 89 | 64 | 25 | 48 | 41 | 33 | 15 | 31 | 10 |
| _032 | 79 | 50 | 29 | 40 | 39 | 24 | 16 | 26 | 13 |
| _033 | 104 | 89 | 15 | 51 | 53 | 42 | 9 | 47 | 6 |
| _034 | 82 | 64 | 18 | 38 | 44 | 30 | 8 | 34 | 10 |
| _035 | 60 | 45 | 15 | 43 | 17 | 34 | 9 | 11 | 6 |
| _036 | 106 | 84 | 22 | 50 | 56 | 39 | 11 | 45 | 11 |
| _037 | 89 | 65 | 24 | 46 | 43 | 34 | 12 | 31 | 12 |
| _038 | 93 | 79 | 14 | 44 | 49 | 39 | 5 | 40 | 9 |
| _039 | 70 | 47 | 23 | 33 | 37 | 19 | 14 | 28 | 9 |
| _040 | 84 | 65 | 19 | 40 | 44 | 32 | 8 | 33 | 11 |
| _041 | 75 | 54 | 21 | 47 | 28 | 34 | 13 | 20 | 8 |
| _042 | 79 | 55 | 24 | 41 | 38 | 33 | 8 | 22 | 16 |
| _043 | 78 | 67 | 11 | 29 | 49 | 24 | 5 | 43 | 6 |
| _044 | 75 | 53 | 22 | 35 | 40 | 20 | 15 | 33 | 7 |
| _045 | 76 | 63 | 13 | 41 | 35 | 33 | 8 | 30 | 5 |
| _046 | 83 | 64 | 19 | 21 | 62 | 21 | 0 | 43 | 19 |
| _047 | 92 | 74 | 18 | 48 | 44 | 39 | 9 | 35 | 9 |
| _048 | 63 | 56 | 7 | 23 | 40 | 21 | 2 | 35 | 5 |
| Mean | 78.27 | 58.67 | 19.60 | 38.31 | 39.96 | 28.69 | 9.63 | 29.98 | 9.98 |
| Median | 80.00 | 58.50 | 19.00 | 37.00 | 40.00 | 29.00 | 9.00 | 32.00 | 9.00 |
| Sd | 14.37 | 14.83 | 5.12 | 8.54 | 10.71 | 8.23 | 4.23 | 9.78 | 3.60 |
| SEM | 2.07 | 2.14 | 0.74 | 1.23 | 1.55 | 1.19 | 0.61 | 1.41 | 0.52 |

**ds*Tfap2c***

| Embryo Number | Total | Outer | Inner | Marked | Unmarked | Marked Outer | Marked Inner | Unmarked Outer | Unmarked Inner |
|---|---|---|---|---|---|---|---|---|---|
| _001 | 87 | 70 | 17 | 30 | 57 | 26 | 4 | 44 | 13 |
| _002 | 49 | 32 | 17 | 13 | 36 | 8 | 5 | 24 | 12 |
| _003 | 64 | 44 | 20 | 18 | 46 | 18 | 0 | 26 | 20 |
| _004 | 91 | 69 | 22 | 30 | 61 | 27 | 3 | 42 | 19 |
| _005 | 82 | 64 | 18 | 21 | 61 | 19 | 2 | 45 | 16 |
| _006 | 66 | 53 | 13 | 22 | 44 | 16 | 6 | 37 | 7 |
| _007 | 73 | 52 | 21 | 17 | 56 | 11 | 6 | 41 | 15 |
| _008 | 54 | 41 | 13 | 19 | 35 | 16 | 3 | 25 | 10 |
| _009 | 64 | 46 | 18 | 24 | 40 | 14 | 10 | 32 | 8 |
| _010 | 78 | 61 | 17 | 36 | 42 | 32 | 4 | 29 | 13 |
| _011 | 58 | 44 | 14 | 20 | 38 | 12 | 8 | 32 | 6 |
| _012 | 65 | 51 | 14 | 31 | 34 | 27 | 4 | 24 | 10 |
| _013 | 60 | 37 | 23 | 21 | 39 | 16 | 5 | 21 | 18 |
| _014 | 82 | 69 | 13 | 42 | 40 | 36 | 6 | 33 | 7 |
| _015 | 58 | 38 | 20 | 21 | 37 | 14 | 7 | 24 | 13 |
| _016 | 79 | 57 | 22 | 25 | 54 | 22 | 3 | 35 | 19 |
| _017 | 57 | 41 | 16 | 17 | 40 | 12 | 5 | 29 | 11 |
| _018 | 81 | 59 | 22 | 37 | 44 | 31 | 6 | 28 | 16 |
| Mean | 69.33 | 51.56 | 17.78 | 24.67 | 44.67 | 19.83 | 4.83 | 31.72 | 12.94 |
| Median | 65.50 | 51.50 | 17.50 | 21.50 | 41.00 | 17.00 | 5.00 | 30.50 | 13.00 |
| Sd | 12.49 | 11.95 | 3.46 | 7.96 | 9.06 | 8.18 | 2.31 | 7.51 | 4.45 |
| SEM | 2.94 | 2.82 | 0.81 | 1.88 | 2.13 | 1.93 | 0.54 | 1.77 | 1.05 |

**Supplementary Tables S3b:** Blastocyst lineage/positional contributions from injected/marked control dsEGFP and ds*Tfap2c* cell clones, versus equivalent non-injected/unmarked clones, specifically at +72 h (E4.5) post-single 2-cell stage microinjection. Data include raw cell counts. Relates to data in Fig. S2.

### Supplementary Tables 4a: +48h (Clonal Cdx2 KO - E3.5)

***Cas9* mRNA+, *Cdx2* sgRNA-**

| Embryo Number | Total | Outer | Inner | Marked | Unmarked | | Marked Outer | Marked Inner | Unmarked Outer | Unmarked Inner |
|---|---|---|---|---|---|---|---|---|---|---|
| _001 | 36 | 20 | 16 | 16 | 20 | | 8 | 8 | 12 | 8 |
| _002 | 30 | 18 | 12 | 16 | 14 | | 10 | 6 | 8 | 6 |
| _003 | 45 | 27 | 18 | 19 | 26 | | 11 | 8 | 16 | 10 |
| _004 | 32 | 21 | 11 | 15 | 17 | | 11 | 4 | 10 | 7 |
| _005 | 27 | 19 | 8 | 14 | 13 | | 9 | 5 | 10 | 3 |
| _006 | 32 | 16 | 16 | 16 | 16 | | 8 | 8 | 8 | 8 |
| _007 | 23 | 15 | 8 | 11 | 12 | | 7 | 4 | 8 | 4 |
| _008 | 38 | 26 | 12 | 17 | 21 | | 10 | 7 | 16 | 5 |
| _009 | 35 | 19 | 16 | 20 | 15 | | 11 | 9 | 8 | 7 |
| Mean | 33.11 | 20.11 | 13.00 | 16.00 | 17.11 | | 9.44 | 6.56 | 10.67 | 6.44 |
| Median | 32.00 | 19.00 | 12.00 | 16.00 | 16.00 | | 10.00 | 7.00 | 10.00 | 7.00 |
| Sd | 6.41 | 4.08 | 3.67 | 2.65 | 4.48 | | 1.51 | 1.88 | 3.32 | 2.19 |
| SEM | 2.14 | 1.36 | 1.22 | 0.88 | 1.49 | | 0.50 | 0.63 | 1.11 | 0.73 |

***Cas9* mRNA-, *Cdx2* sgRNA+**

| Embryo Number | Total | Outer | Inner | Marked | Unmarked | | Marked Outer | Marked Inner | Unmarked Outer | Unmarked Inner |
|---|---|---|---|---|---|---|---|---|---|---|
| _001 | 32 | 16 | 16 | 16 | 16 | | 9 | 7 | 7 | 9 |
| _002 | 30 | 20 | 10 | 13 | 17 | | 9 | 4 | 11 | 6 |
| _003 | 35 | 18 | 17 | 17 | 18 | | 11 | 6 | 7 | 11 |
| _004 | 29 | 15 | 14 | 16 | 13 | | 9 | 7 | 6 | 7 |
| _005 | 33 | 19 | 14 | 15 | 18 | | 10 | 5 | 9 | 9 |
| _006 | 32 | 17 | 15 | 16 | 16 | | 9 | 7 | 8 | 8 |
| _007 | 34 | 21 | 13 | 16 | 18 | | 10 | 6 | 11 | 7 |
| _008 | 41 | 27 | 14 | 17 | 24 | | 11 | 6 | 16 | 8 |
| _009 | 33 | 23 | 10 | 16 | 17 | | 10 | 6 | 13 | 4 |
| _010 | 34 | 22 | 12 | 16 | 18 | | 9 | 7 | 13 | 5 |
| Mean | 33.30 | 19.80 | 13.50 | 15.80 | 17.50 | | 9.70 | 6.10 | 10.10 | 7.40 |
| Median | 33.00 | 19.50 | 14.00 | 16.00 | 17.50 | | 9.50 | 6.00 | 10.00 | 7.50 |
| Sd | 3.27 | 3.61 | 2.32 | 1.14 | 2.76 | | 0.82 | 0.99 | 3.25 | 2.07 |
| SEM | 1.03 | 1.14 | 0.73 | 0.36 | 0.87 | | 0.26 | 0.31 | 1.03 | 0.65 |

**Cas9 mRNA+, *Cdx2* sgRNA+**

| Embryo Number | Total | Outer | Inner | Marked | Unmarked | | Marked Outer | Marked Inner | Unmarked Outer | Unmarked Inner |
|---|---|---|---|---|---|---|---|---|---|---|
| _001 | 32 | 19 | 13 | 16 | 16 | | 8 | 8 | 11 | 5 |
| _002 | 32 | 19 | 13 | 16 | 16 | | 10 | 6 | 9 | 7 |
| _003 | 38 | 23 | 15 | 19 | 19 | | 12 | 7 | 11 | 8 |
| _004 | 34 | 20 | 14 | 15 | 19 | | 10 | 5 | 10 | 9 |
| _005 | 33 | 16 | 17 | 16 | 17 | | 8 | 8 | 8 | 9 |
| _006 | 33 | 16 | 17 | 16 | 17 | | 7 | 9 | 9 | 8 |
| _007 | 30 | 18 | 12 | 15 | 15 | | 8 | 7 | 10 | 5 |
| _008 | 33 | 18 | 15 | 17 | 16 | | 9 | 8 | 9 | 7 |
| _009 | 36 | 23 | 13 | 16 | 20 | | 11 | 5 | 12 | 8 |
| _010 | 35 | 21 | 14 | 16 | 19 | | 9 | 7 | 12 | 7 |
| _011 | 31 | 16 | 15 | 16 | 15 | | 9 | 7 | 7 | 8 |
| Mean | 33.36 | 19.00 | 14.36 | 16.18 | 17.18 | | 9.18 | 7.00 | 9.82 | 7.36 |
| Median | 33.00 | 19.00 | 14.00 | 16.00 | 17.00 | | 9.00 | 7.00 | 10.00 | 8.00 |
| Sd | 2.29 | 2.57 | 1.63 | 1.08 | 1.78 | | 1.47 | 1.26 | 1.60 | 1.36 |
| SEM | 0.69 | 0.77 | 0.49 | 0.33 | 0.54 | | 0.44 | 0.38 | 0.48 | 0.41 |

**Supplementary Tables S4a:** Blastocyst lineage/positional contributions from injected/marked control *Cas9* mRNA+ & *Cdx2* sgRNA- and *Cas9* mRNA+ & *Cdx2* sgRNA- cell clones, plus experimental *Cas9* mRNA+ & *Cdx2* sgRNA+ cell clones, versus equivalent non-injected/unmarked clones, specifically at +48 h (E3.5) post-single 2-cell stage microinjection. Data include raw cell counts. Relates to data in Fig. S3.

### Supplemenatry Tables 4b: +60h (Clonal Cdx2 KO - E4.0)

#### *Cas9* mRNA+, *Cdx2* sgRNA-

| Embryo Number | Total | Outer | Inner | Marked | Unmarked | Marked Outer | Marked Inner | Unmarked Outer | Unmarked Inner |
|---|---|---|---|---|---|---|---|---|---|
| _001 | 66 | 38 | 28 | 31 | 35 | 15 | 16 | 23 | 12 |
| _002 | 62 | 38 | 24 | 30 | 32 | 17 | 13 | 21 | 11 |
| _003 | 73 | 53 | 20 | 25 | 48 | 20 | 5 | 33 | 15 |
| _004 | 94 | 68 | 26 | 41 | 53 | 30 | 11 | 38 | 15 |
| _005 | 51 | 30 | 21 | 27 | 24 | 15 | 12 | 15 | 9 |
| _006 | 73 | 52 | 21 | 28 | 45 | 18 | 10 | 34 | 11 |
| _007 | 68 | 51 | 17 | 25 | 43 | 21 | 4 | 30 | 13 |
| _008 | 62 | 45 | 17 | 33 | 29 | 25 | 8 | 20 | 9 |
| _009 | 96 | 73 | 23 | 51 | 45 | 40 | 11 | 33 | 12 |
| _010 | 58 | 34 | 24 | 24 | 34 | 12 | 12 | 22 | 12 |
| _011 | 48 | 24 | 24 | 24 | 24 | 12 | 12 | 12 | 12 |
| _012 | 54 | 36 | 18 | 20 | 34 | 15 | 5 | 21 | 13 |
| _013 | 76 | 51 | 25 | 37 | 39 | 24 | 13 | 27 | 12 |
| _014 | 67 | 47 | 20 | 26 | 41 | 22 | 4 | 25 | 16 |
| _015 | 71 | 50 | 21 | 29 | 42 | 17 | 12 | 33 | 9 |
| _016 | 59 | 37 | 22 | 30 | 29 | 15 | 15 | 22 | 7 |
| _017 | 60 | 40 | 20 | 35 | 25 | 27 | 8 | 13 | 12 |
| Mean | 66.94 | 45.12 | 21.82 | 30.35 | 36.59 | 20.29 | 10.06 | 24.82 | 11.76 |
| Median | 66.00 | 45.00 | 21.00 | 29.00 | 35.00 | 18.00 | 11.00 | 23.00 | 12.00 |
| Sd | 13.16 | 12.70 | 3.11 | 7.48 | 8.80 | 7.26 | 3.77 | 7.76 | 2.36 |
| SEM | 3.19 | 3.08 | 0.75 | 1.81 | 2.14 | 1.76 | 0.91 | 1.88 | 0.57 |

#### *Cas9* mRNA-, *Cdx2* sgRNA+

| Embryo Number | Total | Outer | Inner | Marked | Unmarked | Marked Outer | Marked Inner | Unmarked Outer | Unmarked Inner |
|---|---|---|---|---|---|---|---|---|---|
| _001 | 60 | 41 | 19 | 22 | 38 | 15 | 7 | 26 | 12 |
| _002 | 98 | 71 | 27 | 43 | 55 | 30 | 13 | 41 | 14 |
| _003 | 56 | 39 | 17 | 28 | 28 | 21 | 7 | 18 | 10 |
| _004 | 84 | 62 | 22 | 45 | 39 | 33 | 12 | 29 | 10 |
| _005 | 79 | 55 | 24 | 43 | 36 | 26 | 17 | 29 | 7 |
| _006 | 75 | 58 | 17 | 40 | 35 | 34 | 6 | 24 | 11 |
| _007 | 74 | 47 | 27 | 32 | 42 | 20 | 12 | 27 | 15 |
| _008 | 87 | 66 | 21 | 37 | 50 | 29 | 8 | 37 | 13 |
| _009 | 83 | 57 | 26 | 44 | 39 | 34 | 10 | 23 | 16 |
| _010 | 96 | 63 | 33 | 48 | 48 | 28 | 20 | 35 | 13 |
| _011 | 66 | 37 | 29 | 28 | 38 | 16 | 12 | 21 | 17 |
| _012 | 89 | 67 | 22 | 41 | 48 | 34 | 7 | 33 | 15 |
| _013 | 82 | 55 | 27 | 42 | 40 | 28 | 14 | 27 | 13 |
| _014 | 59 | 32 | 27 | 28 | 31 | 16 | 12 | 16 | 15 |
| _015 | 69 | 37 | 32 | 26 | 43 | 11 | 15 | 26 | 17 |
| _016 | 91 | 65 | 26 | 44 | 47 | 36 | 8 | 29 | 18 |
| _017 | 60 | 37 | 23 | 28 | 32 | 18 | 10 | 19 | 13 |
| _018 | 80 | 52 | 28 | 39 | 41 | 25 | 14 | 27 | 14 |
| _019 | 58 | 33 | 25 | 26 | 32 | 19 | 7 | 14 | 18 |
| Mean | 76.11 | 51.26 | 24.84 | 36.00 | 40.11 | 24.89 | 11.11 | 26.37 | 13.74 |
| Median | 79.00 | 55.00 | 26.00 | 39.00 | 39.00 | 26.00 | 12.00 | 27.00 | 14.00 |
| Sd | 13.44 | 12.89 | 4.46 | 8.19 | 7.15 | 7.72 | 3.86 | 7.06 | 2.92 |
| SEM | 3.08 | 2.96 | 1.02 | 1.88 | 1.64 | 1.77 | 0.88 | 1.62 | 0.67 |

#### Cas9 mRNA+, *Cdx2* sgRNA+

| Embryo Number | Total | Outer | Inner | Marked | Unmarked | Marked Outer | Marked Inner | Unmarked Outer | Unmarked Inner |
|---|---|---|---|---|---|---|---|---|---|
| _001 | 83 | 63 | 20 | 31 | 52 | 26 | 5 | 37 | 15 |
| _002 | 60 | 45 | 15 | 36 | 24 | 26 | 10 | 19 | 5 |
| _003 | 77 | 50 | 27 | 30 | 47 | 19 | 11 | 31 | 16 |
| _004 | 69 | 47 | 22 | 24 | 45 | 16 | 8 | 31 | 14 |
| _005 | 51 | 31 | 20 | 26 | 25 | 13 | 13 | 18 | 7 |
| _006 | 75 | 48 | 27 | 34 | 41 | 20 | 14 | 28 | 13 |
| _007 | 63 | 42 | 21 | 27 | 36 | 17 | 10 | 25 | 11 |
| _008 | 86 | 56 | 30 | 39 | 47 | 23 | 16 | 33 | 14 |
| _009 | 75 | 55 | 20 | 36 | 39 | 25 | 11 | 30 | 9 |
| _010 | 91 | 58 | 33 | 48 | 43 | 32 | 16 | 26 | 17 |
| _011 | 76 | 54 | 22 | 34 | 42 | 24 | 10 | 30 | 12 |
| _012 | 79 | 51 | 28 | 31 | 48 | 26 | 5 | 25 | 23 |
| _013 | 80 | 59 | 21 | 34 | 46 | 19 | 15 | 40 | 6 |
| _014 | 89 | 66 | 23 | 37 | 52 | 28 | 9 | 38 | 14 |
| _015 | 84 | 49 | 35 | 42 | 42 | 19 | 23 | 30 | 12 |
| Mean | 75.87 | 51.60 | 24.27 | 33.93 | 41.93 | 22.20 | 11.73 | 29.40 | 12.53 |
| Median | 77.00 | 51.00 | 22.00 | 34.00 | 43.00 | 23.00 | 11.00 | 30.00 | 13.00 |
| Sd | 11.13 | 8.76 | 5.52 | 6.26 | 8.33 | 5.14 | 4.65 | 6.29 | 4.63 |
| SEM | 2.87 | 2.26 | 1.43 | 1.62 | 2.15 | 1.33 | 1.20 | 1.62 | 1.19 |

**Supplementary Tables S4b:** Blastocyst lineage/positional contributions from injected/marked control *Cas9* mRNA+ & *Cdx2* sgRNA- and *Cas9* mRNA+ & *Cdx2* sgRNA- cell clones, plus experimental *Cas9* mRNA+ & *Cdx2* sgRNA+ cell clones, versus equivalent non-injected/unmarked clones, specifically at +60 h (E4.0) post-single 2-cell stage microinjection. Data include raw cell counts. Relates to data in Fig. S3.

**Supplementary tables 5a: Summary of observed average (and proportional) internalisation events in live-embryo imaging movies (clonal siNTC & siTead4 treatments) - relates to Fig. 2.**

Note: These numbers represent the total number of cells per embryo at end of the movie (at frame 100 ) and their origin at the start of the movie (i.e. they do not represent the number of internalisations per se)

| Clonal siNTC | Total number of injected cells | Number of marked cells by origin | | | | | | | |
|---|---|---|---|---|---|---|---|---|---|
| | | Outer origin | | | | | | | Inner origin |
| | | Total | Final position: Outer | Final position: Inner | | | | | Final position: Inner |
| | | | | Division mediated internalisation | | | Migration mediated | | |
| | | | | Asymmetric (typical) | Atypical Division | | Constriction | Abscission | |
| Embryo # | | | | | Type I | Type II | | | |
| 1 | 32 | 21 | 19 | 2 | 0 | 0 | 0 | 0 | 11 |
| 2 | 29 | 18 | 17 | 1 | 0 | 0 | 0 | 0 | 11 |
| 3 | 19 | 18 | 12 | 2 | 0 | 0 | 2 | 2 | 1 |
| 4 | 37 | 27 | 27 | 0 | 0 | 0 | 0 | 0 | 10 |
| 5 | 35 | 28 | 25 | 1 | 0 | 0 | 2 | 0 | 7 |
| 6 | 35 | 20 | 19 | 1 | 0 | 0 | 0 | 0 | 15 |
| 7 | 24 | 15 | 15 | 0 | 0 | 0 | 0 | 0 | 9 |
| 8 | 24 | 18 | 12 | 1 | 0 | 0 | 2 | 0 | 8 |
| 9 | 30 | 20 | 15 | 5 | 0 | 0 | 0 | 0 | 10 |
| 10 | 35 | 25 | 25 | 0 | 0 | 0 | 0 | 0 | 10 |
| 11 | 33 | 20 | 19 | 1 | 0 | 0 | 0 | 0 | 13 |
| 12 | 20 | 10 | 19 | 1 | 0 | 0 | 0 | 0 | 10 |
| 13 | 41 | 26 | 26 | 0 | 0 | 0 | 0 | 0 | 15 |
| 14 | 32 | 16 | 14 | 2 | 0 | 0 | 0 | 0 | 16 |
| 15 | 23 | 16 | 14 | 0 | 0 | 0 | 0 | 0 | 9 |
| 16 | 21 | 13 | 11 | 0 | 0 | 0 | 2 | 0 | 8 |
| Average | 29.38 | 19.44 | **18.06** | 1.06 | 0.00 | 0.00 | 0.50 | 0.13 | **10.19** |
| Sum of average | | | | **1.69** | | | | | |
| | | | | **1.06** | | | **0.63** | | |

| Clonal siTead4 | Total number of injected cells | Number of marked cells by origin at the start of the movie | | | | | | | |
|---|---|---|---|---|---|---|---|---|---|
| | | Outer origin | | | | | | | Inner origin |
| | | Total | Final position: Outer | Final position: Inner | | | | | Final position: Inner |
| | | | | Division mediated internalisation | | | Migration mediated | | |
| | | | | Asymmetric (typical) | Atypical Division | | Constriction | Abscission | |
| Embryo # | | | | | Type I | Type II | | | |
| 1 | 17 | 14 | 9 | 2 | 0 | 0 | 2 | 0 | 4 |
| 2 | 38 | 15 | 2 | 1 | 3 | 4 | 0 | 5 | 23 |
| 3 | 26 | 20 | 6 | 1 | 2 | 0 | 11 | 0 | 6 |
| 4 | 16 | 7 | 8 | 4 | 0 | 0 | 0 | 1 | 1 |
| 5 | 11 | 10 | 6 | 0 | 0 | 0 | 1 | 3 | 1 |
| 6 | 35 | 21 | 5 | 6 | 1 | 0 | 8 | 5 | 14 |
| Average | 23.83 | 14.50 | **6.00** | 2.33 | 1.00 | 0.67 | 3.67 | 2.33 | **8.17** |
| Sum of averages | | | | **10.00** | | | | | |
| | | | | **4.00** | | | **6.00** | | |

| Clonal siNTC | Proportion of marked cells by origin at the start of the movie | | | | | | | |
|---|---|---|---|---|---|---|---|---|
| | Outer origin | | | | | | | Inner origin |
| | Total | Final position: Outer | Final position: Inner | | | | | Final position: Inner |
| | | | Division mediated internalisation | | | Migration mediated | | |
| | | | Asymmetric (typical) | Atypical Division | | Constriction | Abscission | |
| Embryo # | | | | Type I | Type II | | | |
| 1 | 0.563 | 0.594 | 0.063 | 0.000 | 0.000 | 0.000 | 0.000 | 0.344 |
| 2 | 0.621 | 0.586 | 0.034 | 0.000 | 0.000 | 0.000 | 0.000 | 0.379 |
| 3 | 1.421 | 0.632 | 0.105 | 0.000 | 0.000 | 0.105 | 0.105 | 0.053 |
| 4 | 0.757 | 0.730 | 0.000 | 0.000 | 0.000 | 0.000 | 0.000 | 0.270 |
| 5 | 0.571 | 0.714 | 0.029 | 0.000 | 0.000 | 0.057 | 0.000 | 0.200 |
| 6 | 0.571 | 0.543 | 0.029 | 0.000 | 0.000 | 0.000 | 0.000 | 0.429 |
| 7 | 0.625 | 0.625 | 0.000 | 0.000 | 0.000 | 0.000 | 0.000 | 0.375 |
| 8 | 0.750 | 0.500 | 0.042 | 0.000 | 0.000 | 0.083 | 0.000 | 0.333 |
| 9 | 0.667 | 0.500 | 0.167 | 0.000 | 0.000 | 0.000 | 0.000 | 0.333 |
| 10 | 0.714 | 0.714 | 0.000 | 0.000 | 0.000 | 0.000 | 0.000 | 0.286 |
| 11 | 0.606 | 0.576 | 0.030 | 0.000 | 0.000 | 0.000 | 0.000 | 0.394 |
| 12 | 0.500 | 0.950 | 0.050 | 0.000 | 0.000 | 0.000 | 0.000 | 0.500 |
| 13 | 0.634 | 0.634 | 0.000 | 0.000 | 0.000 | 0.000 | 0.000 | 0.366 |
| 14 | 0.500 | 0.438 | 0.063 | 0.000 | 0.000 | 0.000 | 0.000 | 0.500 |
| 15 | 0.696 | 0.609 | 0.000 | 0.000 | 0.000 | 0.000 | 0.000 | 0.391 |
| 16 | 0.619 | 0.524 | 0.000 | 0.000 | 0.000 | 0.095 | 0.000 | 0.381 |
| Average | 0.590 | **0.631** | 0.023 | 0.000 | 0.000 | 0.019 | 0.000 | **0.428** |
| Sum of averages | | | **0.042** | | | | | |
| | | | **0.023** | | | **0.019** | | |

| Clonal siTead4 | Proportion of marked cells by origin at the start of the movie | | | | | | | |
|---|---|---|---|---|---|---|---|---|
| | Outer origin | | | | | | | Inner origin |
| | Total | Final position: Outer | Final position: Inner | | | | | Final position: Inner |
| | | | Division mediated internalisation | | | Migration mediated | | |
| | | | Asymmetric (typical) | Atypical Division | | Constriction | Abscission | |
| Embryo # | | | | Type I | Type II | | | |
| 1 | 0.824 | 0.529 | 0.118 | 0.000 | 0.000 | 0.118 | 0.000 | 0.235 |
| 2 | 0.395 | 0.053 | 0.026 | 0.079 | 0.105 | 0.000 | 0.132 | 0.605 |
| 3 | 0.769 | 0.231 | 0.038 | 0.077 | 0.000 | 0.423 | 0.000 | 0.231 |
| 4 | 0.438 | 0.500 | 0.250 | 0.000 | 0.000 | 0.000 | 0.063 | 0.063 |
| 5 | 0.909 | 0.545 | 0.000 | 0.000 | 0.000 | 0.091 | 0.273 | 0.091 |
| 6 | 0.600 | 0.143 | 0.250 | 0.029 | 0.000 | 0.229 | 0.143 | 0.400 |
| Average | 0.656 | **0.334** | 0.114 | 0.031 | 0.018 | 0.143 | 0.102 | **0.271** |
| Sum of averages | | | **0.407** | | | | | |
| | | | **0.162** | | | **0.245** | | |

**Supplementary Tables S5a:** The categorised origin of internalised marked control (siNTC) and marked *Tead4* KD (si*Tead4*) clones at the end of the live-embryo imaging period, per imaged blastocyst (note, total cell numbers and proportion of relative contribution of each method of outer-cell internalisation are given – if a cell was already internalised at the start of the movie, it is designated as having an inner origin). Relates to data in Figs. 2 and S4 & S5.

**Supplementary tables 5b: Summary of observed frequency of internalisation events in live-embryo imaging movies (clonal siNTC & siTead4 treatments) - relates to Fig. 2.**

**Types of observed internalisation events**

| | |
|---|---|
| A | Asymmetric division |
| B | Contriction |
| C | Abscission |
| D | Type I (division associated) |
| E | Type II (division associated) |

**siNTC**
n=16

**Raw (sum total number of internalisation events observed within all embryos in the group. n=16)**

Number of injected cells

| | 1 | 2 | 3 | 4 | 5 | 6 | 7 | 8 | 9 | 10 | 11 | 12 | 13 | 14 | 15 | 16 | 32-cell stage | 17 | 18 | 19 | 20 | 21 | 22 | 23 | 24 | 25 | 26 | 27 | 28 | 29 | 30 | 31 | 32 | 64-cell stage | 33 | 34 | 35 | 36 | 37 | 38 | 39 | 40 | 41 | 42 | 43 |
|---|---|---|---|---|---|---|---|---|---|---|---|---|---|---|---|---|---|---|---|---|---|---|---|---|---|---|---|---|---|---|---|---|---|---|---|---|---|---|---|---|---|---|---|---|---|
| A | 0 | 0 | 0 | 0 | 0 | 0 | 0 | 0 | 0 | 0 | 0 | 1 | 1 | 0 | 1 | 0 | | 0 | 2 | 2 | 2 | 0 | 2 | 1 | 0 | 2 | 1 | 0 | 0 | 0 | 1 | 0 | 0 | | 0 | 0 | 0 | 0 | 0 | 0 | 0 | 0 | 0 | 0 | 0 |
| B | 0 | 0 | 0 | 0 | 0 | 0 | 0 | 0 | 0 | 0 | 0 | 0 | 1 | 0 | 3 | 1 | | 0 | 0 | 0 | 0 | 1 | 0 | 1 | 0 | 0 | 0 | 0 | 0 | 0 | 0 | 0 | 0 | | 0 | 0 | 0 | 0 | 0 | 0 | 0 | 0 | 0 | 0 | 0 |
| C | 0 | 0 | 0 | 0 | 0 | 0 | 0 | 0 | 0 | 0 | 0 | 0 | 0 | 0 | 0 | 0 | | 0 | 0 | 0 | 0 | 0 | 0 | 0 | 0 | 0 | 0 | 0 | 0 | 0 | 0 | 0 | 0 | | 0 | 0 | 0 | 0 | 0 | 0 | 0 | 0 | 0 | 0 | 0 |
| D | 0 | 0 | 0 | 0 | 0 | 0 | 0 | 0 | 0 | 0 | 0 | 0 | 0 | 0 | 0 | 0 | | 0 | 0 | 0 | 0 | 0 | 0 | 0 | 0 | 0 | 0 | 0 | 0 | 0 | 0 | 0 | 0 | | 0 | 0 | 0 | 0 | 0 | 0 | 0 | 0 | 0 | 0 | 0 |
| E | 0 | 0 | 0 | 0 | 0 | 0 | 0 | 0 | 0 | 0 | 0 | 0 | 0 | 0 | 0 | 0 | | 0 | 0 | 0 | 0 | 0 | 0 | 0 | 0 | 0 | 0 | 0 | 0 | 0 | 0 | 0 | 0 | | 0 | 0 | 0 | 0 | 0 | 0 | 0 | 0 | 0 | 0 | 0 |

**Population frequency (i.e. average number of events per embryo)**

| | 1 | 2 | 3 | 4 | 5 | 6 | 7 | 8 | 9 | 10 | 11 | 12 | 13 | 14 | 15 | 16 | 32-cell stage | 17 | 18 | 19 | 20 | 21 | 22 | 23 | 24 | 25 | 26 | 27 | 28 | 29 | 30 | 31 | 32 | 64-cell stage | 33 | 34 | 35 | 36 | 37 | 38 | 39 | 40 | 41 | 42 | 43 |
|---|---|---|---|---|---|---|---|---|---|---|---|---|---|---|---|---|---|---|---|---|---|---|---|---|---|---|---|---|---|---|---|---|---|---|---|---|---|---|---|---|---|---|---|---|---|
| A | 0 | 0 | 0 | 0 | 0 | 0 | 0 | 0 | 0 | 0 | 0 | 0.1 | 0.1 | 0 | 0.1 | 0 | | 0 | 0.1 | 0.1 | 0.1 | 0 | 0.1 | 0.1 | 0 | 0.1 | 0.1 | 0 | 0 | 0 | 0.1 | 0 | 0 | | 0 | 0 | 0 | 0 | 0 | 0 | 0 | 0 | 0 | 0 | 0 |
| B | 0 | 0 | 0 | 0 | 0 | 0 | 0 | 0 | 0 | 0 | 0 | 0 | 0.1 | 0 | 0.2 | 0.1 | | 0 | 0 | 0 | 0 | 0.1 | 0 | 0.1 | 0 | 0 | 0 | 0 | 0 | 0 | 0 | 0 | 0 | | 0 | 0 | 0 | 0 | 0 | 0 | 0 | 0 | 0 | 0 | 0 |
| C | 0 | 0 | 0 | 0 | 0 | 0 | 0 | 0 | 0 | 0 | 0 | 0 | 0 | 0 | 0 | 0 | | 0 | 0 | 0 | 0 | 0 | 0 | 0 | 0 | 0 | 0 | 0 | 0 | 0 | 0 | 0 | 0 | | 0 | 0 | 0 | 0 | 0 | 0 | 0 | 0 | 0 | 0 | 0 |
| D | 0 | 0 | 0 | 0 | 0 | 0 | 0 | 0 | 0 | 0 | 0 | 0 | 0 | 0 | 0 | 0 | | 0 | 0 | 0 | 0 | 0 | 0 | 0 | 0 | 0 | 0 | 0 | 0 | 0 | 0 | 0 | 0 | | 0 | 0 | 0 | 0 | 0 | 0 | 0 | 0 | 0 | 0 | 0 |
| E | 0 | 0 | 0 | 0 | 0 | 0 | 0 | 0 | 0 | 0 | 0 | 0 | 0 | 0 | 0 | 0 | | 0 | 0 | 0 | 0 | 0 | 0 | 0 | 0 | 0 | 0 | 0 | 0 | 0 | 0 | 0 | 0 | | 0 | 0 | 0 | 0 | 0 | 0 | 0 | 0 | 0 | 0 | 0 |

**si*Tead4***
n=6

**Raw (sum total number of internalisation events observed within all embryos. n=6)**

Number of injected cells

| | 1 | 2 | 3 | 4 | 5 | 6 | 7 | 8 | 9 | 10 | 11 | 12 | 13 | 14 | 15 | 16 | 32-cell stage | 17 | 18 | 19 | 20 | 21 | 22 | 23 | 24 | 25 | 26 | 27 | 28 | 29 | 30 | 31 | 32 | 64-cell stage | 33 | 34 | 35 | 36 | 37 | 38 | 39 | 40 | 41 | 42 | 43 |
|---|---|---|---|---|---|---|---|---|---|---|---|---|---|---|---|---|---|---|---|---|---|---|---|---|---|---|---|---|---|---|---|---|---|---|---|---|---|---|---|---|---|---|---|---|---|
| A | 0 | 0 | 0 | 0 | 0 | 0 | 0 | 0 | 0 | 0 | 0 | 1 | 0 | 0 | 0 | 1 | | 0 | 3 | 1 | 2 | 0 | 0 | 1 | 1 | 1 | 0 | 0 | 0 | 1 | 1 | 0 | 1 | | 2 | 1 | 0 | 0 | 0 | 0 | 1 | 0 | 0 | 1 | 0 |
| B | 0 | 0 | 0 | 0 | 0 | 0 | 0 | 0 | 0 | 0 | 0 | 1 | 0 | 0 | 1 | 3 | | 1 | 0 | 0 | 0 | 0 | 3 | 0 | 0 | 1 | 1 | 1 | 1 | 0 | 0 | 0 | 0 | | 0 | 0 | 0 | 0 | 0 | 0 | 1 | 0 | 0 | 0 | 0 |
| C | 0 | 0 | 0 | 0 | 0 | 0 | 0 | 0 | 0 | 1 | 0 | 2 | 0 | 0 | 0 | 0 | | 0 | 1 | 0 | 0 | 0 | 0 | 0 | 0 | 0 | 1 | 0 | 1 | 1 | 0 | 1 | 0 | | 0 | 0 | 0 | 1 | 0 | 0 | 1 | 1 | 0 | 1 | 1 |
| D | 0 | 0 | 0 | 0 | 0 | 0 | 0 | 0 | 0 | 0 | 0 | 0 | 0 | 0 | 0 | 0 | | 0 | 0 | 0 | 0 | 1 | 1 | 0 | 0 | 1 | 0 | 0 | 0 | 1 | 0 | 0 | 0 | | 0 | 0 | 0 | 0 | 0 | 0 | 0 | 0 | 0 | 0 | 0 |
| E | 0 | 0 | 0 | 0 | 0 | 0 | 0 | 0 | 0 | 0 | 0 | 0 | 0 | 0 | 0 | 0 | | 0 | 0 | 0 | 0 | 0 | 0 | 0 | 0 | 2 | 0 | 0 | 0 | 0 | 0 | 0 | 2 | | 0 | 0 | 0 | 0 | 0 | 0 | 0 | 0 | 0 | 0 | 0 |

**Population frequency (i.e. average number of events per embryo)**

| | 1 | 2 | 3 | 4 | 5 | 6 | 7 | 8 | 9 | 10 | 11 | 12 | 13 | 14 | 15 | 16 | 32-cell stage | 17 | 18 | 19 | 20 | 21 | 22 | 23 | 24 | 25 | 26 | 27 | 28 | 29 | 30 | 31 | 32 | 64-cell stage | 33 | 34 | 35 | 36 | 37 | 38 | 39 | 40 | 41 | 42 | 43 |
|---|---|---|---|---|---|---|---|---|---|---|---|---|---|---|---|---|---|---|---|---|---|---|---|---|---|---|---|---|---|---|---|---|---|---|---|---|---|---|---|---|---|---|---|---|---|
| A | 0 | 0 | 0 | 0 | 0 | 0 | 0 | 0 | 0 | 0 | 0 | 0.2 | 0 | 0 | 0 | 0.2 | | 0 | 0.5 | 0.2 | 0.3 | 0 | 0 | 0.2 | 0.2 | 0.2 | 0 | 0 | 0 | 0.2 | 0.2 | 0 | 0.2 | | 0.3 | 0.2 | 0 | 0 | 0 | 0 | 0.2 | 0 | 0 | 0.2 | 0 |
| B | 0 | 0 | 0 | 0 | 0 | 0 | 0 | 0 | 0 | 0 | 0 | 0.2 | 0 | 0 | 0.2 | 0.5 | | 0.2 | 0 | 0 | 0 | 0 | 0.5 | 0 | 0 | 0.2 | 0.2 | 0.2 | 0.2 | 0 | 0 | 0 | 0 | | 0 | 0 | 0 | 0 | 0 | 0 | 0.2 | 0 | 0 | 0 | 0 |
| C | 0 | 0 | 0 | 0 | 0 | 0 | 0 | 0 | 0 | 0.2 | 0 | 0.3 | 0 | 0 | 0 | 0 | | 0 | 0.2 | 0 | 0 | 0 | 0 | 0 | 0 | 0 | 0.2 | 0 | 0.2 | 0.2 | 0 | 0.2 | 0 | | 0 | 0 | 0 | 0.2 | 0 | 0 | 0.2 | 0.2 | 0 | 0.2 | 0.2 |
| D | 0 | 0 | 0 | 0 | 0 | 0 | 0 | 0 | 0 | 0 | 0 | 0 | 0 | 0 | 0 | 0 | | 0 | 0 | 0 | 0 | 0.2 | 0.2 | 0 | 0 | 0.2 | 0 | 0 | 0 | 0.2 | 0 | 0 | 0 | | 0 | 0 | 0 | 0 | 0 | 0 | 0 | 0 | 0 | 0 | 0 |
| E | 0 | 0 | 0 | 0 | 0 | 0 | 0 | 0 | 0 | 0 | 0 | 0 | 0 | 0 | 0 | 0 | | 0 | 0 | 0 | 0 | 0 | 0 | 0 | 0 | 0.3 | 0 | 0 | 0 | 0 | 0 | 0 | 0.3 | | 0 | 0 | 0 | 0 | 0 | 0 | 0 | 0 | 0 | 0 | 0 |

**Supplementary Tables S5b:** Summary matrix of the observed frequency of categorised mechanisms of internalisation of marked control (siNTC - upper) and marked *Tead4* KD (si*Tead4 -lower*) clones during the live-embryo imaging period, per imaged blastocyst. Internalisation frequencies of marked clones are highlighted in graded yellow shading, as a function of the number of marked/injected cells within an embryo (representing 50% of the all embryos due to the presence of the unmarked clone – hence, the notation of the overall 32- and 64-cell stages), as both absolute internalisation events or as a population frequency (i.e. averaged for the number of embryos assayed in each group). Relates to data in Figs. 2 and S4 & S5.

**Supplementary Tables 6a: YAP1 normalised fluorescence intensity (Relative Fluorescence Units - RFU)**

**Whole embryo**

| | siNTC | si*Tead4* |
|---|---|---|
| | 2782242.50 | 1934444.80 |
| | 2395659.60 | 1448502.30 |
| | 2557307.20 | 1218896.60 |
| | 1984819.60 | 1410909.40 |
| | 2846402.40 | 1161238.40 |
| | 2377453.10 | 1079038.30 |
| | 2182748.90 | 1079952.30 |
| | 1725254.20 | 775362.51 |
| | 1925204.10 | 783278.77 |
| | 1626192.00 | 747259.59 |
| | 2563146.60 | 802128.96 |
| | 1620582.50 | 842788.63 |
| | 1981381.40 | 815129.31 |
| | 1959864.20 | 1996425.60 |
| | 1560884.20 | 1779683.20 |
| | 2516521.00 | 1457264.70 |
| | 2712283.60 | 1318256.10 |
| | 2864515.70 | 1592217.90 |
| | 2589613.30 | 1782674.30 |
| | 2957645.60 | 1750060.10 |
| | 2628431.10 | 1862375.00 |
| | 2677088.20 | 1377768.00 |
| | 2713939.30 | |
| | 2645960.00 | |
| | 2783297.40 | |
| n | 25 | 22 |
| Mean | 2367137.51 | 1318893.40 |
| Median | 2557307.20 | 1348012.05 |
| Sd | 440578.82 | 418346.33 |
| SEM | 88115.76 | 89191.74 |

**Individual outer cells**

| | Cytoplasmic YAP1 | | | | Nuclear YAP1 | | | |
|---|---|---|---|---|---|---|---|---|
| | siNTC | | si*Tead4* | | siNTC | | si*Tead4* | |
| | Unmarked | Marked | Unmarked | Marked | Unmarked | Marked | Unmarked | Marked |
| | 55499.07 | 64116.44 | 63543.08 | 65974.12 | 114037.62 | 81027.62 | 99688.24 | 31810.60 |
| | 31742.83 | 51168.51 | 40474.33 | 23538.66 | 85011.06 | 78803.43 | 63300.25 | 13401.23 |
| | 96930.40 | 36480.75 | 30237.16 | 29364.86 | 168855.66 | 70224.33 | 56412.64 | 8471.74 |
| | 51197.80 | 48035.95 | 14727.94 | 31775.02 | 96546.73 | 74217.11 | 29281.95 | 15112.21 |
| | 56358.34 | 57780.02 | 17444.68 | 27854.00 | 127843.29 | 82820.26 | 24163.04 | 21242.83 |
| | 44410.24 | 37364.47 | 15066.03 | 17031.40 | 87733.37 | 60245.93 | 49780.62 | 15277.45 |
| | 46862.12 | 37156.38 | 34417.34 | 23227.07 | 60721.84 | 57593.45 | 40972.89 | 18352.10 |
| | 31870.30 | 32995.69 | 21830.48 | 21674.88 | 58097.48 | 96492.16 | 30053.66 | 12957.99 |
| | 27193.36 | 26053.89 | 16799.80 | 21081.51 | 49945.18 | 85733.52 | 18319.78 | 15058.14 |
| | 36760.35 | 51138.60 | 28305.67 | 42817.33 | 66317.35 | 38483.31 | 105414.94 | 17181.12 |
| | 27625.48 | 37214.13 | 18802.94 | 38633.26 | 70421.66 | 50370.52 | 61771.45 | 6819.65 |
| | 42101.98 | 23461.77 | 21453.28 | 29320.51 | 88940.11 | 33173.97 | 60748.80 | 12428.92 |
| | 34094.56 | 61992.48 | 15399.35 | 14570.87 | 79105.79 | 134753.08 | 71985.62 | 8541.89 |
| | 41121.00 | 37220.42 | 19870.94 | 14453.18 | 94081.39 | 103767.79 | 48341.38 | 12170.44 |
| | 36847.99 | 41706.33 | 19472.06 | 38257.99 | 66040.34 | 88784.26 | 48740.06 | 37671.02 |
| | 34526.92 | 44536.73 | 11441.15 | 17865.41 | 58943.89 | 104445.78 | 34722.39 | 11545.76 |
| | 48330.41 | 35212.54 | 12458.64 | 10336.92 | 74361.44 | 73456.08 | 32641.86 | 7430.71 |
| | 40034.70 | 35411.83 | 9990.62 | 22323.64 | 75804.95 | 68955.11 | 33167.03 | 10514.62 |
| | 28903.71 | 22050.64 | 26543.05 | 14013.97 | 76933.13 | 52481.40 | 38365.52 | 7517.54 |
| | 17598.75 | 22857.55 | 12997.08 | 15258.26 | 38937.22 | 50715.72 | 31231.80 | 6430.63 |
| | 31182.34 | 39470.34 | 44079.35 | 48738.26 | 42960.91 | 79872.90 | 90885.92 | 37373.40 |
| | 28287.80 | 22213.72 | 57990.50 | 78006.55 | 67047.75 | 52390.12 | 102773.39 | 35487.19 |
| | 20072.47 | 24904.01 | 36655.23 | 44151.61 | 25646.57 | 50471.81 | 93842.50 | 24855.11 |
| | 36685.05 | 24207.14 | 45364.88 | 60573.90 | 81342.48 | 73030.46 | 32998.98 | 32222.82 |
| | 14413.30 | 27053.98 | 59537.68 | 38010.19 | 20119.02 | 75583.11 | 63274.89 | 36970.58 |
| | 18530.64 | 32867.77 | 41400.65 | 52304.23 | 27460.76 | 40666.30 | 32568.14 | 45029.80 |
| | 36078.77 | 9151.99 | 37432.44 | 80920.06 | 33137.84 | 18477.58 | 99512.77 | 31379.87 |
| | 24472.43 | 29189.63 | 39339.93 | 25289.73 | 21537.54 | 42979.98 | 83835.36 | 11737.31 |
| | 18911.77 | 15955.42 | 25923.82 | 41659.11 | 61955.09 | 31927.22 | 60953.25 | 26970.75 |
| | 23671.04 | 24011.85 | 35879.90 | 57768.09 | 53832.59 | 48671.58 | 72693.49 | 45176.51 |
| | 15672.27 | 18412.73 | 14547.60 | 41517.71 | 53771.95 | 38309.94 | 56754.83 | 29544.53 |
| | | 17674.40 | 63877.47 | 51922.27 | | 43200.19 | 63399.42 | 33435.08 |
| | | | 32181.93 | 41600.33 | | | 41848.86 | 30951.88 |
| n | 31 | 32 | 33 | 33 | 31 | 32 | 33 | 33 |
| Mean | 35418.97 | 34033.38 | 29863.24 | 35813.18 | 68628.77 | 65066.44 | 56801.39 | 21547.62 |
| Median | 34094.56 | 34104.12 | 26543.05 | 31775.02 | 66317.35 | 64600.52 | 56412.64 | 17181.12 |
| Sd | 16150.59 | 13533.85 | 15751.53 | 18596.85 | 31907.67 | 25273.58 | 25328.72 | 12074.63 |
| SEM | 2900.73 | 2392.47 | 2741.99 | 3237.30 | 5730.79 | 4467.78 | 4409.16 | 2101.92 |

**Individual inner cells**

| | Cytoplasmic YAP1 | | | | Nuclear YAP1 | | | |
|---|---|---|---|---|---|---|---|---|
| | siNTC | | si*Tead4* | | siNTC | | si*Tead4* | |
| | Unmarked | Marked | Unmarked | Marked | Unmarked | Marked | Unmarked | Marked |
| | 39780.64 | 50087.24 | 58948.58 | 54110.87 | 16600.64 | 17047.72 | 14522.18 | 10939.94 |
| | 36602.86 | 30102.50 | 24093.29 | 29478.23 | 16457.89 | 13620.74 | 8691.44 | 7632.33 |
| | 51071.56 | 42444.87 | 26691.47 | 38629.41 | 15619.32 | 19234.05 | 8918.58 | 9334.86 |
| | 59570.43 | 65887.67 | 20463.26 | 36135.82 | 15342.16 | 21494.42 | 13626.12 | 10613.50 |
| | 79910.75 | 49713.44 | 32879.36 | 24368.82 | 40930.12 | 18623.75 | 11631.88 | 11760.48 |
| | 79197.65 | 39391.14 | 26773.38 | 33860.83 | 21487.66 | 14451.19 | 8802.59 | 7997.12 |
| | 70017.25 | 70089.43 | 24955.40 | 33446.72 | 22382.02 | 20993.82 | 8459.47 | 8446.35 |
| | 65273.63 | 44026.78 | 29775.82 | 11953.23 | 19184.65 | 10870.99 | 9510.31 | 3587.65 |
| | 40555.39 | 65189.97 | 20968.32 | 36300.23 | 13912.55 | 18297.21 | 10259.69 | 11805.50 |
| | 33464.88 | 35684.64 | 17140.65 | 38066.70 | 12482.83 | 14428.98 | 9140.34 | 11562.37 |
| | 40562.41 | 48482.60 | 16760.79 | 31747.57 | 11191.00 | 13487.93 | 8653.76 | 9962.65 |
| | 41987.08 | 46960.39 | 21519.26 | 10657.61 | 14205.21 | 15720.96 | 4946.63 | 5764.84 |
| | 51357.04 | 51979.88 | 16858.08 | 14416.11 | 19357.92 | 13491.06 | 4651.90 | 4637.67 |
| | 27324.19 | 23606.99 | 27445.90 | 32776.73 | 9442.95 | 10250.25 | 6234.90 | 8802.75 |
| | 61504.62 | 54604.63 | 15577.95 | 20621.10 | 18160.91 | 23364.98 | 5976.37 | 8389.77 |
| | 44381.31 | 34739.12 | 6931.97 | 13318.25 | 13470.79 | 15319.71 | 2388.02 | 6239.17 |
| | 41489.42 | 44537.03 | 22505.27 | 24247.49 | 13689.55 | 17502.54 | 8090.97 | 6862.26 |
| | 8282.58 | 33457.90 | 30400.73 | 19970.60 | 6710.09 | 10524.05 | 7232.96 | 4807.71 |
| | 18468.68 | 35482.30 | 16124.17 | 15224.39 | 6136.22 | 8356.74 | 6029.91 | 4216.86 |
| | 20937.18 | 29362.64 | 61002.46 | 45476.96 | 9696.15 | 13660.60 | 20682.15 | 11874.96 |
| | 27535.69 | 22896.07 | 35933.07 | 45825.28 | 10756.89 | 11896.83 | 13854.27 | 10405.88 |
| | 18206.01 | 23165.72 | 40880.96 | 36181.23 | 6420.75 | 9557.66 | 16712.99 | 13061.32 |
| | 22120.47 | 38508.05 | 41826.67 | 35302.30 | 6447.54 | 10912.46 | 12418.45 | 16155.24 |
| | 15597.28 | 15455.26 | 67861.30 | 41124.41 | 4352.77 | 5331.67 | 16420.33 | 14412.46 |
| | 15050.54 | 20769.80 | 40345.05 | 31424.21 | 3911.15 | 6174.43 | 14370.43 | 11895.88 |
| | 25513.66 | 18479.80 | 22411.68 | 37476.36 | 6923.03 | 5424.02 | 12231.56 | 14676.53 |
| | 28323.18 | 21302.56 | 27856.10 | 53159.66 | 8527.03 | 5370.63 | 18965.73 | 23775.95 |
| | 17944.48 | 16103.85 | 31504.78 | 45353.21 | 6937.09 | 8665.22 | 10983.64 | 14524.33 |
| | 25161.30 | 17319.36 | 38174.10 | 91655.13 | 7642.00 | 9310.21 | 19898.39 | 25946.10 |
| | 16078.13 | 13170.94 | 47081.57 | 54329.01 | 8886.00 | 5168.46 | 14960.17 | 17553.83 |
| | | | 39234.13 | 62785.98 | | | 12460.99 | 18818.25 |
| | | | 43594.86 | 46023.33 | | | 14173.21 | 12507.53 |
| n | 30 | 30 | 32 | 32 | 30 | 30 | 32 | 32 |
| Mean | 37442.34 | 36766.75 | 31078.76 | 35795.24 | 12908.83 | 12951.78 | 11121.89 | 11217.88 |
| Median | 35033.87 | 35583.47 | 27651.00 | 35719.06 | 11836.92 | 13489.50 | 10621.67 | 10776.72 |
| Sd | 19858.90 | 15942.40 | 14064.80 | 16741.81 | 7398.06 | 5148.82 | 4570.68 | 5244.99 |
| SEM | 3625.72 | 2910.67 | 2486.33 | 2959.56 | 1350.70 | 940.04 | 807.99 | 927.19 |

**Nuclear: Cytoplasmic YAP1 ratio**

| | Outer | | | | Inner | | | |
|---|---|---|---|---|---|---|---|---|
| | siNTC | | si*Tead4* | | siNTC | | si*Tead4* | |
| | Unmarked | Marked | Unmarked | Marked | Unmarked | Marked | Unmarked | Marked |
| | 2.06 | 1.26 | 1.62 | 0.48 | 0.43 | 0.34 | 0.25 | 0.21 |
| | 2.69 | 1.55 | 1.61 | 0.57 | 0.45 | 0.46 | 0.36 | 0.26 |
| | 1.81 | 1.93 | 1.87 | 0.29 | 0.31 | 0.46 | 0.34 | 0.24 |
| | 1.89 | 1.56 | 2.06 | 0.48 | 0.26 | 0.32 | 0.67 | 0.29 |
| | 2.27 | 1.44 | 1.40 | 0.77 | 0.51 | 0.37 | 0.35 | 0.48 |
| | 1.98 | 1.62 | 3.31 | 0.90 | 0.27 | 0.37 | 0.33 | 0.24 |
| | 1.29 | 1.55 | 1.19 | 0.79 | 0.32 | 0.30 | 0.34 | 0.25 |
| | 1.82 | 2.93 | 1.38 | 0.61 | 0.29 | 0.25 | 0.32 | 0.30 |
| | 1.85 | 3.32 | 1.10 | 0.72 | 0.34 | 0.28 | 0.50 | 0.34 |
| | 1.81 | 0.77 | 3.71 | 0.40 | 0.37 | 0.40 | 0.55 | 0.30 |
| | 2.61 | 1.35 | 3.34 | 0.18 | 0.28 | 0.28 | 0.54 | 0.32 |
| | 2.13 | 1.41 | 2.91 | 0.44 | 0.34 | 0.34 | 0.23 | 0.54 |
| | 2.40 | 2.20 | 4.75 | 0.60 | 0.38 | 0.26 | 0.28 | 0.33 |
| | 2.29 | 2.81 | 2.45 | 0.84 | 0.35 | 0.44 | 0.23 | 0.27 |
| | 1.82 | 2.13 | 2.61 | 0.99 | 0.30 | 0.44 | 0.40 | 0.41 |
| | 1.71 | 2.36 | 3.03 | 0.65 | 0.31 | 0.44 | 0.34 | 0.48 |
| | 1.54 | 2.09 | 2.73 | 0.73 | 0.33 | 0.39 | 0.36 | 0.30 |
| | 1.90 | 1.95 | 3.45 | 0.49 | 0.81 | 0.32 | 0.24 | 0.24 |
| | 2.67 | 2.46 | 1.49 | 0.54 | 0.33 | 0.24 | 0.37 | 0.28 |
| | 2.25 | 2.29 | 2.43 | 0.42 | 0.47 | 0.46 | 0.34 | 0.26 |
| | 1.38 | 2.03 | 2.07 | 0.77 | 0.39 | 0.52 | 0.39 | 0.23 |
| | 2.39 | 2.39 | 1.78 | 0.46 | 0.35 | 0.41 | 0.41 | 0.36 |
| | 1.28 | 2.03 | 2.58 | 0.59 | 0.29 | 0.28 | 0.29 | 0.48 |
| | 2.22 | 3.02 | 0.73 | 0.55 | 0.28 | 0.35 | 0.24 | 0.35 |
| | 1.43 | 2.86 | 1.07 | 0.98 | 0.26 | 0.30 | 0.36 | 0.39 |
| | 1.49 | 1.24 | 0.79 | 0.87 | 0.27 | 0.30 | 0.55 | 0.39 |
| | 0.93 | 2.21 | 2.66 | 0.39 | 0.31 | 0.25 | 0.68 | 0.46 |
| | 0.89 | 1.49 | 2.13 | 0.46 | 0.39 | 0.55 | 0.34 | 0.32 |
| | 3.29 | 2.02 | 2.36 | 0.65 | 0.30 | 0.54 | 0.53 | 0.28 |
| | 2.27 | 2.04 | 2.11 | 0.80 | 0.56 | 0.39 | 0.32 | 0.32 |
| | 3.43 | 2.13 | 4.00 | 0.71 | | | 0.32 | 0.30 |
| | | 2.46 | 0.99 | 0.64 | | | 0.33 | 0.28 |
| | | | 1.31 | 0.74 | | | | |
| n | 31 | 32 | 33 | 33 | 30 | 30 | 32 | 32 |
| Mean | 1.99 | 2.03 | 2.21 | 0.62 | 0.36 | 0.37 | 0.38 | 0.33 |
| Median | 1.90 | 2.04 | 2.11 | 0.61 | 0.33 | 0.36 | 0.34 | 0.30 |
| Sd | 0.59 | 0.59 | 0.98 | 0.19 | 0.11 | 0.09 | 0.12 | 0.09 |
| SEM | 0.11 | 0.10 | 0.17 | 0.03 | 0.02 | 0.02 | 0.02 | 0.02 |

**Supplementary Tables S6a:** Normalised fluorescence intensity (Relative Fluorescence Units; RFU) quantification of whole embryo YAP1 expression in control siNTC and si*Tead4*-microinjected blastocysts, and cytoplasmic and nuclear YAP1 levels in unmarked/marked cell clones located in inner and outer positions. Also shown are the associated Nuclear:Cytoplasmic (N:C) expression ratios. Relates to data in Figs. 3 and S6.

| Supplementary Tables 6b: PARD6B normalised fluorescence intensity (Relative Fluorescence Units - RFU) | | | | | | |
|---|---|---|---|---|---|---|
| | Whole embryo | | Individual outer-cells, apical PARD6B | | | |
| | siNTC | si*Tead4* | siNTC | | si*Tead4* | |
| | | | Unmarked | Marked | Unmarked | Marked |
| | 3311613.00 | 2612431.40 | 69834.16 | 91812.67 | 16407.88 | 48322.00 |
| | 2439579.00 | 2893339.80 | 37210.84 | 84833.87 | 36059.94 | 41166.79 |
| | 3329601.00 | 1520673.50 | 31171.70 | 36053.47 | 49529.47 | 40969.44 |
| | 3522367.00 | 2692245.60 | 47056.90 | 39966.20 | 132840.10 | 72931.59 |
| | 3735861.00 | 1779143.70 | 13726.55 | 47324.80 | 27641.53 | 23202.24 |
| | 3391352.00 | 2435475.50 | 54680.77 | 29258.17 | 41303.48 | 28055.41 |
| | 3219908.00 | 2983450.40 | 37950.26 | 30537.66 | 63327.55 | 38076.30 |
| | 2455310.00 | 3043421.30 | 46417.19 | 26240.68 | 70959.62 | 58510.41 |
| | 3665803.00 | 1923615.70 | 66706.12 | 45582.25 | 79807.53 | 18485.28 |
| | 2788326.00 | 3180340.80 | 23554.19 | 45953.64 | 93019.41 | 63696.37 |
| | 2296299.00 | 2471961.20 | 52349.74 | 27897.91 | 26136.57 | 26664.78 |
| | 2888279.00 | 2092600.20 | 37007.78 | 34119.09 | 25030.52 | 12290.12 |
| | 2898946.00 | 2873817.10 | 118842.40 | 63624.87 | 73715.80 | 49013.48 |
| | 3278286.00 | 1635152.50 | 78029.26 | 36460.19 | 38074.07 | 21199.73 |
| | 2460233.00 | 2201125.60 | 32806.29 | 44092.33 | 26633.86 | 10599.13 |
| | 2667170.00 | 1425461.80 | 42335.60 | 44475.29 | 36933.09 | 35558.85 |
| | 2344369.00 | 2100214.70 | 25399.12 | 35806.14 | 30013.30 | 22198.61 |
| | 2520134.00 | 1887239.30 | 42813.75 | 36995.97 | 38693.23 | 35774.72 |
| | 3239904.00 | 1929655.40 | 68322.08 | 31242.81 | 84967.23 | 42831.95 |
| | 2346632.00 | 2414912.60 | 29086.82 | 19319.78 | 69743.42 | 41991.50 |
| | 2318343.00 | 2201230.30 | 62648.63 | 29710.02 | 68624.63 | 15870.56 |
| | 2987719.00 | 1905317.10 | 60975.35 | 35060.84 | 47731.07 | 12642.59 |
| | 2800621.00 | | 48146.17 | 30962.76 | 17526.66 | 38867.50 |
| | 3186036.00 | | 65143.38 | 42634.97 | 19362.00 | 21555.38 |
| | 2880221.00 | | 106815.70 | 117161.30 | 60510.87 | 38799.97 |
| | | | 88532.66 | 104744.70 | 29588.84 | 29251.42 |
| | | | 86320.91 | 63081.08 | 34646.66 | 89973.29 |
| | | | 46290.56 | 69154.81 | 58055.73 | 45250.83 |
| | | | 66112.99 | 32118.54 | 42328.42 | 25843.66 |
| | | | 32882.23 | 62018.57 | 35606.17 | 43916.60 |
| | | | 34510.85 | 31659.68 | 28934.33 | 34206.06 |
| | | | 37638.80 | 42625.87 | 47342.96 | 50009.80 |
| | | | 40558.49 | 125805.40 | 51071.99 | 73440.82 |
| | | | 88636.78 | 56631.06 | 39694.95 | 44570.13 |
| | | | 40046.99 | 45139.18 | 45208.65 | 16473.72 |
| | | | 36755.12 | 29729.09 | 40775.84 | 20392.71 |
| | | | 56295.56 | 36254.64 | 58664.74 | 80081.88 |
| | | | 102231.50 | 52947.58 | 72655.03 | 28915.94 |
| | | | 46115.77 | 25053.46 | 24752.68 | 40798.04 |
| | | | 30228.38 | 56731.75 | 22337.65 | 16088.40 |
| | | | 56354.17 | 37381.01 | 22386.87 | 21712.76 |
| | | | 29906.98 | 69110.53 | 29038.74 | 18167.01 |
| | | | 57142.80 | 78663.24 | 27787.34 | 29397.69 |
| | | | 101288.90 | 75715.72 | 40035.32 | 19864.77 |
| | | | 63193.31 | 77450.85 | 41999.42 | 13780.44 |
| | | | 32594.38 | 48074.08 | 23677.45 | 25574.06 |
| | | | 42686.05 | 37830.69 | 18926.42 | 20076.16 |
| | | | 36192.93 | 48678.27 | 9444.28 | 30535.98 |
| | | | 52267.07 | 72003.24 | 29454.72 | 29040.05 |
| | | | 32531.78 | 30576.53 | 36051.80 | 35637.05 |
| | | | 38814.78 | 21688.90 | 26992.55 | 12234.71 |
| | | | 26809.62 | 55918.80 | 45615.77 | 38906.95 |
| | | | 13154.08 | 57643.46 | 28931.41 | 15755.61 |
| | | | 24596.31 | 38086.75 | 38831.07 | 22138.06 |
| | | | 26980.36 | 21849.98 | 44063.60 | 37912.09 |
| | | | 13697.09 | 20224.49 | 18032.24 | 13371.55 |
| | | | 26903.28 | 22146.94 | 29827.81 | 10929.09 |
| | | | 52176.02 | 41070.90 | 46493.86 | 38626.05 |
| | | | 8924.25 | 28932.27 | 30722.36 | 43671.25 |
| | | | 32869.60 | 25990.54 | 21795.44 | 14920.89 |
| | | | 30238.76 | 95201.74 | 42859.69 | 20013.27 |
| | | | 36598.84 | 19241.67 | 26984.69 | 33686.34 |
| | | | 29711.78 | 33671.50 | 25848.65 | 34976.52 |
| | | | 36725.63 | 40952.60 | 19216.55 | 19594.33 |
| | | | 26395.07 | 51982.16 | 23033.31 | 15483.99 |
| | | | 39411.26 | 6658.84 | 8108.80 | 13701.99 |
| | | | 45329.08 | 31308.36 | | |
| | | | 25411.52 | 20456.01 | | |
| | | | 44903.01 | 35826.18 | | |
| | | | 28813.39 | 52666.90 | | |
| | | | 38492.12 | 26005.74 | | |
| | | | 20684.43 | 21604.21 | | |
| | | | 59246.50 | 30015.35 | | |
| | | | 18514.59 | 26366.34 | | |
| | | | 19235.87 | 42289.60 | | |
| | | | 59141.29 | 21193.81 | | |
| | | | 16563.12 | 42720.05 | | |
| | | | 47686.90 | 33015.75 | | |
| n | 25 | 22 | 78 | 78 | 66 | 66 |
| Mean | 2918916.48 | 2281946.61 | 45184.30 | 44628.68 | 40339.66 | 32245.40 |
| Median | 2888279.00 | 2201177.95 | 39113.02 | 37605.85 | 36055.87 | 29145.74 |
| Sd | 449613.14 | 515217.64 | 22776.04 | 23045.07 | 21817.75 | 17339.18 |
| SEM | 89922.63 | 109844.77 | 2578.88 | 2609.34 | 2685.58 | 2134.31 |

**Supplementary Tables S6b:** Normalised fluorescence intensity (Relative Fluorescence Units; RFU) quantification of whole embryo PARD6B expression in control siNTC and si*Tead4*-microinjected embryos, apically-localised PARD6B levels in unmarked/marked outer cell clones. Relates to data in Fig. 3.

**Supplementary Tables 7a: AMOT normalised fluorescence intensity (Relative Fluorescence Units - RFU)**

| | Whole embryo | | Individual outer cells | | | | | | | | Individual inner cell membrane | | | |
|---|---|---|---|---|---|---|---|---|---|---|---|---|---|---|
| | | | Outer, apical domain | | | | Outer, lateral domain | | | | | | | |
| | siNTC | si*Tead4* | siNTC | | si*Tead4* | | siNTC | | si*Tead4* | | siNTC | | si*Tead4* | |
| | | | Unmarked | Marked | Unmarked | Marked | Unmarked | Marked | Unmarked | Marked | Unmarked | Marked | Unmarked | Marked |
| | 17881790.07 | 14838070.27 | 89483.17 | 99160.32 | 88605.56 | 141485.60 | 21335.58 | 21035.40 | 24315.38 | 27879.07 | 4323.21 | 3080.44 | 2256.96 | 997.37 |
| | 11192581.10 | 11761102.04 | 36155.78 | 137549.30 | 54262.56 | 74452.84 | 19747.41 | 20195.10 | 14877.88 | 12435.82 | 3073.82 | 3288.02 | 2560.05 | 1661.70 |
| | 19593142.39 | 11136246.57 | 137327.00 | 138017.10 | 53592.19 | 68336.91 | 11586.82 | 13229.69 | 3733.29 | 15918.28 | 2096.23 | 4020.12 | 1924.86 | 3018.94 |
| | 18593956.67 | 6041542.01 | 39004.13 | 114315.40 | 33525.86 | 64818.00 | 14637.64 | 8649.76 | 8355.58 | 11856.64 | 2065.01 | 3429.62 | 2582.27 | 1267.05 |
| | 11702564.02 | 8983200.24 | 51902.82 | 65019.06 | 54864.53 | 65340.78 | 18134.39 | 12777.24 | 2104.77 | 13055.96 | 1673.57 | 1762.97 | 2470.38 | 1315.20 |
| | 13228080.93 | 9105248.33 | 19616.36 | 58100.90 | 80051.54 | 63877.52 | 8146.97 | 2670.66 | 17077.98 | 25306.92 | 1464.42 | 1437.06 | 2213.27 | 815.86 |
| | 18297270.64 | 9372249.62 | 60785.53 | 153122.10 | 119309.90 | 53208.98 | 10463.95 | 13234.94 | 7461.18 | 9928.54 | 4215.21 | 4101.49 | 3955.41 | 1691.28 |
| | 12524396.06 | 8378518.89 | 53531.52 | 116307.60 | 51560.50 | 65515.00 | 4766.29 | 13840.06 | 17501.81 | 11069.87 | 3442.02 | 3767.96 | 2056.00 | 1971.02 |
| | 12480893.90 | 11527759.62 | 85769.13 | 93699.86 | 113924.20 | 50159.93 | 11470.74 | 29289.41 | 11996.85 | 20236.56 | 2965.28 | 2235.44 | 1186.98 | 858.60 |
| | 10894225.38 | 9518943.32 | 80212.19 | 80937.17 | 43437.35 | 99988.59 | 9965.59 | 56856.48 | 8069.04 | 34239.06 | 2872.80 | 3642.23 | 2163.83 | 1019.79 |
| | 12992070.46 | 9253819.50 | 58603.11 | 106260.80 | 62350.24 | 34848.54 | 7649.93 | 17570.14 | 3433.74 | 22459.37 | 3070.26 | 3380.67 | 807.83 | 1813.47 |
| | 18990064.19 | 8556417.16 | 65800.65 | 30139.44 | 62539.39 | 93159.89 | 9354.24 | 5184.13 | 6170.89 | 3344.08 | 1929.07 | 2490.64 | 2104.66 | 2416.41 |
| | 19049932.28 | 7461123.77 | 44373.26 | 58472.81 | 72645.47 | 27310.75 | 7953.52 | 8942.30 | 3237.09 | 10303.05 | 1502.35 | 2544.69 | 1600.38 | 1309.67 |
| | 15310364.89 | 8939922.48 | 120837.00 | 53437.33 | 48318.04 | 34302.36 | 11901.28 | 27536.49 | 6779.15 | 5369.72 | 1383.04 | 2201.49 | 1243.42 | 1051.92 |
| | 21561386.32 | 11790451.61 | 134786.40 | 49281.44 | 87618.54 | 21712.53 | 21093.00 | 8493.73 | 4472.44 | 21875.69 | 4829.89 | 2237.14 | 986.19 | 500.61 |
| | 18810613.20 | 7997401.20 | 118032.00 | 124524.90 | 46038.39 | 77225.68 | 11580.25 | 6147.93 | 15046.74 | 9290.99 | 4104.61 | 3236.76 | 1015.93 | 1506.83 |
| | 18164621.97 | 18363100.24 | 59374.66 | 54504.53 | 55379.25 | 39510.51 | 7912.32 | 9945.29 | 12808.15 | 10787.95 | 2513.93 | 4937.66 | 1323.38 | 1333.44 |
| | 14766985.24 | 9852139.83 | 66436.19 | 72922.29 | 52435.29 | 61348.39 | 5933.26 | 3392.20 | 7916.71 | 17845.47 | 1966.41 | 2853.14 | 2119.05 | 849.44 |
| | 13830820.06 | 9386249.64 | 76004.29 | 96871.92 | 32364.53 | 117067.70 | 4919.59 | 9753.39 | 18510.12 | 15092.40 | 1291.18 | 1539.50 | 2551.10 | 3170.09 |
| | 13710672.41 | 8387931.43 | 112131.10 | 81526.76 | 53802.34 | 66158.95 | 7236.97 | 12483.24 | 7342.29 | 21853.21 | 2774.66 | 1468.95 | 1779.38 | 1630.62 |
| | 12142343.75 | 7408919.87 | 122277.50 | 83363.80 | 55464.84 | 55043.37 | 15472.31 | 10110.91 | 16941.48 | 19226.61 | 3129.44 | 1855.67 | 1336.36 | 484.62 |
| | 14214304.45 | 8796763.70 | 113942.70 | 122230.60 | 54412.45 | 34760.32 | 7955.16 | 2626.94 | 4653.37 | 5739.98 | 2651.68 | 2497.16 | 927.56 | 2673.52 |
| | 14161696.74 | 8545908.26 | 57795.23 | 81377.62 | 57143.57 | 55905.90 | 21391.04 | 5901.36 | 10162.39 | 3320.63 | 2517.02 | 1773.84 | 1214.85 | 2464.16 |
| | | 9196729.18 | 65287.62 | 93812.32 | 60062.98 | 83977.42 | 12448.54 | 7588.34 | 14839.59 | 15110.05 | 3073.85 | 1850.27 | 1279.47 | 1501.17 |
| | | 10515644.28 | 72951.53 | 97036.97 | 40350.87 | 75758.53 | 8085.14 | 12664.66 | 8201.78 | 10965.42 | 2974.96 | 2981.33 | 1372.51 | 1224.45 |
| | | 7168667.77 | 61818.31 | 91114.86 | 37474.78 | 43660.65 | 3385.52 | 18023.28 | 15604.05 | 4428.11 | 2237.97 | 1449.52 | 1476.33 | 872.66 |
| | | 9481387.37 | 93901.52 | 48435.98 | 52882.69 | 46950.21 | 22257.59 | 8539.24 | 13707.61 | 9194.50 | 2561.45 | 1003.96 | 2138.35 | 1542.54 |
| | | 7346109.47 | 77295.26 | 41412.53 | 93569.86 | 56683.40 | 5306.84 | 10834.30 | 4947.11 | 12813.87 | 2291.53 | 3944.13 | 1566.22 | 751.23 |
| | | 8301735.39 | | | 31517.22 | 15700.63 | 8161.66 | 14422.10 | 3011.29 | 26515.94 | 2251.15 | 2949.72 | 1023.99 | 1056.66 |
| | | 9938450.17 | | | 60629.62 | 35173.40 | 24115.50 | 22547.90 | | 13952.39 | 1933.94 | 1775.63 | 1810.36 | 980.45 |
| | | 14830866.60 | | | 60398.55 | 60777.40 | 13453.13 | 12211.21 | | 3391.51 | 2321.31 | 3691.00 | 1048.05 | 1616.73 |
| | | 16401597.94 | | | 38960.99 | 39207.70 | 19989.44 | 16514.88 | | 4728.24 | 1837.04 | 2706.10 | 1163.54 | 1755.48 |
| | | 11103469.03 | | | | 46769.05 | 11704.96 | 11042.29 | | 4651.48 | | 2944.02 | | 1314.19 |
| | | 12776487.86 | | | | | 23315.59 | 5511.35 | | 1497.40 | | | | |
| | | 11839237.10 | | | | | 8140.41 | 10677.53 | | | | | | |
| | | 9605922.17 | | | | | 9439.78 | 6634.81 | | | | | | |
| | | 8226209.66 | | | | | 9197.78 | 13207.81 | | | | | | |
| | | 10927081.97 | | | | | 11106.30 | 13608.01 | | | | | | |
| | | 9305571.86 | | | | | 11840.70 | 8594.81 | | | | | | |
| | | 11560612.98 | | | | | 7325.03 | 15065.87 | | | | | | |
| | | 8177238.57 | | | | | 12582.62 | 2829.42 | | | | | | |
| | | 9236989.22 | | | | | 5404.34 | 4658.63 | | | | | | |
| | | 9231989.62 | | | | | 13115.47 | | | | | | | |
| | | 7791779.99 | | | | | 4415.32 | | | | | | | |
| | | 8938885.91 | | | | | 10513.52 | | | | | | | |
| | | | | | | | 14673.86 | | | | | | | |
| n | 23 | 45 | 28 | 28 | 32 | 33 | 46 | 42 | 29 | 34 | 32 | 33 | 32 | 33 |
| Mean | 15395425.09 | 9940126.53 | 77694.14 | 87248.42 | 59671.69 | 59702.95 | 11751.90 | 12739.12 | 10113.09 | 13402.49 | 2604.32 | 2699.34 | 1726.84 | 1467.79 |
| Median | 14214304.45 | 9253819.50 | 69693.86 | 87239.33 | 54638.49 | 56683.40 | 10809.91 | 10938.30 | 8201.78 | 12146.23 | 2515.47 | 2706.10 | 1583.30 | 1315.20 |
| Sd | 3191202.26 | 2452744.25 | 31210.81 | 31887.25 | 21493.74 | 26711.04 | 5558.70 | 9334.96 | 5787.30 | 8116.72 | 884.65 | 951.67 | 679.16 | 670.57 |
| SEM | 665411.67 | 365633.52 | 5898.29 | 6026.12 | 3799.59 | 4649.79 | 819.59 | 1440.42 | 1074.68 | 1392.01 | 156.38 | 165.66 | 120.06 | 116.73 |

**Supplementary Tables S7a:** Normalised fluorescence intensity (Relative Fluorescence Units; RFU) quantification of whole-embryo AMOT levels in control siNTC and si*Tead4*–microinjected blastocysts, including expression at the outer-cell apical and lateral domains and at inner-cell plasma membranes, for unmarked/marked cell clones. Relates to data in Figs. 3 and S7.

| Supplementary Tables 7b: CTNNB1 normalised fluorescence intensity (Relative Fluorescence Units - RFU) | | | | | | | | | | | | | | |
|---|---|---|---|---|---|---|---|---|---|---|---|---|---|---|
| | Whole embryo | | Individual outer cells | | | | | | | | Individual inner cell membrane | | | |
| | | | Outer, apical domain | | | | Outer, lateral domain | | | | | | | |
| | siNTC | *siTead4* | siNTC | | *siTead4* | | siNTC | | *siTead4* | | siNTC | | *siTead4* | |
| | | | Unmarked | Marked | Unmarked | Marked | Unmarked | Marked | Unmarked | Marked | Unmarked | Marked | Unmarked | Marked |
| | 4083647.55 | 8181135.66 | 16640.08 | 10368.59 | 18255.35 | 9700.71 | 32447.13 | 18320.96 | 44925.85 | 54564.88 | 887.58 | 1150.33 | 2889.02 | 2394.01 |
| | 6281086.48 | 5395483.41 | 6462.66 | 8085.57 | 8194.13 | 5245.79 | 28403.09 | 26471.76 | 32034.18 | 37069.15 | 1328.12 | 970.87 | 1642.01 | 1124.92 |
| | 4707746.48 | 7215794.58 | 17713.48 | 14262.80 | 24462.71 | 10885.64 | 21158.87 | 31636.42 | 11534.46 | 39467.98 | 680.53 | 1200.20 | 2037.87 | 2402.01 |
| | 4726542.34 | 4895962.83 | 10257.57 | 22102.82 | 10033.02 | 7447.03 | 39849.51 | 36887.21 | 14337.48 | 44099.55 | 1281.74 | 2015.33 | 1303.42 | 1626.34 |
| | 4888123.15 | 4478221.88 | 26957.35 | 11134.70 | 7637.55 | 8124.34 | 51452.98 | 31462.23 | 6764.21 | 12340.44 | 1265.25 | 1999.98 | 1861.53 | 1960.36 |
| | 7246036.52 | 5284720.37 | 21303.85 | 16592.80 | 14030.00 | 10740.17 | 16052.41 | 9641.88 | 38611.97 | 44641.83 | 1433.60 | 1672.86 | 2006.38 | 1258.54 |
| | 7006818.29 | 7315652.25 | 6783.40 | 7559.78 | 14179.45 | 13595.55 | 15315.55 | 29214.90 | 17791.31 | 17445.64 | 1038.81 | 1224.88 | 2439.48 | 1624.29 |
| | 4683334.02 | 4565577.72 | 6884.14 | 10338.60 | 19157.85 | 15389.72 | 22413.62 | 16702.52 | 57233.27 | 16659.96 | 971.68 | 1009.73 | 1564.36 | 2333.23 |
| | 4819599.48 | 5626568.67 | 32555.90 | 13247.97 | 22064.17 | 13439.98 | 20158.35 | 18560.99 | 13060.82 | 34188.01 | 1387.49 | 892.27 | 1688.20 | 1227.04 |
| | 7637066.39 | 3310994.23 | 13005.91 | 14300.11 | 12376.83 | 15478.87 | 13538.83 | 19288.41 | 12933.68 | 41966.97 | 1298.32 | 1011.90 | 1563.97 | 2143.95 |
| | 8169187.23 | 6610923.46 | 15429.54 | 13012.01 | 11630.24 | 15032.57 | 12981.80 | 11656.87 | 21357.65 | 47391.85 | 862.17 | 765.11 | 1086.06 | 1450.04 |
| | 8929066.15 | 5364449.43 | 16404.62 | 9086.69 | 14829.90 | 18151.67 | 7708.05 | 7938.49 | 22075.26 | 10437.85 | 1150.68 | 880.65 | 1775.69 | 2094.10 |
| | 13666388.21 | 6949893.09 | 10024.89 | 16295.87 | 9152.58 | 13744.38 | 15597.87 | 9985.06 | 18018.87 | 12533.00 | 855.13 | 2784.33 | 1372.42 | 1908.17 |
| | 7042725.17 | 6387943.03 | 27640.72 | 7135.22 | 14188.76 | 16442.73 | 44319.60 | 29478.85 | 33790.46 | 20885.39 | 1795.34 | 2550.23 | 1404.37 | 1697.77 |
| | 8435807.49 | 5374566.03 | 20179.39 | 6746.53 | 15658.56 | 30822.61 | 67558.75 | 44401.36 | 12581.59 | 45395.68 | 2178.27 | 2598.22 | 1971.78 | 1041.52 |
| | 6617620.44 | 5618936.94 | 13878.09 | 13885.67 | 12756.63 | 16685.91 | 40577.23 | 37071.28 | 45270.69 | 30376.31 | 1776.08 | 1709.61 | 1092.02 | 1475.94 |
| | 5353371.45 | 7267363.08 | 10775.31 | 18216.81 | 14476.23 | 6073.95 | 30797.77 | 44012.00 | 42382.92 | 25391.37 | 2342.11 | 1995.61 | 1112.06 | 1241.40 |
| | 9382457.24 | 6481922.80 | 10532.01 | 11054.35 | 6408.67 | 18240.75 | 21336.30 | 16370.51 | 41561.10 | 22055.05 | 1085.37 | 1150.58 | 1175.13 | 1224.91 |
| | 8956011.45 | 3978713.00 | 9995.04 | 13765.41 | 11575.75 | 6763.38 | 30814.34 | 44005.20 | 33448.48 | 47371.17 | 1364.55 | 1264.22 | 1052.74 | 1043.06 |
| | 7664434.62 | 2876487.64 | 9728.82 | 8307.64 | 14912.02 | 15527.86 | 25810.52 | 21511.71 | 19836.92 | 30857.57 | 1486.32 | 698.40 | 575.94 | 1594.05 |
| | 6219130.07 | 4500619.87 | 6445.28 | 7514.97 | 10549.35 | 8231.45 | 5065.49 | 33280.70 | 21550.53 | 24331.70 | 2426.48 | 895.82 | 1183.82 | 697.49 |
| | 7425067.64 | 4325739.41 | 9753.83 | 12425.15 | 15924.01 | 9199.35 | 16116.02 | 38978.10 | 26482.96 | 11416.49 | 1325.65 | 1222.06 | 886.58 | 1427.51 |
| | 5288103.85 | 5257821.05 | 8111.12 | 7677.17 | 13359.52 | 8870.81 | 54582.03 | 33186.01 | 22547.31 | 7167.52 | 1061.55 | 1238.07 | 568.17 | 1281.10 |
| | | 6361676.39 | 11841.28 | 14474.05 | 10186.18 | 12470.04 | 39794.39 | 36824.60 | 61505.58 | 18689.32 | 1374.02 | 1280.26 | 1306.62 | 930.46 |
| | | 7497173.47 | 57123.06 | 14351.47 | 10483.44 | 15126.77 | 42742.89 | 18384.08 | 16976.84 | 21793.85 | 977.55 | 739.79 | 1444.99 | 2007.77 |
| | | 5051966.52 | 12772.93 | 12405.77 | 6908.26 | 9313.48 | 23561.05 | 40248.35 | 25718.12 | 15562.36 | 1359.44 | 1062.20 | 982.51 | 1950.99 |
| | | 9977538.63 | 19148.72 | 7734.95 | 12288.06 | 13856.97 | 25991.61 | 22335.46 | 35918.99 | 39453.47 | 2350.38 | 999.99 | 1643.11 | 1481.11 |
| | | 3290289.44 | 11361.80 | 9464.01 | 27165.45 | 7083.73 | 13821.54 | 49704.46 | 23051.89 | 60076.82 | 1714.56 | 3252.28 | 1029.51 | 1281.35 |
| | | 5191105.39 | | | 13081.61 | 7782.05 | 33314.10 | 45446.93 | 18597.01 | 37992.43 | 2783.52 | 1903.36 | 1100.09 | 1661.23 |
| | | 6798481.28 | | | 13914.54 | 18741.84 | 33152.38 | 62202.47 | | 12524.41 | 1036.82 | 3897.64 | 2208.27 | 1797.39 |
| | | 8861624.32 | | | 13371.33 | 13823.22 | 43368.18 | 21132.91 | | 12742.74 | 1071.25 | 1727.79 | 1781.71 | 1862.01 |
| | | 8845830.88 | | | 7643.63 | 7829.78 | 55970.49 | 13554.55 | | 10180.47 | 1681.41 | 1136.14 | 1075.14 | 1789.71 |
| | | 6027081.80 | | | | 12255.73 | 9651.66 | 19172.95 | | 21447.01 | | 1116.48 | | 1892.05 |
| | | 6090874.13 | | | | | 23884.32 | 19028.39 | | 7441.07 | | | | |
| | | 7516419.01 | | | | | 7757.58 | 29608.35 | | | | | | |
| | | 5518200.32 | | | | | 32980.81 | 41508.36 | | | | | | |
| | | 5240551.09 | | | | | 25231.59 | 59542.29 | | | | | | |
| | | 6511408.54 | | | | | 17671.83 | 50116.75 | | | | | | |
| | | 3343589.58 | | | | | 29603.81 | 33818.08 | | | | | | |
| | | 7403435.07 | | | | | 28319.63 | 37136.63 | | | | | | |
| | | 5373964.02 | | | | | 33040.36 | 5748.35 | | | | | | |
| | | 5174983.68 | | | | | 16754.27 | 20678.94 | | | | | | |
| | | 4184673.13 | | | | | 30786.04 | | | | | | | |
| | | 8277004.85 | | | | | 20808.21 | | | | | | | |
| | | 4889037.36 | | | | | 20411.20 | | | | | | | |
| | | | | | | | 43551.04 | | | | | | | |
| n | 23 | 45 | 28 | 28 | 32 | 33 | 46 | 42 | 29 | 34 | 32 | 33 | 32 | 33 |
| Mean | 6923016.16 | 5882053.32 | 15703.96 | 11840.98 | 13464.24 | 12488.45 | 27961.42 | 29434.67 | 27306.91 | 27528.22 | 1425.99 | 1515.67 | 1463.28 | 1603.81 |
| Median | 7006818.29 | 5518200.32 | 12307.11 | 11770.24 | 13220.57 | 12470.04 | 25901.07 | 29543.60 | 22547.31 | 23193.38 | 1326.88 | 1222.06 | 1388.40 | 1624.29 |
| Sd | 2158185.27 | 1595696.36 | 10528.91 | 3801.22 | 4793.98 | 5094.48 | 13866.34 | 13972.81 | 13955.55 | 14967.12 | 516.14 | 767.24 | 518.55 | 434.07 |
| SEM | 450012.74 | 319139.27 | 2105.78 | 760.24 | 958.80 | 1018.90 | 2773.27 | 2794.56 | 2791.11 | 2993.42 | 103.23 | 153.45 | 103.71 | 86.81 |

**Supplementary Tables S7b:** Normalised fluorescence intensity (Relative Fluorescence Units; RFU) quantification of whole-embryo CTNNB1 levels in control siNTC and si*Tead4*–microinjected blastocysts, including expression at the outer-cell apical and lateral domains and at inner-cell plasma membranes, for unmarked/marked cell clones. Relates to data in Figs. 3 and S7.

**Supplementary table 8: Comparison of common DEGs in marked *Tead4* KD outer clones (E3.5) with those identified in gene-edited *Tead4* knockout whole embryos (E2.5);** *as reported by Wu et al., 2024 – Reproduction:167, e230322 - TEAD4 regulates KRT8 and YAP in preimplantation embryos in mice but not in cattle).*

| **Comparing Wu *et al.,* 2024 with Collier *et al.,* 2025** (RPKM >0.5 & fold change >2.0) | | |
|---|---|---|
| | **Wu *et al.,* 2024** | **Also in Collier *et al.*, 2025** |
| Downregulated DEGs | 310 | 125 |
| Upregulated DEGs | 360 | 105 |
| **Comparing Collier *et al.,* 2025 with Wu *et al.,* 2024** (RPKM >0.5 & fold change >2.0) | | |
| | **Collier *et al.*, 2025** | **Also in Wu *et al.,* 2024** |
| Downregulated DEGs | 604 | 125 (inc. 6 upregulated) |
| Upregulated DEGs | 677 | 99 |

*Note, Wu et al., 2024 data were derived from whole embryos at the E3.25 stage and RNA-Seq libraries that were prepared from amplified cDNA (contrasting with Collier et al., 2025 data derived from only outer-cell clones at E3.5 and RNA-Seq libraries prepared from non-amplified cDNA).*

**Supplementary Table 8:** Comparison of common DEGs exhibiting >2-fold gene expression differences in marked *Tead4* KD outer clones (E3.5) with those identified in gene-edited *Tead4* knockout whole embryos (E2.5); as reported by Wu et al., 2024[49]. Relates to data in Fig. 4.

**Supplementary table 9: Relationship between identified DEGs in marked *Tead4* KD outer clones (E3.5) with TEAD4-associated chromatin immunoprecipitation (ChIP) peaks in mouse Trophoblastic Stem Cells (TSCs);** *as reported by Home et al., 2012 – Proc. Natl. Acad. Sci. USA:109, 7362-7367 - Altered subcellular localization of transcription factor TEAD4 regulates first mammalian cell lineage commitment).*

| | | |
|---|---|---|
| **Collier *et al.,* 2025 identified DEGs** (RPKM >0.5 & fold change >2.0 – total; 1271) **associated with a TEAD4 ChIP peak in mouse TSCs; Home et al., 2012** | | |
| | **Genomic location of TEAD4 CHIP peak relative to DEG** | |
| | Promoter, gene body or 10Kb upstream of transcriptional start site | Promoter |
| Downregulated DEGs (604) | 110 | 14 |
| Upregulated DEGs (667) | 78 | 8 |
| **Collier *et al.,* 2025 identified DEGs** (RPKM >0.5 & fold change >10.0 – total; 247) **associated with a TEAD4 ChIP peak in mouse TSCs; Home et al., 2012** | | |
| | Promoter, gene body or 10Kb upstream of transcriptional start site | Promoter |
| Downregulated DEGs (92) | 8 | 3 |
| Upregulated DEGs (155) | 9 | 0 |

*Note, supplementary data Excel workbooks 2 & 3 list all DEGs identified in this study (Collier et al., 2025) and their association with previously associated mouse TSC TEAD4 ChIP peaks (Home et al., 2012) using RPKM >0.5 & fold change >2.0 and RPKM >0.5 & fold change >10.0 cutoffs, respectively.*

**Supplementary Table 9:** Relationship between identified DEGs >2-fold or >10-fold gene expression differences in marked *Tead4* KD outer clones (E3.5) with TEAD4-associated chromatin immunoprecipitation (ChIP-Seq) peaks in mouse Trophoblastic Stem Cells (TSCs); as reported by Home et al., 2012[50]. Relates to data in Fig. 4.

**Supplementary Table 10:**

| ID | Description | GeneRatio | BgRatio | RichFactor | FoldEnrichment | zScore | pvalue | p.adjust | qvalue |
|---|---|---|---|---|---|---|---|---|---|
| GO:0051015 | actin filament binding | 39/1062 | 221/28366 | 0.176470588 | 4.713526088 | 10.93005895 | 6.33E-16 | 6.73E-13 | 4.91E-13 |
| GO:0050839 | cell adhesion molecule binding | 41/1062 | 313/28366 | 0.130990415 | 3.498751527 | 8.766900955 | 3.50E-12 | 1.24E-09 | 9.06E-10 |
| GO:0030695 | GTPase regulator activity | 46/1062 | 430/28366 | 0.106976744 | 2.857346823 | 7.653950314 | 1.98E-10 | 4.21E-08 | 3.07E-08 |
| GO:0008514 | organic anion transmembrane transporter activity | 33/1062 | 279/28366 | 0.11827957 | 3.159245084 | 7.148120726 | 6.14E-09 | 1.09E-06 | 7.94E-07 |
| GO:0022804 | active transmembrane transporter activity | 42/1062 | 439/28366 | 0.095671982 | 2.555396832 | 6.477416783 | 3.21E-08 | 4.88E-06 | 3.56E-06 |
| GO:0005178 | integrin binding | 22/1062 | 155/28366 | 0.141935484 | 3.791094101 | 6.871817264 | 8.34E-08 | 1.11E-05 | 8.09E-06 |
| GO:0019838 | growth factor binding | 21/1062 | 148/28366 | 0.141891892 | 3.78992976 | 6.7112208 | 1.64E-07 | 1.94E-05 | 1.42E-05 |
| GO:0061629 | RNA polymerase II-specific DNA-binding transcription factor binding | 36/1062 | 387/28366 | 0.093023256 | 2.484649411 | 5.799677083 | 6.00E-07 | 5.95E-05 | 4.34E-05 |
| GO:0042578 | phosphoric ester hydrolase activity | 34/1062 | 356/28366 | 0.095505618 | 2.550953258 | 5.807746725 | 6.72E-07 | 5.95E-05 | 4.34E-05 |
| GO:0001217 | DNA-binding transcription repressor activity | 31/1062 | 310/28366 | 0.1 | 2.670998117 | 5.834242084 | 7.76E-07 | 5.98E-05 | 4.37E-05 |
| GO:0005319 | lipid transporter activity | 22/1062 | 176/28366 | 0.125 | 3.338747646 | 6.138081831 | 7.88E-07 | 5.98E-05 | 4.37E-05 |
| GO:0005006 | epidermal growth factor receptor activity | 11/1062 | 51/28366 | 0.215686275 | 5.76097633 | 6.711403377 | 2.32E-06 | 0.00015394 | 0.000112347 |
| GO:0140793 | histone H2AXY142 phosphatase activity | 14/1062 | 86/28366 | 0.162790698 | 4.348136469 | 6.132720741 | 3.59E-06 | 0.000188133 | 0.000137302 |
| GO:0017017 | MAP kinase tyrosine/serine/threonine phosphatase activity | 6/1062 | 13/28366 | 0.461538462 | 12.32768362 | 8.056636926 | 3.72E-06 | 0.000188133 | 0.000137302 |
| GO:0004620 | phospholipase activity | 15/1062 | 99/28366 | 0.151515152 | 4.046966844 | 5.989436721 | 4.10E-06 | 0.000198119 | 0.000144589 |
| GO:0051020 | GTPase binding | 31/1062 | 337/28366 | 0.091988131 | 2.457001235 | 5.306541723 | 4.48E-06 | 0.000198523 | 0.000144885 |
| GO:0005516 | calmodulin binding | 22/1062 | 198/28366 | 0.111111111 | 2.967775685 | 5.479875526 | 5.66E-06 | 0.000240498 | 0.000175518 |
| GO:0046943 | carboxylic acid transmembrane transporter activity | 22/1062 | 199/28366 | 0.110552764 | 2.95286224 | 5.452157135 | 6.14E-06 | 0.000250959 | 0.000183153 |
| GO:0044548 | S100 protein binding | 6/1062 | 15/28366 | 0.4 | 10.68399247 | 7.398716663 | 1.02E-05 | 0.000372514 | 0.000271865 |
| GO:0016757 | glycosyltransferase activity | 26/1062 | 271/28366 | 0.095940959 | 2.562581219 | 5.09745314 | 1.24E-05 | 0.000410457 | 0.000299556 |
| GO:0001221 | transcription coregulator binding | 18/1062 | 150/28366 | 0.12 | 3.20519774 | 5.340549683 | 1.37E-05 | 0.000441509 | 0.000322218 |
| GO:0005030 | neurotrophin receptor activity | 10/1062 | 51/28366 | 0.196078431 | 5.237251209 | 5.97312402 | 1.64E-05 | 0.000512341 | 0.000373912 |
| GO:0030246 | carbohydrate binding | 27/1062 | 293/28366 | 0.092150171 | 2.461329323 | 4.958816194 | 1.76E-05 | 0.000533623 | 0.000389444 |
| GO:0046873 | metal ion transmembrane transporter activity | 36/1062 | 450/28366 | 0.08 | 2.136798493 | 4.794062218 | 1.83E-05 | 0.000540437 | 0.000394417 |
| GO:0140789 | histone phosphatase activity | 16/1062 | 126/28366 | 0.126984127 | 3.39174364 | 5.306498782 | 2.00E-05 | 0.000559847 | 0.000408583 |
| GO:0140327 | flippase activity | 6/1062 | 17/28366 | 0.352941176 | 9.427052177 | 6.854436956 | 2.36E-05 | 0.000626133 | 0.000456959 |
| GO:0005201 | extracellular matrix structural constituent | 17/1062 | 142/28366 | 0.11971831 | 3.197673802 | 5.177720896 | 2.43E-05 | 0.000630124 | 0.000459872 |
| GO:0140791 | histone H2AXS140 phosphatase activity | 11/1062 | 71/28366 | 0.154929577 | 4.138166096 | 5.221445701 | 6.29E-05 | 0.001114836 | 0.00081362 |
| GO:0180004 | RNA polymerase II CTD heptapeptide repeat Y1 phosphatase activity | 11/1062 | 71/28366 | 0.154929577 | 4.138166096 | 5.221445701 | 6.29E-05 | 0.001114836 | 0.00081362 |
| GO:0180005 | RNA polymerase II CTD heptapeptide repeat T4 phosphatase activity | 11/1062 | 71/28366 | 0.154929577 | 4.138166096 | 5.221445701 | 6.29E-05 | 0.001114836 | 0.00081362 |
| GO:0180006 | RNA polymerase II CTD heptapeptide repeat S2 phosphatase activity | 11/1062 | 71/28366 | 0.154929577 | 4.138166096 | 5.221445701 | 6.29E-05 | 0.001114836 | 0.00081362 |
| GO:0180007 | RNA polymerase II CTD heptapeptide repeat S5 phosphatase activity | 11/1062 | 71/28366 | 0.154929577 | 4.138166096 | 5.221445701 | 6.29E-05 | 0.001114836 | 0.00081362 |
| GO:0180008 | RNA polymerase II CTD heptapeptide repeat S7 phosphatase activity | 11/1062 | 71/28366 | 0.154929577 | 4.138166096 | 5.221445701 | 6.29E-05 | 0.001114836 | 0.00081362 |
| GO:0003712 | transcription coregulator activity | 36/1062 | 478/28366 | 0.075313808 | 2.011630381 | 4.399138563 | 6.45E-05 | 0.001114836 | 0.00081362 |
| GO:0000146 | microfilament motor activity | 8/1062 | 38/28366 | 0.210526316 | 5.62315393 | 5.624225302 | 6.76E-05 | 0.001114836 | 0.00081362 |
| GO:0015294 | solute:monoatomic cation symporter activity | 14/1062 | 115/28366 | 0.12173913 | 3.251649881 | 4.771702768 | 0.000103234 | 0.001406898 | 0.00102677 |
| GO:0072341 | modified amino acid binding | 14/1062 | 115/28366 | 0.12173913 | 3.251649881 | 4.771702768 | 0.000103234 | 0.001406898 | 0.00102677 |
| GO:0005544 | calcium-dependent phospholipid binding | 9/1062 | 53/28366 | 0.169811321 | 4.535657179 | 5.081075566 | 0.000140489 | 0.001788267 | 0.001305098 |
| GO:0038064 | collagen receptor activity | 9/1062 | 53/28366 | 0.169811321 | 4.535657179 | 5.081075566 | 0.000140489 | 0.001788267 | 0.001305098 |
| GO:0005109 | frizzled binding | 8/1062 | 42/28366 | 0.19047619 | 5.08761546 | 5.228268708 | 0.000142886 | 0.001788267 | 0.001305098 |
| GO:0004364 | glutathione transferase activity | 7/1062 | 33/28366 | 0.212121212 | 5.665753581 | 5.288993117 | 0.000183855 | 0.002269732 | 0.001656476 |
| GO:0031406 | carboxylic acid binding | 19/1062 | 204/28366 | 0.093137255 | 2.487694324 | 4.205692902 | 0.000257337 | 0.003073583 | 0.002243136 |
| GO:0016500 | protein-hormone receptor activity | 10/1062 | 72/28366 | 0.138888889 | 3.709719607 | 4.540293647 | 0.000338332 | 0.003952166 | 0.002884335 |
| GO:0043177 | organic acid binding | 19/1062 | 210/28366 | 0.09047619 | 2.416617344 | 4.063658824 | 0.000371297 | 0.004269812 | 0.003116158 |
| GO:0140801 | histone H2AXY142 kinase activity | 13/1062 | 115/28366 | 0.113043478 | 3.019389175 | 4.279495226 | 0.000377575 | 0.004269812 | 0.003116158 |
| GO:0043425 | bHLH transcription factor binding | 7/1062 | 37/28366 | 0.189189189 | 5.05323968 | 4.865506311 | 0.000388933 | 0.004306628 | 0.003143026 |
| GO:0005200 | structural constituent of cytoskeleton | 10/1062 | 74/28366 | 0.135135135 | 3.609456915 | 4.432764717 | 0.000423801 | 0.004550513 | 0.003321016 |
| GO:1901681 | sulfur compound binding | 24/1062 | 303/28366 | 0.079207921 | 2.115642073 | 3.850522493 | 0.000496997 | 0.005283082 | 0.003855653 |
| GO:0001730 | 2'-5'-oligoadenylate synthetase activity | 4/1062 | 11/28366 | 0.363636364 | 9.712720425 | 5.700012525 | 0.000522042 | 0.00539263 | 0.003935603 |
| GO:0016671 | oxidoreductase activity, acting on a sulfur group of donors, disulfide as acceptor | 4/1062 | 11/28366 | 0.363636364 | 9.712720425 | 5.700012525 | 0.000522042 | 0.00539263 | 0.003935603 |
| GO:0016829 | lyase activity | 18/1062 | 199/28366 | 0.090452261 | 2.415978196 | 3.953241283 | 0.000525085 | 0.00539263 | 0.003935603 |
| GO:0170055 | lipid transmembrane transporter activity | 9/1062 | 63/28366 | 0.142857143 | 3.815711595 | 4.412479727 | 0.000537373 | 0.005440259 | 0.003970363 |
| GO:0048029 | monosaccharide binding | 11/1062 | 91/28366 | 0.120879121 | 3.228679042 | 4.199595224 | 0.00059006 | 0.005917298 | 0.004318511 |
| GO:0019207 | kinase regulator activity | 23/1062 | 289/28366 | 0.079584775 | 2.125707844 | 3.793497282 | 0.000603442 | 0.005970578 | 0.004357396 |
| GO:0070851 | growth factor receptor binding | 15/1062 | 152/28366 | 0.098684211 | 2.635853405 | 3.988176406 | 0.000606606 | 0.005970578 | 0.004357396 |
| GO:0004602 | glutathione peroxidase activity | 6/1062 | 29/28366 | 0.206896552 | 5.526203 | 4.809463207 | 0.000615519 | 0.006002724 | 0.004380856 |
| GO:0005452 | solute:inorganic anion antiporter activity | 6/1062 | 30/28366 | 0.2 | 5.341996234 | 4.692684024 | 0.000745241 | 0.007201738 | 0.00525591 |
| GO:0016903 | oxidoreductase activity, acting on the aldehyde or oxo group of donors | 9/1062 | 66/28366 | 0.136363636 | 3.642270159 | 4.238346752 | 0.000760542 | 0.007283392 | 0.005315503 |
| GO:0042626 | ATPase-coupled transmembrane transporter activity | 12/1062 | 109/28366 | 0.110091743 | 2.940548385 | 4.003273057 | 0.000794081 | 0.007536681 | 0.005500355 |
| GO:0005546 | phosphatidylinositol-4,5-bisphosphate binding | 11/1062 | 97/28366 | 0.113402062 | 3.028966937 | 3.947718064 | 0.001012619 | 0.009122151 | 0.006657449 |
| GO:0046982 | protein heterodimerization activity | 24/1062 | 319/28366 | 0.07523511 | 2.009528364 | 3.576112514 | 0.001021351 | 0.009123494 | 0.00665843 |
| GO:0004861 | cyclin-dependent protein serine/threonine kinase inhibitor activity | 4/1062 | 13/28366 | 0.307692308 | 8.218455744 | 5.13401319 | 0.001065218 | 0.00935807 | 0.006829626 |
| GO:0052650 | all-trans-retinol dehydrogenase (NADP+) activity | 4/1062 | 13/28366 | 0.307692308 | 8.218455744 | 5.13401319 | 0.001065218 | 0.00935807 | 0.006829626 |
| GO:0016004 | phospholipase activator activity | 5/1062 | 22/28366 | 0.227272727 | 6.070450265 | 4.692105977 | 0.001127208 | 0.009821492 | 0.007167836 |
| GO:0042277 | peptide binding | 23/1062 | 307/28366 | 0.074918567 | 2.001073508 | 3.478080074 | 0.001359108 | 0.011466123 | 0.008368106 |
| GO:0004467 | long-chain fatty acid-CoA ligase activity | 4/1062 | 15/28366 | 0.266666667 | 7.122661645 | 4.677806069 | 0.001915475 | 0.015662692 | 0.01143081 |
| GO:0016717 | oxidoreductase activity, acting on paired donors, with oxidation of a pair of donors resulting in the reduction of molecular oxygen to two molecules of water | 4/1062 | 15/28366 | 0.266666667 | 7.122661645 | 4.677806069 | 0.001915475 | 0.015662692 | 0.01143081 |
| GO:0015605 | organophosphate ester transmembrane transporter activity | 7/1062 | 48/28366 | 0.145833333 | 3.895205587 | 3.959216519 | 0.001946879 | 0.015678272 | 0.011442181 |
| GO:0043178 | alcohol binding | 10/1062 | 90/28366 | 0.111111111 | 2.967775685 | 3.687468957 | 0.00197398 | 0.015777 | 0.011514234 |
| GO:0005518 | collagen binding | 9/1062 | 76/28366 | 0.118421053 | 3.163024086 | 3.723850117 | 0.002100667 | 0.016540806 | 0.012071668 |
| GO:0141003 | histone H2AX kinase activity | 25/1062 | 359/28366 | 0.069637883 | 1.860026544 | 3.234195725 | 0.002352186 | 0.01811865 | 0.013223197 |
| GO:0005539 | glycosaminoglycan binding | 19/1062 | 246/28366 | 0.077235772 | 2.062966025 | 3.302324859 | 0.002430367 | 0.018450574 | 0.013465438 |
| GO:0140996 | histone H3 kinase activity | 25/1062 | 362/28366 | 0.069060773 | 1.844611959 | 3.189642371 | 0.002625692 | 0.019117193 | 0.013951945 |
| GO:0019842 | vitamin binding | 13/1062 | 142/28366 | 0.091549296 | 2.445279966 | 3.405080533 | 0.002676406 | 0.019353872 | 0.014124676 |
| GO:0140994 | RNA polymerase II CTD heptapeptide repeat modifying activity | 11/1062 | 110/28366 | 0.1 | 2.670998117 | 3.463039752 | 0.002801633 | 0.019987492 | 0.014587099 |
| GO:0004303 | estradiol 17-beta-dehydrogenase [NAD(P)+] activity | 4/1062 | 17/28366 | 0.235294118 | 6.284701451 | 4.298496338 | 0.003146327 | 0.021799675 | 0.015909651 |
| GO:0017124 | SH3 domain binding | 12/1062 | 128/28366 | 0.09375 | 2.504060734 | 3.363519161 | 0.003158184 | 0.021799675 | 0.015909651 |
| GO:0004175 | endopeptidase activity | 29/1062 | 452/28366 | 0.064159292 | 1.713693482 | 3.016549286 | 0.003684358 | 0.024945688 | 0.018205646 |
| GO:1990782 | protein tyrosine kinase binding | 12/1062 | 132/28366 | 0.090909091 | 2.428180106 | 3.243577315 | 0.004058648 | 0.026964645 | 0.019679104 |
| GO:0030145 | manganese ion binding | 8/1062 | 69/28366 | 0.115942029 | 3.096809411 | 3.439171394 | 0.004168193 | 0.027520429 | 0.020084722 |
| GO:0016887 | ATP hydrolysis activity | 26/1062 | 396/28366 | 0.065656566 | 1.753685632 | 2.978736662 | 0.004280417 | 0.028086936 | 0.020498165 |
| GO:0042805 | actinin binding | 6/1062 | 42/28366 | 0.142857143 | 3.815711595 | 3.601438777 | 0.004498797 | 0.029044208 | 0.021196793 |
| GO:0008013 | beta-catenin binding | 11/1062 | 117/28366 | 0.094017094 | 2.511194811 | 3.23037068 | 0.004508273 | 0.029044208 | 0.021196793 |
| GO:0050840 | extracellular matrix binding | 8/1062 | 70/28366 | 0.114285714 | 3.052569276 | 3.390976931 | 0.004555063 | 0.029168868 | 0.021287771 |
| GO:0001786 | phosphatidylserine binding | 8/1062 | 71/28366 | 0.112676056 | 3.009575343 | 3.343637084 | 0.004968907 | 0.031628432 | 0.023082789 |

| | | | | | | | | | |
|---|---|---|---|---|---|---|---|---|---|
| GO:0005391 | P-type sodium:potassium-exchanging transporter activity | 3/1062 | 10/28366 | 0.3 | 8.01299435 | 4.374430489 | 0.005153896 | 0.031852278 | 0.023246155 |
| GO:0016813 | hydrolase activity, acting on carbon-nitrogen (but not peptide) bonds, in linear amidines | 3/1062 | 10/28366 | 0.3 | 8.01299435 | 4.374430489 | 0.005153896 | 0.031852278 | 0.023246155 |
| GO:0045159 | myosin II binding | 3/1062 | 10/28366 | 0.3 | 8.01299435 | 4.374430489 | 0.005153896 | 0.031852278 | 0.023246155 |
| GO:0043394 | proteoglycan binding | 6/1062 | 44/28366 | 0.136363636 | 3.642270159 | 3.459251307 | 0.005683665 | 0.0341341 | 0.024911454 |
| GO:0004629 | phospholipase C activity | 4/1062 | 20/28366 | 0.2 | 5.341996234 | 3.830884546 | 0.005858432 | 0.034406148 | 0.025109998 |
| GO:0032794 | GTPase activating protein binding | 4/1062 | 20/28366 | 0.2 | 5.341996234 | 3.830884546 | 0.005858432 | 0.034406148 | 0.025109998 |
| GO:0043295 | glutathione binding | 4/1062 | 20/28366 | 0.2 | 5.341996234 | 3.830884546 | 0.005858432 | 0.034406148 | 0.025109998 |
| GO:0046875 | ephrin receptor binding | 5/1062 | 32/28366 | 0.15625 | 4.173434557 | 3.542347036 | 0.006328726 | 0.036123011 | 0.026362984 |
| GO:0090482 | vitamin transmembrane transporter activity | 5/1062 | 32/28366 | 0.15625 | 4.173434557 | 3.542347036 | 0.006328726 | 0.036123011 | 0.026362984 |
| GO:0070888 | E-box binding | 7/1062 | 59/28366 | 0.118644068 | 3.168980816 | 3.289086604 | 0.006330442 | 0.036123011 | 0.026362984 |
| GO:0001618 | virus receptor activity | 6/1062 | 45/28366 | 0.133333333 | 3.561330822 | 3.391237104 | 0.00635466 | 0.036123011 | 0.026362984 |
| GO:0030165 | PDZ domain binding | 11/1062 | 123/28366 | 0.089430894 | 2.388697503 | 3.044004653 | 0.006551144 | 0.037041839 | 0.027033555 |
| GO:0016922 | nuclear receptor binding | 13/1062 | 158/28366 | 0.082278481 | 2.197656678 | 2.977249639 | 0.00661248 | 0.037190826 | 0.027142287 |
| GO:0008307 | structural constituent of muscle | 4/1062 | 21/28366 | 0.19047619 | 5.08761546 | 3.695574523 | 0.007025471 | 0.038102427 | 0.027807584 |
| GO:0019200 | carbohydrate kinase activity | 4/1062 | 21/28366 | 0.19047619 | 5.08761546 | 3.695574523 | 0.007025471 | 0.038102427 | 0.027807584 |
| GO:0070006 | metalloaminopeptidase activity | 4/1062 | 21/28366 | 0.19047619 | 5.08761546 | 3.695574523 | 0.007025471 | 0.038102427 | 0.027807584 |
| GO:0016798 | hydrolase activity, acting on glycosyl bonds | 12/1062 | 142/28366 | 0.084507042 | 2.257181507 | 2.961920443 | 0.007226547 | 0.038833033 | 0.028340788 |
| GO:0042910 | xenobiotic transmembrane transporter activity | 5/1062 | 33/28366 | 0.151515152 | 4.046966844 | 3.453972951 | 0.007233246 | 0.038833033 | 0.028340788 |
| GO:0016209 | antioxidant activity | 9/1062 | 92/28366 | 0.097826087 | 2.61293294 | 3.05602648 | 0.007558431 | 0.040374936 | 0.029466087 |
| GO:0048306 | calcium-dependent protein binding | 9/1062 | 93/28366 | 0.096774194 | 2.584836887 | 3.019121655 | 0.008098027 | 0.042404938 | 0.030947606 |
| GO:0015662 | P-type ion transporter activity | 5/1062 | 34/28366 | 0.147058824 | 3.927938407 | 3.369017704 | 0.008224128 | 0.042602771 | 0.031091986 |
| GO:0005080 | protein kinase C binding | 7/1062 | 62/28366 | 0.112903226 | 3.015643035 | 3.133473844 | 0.008289666 | 0.042602771 | 0.031091986 |
| GO:0140103 | catalytic activity, acting on a glycoprotein | 4/1062 | 22/28366 | 0.181818182 | 4.856360212 | 3.56860828 | 0.008336196 | 0.042602771 | 0.031091986 |
| GO:0140784 | metal ion sensor activity | 4/1062 | 22/28366 | 0.181818182 | 4.856360212 | 3.56860828 | 0.008336196 | 0.042602771 | 0.031091986 |
| GO:0051219 | phosphoprotein binding | 10/1062 | 111/28366 | 0.09009009 | 2.40630461 | 2.927748375 | 0.008846573 | 0.04438019 | 0.032389167 |
| GO:0035251 | UDP-glucosyltransferase activity | 3/1062 | 12/28366 | 0.25 | 6.677495292 | 3.879544636 | 0.008934488 | 0.04438019 | 0.032389167 |
| GO:0140031 | phosphorylation-dependent protein binding | 3/1062 | 12/28366 | 0.25 | 6.677495292 | 3.879544636 | 0.008934488 | 0.04438019 | 0.032389167 |
| GO:0120020 | cholesterol transfer activity | 4/1062 | 23/28366 | 0.173913043 | 4.645214116 | 3.449090324 | 0.009797227 | 0.048116535 | 0.035115994 |
| GO:0018455 | alcohol dehydrogenase [NAD(P)+] activity | 7/1062 | 64/28366 | 0.109375 | 2.92140419 | 3.034874048 | 0.009822472 | 0.048116535 | 0.035115994 |

**Supplementary Table 10:** Summary of GO function terms associated with DEGs exhibiting >2 fold down- or upregulation in marked *Tead4* KD versus unmarked outer control cell clones at the E3.5 stage (RNA-Seq data). Relates to data in Fig. 4.

**Supplementary Table 11:**

| ID | Description | GeneRatio | BgRatio | RichFactor | FoldEnrichment | zScore | pvalue | p.adjust | qvalue |
|---|---|---|---|---|---|---|---|---|---|
| GO:0007015 | actin filament organization | 64/1052 | 474/28832 | 0.135021097 | 3.700502158 | 11.53632554 | 1.63E-19 | 8.91E-16 | 5.44E-16 |
| GO:0060070 | canonical Wnt signaling pathway | 47/1052 | 319/28832 | 0.147335423 | 4.037998975 | 10.61776502 | 3.82E-16 | 1.05E-12 | 6.38E-13 |
| GO:0022604 | regulation of cell morphogenesis | 42/1052 | 283/28832 | 0.148409894 | 4.067446829 | 10.09128364 | 1.11E-14 | 1.10E-11 | 6.73E-12 |
| GO:0042060 | wound healing | 51/1052 | 403/28832 | 0.126550868 | 3.468359924 | 9.710728686 | 1.19E-14 | 1.10E-11 | 6.73E-12 |
| GO:0010717 | regulation of epithelial to mesenchymal transition | 26/1052 | 108/28832 | 0.240740741 | 6.597943952 | 11.34199801 | 1.21E-14 | 1.10E-11 | 6.73E-12 |
| GO:0072001 | renal system development | 50/1052 | 392/28832 | 0.12755102 | 3.495770932 | 9.681765555 | 1.59E-14 | 1.24E-11 | 7.57E-12 |
| GO:0010721 | negative regulation of cell development | 44/1052 | 315/28832 | 0.13968254 | 3.828257589 | 9.821864141 | 2.38E-14 | 1.60E-11 | 9.74E-12 |
| GO:0007264 | small GTPase-mediated signal transduction | 52/1052 | 440/28832 | 0.118181818 | 3.238990667 | 9.20984308 | 1.04E-13 | 4.39E-11 | 2.68E-11 |
| GO:0050767 | regulation of neurogenesis | 54/1052 | 473/28832 | 0.114164905 | 3.12889975 | 9.084727966 | 1.44E-13 | 5.63E-11 | 3.43E-11 |
| GO:0031589 | cell-substrate adhesion | 47/1052 | 376/28832 | 0.125 | 3.425855513 | 9.21390073 | 2.08E-13 | 7.59E-11 | 4.63E-11 |
| GO:0050678 | regulation of epithelial cell proliferation | 51/1052 | 434/28832 | 0.117511521 | 3.22061993 | 9.070825719 | 2.25E-13 | 7.71E-11 | 4.70E-11 |
| GO:0031346 | positive regulation of cell projection organization | 53/1052 | 465/28832 | 0.113978495 | 3.123790834 | 8.984690398 | 2.59E-13 | 7.88E-11 | 4.81E-11 |
| GO:0032970 | regulation of actin filament-based process | 49/1052 | 416/28832 | 0.117788462 | 3.228210003 | 8.908260636 | 6.12E-13 | 1.76E-10 | 1.08E-10 |
| GO:0045859 | regulation of protein kinase activity | 53/1052 | 477/28832 | 0.111111111 | 3.045204901 | 8.765017564 | 7.13E-13 | 1.95E-10 | 1.19E-10 |
| GO:0007162 | negative regulation of cell adhesion | 42/1052 | 322/28832 | 0.130434783 | 3.574805753 | 9.041602297 | 9.63E-13 | 2.51E-10 | 1.53E-10 |
| GO:0045165 | cell fate commitment | 41/1052 | 310/28832 | 0.132258065 | 3.624776156 | 9.04179661 | 1.14E-12 | 2.83E-10 | 1.73E-10 |
| GO:0070372 | regulation of ERK1 and ERK2 cascade | 42/1052 | 333/28832 | 0.126126126 | 3.456719076 | 8.774743021 | 2.96E-12 | 7.04E-10 | 4.30E-10 |
| GO:0042063 | gliogenesis | 47/1052 | 412/28832 | 0.11407767 | 3.126508915 | 8.460097755 | 5.71E-12 | 1.28E-09 | 7.83E-10 |
| GO:0032102 | negative regulation of response to external stimulus | 48/1052 | 429/28832 | 0.111888112 | 3.06650004 | 8.391769752 | 6.82E-12 | 1.44E-09 | 8.76E-10 |
| GO:0090287 | regulation of cellular response to growth factor stimulus | 41/1052 | 336/28832 | 0.12202381 | 3.344287525 | 8.411242738 | 1.56E-11 | 3.15E-09 | 1.92E-09 |
| GO:0015711 | organic anion transport | 50/1052 | 477/28832 | 0.104821803 | 2.872834812 | 8.026300942 | 2.72E-11 | 5.32E-09 | 3.24E-09 |
| GO:0060562 | epithelial tube morphogenesis | 45/1052 | 403/28832 | 0.111662531 | 3.06031758 | 8.105456919 | 3.33E-11 | 6.28E-09 | 3.83E-09 |
| GO:0030198 | extracellular matrix organization | 40/1052 | 337/28832 | 0.118694362 | 3.253037876 | 8.096004014 | 6.48E-11 | 1.14E-08 | 6.98E-09 |
| GO:0034446 | substrate adhesion-dependent cell spreading | 21/1052 | 101/28832 | 0.207920792 | 5.698452735 | 9.204745397 | 8.45E-11 | 1.36E-08 | 8.30E-09 |
| GO:0048705 | skeletal system morphogenesis | 35/1052 | 270/28832 | 0.12962963 | 3.552739051 | 8.200977097 | 8.95E-11 | 1.40E-08 | 8.54E-09 |
| GO:0002064 | epithelial cell development | 34/1052 | 262/28832 | 0.129770992 | 3.556613357 | 8.089695621 | 1.61E-10 | 2.33E-08 | 1.42E-08 |
| GO:0043086 | negative regulation of catalytic activity | 42/1052 | 377/28832 | 0.111405836 | 3.053282367 | 7.809291965 | 1.62E-10 | 2.33E-08 | 1.42E-08 |
| GO:0048562 | embryonic organ morphogenesis | 40/1052 | 348/28832 | 0.114942529 | 3.150211966 | 7.853115956 | 1.73E-10 | 2.37E-08 | 1.45E-08 |
| GO:1902106 | negative regulation of leukocyte differentiation | 23/1052 | 126/28832 | 0.182539683 | 5.002836623 | 8.762709194 | 1.74E-10 | 2.37E-08 | 1.45E-08 |
| GO:0045936 | negative regulation of phosphate metabolic process | 42/1052 | 381/28832 | 0.11023622 | 3.021226909 | 7.728592227 | 2.25E-10 | 2.75E-08 | 1.68E-08 |
| GO:0032956 | regulation of actin cytoskeleton organization | 41/1052 | 366/28832 | 0.112021858 | 3.070165597 | 7.756289479 | 2.26E-10 | 2.75E-08 | 1.68E-08 |
| GO:0018212 | peptidyl-tyrosine modification | 36/1052 | 300/28832 | 0.12 | 3.288821293 | 7.754932232 | 4.29E-10 | 4.89E-08 | 2.98E-08 |
| GO:0001894 | tissue homeostasis | 36/1052 | 303/28832 | 0.118811881 | 3.256258706 | 7.683136287 | 5.66E-10 | 6.01E-08 | 3.67E-08 |
| GO:0060249 | anatomical structure homeostasis | 36/1052 | 303/28832 | 0.118811881 | 3.256258706 | 7.683136287 | 5.66E-10 | 6.01E-08 | 3.67E-08 |
| GO:0008544 | epidermis development | 43/1052 | 410/28832 | 0.104878049 | 2.874376333 | 7.438630818 | 6.59E-10 | 6.67E-08 | 4.07E-08 |
| GO:0046434 | organophosphate catabolic process | 33/1052 | 264/28832 | 0.125 | 3.425855513 | 7.705460125 | 7.97E-10 | 7.93E-08 | 4.84E-08 |
| GO:0043603 | amide metabolic process | 47/1052 | 479/28832 | 0.098121086 | 2.689189296 | 7.2546785 | 9.70E-10 | 9.48E-08 | 5.78E-08 |
| GO:0032535 | regulation of cellular component size | 42/1052 | 404/28832 | 0.103960396 | 2.849226368 | 7.284155505 | 1.36E-09 | 1.30E-07 | 7.94E-08 |
| GO:0003002 | regionalization | 45/1052 | 452/28832 | 0.099557522 | 2.728557488 | 7.208060979 | 1.40E-09 | 1.30E-07 | 7.94E-08 |
| GO:0030336 | negative regulation of cell migration | 37/1052 | 328/28832 | 0.112804878 | 3.091625707 | 7.413778223 | 1.40E-09 | 1.30E-07 | 7.94E-08 |
| GO:0040013 | negative regulation of locomotion | 39/1052 | 362/28832 | 0.107734807 | 2.95267105 | 7.275473137 | 1.91E-09 | 1.71E-07 | 1.05E-07 |
| GO:0061448 | connective tissue development | 37/1052 | 335/28832 | 0.110447761 | 3.027024573 | 7.261953239 | 2.51E-09 | 2.22E-07 | 1.35E-07 |
| GO:0043588 | skin development | 38/1052 | 352/28832 | 0.107954545 | 2.958693398 | 7.195142523 | 2.91E-09 | 2.48E-07 | 1.52E-07 |
| GO:0050878 | regulation of body fluid levels | 40/1052 | 384/28832 | 0.104166667 | 2.854879594 | 7.120771804 | 3.16E-09 | 2.66E-07 | 1.62E-07 |
| GO:0045216 | cell-cell junction organization | 30/1052 | 239/28832 | 0.125523013 | 3.440189637 | 7.371642564 | 4.21E-09 | 3.29E-07 | 2.01E-07 |
| GO:0003229 | ventricular cardiac muscle tissue development | 16/1052 | 71/28832 | 0.225352113 | 6.176190221 | 8.497849074 | 4.28E-09 | 3.30E-07 | 2.01E-07 |
| GO:0001666 | response to hypoxia | 37/1052 | 342/28832 | 0.108187135 | 2.96506793 | 7.114035018 | 4.42E-09 | 3.34E-07 | 2.04E-07 |
| GO:0001890 | placenta development | 27/1052 | 199/28832 | 0.135678392 | 3.718516537 | 7.488532053 | 4.60E-09 | 3.37E-07 | 2.06E-07 |
| GO:0006869 | lipid transport | 44/1052 | 456/28832 | 0.096491228 | 2.644520045 | 6.888388094 | 5.53E-09 | 3.88E-07 | 2.37E-07 |
| GO:0051336 | regulation of hydrolase activity | 42/1052 | 424/28832 | 0.099056604 | 2.714828897 | 6.92237296 | 5.72E-09 | 3.96E-07 | 2.42E-07 |
| GO:0007159 | leukocyte cell-cell adhesion | 42/1052 | 427/28832 | 0.098360656 | 2.695755158 | 6.86991383 | 7.03E-09 | 4.60E-07 | 2.81E-07 |
| GO:0007517 | muscle organ development | 40/1052 | 395/28832 | 0.101265823 | 2.775376618 | 6.91383096 | 7.07E-09 | 4.60E-07 | 2.81E-07 |
| GO:0030324 | lung development | 31/1052 | 259/28832 | 0.11969112 | 3.280355858 | 7.1737492 | 7.37E-09 | 4.69E-07 | 2.86E-07 |
| GO:0030048 | actin filament-based movement | 21/1052 | 128/28832 | 0.1640625 | 4.496435361 | 7.714910287 | 7.93E-09 | 4.92E-07 | 3.01E-07 |
| GO:0007009 | plasma membrane organization | 25/1052 | 178/28832 | 0.140449438 | 3.849275858 | 7.420334332 | 8.36E-09 | 5.03E-07 | 3.07E-07 |
| GO:0010876 | lipid localization | 45/1052 | 485/28832 | 0.092783505 | 2.542903061 | 6.668490863 | 1.23E-08 | 6.67E-07 | 4.07E-07 |
| GO:0019216 | regulation of lipid metabolic process | 39/1052 | 389/28832 | 0.100257069 | 2.747729872 | 6.75356387 | 1.43E-08 | 7.69E-07 | 4.69E-07 |
| GO:0050807 | regulation of synapse organization | 39/1052 | 393/28832 | 0.099236641 | 2.719763156 | 6.680044524 | 1.90E-08 | 9.97E-07 | 6.09E-07 |
| GO:0048814 | regulation of dendrite morphogenesis | 18/1052 | 100/28832 | 0.18 | 4.933231939 | 7.667225395 | 2.07E-08 | 1.06E-06 | 6.46E-07 |
| GO:0006631 | fatty acid metabolic process | 44/1052 | 477/28832 | 0.092243187 | 2.528094635 | 6.548867698 | 2.12E-08 | 1.07E-06 | 6.53E-07 |
| GO:0016042 | lipid catabolic process | 36/1052 | 347/28832 | 0.103746398 | 2.843361348 | 6.72262219 | 2.13E-08 | 1.07E-06 | 6.53E-07 |
| GO:0043254 | regulation of protein-containing complex assembly | 43/1052 | 461/28832 | 0.093275488 | 2.55638676 | 6.555446401 | 2.22E-08 | 1.11E-06 | 6.75E-07 |
| GO:0022409 | positive regulation of cell-cell adhesion | 35/1052 | 334/28832 | 0.104790419 | 2.871974682 | 6.696341833 | 2.59E-08 | 1.26E-06 | 7.66E-07 |
| GO:0006942 | regulation of striated muscle contraction | 17/1052 | 91/28832 | 0.186813187 | 5.119959888 | 7.660098406 | 2.86E-08 | 1.37E-06 | 8.38E-07 |
| GO:0051962 | positive regulation of nervous system development | 38/1052 | 384/28832 | 0.098958333 | 2.712135615 | 6.572786205 | 3.10E-08 | 1.46E-06 | 8.93E-07 |
| GO:0050803 | regulation of synapse structure or activity | 39/1052 | 401/28832 | 0.097256858 | 2.665503542 | 6.535717642 | 3.28E-08 | 1.52E-06 | 9.27E-07 |
| GO:0010038 | response to metal ion | 38/1052 | 386/28832 | 0.098445596 | 2.698083099 | 6.536023291 | 3.56E-08 | 1.64E-06 | 9.99E-07 |
| GO:1902903 | regulation of supramolecular fiber organization | 39/1052 | 403/28832 | 0.096774194 | 2.652275236 | 6.500185153 | 3.75E-08 | 1.71E-06 | 1.04E-06 |
| GO:0034109 | homotypic cell-cell adhesion | 17/1052 | 93/28832 | 0.182795699 | 5.009853224 | 7.537125147 | 4.01E-08 | 1.81E-06 | 1.11E-06 |
| GO:0002573 | myeloid leukocyte differentiation | 32/1052 | 295/28832 | 0.108474576 | 2.972945801 | 6.628169525 | 4.54E-08 | 2.02E-06 | 1.23E-06 |

| | | | | | | | | | |
|---|---|---|---|---|---|---|---|---|---|
| GO:0007369 | gastrulation | 26/1052 | 209/28832 | 0.124401914 | 3.40946386 | 6.803111465 | 5.39E-08 | 2.36E-06 | 1.44E-06 |
| GO:1903035 | negative regulation of response to wounding | 17/1052 | 96/28832 | 0.177083333 | 4.853295311 | 7.359127973 | 6.54E-08 | 2.80E-06 | 1.71E-06 |
| GO:0050866 | negative regulation of cell activation | 27/1052 | 226/28832 | 0.119469027 | 3.274268986 | 6.679425091 | 7.03E-08 | 2.91E-06 | 1.78E-06 |
| GO:0048010 | vascular endothelial growth factor receptor signaling pathway | 17/1052 | 97/28832 | 0.175257732 | 4.803261338 | 7.301431873 | 7.66E-08 | 3.13E-06 | 1.91E-06 |
| GO:0048638 | regulation of developmental growth | 38/1052 | 398/28832 | 0.095477387 | 2.616733859 | 6.320227229 | 7.94E-08 | 3.22E-06 | 1.96E-06 |
| GO:0051961 | negative regulation of nervous system development | 23/1052 | 173/28832 | 0.132947977 | 3.643684477 | 6.786936073 | 9.19E-08 | 3.62E-06 | 2.21E-06 |
| GO:1901292 | nucleoside phosphate catabolic process | 25/1052 | 201/28832 | 0.124378109 | 3.408811456 | 6.668899619 | 9.70E-08 | 3.79E-06 | 2.31E-06 |
| GO:0141091 | transforming growth factor beta receptor superfamily signaling pathway | 36/1052 | 370/28832 | 0.097297297 | 2.666611859 | 6.278764945 | 1.09E-07 | 4.16E-06 | 2.54E-06 |
| GO:0007178 | cell surface receptor protein serine/threonine kinase signaling pathway | 38/1052 | 404/28832 | 0.094059406 | 2.577871475 | 6.21528083 | 1.17E-07 | 4.41E-06 | 2.69E-06 |
| GO:0019827 | stem cell population maintenance | 26/1052 | 219/28832 | 0.118721461 | 3.253780579 | 6.51513763 | 1.39E-07 | 5.06E-06 | 3.09E-06 |
| GO:0006936 | muscle contraction | 33/1052 | 330/28832 | 0.1 | 2.740684411 | 6.188864071 | 1.94E-07 | 6.89E-06 | 4.20E-06 |
| GO:0045926 | negative regulation of growth | 28/1052 | 253/28832 | 0.110671937 | 3.033168518 | 6.32094083 | 2.05E-07 | 7.23E-06 | 4.41E-06 |
| GO:0032370 | positive regulation of lipid transport | 17/1052 | 104/28832 | 0.163461538 | 4.479964902 | 6.918476015 | 2.19E-07 | 7.59E-06 | 4.63E-06 |
| GO:2000241 | regulation of reproductive process | 26/1052 | 225/28832 | 0.115555556 | 3.167013097 | 6.35021222 | 2.37E-07 | 8.17E-06 | 4.98E-06 |
| GO:0098727 | maintenance of cell number | 26/1052 | 227/28832 | 0.114537445 | 3.139109898 | 6.296462947 | 2.83E-07 | 9.54E-06 | 5.82E-06 |
| GO:0023019 | signal transduction involved in regulation of gene expression | 9/1052 | 27/28832 | 0.333333333 | 9.135614702 | 8.230191056 | 2.87E-07 | 9.63E-06 | 5.88E-06 |
| GO:0044282 | small molecule catabolic process | 35/1052 | 369/28832 | 0.094850949 | 2.599565159 | 6.017919652 | 3.00E-07 | 1.00E-05 | 6.11E-06 |
| GO:0007163 | establishment or maintenance of cell polarity | 27/1052 | 243/28832 | 0.111111111 | 3.045204901 | 6.230344791 | 3.11E-07 | 1.03E-05 | 6.29E-06 |
| GO:0035924 | cellular response to vascular endothelial growth factor stimulus | 17/1052 | 107/28832 | 0.158878505 | 4.354358409 | 6.764612707 | 3.34E-07 | 1.10E-05 | 6.72E-06 |
| GO:0006935 | chemotaxis | 41/1052 | 475/28832 | 0.086315789 | 2.365643386 | 5.840171104 | 3.81E-07 | 1.23E-05 | 7.49E-06 |
| GO:0050863 | regulation of T cell activation | 36/1052 | 391/28832 | 0.092071611 | 2.523392296 | 5.902001355 | 4.20E-07 | 1.34E-05 | 8.15E-06 |
| GO:0050900 | leukocyte migration | 37/1052 | 408/28832 | 0.090686275 | 2.485424588 | 5.880438266 | 4.23E-07 | 1.34E-05 | 8.15E-06 |
| GO:0045664 | regulation of neuron differentiation | 27/1052 | 247/28832 | 0.109311741 | 2.995889842 | 6.130382155 | 4.31E-07 | 1.35E-05 | 8.22E-06 |
| GO:0043583 | ear development | 29/1052 | 279/28832 | 0.103942652 | 2.848740068 | 6.03842512 | 4.73E-07 | 1.44E-05 | 8.82E-06 |
| GO:0072089 | stem cell proliferation | 20/1052 | 149/28832 | 0.134228188 | 3.678771021 | 6.37952057 | 5.34E-07 | 1.59E-05 | 9.72E-06 |
| GO:0061383 | trabecula morphogenesis | 12/1052 | 55/28832 | 0.218181818 | 5.979675078 | 7.193350047 | 5.39E-07 | 1.59E-05 | 9.72E-06 |
| GO:0099563 | modification of synaptic structure | 12/1052 | 55/28832 | 0.218181818 | 5.979675078 | 7.193350047 | 5.39E-07 | 1.59E-05 | 9.72E-06 |
| GO:0038084 | vascular endothelial growth factor signaling pathway | 15/1052 | 88/28832 | 0.170454545 | 4.671621155 | 6.712699402 | 6.43E-07 | 1.83E-05 | 1.12E-05 |
| GO:0030217 | T cell differentiation | 33/1052 | 348/28832 | 0.094827586 | 2.598924872 | 5.839676564 | 6.53E-07 | 1.85E-05 | 1.13E-05 |
| GO:0062012 | regulation of small molecule metabolic process | 36/1052 | 401/28832 | 0.089775561 | 2.460464808 | 5.731110965 | 7.68E-07 | 2.08E-05 | 1.27E-05 |
| GO:0010770 | positive regulation of cell morphogenesis | 13/1052 | 67/28832 | 0.194029851 | 5.317745871 | 6.885474782 | 7.69E-07 | 2.08E-05 | 1.27E-05 |
| GO:0006790 | sulfur compound metabolic process | 33/1052 | 351/28832 | 0.094017094 | 2.576711839 | 5.783621616 | 7.92E-07 | 2.13E-05 | 1.30E-05 |
| GO:1901136 | carbohydrate derivative catabolic process | 28/1052 | 271/28832 | 0.103321033 | 2.83170345 | 5.895555153 | 8.36E-07 | 2.22E-05 | 1.36E-05 |
| GO:1905954 | positive regulation of lipid localization | 18/1052 | 127/28832 | 0.141732283 | 3.884434598 | 6.339505504 | 8.71E-07 | 2.30E-05 | 1.41E-05 |
| GO:0006066 | alcohol metabolic process | 34/1052 | 373/28832 | 0.091152815 | 2.498210991 | 5.667469337 | 1.10E-06 | 2.82E-05 | 1.72E-05 |
| GO:0044344 | cellular response to fibroblast growth factor stimulus | 20/1052 | 156/28832 | 0.128205128 | 3.513697962 | 6.126151161 | 1.12E-06 | 2.86E-05 | 1.75E-05 |
| GO:0050868 | negative regulation of T cell activation | 19/1052 | 143/28832 | 0.132867133 | 3.641468797 | 6.16208097 | 1.19E-06 | 3.02E-05 | 1.84E-05 |
| GO:0051495 | positive regulation of cytoskeleton organization | 23/1052 | 201/28832 | 0.114427861 | 3.13610654 | 5.913904159 | 1.37E-06 | 3.39E-05 | 2.07E-05 |
| GO:0061053 | somite development | 15/1052 | 94/28832 | 0.159574468 | 4.37343257 | 6.374986839 | 1.54E-06 | 3.75E-05 | 2.29E-05 |
| GO:0009132 | nucleoside diphosphate metabolic process | 19/1052 | 146/28832 | 0.130136986 | 3.566644096 | 6.050324729 | 1.64E-06 | 3.98E-05 | 2.43E-05 |
| GO:0031099 | regeneration | 25/1052 | 236/28832 | 0.105932203 | 2.903267384 | 5.713138662 | 2.03E-06 | 4.72E-05 | 2.88E-05 |
| GO:0050920 | regulation of chemotaxis | 25/1052 | 236/28832 | 0.105932203 | 2.903267384 | 5.713138662 | 2.03E-06 | 4.72E-05 | 2.88E-05 |
| GO:0001763 | morphogenesis of a branching structure | 26/1052 | 252/28832 | 0.103174603 | 2.827690265 | 5.670787313 | 2.09E-06 | 4.85E-05 | 2.96E-05 |
| GO:0071774 | response to fibroblast growth factor | 20/1052 | 163/28832 | 0.122699387 | 3.362802958 | 5.886899721 | 2.24E-06 | 5.09E-05 | 3.11E-05 |
| GO:0016052 | carbohydrate catabolic process | 22/1052 | 192/28832 | 0.114583333 | 3.140367554 | 5.790606701 | 2.26E-06 | 5.11E-05 | 3.12E-05 |
| GO:0050890 | cognition | 35/1052 | 403/28832 | 0.086848635 | 2.380247007 | 5.430003976 | 2.32E-06 | 5.18E-05 | 3.16E-05 |
| GO:0051345 | positive regulation of hydrolase activity | 24/1052 | 223/28832 | 0.107623318 | 2.949615509 | 5.687503356 | 2.44E-06 | 5.41E-05 | 3.30E-05 |
| GO:0001570 | vasculogenesis | 15/1052 | 99/28832 | 0.151515152 | 4.152552137 | 6.114500585 | 3.00E-06 | 6.57E-05 | 4.01E-05 |
| GO:0032868 | response to insulin | 32/1052 | 356/28832 | 0.08988764 | 2.463536549 | 5.407053686 | 3.03E-06 | 6.57E-05 | 4.01E-05 |
| GO:0048545 | response to steroid hormone | 32/1052 | 356/28832 | 0.08988764 | 2.463536549 | 5.407053686 | 3.03E-06 | 6.57E-05 | 4.01E-05 |
| GO:0051258 | protein polymerization | 28/1052 | 290/28832 | 0.096551724 | 2.646178052 | 5.482835389 | 3.18E-06 | 6.85E-05 | 4.18E-05 |
| GO:0035265 | organ growth | 24/1052 | 227/28832 | 0.105726872 | 2.897639906 | 5.58569682 | 3.34E-06 | 7.16E-05 | 4.37E-05 |
| GO:0072523 | purine-containing compound catabolic process | 21/1052 | 182/28832 | 0.115384615 | 3.162328166 | 5.694644119 | 3.42E-06 | 7.27E-05 | 4.44E-05 |
| GO:0001756 | somitogenesis | 13/1052 | 76/28832 | 0.171052632 | 4.688012808 | 6.26479092 | 3.42E-06 | 7.27E-05 | 4.44E-05 |
| GO:0007626 | locomotory behavior | 28/1052 | 292/28832 | 0.095890411 | 2.628053544 | 5.44132577 | 3.63E-06 | 7.67E-05 | 4.68E-05 |
| GO:1900744 | regulation of p38MAPK cascade | 10/1052 | 45/28832 | 0.222222222 | 6.090409801 | 6.650164706 | 4.00E-06 | 8.38E-05 | 5.11E-05 |
| GO:0098742 | cell-cell adhesion via plasma-membrane adhesion molecules | 24/1052 | 232/28832 | 0.103448276 | 2.83519077 | 5.461523345 | 4.87E-06 | 9.84E-05 | 6.01E-05 |
| GO:0017145 | stem cell division | 10/1052 | 46/28832 | 0.217391304 | 5.958009588 | 6.548882824 | 4.94E-06 | 9.94E-05 | 6.06E-05 |
| GO:0150105 | protein localization to cell-cell junction | 8/1052 | 28/28832 | 0.285714286 | 7.830526888 | 7.036852082 | 4.96E-06 | 9.94E-05 | 6.06E-05 |
| GO:0006643 | membrane lipid metabolic process | 22/1052 | 202/28832 | 0.108910891 | 2.984903814 | 5.509041946 | 5.22E-06 | 0.000103921 | 6.34E-05 |
| GO:0055074 | calcium ion homeostasis | 32/1052 | 366/28832 | 0.087431694 | 2.396226807 | 5.231243004 | 5.40E-06 | 0.000106736 | 6.51E-05 |
| GO:0033674 | positive regulation of kinase activity | 28/1052 | 300/28832 | 0.093333333 | 2.557972117 | 5.27868568 | 6.09E-06 | 0.000118066 | 7.21E-05 |
| GO:0030100 | regulation of endocytosis | 31/1052 | 352/28832 | 0.088068182 | 2.41367093 | 5.193035231 | 6.51E-06 | 0.000125016 | 7.63E-05 |
| GO:0008016 | regulation of heart contraction | 22/1052 | 205/28832 | 0.107317073 | 2.941222294 | 5.427950572 | 6.63E-06 | 0.000126906 | 7.75E-05 |
| GO:0050870 | positive regulation of T cell activation | 25/1052 | 253/28832 | 0.098814229 | 2.708186177 | 5.310599517 | 7.03E-06 | 0.000134017 | 8.18E-05 |
| GO:0045765 | regulation of angiogenesis | 29/1052 | 320/28832 | 0.090625 | 2.483745247 | 5.193882148 | 7.41E-06 | 0.000140352 | 8.57E-05 |
| GO:0050905 | neuromuscular process | 23/1052 | 222/28832 | 0.103603604 | 2.839447813 | 5.353978607 | 7.44E-06 | 0.000140352 | 8.57E-05 |
| GO:0048771 | tissue remodeling | 23/1052 | 223/28832 | 0.103139013 | 2.826714863 | 5.328972233 | 8.01E-06 | 0.000150154 | 9.16E-05 |
| GO:0007548 | sex differentiation | 32/1052 | 375/28832 | 0.085333333 | 2.338717364 | 5.07786997 | 8.90E-06 | 0.000162878 | 9.94E-05 |
| GO:0043542 | endothelial cell migration | 24/1052 | 241/28832 | 0.099585062 | 2.729312276 | 5.24612791 | 9.34E-06 | 0.000168654 | 0.00010293 |
| GO:0098773 | skin epidermis development | 19/1052 | 167/28832 | 0.113772455 | 3.11814394 | 5.342067046 | 1.21E-05 | 0.000212798 | 0.00012987 |
| GO:0150104 | transport across blood-brain barrier | 7/1052 | 23/28832 | 0.304347826 | 8.341213424 | 6.853923907 | 1.24E-05 | 0.000217886 | 0.000132976 |
| GO:0050819 | negative regulation of coagulation | 10/1052 | 51/28832 | 0.196078431 | 5.373891001 | 6.083754866 | 1.31E-05 | 0.000230165 | 0.000140469 |

| | | | | | | | | | |
|---|---|---|---|---|---|---|---|---|---|
| GO:0050867 | positive regulation of cell activation | 33/1052 | 401/28832 | 0.082294264 | 2.255426074 | 4.926504287 | 1.37E-05 | 0.000237095 | 0.000144699 |
| GO:0034103 | regulation of tissue remodeling | 13/1052 | 86/28832 | 0.151162791 | 4.142895039 | 5.680178645 | 1.39E-05 | 0.000237095 | 0.000144699 |
| GO:1904888 | cranial skeletal system development | 13/1052 | 86/28832 | 0.151162791 | 4.142895039 | 5.680178645 | 1.39E-05 | 0.000237095 | 0.000144699 |
| GO:0048144 | fibroblast proliferation | 18/1052 | 154/28832 | 0.116883117 | 3.203397363 | 5.335195742 | 1.40E-05 | 0.000237692 | 0.000145063 |
| GO:0060047 | heart contraction | 24/1052 | 247/28832 | 0.097165992 | 2.663013193 | 5.107950442 | 1.41E-05 | 0.000237692 | 0.000145063 |
| GO:0055117 | regulation of cardiac muscle contraction | 12/1052 | 74/28832 | 0.162162162 | 4.444353098 | 5.773189183 | 1.43E-05 | 0.000239195 | 0.000145981 |
| GO:0036005 | response to macrophage colony-stimulating factor | 11/1052 | 63/28832 | 0.174603175 | 4.785321987 | 5.853051174 | 1.57E-05 | 0.000257256 | 0.000157003 |
| GO:0038066 | p38MAPK cascade | 11/1052 | 63/28832 | 0.174603175 | 4.785321987 | 5.853051174 | 1.57E-05 | 0.000257256 | 0.000157003 |
| GO:0006644 | phospholipid metabolic process | 31/1052 | 368/28832 | 0.08423913 | 2.308728715 | 4.91696569 | 1.58E-05 | 0.000257275 | 0.000157015 |
| GO:0048145 | regulation of fibroblast proliferation | 16/1052 | 127/28832 | 0.125984252 | 3.452830754 | 5.390912339 | 1.62E-05 | 0.000261844 | 0.000159803 |
| GO:1905330 | regulation of morphogenesis of an epithelium | 12/1052 | 75/28832 | 0.16 | 4.385095057 | 5.712172497 | 1.64E-05 | 0.00026483 | 0.000161626 |
| GO:0016053 | organic acid biosynthetic process | 29/1052 | 335/28832 | 0.086567164 | 2.372532773 | 4.917191904 | 1.77E-05 | 0.000282525 | 0.000172425 |
| GO:2000027 | regulation of animal organ morphogenesis | 14/1052 | 101/28832 | 0.138613861 | 3.79896849 | 5.483463127 | 1.82E-05 | 0.000290054 | 0.00017702 |
| GO:0007589 | body fluid secretion | 16/1052 | 129/28832 | 0.124031008 | 3.399298494 | 5.314802043 | 1.97E-05 | 0.000309127 | 0.00018866 |
| GO:0030900 | forebrain development | 37/1052 | 482/28832 | 0.076763485 | 2.10384488 | 4.75583345 | 2.00E-05 | 0.000311098 | 0.000189863 |
| GO:0031345 | negative regulation of cell projection organization | 22/1052 | 220/28832 | 0.1 | 2.740684411 | 5.043463409 | 2.03E-05 | 0.00031457 | 0.000191982 |
| GO:0141176 | gene silencing by piRNA-directed DNA methylation | 6/1052 | 17/28832 | 0.352941176 | 9.673003802 | 6.960754034 | 2.04E-05 | 0.00031457 | 0.000191982 |
| GO:0044843 | cell cycle G1/S phase transition | 24/1052 | 253/28832 | 0.09486166 | 2.599858729 | 4.97381908 | 2.10E-05 | 0.000322785 | 0.000196996 |
| GO:0008344 | adult locomotory behavior | 15/1052 | 116/28832 | 0.129310345 | 3.543988462 | 5.342613557 | 2.16E-05 | 0.000328862 | 0.000200704 |
| GO:0038145 | macrophage colony-stimulating factor signaling pathway | 10/1052 | 54/28832 | 0.185185185 | 5.075341501 | 5.833137115 | 2.23E-05 | 0.000338358 | 0.0002065 |
| GO:2000736 | regulation of stem cell differentiation | 13/1052 | 90/28832 | 0.144444444 | 3.958766371 | 5.470726939 | 2.30E-05 | 0.000346095 | 0.000211222 |
| GO:0045862 | positive regulation of proteolysis | 22/1052 | 222/28832 | 0.099099099 | 2.71599356 | 4.994647169 | 2.34E-05 | 0.000350688 | 0.000214025 |
| GO:1990845 | adaptive thermogenesis | 19/1052 | 175/28832 | 0.108571429 | 2.975600217 | 5.101223245 | 2.35E-05 | 0.000350761 | 0.000214069 |
| GO:0051098 | regulation of binding | 29/1052 | 342/28832 | 0.084795322 | 2.323972161 | 4.793108764 | 2.59E-05 | 0.000375886 | 0.000229403 |
| GO:0032869 | cellular response to insulin stimulus | 26/1052 | 290/28832 | 0.089655172 | 2.457165334 | 4.853301046 | 2.60E-05 | 0.000375886 | 0.000229403 |
| GO:0006814 | sodium ion transport | 24/1052 | 258/28832 | 0.093023256 | 2.54947387 | 4.864970285 | 2.89E-05 | 0.000409385 | 0.000249847 |
| GO:0007492 | endoderm development | 13/1052 | 92/28832 | 0.141304348 | 3.872706232 | 5.370483055 | 2.92E-05 | 0.000411246 | 0.000250983 |
| GO:0140966 | piRNA-mediated heterochromatin formation | 6/1052 | 18/28832 | 0.333333333 | 9.135614702 | 6.718873297 | 2.96E-05 | 0.000414345 | 0.000252875 |
| GO:0030162 | regulation of proteolysis | 36/1052 | 473/28832 | 0.076109937 | 2.085933167 | 4.634040381 | 3.08E-05 | 0.000428364 | 0.00026143 |
| GO:0007605 | sensory perception of sound | 26/1052 | 293/28832 | 0.088737201 | 2.432006644 | 4.794364729 | 3.10E-05 | 0.000429545 | 0.000262151 |
| GO:0048872 | homeostasis of number of cells | 34/1052 | 436/28832 | 0.077981651 | 2.137230962 | 4.656235509 | 3.11E-05 | 0.00042964 | 0.000262209 |
| GO:0001649 | osteoblast differentiation | 23/1052 | 243/28832 | 0.094650206 | 2.594063434 | 4.856024357 | 3.21E-05 | 0.000441843 | 0.000269656 |
| GO:0048511 | rhythmic process | 30/1052 | 364/28832 | 0.082417582 | 2.258805833 | 4.703298712 | 3.24E-05 | 0.000443844 | 0.000270878 |
| GO:0003158 | endothelium development | 17/1052 | 149/28832 | 0.11409396 | 3.126955368 | 5.065365951 | 3.29E-05 | 0.000448719 | 0.000273853 |
| GO:0007411 | axon guidance | 24/1052 | 261/28832 | 0.091954023 | 2.520169573 | 4.80088354 | 3.49E-05 | 0.000472269 | 0.000288225 |
| GO:0072091 | regulation of stem cell proliferation | 14/1052 | 107/28832 | 0.130841121 | 3.58594222 | 5.214975701 | 3.52E-05 | 0.00047576 | 0.000290356 |
| GO:0051347 | positive regulation of transferase activity | 29/1052 | 348/28832 | 0.083333333 | 2.283903676 | 4.689139768 | 3.56E-05 | 0.00047812 | 0.000291796 |
| GO:0007596 | blood coagulation | 20/1052 | 197/28832 | 0.101522843 | 2.782420721 | 4.885019976 | 3.84E-05 | 0.000499789 | 0.000305021 |
| GO:0042632 | cholesterol homeostasis | 14/1052 | 109/28832 | 0.128440367 | 3.520145115 | 5.129741743 | 4.34E-05 | 0.000558717 | 0.000340984 |
| GO:0014855 | striated muscle cell proliferation | 13/1052 | 96/28832 | 0.135416667 | 3.711343473 | 5.178197357 | 4.61E-05 | 0.000588092 | 0.000358912 |
| GO:0050817 | coagulation | 20/1052 | 200/28832 | 0.1 | 2.740684411 | 4.807073884 | 4.76E-05 | 0.000605298 | 0.000369413 |
| GO:0035335 | peptidyl-tyrosine dephosphorylation | 7/1052 | 28/28832 | 0.25 | 6.851711027 | 6.028469779 | 5.10E-05 | 0.000637701 | 0.000389189 |
| GO:0098901 | regulation of cardiac muscle cell action potential | 7/1052 | 28/28832 | 0.25 | 6.851711027 | 6.028469779 | 5.10E-05 | 0.000637701 | 0.000389189 |
| GO:0003143 | embryonic heart tube morphogenesis | 11/1052 | 72/28832 | 0.152777778 | 4.187156738 | 5.269226065 | 5.70E-05 | 0.000698199 | 0.00042611 |
| GO:2000179 | positive regulation of neural precursor cell proliferation | 11/1052 | 72/28832 | 0.152777778 | 4.187156738 | 5.269226065 | 5.70E-05 | 0.000698199 | 0.00042611 |
| GO:0022405 | hair cycle process | 15/1052 | 126/28832 | 0.119047619 | 3.262719536 | 4.953375607 | 5.75E-05 | 0.000698199 | 0.00042611 |
| GO:0031529 | ruffle organization | 10/1052 | 60/28832 | 0.166666667 | 4.567807351 | 5.383486119 | 5.76E-05 | 0.000698199 | 0.00042611 |
| GO:0030195 | negative regulation of blood coagulation | 9/1052 | 49/28832 | 0.183673469 | 5.033910142 | 5.49955859 | 6.11E-05 | 0.000734969 | 0.000448551 |
| GO:0046598 | positive regulation of viral entry into host cell | 5/1052 | 13/28832 | 0.384615385 | 10.54109389 | 6.69579262 | 6.45E-05 | 0.000765976 | 0.000467475 |
| GO:0075294 | positive regulation by symbiont of entry into host | 5/1052 | 13/28832 | 0.384615385 | 10.54109389 | 6.69579262 | 6.45E-05 | 0.000765976 | 0.000467475 |
| GO:0006096 | glycolytic process | 14/1052 | 113/28832 | 0.123893805 | 3.395538208 | 4.965114512 | 6.49E-05 | 0.000767805 | 0.000468591 |
| GO:0060491 | regulation of cell projection assembly | 21/1052 | 221/28832 | 0.095022624 | 2.604270254 | 4.658850255 | 6.57E-05 | 0.000774631 | 0.000472757 |
| GO:0034113 | heterotypic cell-cell adhesion | 10/1052 | 61/28832 | 0.163934426 | 4.492925263 | 5.314327744 | 6.66E-05 | 0.000778698 | 0.000475239 |
| GO:0034204 | lipid translocation | 10/1052 | 61/28832 | 0.163934426 | 4.492925263 | 5.314327744 | 6.66E-05 | 0.000778698 | 0.000475239 |
| GO:0009636 | response to toxic substance | 22/1052 | 238/28832 | 0.092436975 | 2.533405758 | 4.622529741 | 6.73E-05 | 0.000784671 | 0.000478884 |
| GO:0070661 | leukocyte proliferation | 30/1052 | 379/28832 | 0.079155673 | 2.169407185 | 4.459569466 | 6.80E-05 | 0.000791124 | 0.000482822 |
| GO:0120192 | tight junction assembly | 13/1052 | 100/28832 | 0.13 | 3.562889734 | 4.995955885 | 7.10E-05 | 0.000823466 | 0.000502561 |
| GO:0006979 | response to oxidative stress | 33/1052 | 437/28832 | 0.075514874 | 2.069624383 | 4.384526294 | 7.53E-05 | 0.000867828 | 0.000529635 |
| GO:1903532 | positive regulation of secretion by cell | 31/1052 | 400/28832 | 0.0775 | 2.124030418 | 4.405304745 | 7.66E-05 | 0.000878824 | 0.000536346 |
| GO:0042698 | ovulation cycle | 12/1052 | 88/28832 | 0.136363636 | 3.737296924 | 5.00450636 | 8.36E-05 | 0.000934349 | 0.000570232 |
| GO:0006090 | pyruvate metabolic process | 16/1052 | 147/28832 | 0.108843537 | 2.983057862 | 4.690706166 | 9.80E-05 | 0.00108043 | 0.000659386 |
| GO:0036215 | response to stem cell factor | 9/1052 | 52/28832 | 0.173076923 | 4.743492249 | 5.257809146 | 9.93E-05 | 0.001081691 | 0.000660155 |
| GO:0036216 | cellular response to stem cell factor stimulus | 9/1052 | 52/28832 | 0.173076923 | 4.743492249 | 5.257809146 | 9.93E-05 | 0.001081691 | 0.000660155 |
| GO:0001778 | plasma membrane repair | 7/1052 | 31/28832 | 0.225806452 | 6.188642218 | 5.624739201 | 0.000102924 | 0.001108461 | 0.000676493 |
| GO:0030879 | mammary gland development | 18/1052 | 179/28832 | 0.100558659 | 2.755995497 | 4.586025505 | 0.000103947 | 0.001117282 | 0.000681876 |
| GO:0050727 | regulation of inflammatory response | 31/1052 | 407/28832 | 0.076167076 | 2.087499183 | 4.299792614 | 0.000105135 | 0.001127828 | 0.000688313 |
| GO:0001824 | blastocyst development | 17/1052 | 164/28832 | 0.103658537 | 2.840953352 | 4.600845108 | 0.000110429 | 0.001179995 | 0.00072015 |
| GO:0044403 | biological process involved in symbiotic interaction | 18/1052 | 180/28832 | 0.1 | 2.740684411 | 4.55879878 | 0.000111633 | 0.001190532 | 0.000726581 |
| GO:0009914 | hormone transport | 32/1052 | 428/28832 | 0.074766355 | 2.04910984 | 4.255246225 | 0.000115596 | 0.001226446 | 0.000748499 |
| GO:0060021 | roof of mouth development | 13/1052 | 105/28832 | 0.123809524 | 3.393228318 | 4.780851428 | 0.00011799 | 0.001239652 | 0.000756558 |
| GO:0070664 | negative regulation of leukocyte proliferation | 13/1052 | 105/28832 | 0.123809524 | 3.393228318 | 4.780851428 | 0.00011799 | 0.001239652 | 0.000756558 |
| GO:0044409 | symbiont entry into host | 12/1052 | 92/28832 | 0.130434783 | 3.574805753 | 4.813562386 | 0.000129246 | 0.001346862 | 0.000821989 |
| GO:0045682 | regulation of epidermis development | 10/1052 | 66/28832 | 0.151515152 | 4.152552137 | 4.98960435 | 0.00013221 | 0.001364757 | 0.00083291 |

| | | | | | | | | | |
|---|---|---|---|---|---|---|---|---|---|
| GO:0060688 | regulation of morphogenesis of a branching structure | 10/1052 | 66/28832 | 0.151515152 | 4.152552137 | 4.98960435 | 0.00013221 | 0.001364757 | 0.00083291 |
| GO:0097035 | regulation of membrane lipid distribution | 10/1052 | 66/28832 | 0.151515152 | 4.152552137 | 4.98960435 | 0.00013221 | 0.001364757 | 0.00083291 |
| GO:0120161 | regulation of cold-induced thermogenesis | 16/1052 | 151/28832 | 0.105960265 | 2.904036462 | 4.564972776 | 0.000134546 | 0.001385903 | 0.000845816 |
| GO:0055062 | phosphate ion homeostasis | 6/1052 | 23/28832 | 0.260869565 | 7.149611506 | 5.741417241 | 0.000137699 | 0.001402567 | 0.000855985 |
| GO:0070254 | mucus secretion | 6/1052 | 23/28832 | 0.260869565 | 7.149611506 | 5.741417241 | 0.000137699 | 0.001402567 | 0.000855985 |
| GO:0051147 | regulation of muscle cell differentiation | 17/1052 | 167/28832 | 0.101796407 | 2.789918262 | 4.514265147 | 0.000137924 | 0.001402567 | 0.000855985 |
| GO:0005996 | monosaccharide metabolic process | 25/1052 | 303/28832 | 0.082508251 | 2.261290768 | 4.295016721 | 0.000140079 | 0.001421843 | 0.00086775 |
| GO:0006575 | modified amino acid metabolic process | 19/1052 | 202/28832 | 0.094059406 | 2.577871475 | 4.379335706 | 0.000163989 | 0.001604977 | 0.000979516 |
| GO:0042552 | myelination | 19/1052 | 202/28832 | 0.094059406 | 2.577871475 | 4.379335706 | 0.000163989 | 0.001604977 | 0.000979516 |
| GO:0006909 | phagocytosis | 21/1052 | 236/28832 | 0.088983051 | 2.438744603 | 4.318755989 | 0.000166462 | 0.001622067 | 0.000989946 |
| GO:0061351 | neural precursor cell proliferation | 20/1052 | 219/28832 | 0.091324201 | 2.502908138 | 4.344545983 | 0.000166624 | 0.001622067 | 0.000989946 |
| GO:0022411 | cellular component disassembly | 32/1052 | 437/28832 | 0.073226545 | 2.006908493 | 4.127445888 | 0.000168893 | 0.001638322 | 0.000999867 |
| GO:0141005 | transposable element silencing by heterochromatin formation | 6/1052 | 24/28832 | 0.25 | 6.851711027 | 5.580890998 | 0.000177954 | 0.001702074 | 0.001038775 |
| GO:0071248 | cellular response to metal ion | 19/1052 | 204/28832 | 0.093137255 | 2.552598226 | 4.330621894 | 0.000186361 | 0.001770104 | 0.001080293 |
| GO:0045913 | positive regulation of carbohydrate metabolic process | 13/1052 | 110/28832 | 0.118181818 | 3.238990667 | 4.578391106 | 0.000189572 | 0.00178417 | 0.001088878 |
| GO:0035050 | embryonic heart tube development | 12/1052 | 96/28832 | 0.125 | 3.425855513 | 4.632964703 | 0.000194788 | 0.001818571 | 0.001109873 |
| GO:1903825 | organic acid transmembrane transport | 16/1052 | 156/28832 | 0.102564103 | 2.81095837 | 4.413499508 | 0.000196639 | 0.001832727 | 0.001118512 |
| GO:0141196 | transposable element silencing by piRNA-mediated DNA methylation | 5/1052 | 16/28832 | 0.3125 | 8.564638783 | 5.889832455 | 0.00019991 | 0.001844418 | 0.001125647 |
| GO:0050777 | negative regulation of immune response | 20/1052 | 222/28832 | 0.09009009 | 2.469085055 | 4.275984292 | 0.000199916 | 0.001844418 | 0.001125647 |
| GO:0032941 | secretion by tissue | 9/1052 | 57/28832 | 0.157894737 | 4.327396438 | 4.893345845 | 0.000206219 | 0.001891321 | 0.001154272 |
| GO:1902806 | regulation of cell cycle G1/S phase transition | 18/1052 | 189/28832 | 0.095238095 | 2.610175629 | 4.321818278 | 0.000206566 | 0.001891321 | 0.001154272 |
| GO:0008202 | steroid metabolic process | 28/1052 | 366/28832 | 0.076502732 | 2.096698456 | 4.109000126 | 0.000206728 | 0.001891321 | 0.001154272 |
| GO:0021700 | developmental maturation | 31/1052 | 423/28832 | 0.073286052 | 2.008539403 | 4.066368036 | 0.000208949 | 0.001908452 | 0.001164727 |
| GO:0032355 | response to estradiol | 14/1052 | 126/28832 | 0.111111111 | 3.045204901 | 4.477208909 | 0.000210645 | 0.001917532 | 0.001170269 |
| GO:0031644 | regulation of nervous system process | 17/1052 | 173/28832 | 0.098265896 | 2.693158091 | 4.346719826 | 0.00021135 | 0.001917709 | 0.001170376 |
| GO:0046631 | alpha-beta T cell activation | 19/1052 | 206/28832 | 0.09223301 | 2.527815719 | 4.282485023 | 0.000211365 | 0.001917709 | 0.001170376 |
| GO:0000082 | G1/S transition of mitotic cell cycle | 20/1052 | 223/28832 | 0.089686099 | 2.458012924 | 4.253378864 | 0.000212243 | 0.001922484 | 0.00117329 |
| GO:0002040 | sprouting angiogenesis | 15/1052 | 142/28832 | 0.105633803 | 2.895089166 | 4.40534727 | 0.000222933 | 0.002006027 | 0.001224277 |
| GO:0048730 | epidermis morphogenesis | 8/1052 | 46/28832 | 0.173913043 | 4.766407671 | 4.974932393 | 0.000232606 | 0.002072617 | 0.001264917 |
| GO:0010565 | regulation of ketone metabolic process | 16/1052 | 159/28832 | 0.100628931 | 2.75792142 | 4.325467517 | 0.000244767 | 0.00216665 | 0.001322305 |
| GO:0061371 | determination of heart left/right asymmetry | 10/1052 | 71/28832 | 0.14084507 | 3.860118888 | 4.695511101 | 0.000245535 | 0.00216665 | 0.001322305 |
| GO:0042593 | glucose homeostasis | 26/1052 | 334/28832 | 0.077844311 | 2.133466906 | 4.054585746 | 0.00026204 | 0.002297472 | 0.001402146 |
| GO:0051014 | actin filament severing | 5/1052 | 17/28832 | 0.294117647 | 8.060836502 | 5.666865516 | 0.000274718 | 0.002374381 | 0.001449083 |
| GO:0009266 | response to temperature stimulus | 17/1052 | 177/28832 | 0.096045198 | 2.632295762 | 4.238936219 | 0.000277423 | 0.002393887 | 0.001460988 |
| GO:0010526 | transposable element silencing | 7/1052 | 36/28832 | 0.194444444 | 5.329108576 | 5.057725424 | 0.000278725 | 0.002393887 | 0.001460988 |
| GO:0031048 | regulatory ncRNA-mediated heterochromatin formation | 6/1052 | 26/28832 | 0.230769231 | 6.324656332 | 5.285771446 | 0.000285984 | 0.002437097 | 0.001487359 |
| GO:1901879 | regulation of protein depolymerization | 12/1052 | 100/28832 | 0.12 | 3.288821293 | 4.461701983 | 0.000286771 | 0.002440005 | 0.001489133 |
| GO:0048844 | artery morphogenesis | 11/1052 | 86/28832 | 0.127906977 | 3.505526572 | 4.528257657 | 0.000290548 | 0.002464474 | 0.001504067 |
| GO:0007498 | mesoderm development | 14/1052 | 130/28832 | 0.107692308 | 2.951506288 | 4.339674472 | 0.000292203 | 0.002474681 | 0.001510296 |
| GO:0046651 | lymphocyte proliferation | 26/1052 | 337/28832 | 0.077151335 | 2.114474619 | 4.004722286 | 0.000300922 | 0.002536741 | 0.001548171 |
| GO:0060326 | cell chemotaxis | 26/1052 | 337/28832 | 0.077151335 | 2.114474619 | 4.004722286 | 0.000300922 | 0.002536741 | 0.001548171 |
| GO:0030308 | negative regulation of cell growth | 18/1052 | 195/28832 | 0.092307692 | 2.529862533 | 4.171358891 | 0.000303656 | 0.002555852 | 0.001559835 |
| GO:0030865 | cortical cytoskeleton organization | 9/1052 | 60/28832 | 0.15 | 4.111026616 | 4.694246902 | 0.00030736 | 0.002562995 | 0.001564194 |
| GO:1902930 | regulation of alcohol biosynthetic process | 9/1052 | 60/28832 | 0.15 | 4.111026616 | 4.694246902 | 0.00030736 | 0.002562995 | 0.001564194 |
| GO:0097006 | regulation of plasma lipoprotein particle levels | 10/1052 | 73/28832 | 0.136986301 | 3.754362206 | 4.585294058 | 0.000309422 | 0.002568813 | 0.001567744 |
| GO:0140895 | cell surface toll-like receptor signaling pathway | 10/1052 | 73/28832 | 0.136986301 | 3.754362206 | 4.585294058 | 0.000309422 | 0.002568813 | 0.001567744 |
| GO:0034612 | response to tumor necrosis factor | 20/1052 | 230/28832 | 0.086956522 | 2.383203835 | 4.09848645 | 0.000318813 | 0.002607217 | 0.001591183 |
| GO:0002768 | immune response-regulating cell surface receptor signaling pathway | 25/1052 | 320/28832 | 0.078125 | 2.141159696 | 3.994654058 | 0.000323012 | 0.002637606 | 0.001609729 |
| GO:0006874 | intracellular calcium ion homeostasis | 26/1052 | 339/28832 | 0.076696165 | 2.101999843 | 3.971768206 | 0.000329594 | 0.002675388 | 0.001632787 |
| GO:0046466 | membrane lipid catabolic process | 7/1052 | 37/28832 | 0.189189189 | 5.185078615 | 4.956984299 | 0.000333025 | 0.002687248 | 0.001640026 |
| GO:0019693 | ribose phosphate metabolic process | 29/1052 | 396/28832 | 0.073232323 | 2.007066866 | 3.926840395 | 0.000333512 | 0.002687248 | 0.001640026 |
| GO:0070588 | calcium ion transmembrane transport | 27/1052 | 359/28832 | 0.075208914 | 2.061238972 | 3.93745318 | 0.000348238 | 0.002785395 | 0.001699925 |
| GO:0042987 | amyloid precursor protein catabolic process | 9/1052 | 61/28832 | 0.147540984 | 4.043632737 | 4.630749494 | 0.000349028 | 0.002787639 | 0.001701294 |
| GO:0032878 | regulation of establishment or maintenance of cell polarity | 6/1052 | 27/28832 | 0.222222222 | 6.090409801 | 5.149585711 | 0.000356412 | 0.002821897 | 0.001722201 |
| GO:0016311 | dephosphorylation | 17/1052 | 181/28832 | 0.093922652 | 2.57412348 | 4.134088736 | 0.000360724 | 0.002851913 | 0.00174052 |
| GO:0097191 | extrinsic apoptotic signaling pathway | 21/1052 | 250/28832 | 0.084 | 2.302174905 | 4.024061604 | 0.000364767 | 0.002877703 | 0.00175626 |
| GO:1903115 | regulation of actin filament-based movement | 8/1052 | 49/28832 | 0.163265306 | 4.474586793 | 4.737015197 | 0.000365039 | 0.002877703 | 0.00175626 |
| GO:0031269 | pseudopodium assembly | 5/1052 | 18/28832 | 0.277777778 | 7.613012252 | 5.46141786 | 0.00036899 | 0.00289633 | 0.001767628 |
| GO:0042180 | ketone metabolic process | 21/1052 | 251/28832 | 0.083665339 | 2.293002893 | 4.003771186 | 0.000384707 | 0.003015376 | 0.001840282 |
| GO:0034248 | regulation of amide metabolic process | 10/1052 | 75/28832 | 0.133333333 | 3.654245881 | 4.478902384 | 0.000386621 | 0.003021722 | 0.001844155 |
| GO:0032197 | retrotransposition | 7/1052 | 38/28832 | 0.184210526 | 5.048629178 | 4.859822429 | 0.000395483 | 0.003064712 | 0.001870392 |
| GO:0002832 | negative regulation of response to biotic stimulus | 15/1052 | 150/28832 | 0.1 | 2.740684411 | 4.159417895 | 0.000404249 | 0.003125902 | 0.001907735 |
| GO:0051259 | protein complex oligomerization | 23/1052 | 288/28832 | 0.079861111 | 2.188741022 | 3.945460319 | 0.000404522 | 0.003125902 | 0.001907735 |
| GO:0006509 | membrane protein ectodomain proteolysis | 8/1052 | 50/28832 | 0.16 | 4.385095057 | 4.661943316 | 0.000420827 | 0.003238181 | 0.001976259 |
| GO:1905207 | regulation of cardiocyte differentiation | 6/1052 | 28/28832 | 0.214285714 | 5.872895166 | 5.020087477 | 0.000439708 | 0.003364537 | 0.002053375 |
| GO:0045786 | negative regulation of cell cycle | 29/1052 | 404/28832 | 0.071782178 | 1.967322968 | 3.810312813 | 0.000461718 | 0.003513296 | 0.002144162 |
| GO:0035725 | sodium ion transmembrane transport | 17/1052 | 185/28832 | 0.091891892 | 2.518466756 | 4.03202467 | 0.000464827 | 0.003523773 | 0.002150556 |
| GO:0008354 | germ cell migration | 4/1052 | 11/28832 | 0.363636364 | 9.96612513 | 5.787862666 | 0.000473479 | 0.003553364 | 0.002168616 |
| GO:2000047 | regulation of cell-cell adhesion mediated by cadherin | 5/1052 | 19/28832 | 0.263157895 | 7.212327396 | 5.271187912 | 0.000485792 | 0.00361742 | 0.002207709 |
| GO:0043279 | response to alkaloid | 13/1052 | 121/28832 | 0.107438017 | 2.94453697 | 4.171155037 | 0.000485981 | 0.00361742 | 0.002207709 |
| GO:0034142 | toll-like receptor 4 signaling pathway | 9/1052 | 64/28832 | 0.140625 | 3.854087452 | 4.448094758 | 0.000502847 | 0.003722701 | 0.002271962 |
| GO:0002697 | regulation of immune effector process | 33/1052 | 485/28832 | 0.068041237 | 1.864795578 | 3.737682267 | 0.000504372 | 0.003728945 | 0.002275773 |
| GO:0098869 | cellular oxidant detoxification | 10/1052 | 78/28832 | 0.128205128 | 3.513697962 | 4.325963617 | 0.000531985 | 0.003901461 | 0.002381059 |

| | | | | | | | | | |
|---|---|---|---|---|---|---|---|---|---|
| GO:0062197 | cellular response to chemical stress | 26/1052 | 350/28832 | 0.074285714 | 2.035936991 | 3.794492723 | 0.000534588 | 0.003913597 | 0.002388465 |
| GO:0072567 | chemokine (C-X-C motif) ligand 2 production | 6/1052 | 29/28832 | 0.206896552 | 5.670381539 | 4.896706794 | 0.00053743 | 0.003913597 | 0.002388465 |
| GO:2000341 | regulation of chemokine (C-X-C motif) ligand 2 production | 6/1052 | 29/28832 | 0.206896552 | 5.670381539 | 4.896706794 | 0.00053743 | 0.003913597 | 0.002388465 |
| GO:0015695 | organic cation transport | 14/1052 | 138/28832 | 0.101449275 | 2.780404475 | 4.079756806 | 0.000538647 | 0.003913597 | 0.002388465 |
| GO:1990778 | protein localization to cell periphery | 29/1052 | 408/28832 | 0.071078431 | 1.948035488 | 3.753044218 | 0.000540796 | 0.003924 | 0.002394814 |
| GO:0086004 | regulation of cardiac muscle cell contraction | 7/1052 | 40/28832 | 0.175 | 4.796197719 | 4.675354058 | 0.000548366 | 0.003968401 | 0.002421913 |
| GO:0045995 | regulation of embryonic development | 11/1052 | 93/28832 | 0.11827957 | 3.241669733 | 4.213557049 | 0.000574742 | 0.004121122 | 0.002515118 |
| GO:0030239 | myofibril assembly | 10/1052 | 79/28832 | 0.126582278 | 3.469220773 | 4.276647819 | 0.00058948 | 0.004196954 | 0.002561398 |
| GO:0006816 | calcium ion transport | 32/1052 | 470/28832 | 0.068085106 | 1.865997897 | 3.683570125 | 0.000602113 | 0.004274758 | 0.002608882 |
| GO:0051235 | maintenance of location | 20/1052 | 242/28832 | 0.082644628 | 2.265028439 | 3.845674268 | 0.000611739 | 0.004329653 | 0.002642384 |
| GO:0106027 | neuron projection organization | 13/1052 | 124/28832 | 0.10483871 | 2.873298172 | 4.068065019 | 0.000614827 | 0.004345889 | 0.002652293 |
| GO:0031268 | pseudopodium organization | 5/1052 | 20/28832 | 0.25 | 6.851711027 | 5.094279492 | 0.000628365 | 0.004413071 | 0.002693294 |
| GO:1900424 | regulation of defense response to bacterium | 5/1052 | 20/28832 | 0.25 | 6.851711027 | 5.094279492 | 0.000628365 | 0.004413071 | 0.002693294 |
| GO:0046395 | carboxylic acid catabolic process | 20/1052 | 243/28832 | 0.082304527 | 2.255707334 | 3.825284031 | 0.000644294 | 0.004478952 | 0.002733501 |
| GO:0071695 | anatomical structure maturation | 25/1052 | 336/28832 | 0.074404762 | 2.03919971 | 3.728621564 | 0.000661788 | 0.004583089 | 0.002797056 |
| GO:0003014 | renal system process | 14/1052 | 141/28832 | 0.09929078 | 2.721246933 | 3.987048285 | 0.000668334 | 0.004616734 | 0.002817589 |
| GO:0120254 | olefinic compound metabolic process | 17/1052 | 192/28832 | 0.088541667 | 2.426647655 | 3.859690107 | 0.000709739 | 0.0048598 | 0.002965932 |
| GO:0051494 | negative regulation of cytoskeleton organization | 16/1052 | 175/28832 | 0.091428571 | 2.505768604 | 3.888064871 | 0.000712164 | 0.004864231 | 0.002968637 |
| GO:1901568 | fatty acid derivative metabolic process | 8/1052 | 54/28832 | 0.148148148 | 4.060273201 | 4.380244744 | 0.000717306 | 0.004868959 | 0.002971522 |
| GO:0043113 | receptor clustering | 10/1052 | 81/28832 | 0.12345679 | 3.383561001 | 4.180362135 | 0.000719924 | 0.004880673 | 0.002978671 |
| GO:0006887 | exocytosis | 28/1052 | 397/28832 | 0.070528967 | 1.932976411 | 3.642595129 | 0.000752959 | 0.005081138 | 0.003101015 |
| GO:0071805 | potassium ion transmembrane transport | 18/1052 | 211/28832 | 0.085308057 | 2.338024616 | 3.796072789 | 0.00077743 | 0.005212402 | 0.003181125 |
| GO:0046879 | hormone secretion | 29/1052 | 418/28832 | 0.06937799 | 1.901431768 | 3.612654376 | 0.000792735 | 0.005268841 | 0.00321557 |
| GO:0098739 | import across plasma membrane | 18/1052 | 212/28832 | 0.08490566 | 2.326996198 | 3.773761039 | 0.000821279 | 0.005399061 | 0.003295043 |
| GO:0006813 | potassium ion transport | 20/1052 | 248/28832 | 0.080645161 | 2.210229363 | 3.724805732 | 0.000830504 | 0.005435034 | 0.003316997 |
| GO:0090132 | epithelium migration | 12/1052 | 113/28832 | 0.10619469 | 2.910461321 | 3.959719448 | 0.000876695 | 0.005696435 | 0.00347653 |
| GO:0044091 | membrane biogenesis | 9/1052 | 69/28832 | 0.130434783 | 3.574805753 | 4.167000262 | 0.000879929 | 0.005697149 | 0.003476966 |
| GO:0150116 | regulation of cell-substrate junction organization | 9/1052 | 69/28832 | 0.130434783 | 3.574805753 | 4.167000262 | 0.000879929 | 0.005697149 | 0.003476966 |
| GO:0048678 | response to axon injury | 11/1052 | 98/28832 | 0.112244898 | 3.07627842 | 4.006564942 | 0.000896017 | 0.005787615 | 0.003532177 |
| GO:0060560 | developmental growth involved in morphogenesis | 22/1052 | 287/28832 | 0.076655052 | 2.100873067 | 3.647411591 | 0.00092096 | 0.005927734 | 0.003617692 |
| GO:0051446 | positive regulation of meiotic cell cycle | 6/1052 | 32/28832 | 0.1875 | 5.13878327 | 4.558511266 | 0.000934013 | 0.005983589 | 0.00365178 |
| GO:0060071 | Wnt signaling pathway, planar cell polarity pathway | 6/1052 | 32/28832 | 0.1875 | 5.13878327 | 4.558511266 | 0.000934013 | 0.005983589 | 0.00365178 |
| GO:0034764 | positive regulation of transmembrane transport | 20/1052 | 251/28832 | 0.079681275 | 2.183812279 | 3.665663486 | 0.00096311 | 0.006148422 | 0.003752378 |
| GO:0007584 | response to nutrient | 16/1052 | 180/28832 | 0.088888889 | 2.436163921 | 3.76126901 | 0.000963878 | 0.006148422 | 0.003752378 |
| GO:0033689 | negative regulation of osteoblast proliferation | 4/1052 | 13/28832 | 0.307692308 | 8.43287511 | 5.216277179 | 0.000967609 | 0.006148422 | 0.003752378 |
| GO:0050910 | detection of mechanical stimulus involved in sensory perception of sound | 5/1052 | 22/28832 | 0.227272727 | 6.228828206 | 4.774365759 | 0.001004545 | 0.006353599 | 0.003877597 |
| GO:0030278 | regulation of ossification | 14/1052 | 148/28832 | 0.094594595 | 2.592539307 | 3.779834316 | 0.00107684 | 0.00676394 | 0.004128028 |
| GO:0071514 | genomic imprinting | 6/1052 | 33/28832 | 0.181818182 | 4.983062565 | 4.455095279 | 0.001106704 | 0.006903964 | 0.004213484 |
| GO:0006734 | NADH metabolic process | 7/1052 | 45/28832 | 0.155555556 | 4.263286861 | 4.263192151 | 0.001139612 | 0.007085023 | 0.004323984 |
| GO:0045777 | positive regulation of blood pressure | 8/1052 | 58/28832 | 0.137931034 | 3.78025436 | 4.124487672 | 0.001162565 | 0.007195017 | 0.004391113 |
| GO:1900024 | regulation of substrate adhesion-dependent cell spreading | 8/1052 | 59/28832 | 0.13559322 | 3.716182252 | 4.064095852 | 0.001302593 | 0.007946307 | 0.004849626 |
| GO:0001765 | membrane raft assembly | 4/1052 | 14/28832 | 0.285714286 | 7.830526888 | 4.974597037 | 0.0013157 | 0.007962605 | 0.004859572 |
| GO:0007343 | egg activation | 4/1052 | 14/28832 | 0.285714286 | 7.830526888 | 4.974597037 | 0.0013157 | 0.007962605 | 0.004859572 |
| GO:0030166 | proteoglycan biosynthetic process | 9/1052 | 73/28832 | 0.123287671 | 3.378925986 | 3.960290757 | 0.001324411 | 0.007980016 | 0.004870198 |
| GO:0042445 | hormone metabolic process | 21/1052 | 277/28832 | 0.075812274 | 2.077775185 | 3.507507308 | 0.001364959 | 0.008215279 | 0.005013779 |
| GO:0003018 | vascular process in circulatory system | 20/1052 | 259/28832 | 0.077220077 | 2.116358618 | 3.511941663 | 0.001408821 | 0.008414473 | 0.005135347 |
| GO:0005978 | glycogen biosynthetic process | 7/1052 | 47/28832 | 0.14893617 | 4.081870399 | 4.114828942 | 0.001482828 | 0.008789328 | 0.005364121 |
| GO:0150115 | cell-substrate junction organization | 11/1052 | 105/28832 | 0.104761905 | 2.871193192 | 3.738003841 | 0.001583094 | 0.009283073 | 0.005665453 |
| GO:0071356 | cellular response to tumor necrosis factor | 17/1052 | 207/28832 | 0.082125604 | 2.250803622 | 3.514577769 | 0.00162302 | 0.009496837 | 0.005795913 |
| GO:0045576 | mast cell activation | 10/1052 | 90/28832 | 0.111111111 | 3.045204901 | 3.781561686 | 0.00163461 | 0.009544237 | 0.005824841 |
| GO:0006091 | generation of precursor metabolites and energy | 32/1052 | 500/28832 | 0.064 | 1.754038023 | 3.309872266 | 0.001653552 | 0.009644544 | 0.005886059 |
| GO:0045787 | positive regulation of cell cycle | 25/1052 | 359/28832 | 0.069637883 | 1.908554604 | 3.370957256 | 0.001672316 | 0.00974019 | 0.005944431 |
| GO:0010927 | cellular component assembly involved in morphogenesis | 14/1052 | 155/28832 | 0.090322581 | 2.475456887 | 3.584240288 | 0.001677072 | 0.00974019 | 0.005944431 |
| GO:0044331 | cell-cell adhesion mediated by cadherin | 7/1052 | 48/28832 | 0.145833333 | 3.996831432 | 4.043700326 | 0.001682141 | 0.009759273 | 0.005956077 |
| GO:0031100 | animal organ regeneration | 9/1052 | 76/28832 | 0.118421053 | 3.245547328 | 3.814489062 | 0.001764083 | 0.010148581 | 0.006193672 |
| GO:2000108 | positive regulation of leukocyte apoptotic process | 6/1052 | 36/28832 | 0.166666667 | 4.567807351 | 4.168292301 | 0.001773583 | 0.010171147 | 0.006207444 |
| GO:0015867 | ATP transport | 5/1052 | 25/28832 | 0.2 | 5.481368821 | 4.36217695 | 0.001851042 | 0.01056001 | 0.006444767 |
| GO:0002367 | cytokine production involved in immune response | 14/1052 | 157/28832 | 0.089171975 | 2.443922404 | 3.530315913 | 0.001892393 | 0.010762249 | 0.006568193 |
| GO:0071825 | protein-lipid complex organization | 7/1052 | 49/28832 | 0.142857143 | 3.915263444 | 3.974471803 | 0.001901658 | 0.010781318 | 0.006579831 |
| GO:0046034 | ATP metabolic process | 19/1052 | 247/28832 | 0.076923077 | 2.108218777 | 3.403897588 | 0.001912275 | 0.010830288 | 0.006609717 |
| GO:0006694 | steroid biosynthetic process | 15/1052 | 175/28832 | 0.085714286 | 2.349158066 | 3.483678747 | 0.001963305 | 0.011073443 | 0.006758114 |
| GO:0021675 | nerve development | 11/1052 | 108/28832 | 0.101851852 | 2.791437826 | 3.629633386 | 0.001986676 | 0.011193723 | 0.006831521 |
| GO:0001101 | response to acid chemical | 20/1052 | 267/28832 | 0.074906367 | 2.052947124 | 3.363695556 | 0.002019448 | 0.011340577 | 0.006921146 |
| GO:0033687 | osteoblast proliferation | 6/1052 | 37/28832 | 0.162162162 | 4.444353098 | 4.079637636 | 0.002052401 | 0.011481274 | 0.007007014 |
| GO:0042113 | B cell activation | 23/1052 | 326/28832 | 0.070552147 | 1.933611701 | 3.29897138 | 0.002126161 | 0.011857521 | 0.007236637 |
| GO:0006885 | regulation of pH | 11/1052 | 109/28832 | 0.100917431 | 2.765828304 | 3.594333958 | 0.002138433 | 0.011913457 | 0.007270774 |
| GO:0014910 | regulation of smooth muscle cell migration | 11/1052 | 110/28832 | 0.1 | 2.740684411 | 3.559431551 | 0.002299463 | 0.012580362 | 0.007677786 |
| GO:0043467 | regulation of generation of precursor metabolites and energy | 14/1052 | 161/28832 | 0.086956522 | 2.383203835 | 3.424911111 | 0.002392082 | 0.013047938 | 0.007963147 |
| GO:0002090 | regulation of receptor internalization | 10/1052 | 95/28832 | 0.105263158 | 2.884930959 | 3.581031476 | 0.002453865 | 0.013331775 | 0.008136373 |
| GO:0043523 | regulation of neuron apoptotic process | 23/1052 | 330/28832 | 0.06969697 | 1.910173983 | 3.236050733 | 0.002480116 | 0.013461028 | 0.008215256 |
| GO:0071383 | cellular response to steroid hormone stimulus | 17/1052 | 216/28832 | 0.078703704 | 2.157020138 | 3.321505166 | 0.002544162 | 0.013754064 | 0.008394095 |
| GO:0032528 | microvillus organization | 5/1052 | 27/28832 | 0.185185185 | 5.075341501 | 4.122717262 | 0.002648053 | 0.014189516 | 0.008659851 |
| GO:0098743 | cell aggregation | 5/1052 | 27/28832 | 0.185185185 | 5.075341501 | 4.122717262 | 0.002648053 | 0.014189516 | 0.008659851 |

| | | | | | | | | | |
|---|---|---|---|---|---|---|---|---|---|
| GO:2000773 | negative regulation of cellular senescence | 5/1052 | 27/28832 | 0.185185185 | 5.075341501 | 4.122717262 | 0.002648053 | 0.014189516 | 0.008659851 |
| GO:0071887 | leukocyte apoptotic process | 14/1052 | 163/28832 | 0.085889571 | 2.35396207 | 3.373382833 | 0.00268001 | 0.014319181 | 0.008738986 |
| GO:0002237 | response to molecule of bacterial origin | 31/1052 | 495/28832 | 0.062626263 | 1.716388217 | 3.128568732 | 0.002689369 | 0.014319181 | 0.008738986 |
| GO:0033628 | regulation of cell adhesion mediated by integrin | 7/1052 | 52/28832 | 0.134615385 | 3.68938286 | 3.777291593 | 0.002695044 | 0.014319181 | 0.008738986 |
| GO:0051180 | vitamin transport | 7/1052 | 52/28832 | 0.134615385 | 3.68938286 | 3.777291593 | 0.002695044 | 0.014319181 | 0.008738986 |
| GO:0043616 | keratinocyte proliferation | 8/1052 | 66/28832 | 0.121212121 | 3.32204171 | 3.675139669 | 0.002698424 | 0.014319181 | 0.008738986 |
| GO:0141193 | nuclear receptor-mediated signaling pathway | 15/1052 | 181/28832 | 0.082872928 | 2.271285423 | 3.338751258 | 0.002720613 | 0.014381136 | 0.008776797 |
| GO:0009743 | response to carbohydrate | 21/1052 | 296/28832 | 0.070945946 | 1.944404481 | 3.178183352 | 0.003033766 | 0.015702682 | 0.009583335 |
| GO:0007044 | cell-substrate junction assembly | 10/1052 | 98/28832 | 0.102040816 | 2.796616746 | 3.466905785 | 0.003085309 | 0.015894278 | 0.009700266 |
| GO:0006007 | glucose catabolic process | 5/1052 | 28/28832 | 0.178571429 | 4.894079305 | 4.011705175 | 0.003128127 | 0.016024331 | 0.009779638 |
| GO:0006883 | intracellular sodium ion homeostasis | 5/1052 | 28/28832 | 0.178571429 | 4.894079305 | 4.011705175 | 0.003128127 | 0.016024331 | 0.009779638 |
| GO:0033003 | regulation of mast cell activation | 5/1052 | 28/28832 | 0.178571429 | 4.894079305 | 4.011705175 | 0.003128127 | 0.016024331 | 0.009779638 |
| GO:0031669 | cellular response to nutrient levels | 20/1052 | 278/28832 | 0.071942446 | 1.971715403 | 3.168105312 | 0.003212824 | 0.016427439 | 0.010025654 |
| GO:0002275 | myeloid cell activation involved in immune response | 11/1052 | 115/28832 | 0.095652174 | 2.621524219 | 3.390583373 | 0.003258379 | 0.016599743 | 0.010130811 |
| GO:0031128 | developmental induction | 6/1052 | 41/28832 | 0.146341463 | 4.010757674 | 3.754144266 | 0.0035086 | 0.017724421 | 0.010817202 |
| GO:0052652 | cyclic purine nucleotide metabolic process | 6/1052 | 41/28832 | 0.146341463 | 4.010757674 | 3.754144266 | 0.0035086 | 0.017724421 | 0.010817202 |
| GO:0002718 | regulation of cytokine production involved in immune response | 12/1052 | 133/28832 | 0.090225564 | 2.472797964 | 3.312891525 | 0.003518081 | 0.017747375 | 0.01083121 |
| GO:0051302 | regulation of cell division | 15/1052 | 186/28832 | 0.080645161 | 2.210229363 | 3.222283641 | 0.00352271 | 0.017747375 | 0.01083121 |
| GO:0010907 | positive regulation of glucose metabolic process | 8/1052 | 69/28832 | 0.115942029 | 3.177605114 | 3.524180832 | 0.003570624 | 0.01783992 | 0.01088769 |
| GO:0006068 | ethanol catabolic process | 4/1052 | 18/28832 | 0.222222222 | 6.090409801 | 4.203962422 | 0.003580375 | 0.01783992 | 0.01088769 |
| GO:0034375 | high-density lipoprotein particle remodeling | 4/1052 | 18/28832 | 0.222222222 | 6.090409801 | 4.203962422 | 0.003580375 | 0.01783992 | 0.01088769 |
| GO:0070365 | hepatocyte differentiation | 4/1052 | 18/28832 | 0.222222222 | 6.090409801 | 4.203962422 | 0.003580375 | 0.01783992 | 0.01088769 |
| GO:0010959 | regulation of metal ion transport | 28/1052 | 442/28832 | 0.063348416 | 1.73618017 | 3.035177929 | 0.003589895 | 0.017871079 | 0.010906707 |
| GO:0031579 | membrane raft organization | 5/1052 | 29/28832 | 0.172413793 | 4.725317949 | 3.905845713 | 0.003667856 | 0.018143617 | 0.011073037 |
| GO:0070193 | synaptonemal complex organization | 5/1052 | 29/28832 | 0.172413793 | 4.725317949 | 3.905845713 | 0.003667856 | 0.018143617 | 0.011073037 |
| GO:0044703 | multi-organism reproductive process | 19/1052 | 262/28832 | 0.072519084 | 1.987519229 | 3.124731379 | 0.003680943 | 0.018191906 | 0.011102507 |
| GO:0044848 | biological phase | 7/1052 | 55/28832 | 0.127272727 | 3.488143795 | 3.59422834 | 0.003720543 | 0.018213373 | 0.011115608 |
| GO:0045824 | negative regulation of innate immune response | 11/1052 | 117/28832 | 0.094017094 | 2.576711839 | 3.32554211 | 0.003721907 | 0.018213373 | 0.011115608 |
| GO:0051261 | protein depolymerization | 12/1052 | 134/28832 | 0.089552239 | 2.454344248 | 3.283714597 | 0.003739149 | 0.018281399 | 0.011157125 |
| GO:0045732 | positive regulation of protein catabolic process | 16/1052 | 206/28832 | 0.077669903 | 2.128686921 | 3.163722415 | 0.003813457 | 0.018604462 | 0.01135429 |
| GO:0016485 | protein processing | 19/1052 | 263/28832 | 0.072243346 | 1.979962122 | 3.106785334 | 0.003835908 | 0.018687671 | 0.011405072 |
| GO:0060759 | regulation of response to cytokine stimulus | 14/1052 | 170/28832 | 0.082352941 | 2.257034221 | 3.198820848 | 0.003920552 | 0.019066078 | 0.011636014 |
| GO:0060325 | face morphogenesis | 6/1052 | 42/28832 | 0.142857143 | 3.915263444 | 3.679198519 | 0.003969223 | 0.019149184 | 0.011686733 |
| GO:2000677 | regulation of transcription regulatory region DNA binding | 6/1052 | 42/28832 | 0.142857143 | 3.915263444 | 3.679198519 | 0.003969223 | 0.019149184 | 0.011686733 |
| GO:1903169 | regulation of calcium ion transmembrane transport | 15/1052 | 189/28832 | 0.079365079 | 2.175146358 | 3.154170835 | 0.004090569 | 0.01968294 | 0.012012484 |
| GO:0005976 | polysaccharide metabolic process | 11/1052 | 119/28832 | 0.092436975 | 2.533405758 | 3.26184175 | 0.004236661 | 0.020261167 | 0.012365376 |
| GO:0042542 | response to hydrogen peroxide | 11/1052 | 119/28832 | 0.092436975 | 2.533405758 | 3.26184175 | 0.004236661 | 0.020261167 | 0.012365376 |
| GO:0051090 | regulation of DNA-binding transcription factor activity | 24/1052 | 365/28832 | 0.065753425 | 1.802093859 | 3.001044085 | 0.004246335 | 0.020266818 | 0.012368825 |
| GO:0042594 | response to starvation | 18/1052 | 247/28832 | 0.072874494 | 1.997259894 | 3.063087017 | 0.004393529 | 0.02068929 | 0.012626659 |
| GO:0007040 | lysosome organization | 10/1052 | 103/28832 | 0.097087379 | 2.660858651 | 3.285962728 | 0.004419094 | 0.020699368 | 0.012632809 |
| GO:0070228 | regulation of lymphocyte apoptotic process | 9/1052 | 87/28832 | 0.103448276 | 2.83519077 | 3.3360403 | 0.00445736 | 0.020827499 | 0.012711008 |
| GO:0044706 | multi-multicellular organism process | 19/1052 | 267/28832 | 0.071161049 | 1.950299768 | 3.035783121 | 0.00451088 | 0.020984342 | 0.012806729 |
| GO:0140719 | constitutive heterochromatin formation | 7/1052 | 57/28832 | 0.122807018 | 3.365752785 | 3.479130538 | 0.004552878 | 0.021109149 | 0.012882898 |
| GO:0007586 | digestion | 12/1052 | 138/28832 | 0.086956522 | 2.383203835 | 3.169580475 | 0.004737779 | 0.021855303 | 0.013338276 |
| GO:0043270 | positive regulation of monoatomic ion transport | 18/1052 | 249/28832 | 0.072289157 | 1.981217646 | 3.026096137 | 0.004774356 | 0.021878466 | 0.013352412 |
| GO:0021603 | cranial nerve formation | 3/1052 | 10/28832 | 0.3 | 8.222053232 | 4.444984291 | 0.004794787 | 0.021878466 | 0.013352412 |
| GO:0035338 | long-chain fatty-acyl-CoA biosynthetic process | 3/1052 | 10/28832 | 0.3 | 8.222053232 | 4.444984291 | 0.004794787 | 0.021878466 | 0.013352412 |
| GO:0046129 | purine ribonucleoside biosynthetic process | 3/1052 | 10/28832 | 0.3 | 8.222053232 | 4.444984291 | 0.004794787 | 0.021878466 | 0.013352412 |
| GO:0140928 | inhibition of non-skeletal tissue mineralization | 3/1052 | 10/28832 | 0.3 | 8.222053232 | 4.444984291 | 0.004794787 | 0.021878466 | 0.013352412 |
| GO:0042886 | amide transport | 26/1052 | 410/28832 | 0.063414634 | 1.737994992 | 2.928799406 | 0.004824521 | 0.021940945 | 0.013390543 |
| GO:0060561 | apoptotic process involved in morphogenesis | 5/1052 | 31/28832 | 0.161290323 | 4.420458727 | 3.707942749 | 0.004942023 | 0.022308423 | 0.013614814 |
| GO:0045022 | early endosome to late endosome transport | 6/1052 | 44/28832 | 0.136363636 | 3.737296924 | 3.536015034 | 0.005022553 | 0.022523267 | 0.013745934 |
| GO:0008333 | endosome to lysosome transport | 8/1052 | 73/28832 | 0.109589041 | 3.003489765 | 3.335287456 | 0.005055372 | 0.022614835 | 0.013801817 |
| GO:0170040 | proteinogenic amino acid catabolic process | 8/1052 | 73/28832 | 0.109589041 | 3.003489765 | 3.335287456 | 0.005055372 | 0.022614835 | 0.013801817 |
| GO:0009612 | response to mechanical stimulus | 19/1052 | 270/28832 | 0.07037037 | 1.92862977 | 2.983333324 | 0.005079 | 0.022701968 | 0.013854995 |
| GO:0016050 | vesicle organization | 25/1052 | 391/28832 | 0.063938619 | 1.752355761 | 2.914813691 | 0.005088996 | 0.02270954 | 0.013859616 |
| GO:0051607 | defense response to virus | 22/1052 | 330/28832 | 0.066666667 | 1.82712294 | 2.940769399 | 0.005133864 | 0.022872451 | 0.01395904 |
| GO:0034587 | piRNA processing | 4/1052 | 20/28832 | 0.2 | 5.481368821 | 3.901311117 | 0.005349905 | 0.023604301 | 0.014405688 |
| GO:0045725 | positive regulation of glycogen biosynthetic process | 4/1052 | 20/28832 | 0.2 | 5.481368821 | 3.901311117 | 0.005349905 | 0.023604301 | 0.014405688 |
| GO:0022600 | digestive system process | 11/1052 | 123/28832 | 0.089430894 | 2.451018579 | 3.138254053 | 0.005435439 | 0.023943063 | 0.014612434 |
| GO:0009615 | response to virus | 26/1052 | 414/28832 | 0.062801932 | 1.72120277 | 2.876288229 | 0.005460561 | 0.024034374 | 0.014668161 |
| GO:0010951 | negative regulation of endopeptidase activity | 7/1052 | 59/28832 | 0.118644068 | 3.25165947 | 3.369052245 | 0.005518395 | 0.024230448 | 0.014787825 |
| GO:0046677 | response to antibiotic | 6/1052 | 45/28832 | 0.133333333 | 3.654245881 | 3.467534632 | 0.005619928 | 0.024597301 | 0.015011715 |
| GO:0042178 | xenobiotic catabolic process | 5/1052 | 32/28832 | 0.15625 | 4.282319392 | 3.615190492 | 0.005684247 | 0.024642248 | 0.015039146 |
| GO:0021915 | neural tube development | 15/1052 | 196/28832 | 0.076530612 | 2.097462559 | 3.000082336 | 0.005709081 | 0.024730312 | 0.015092892 |
| GO:0050886 | endocrine process | 11/1052 | 124/28832 | 0.088709677 | 2.4312523 | 3.108115675 | 0.005773332 | 0.024949369 | 0.015226582 |
| GO:0170035 | L-amino acid catabolic process | 8/1052 | 75/28832 | 0.106666667 | 2.923396705 | 3.245632271 | 0.005955073 | 0.025613368 | 0.01563182 |
| GO:0002532 | production of molecular mediator involved in inflammatory response | 10/1052 | 108/28832 | 0.092592593 | 2.537670751 | 3.115475744 | 0.006170035 | 0.026434035 | 0.016132672 |
| GO:0071456 | cellular response to hypoxia | 13/1052 | 161/28832 | 0.080745342 | 2.21297499 | 3.003412378 | 0.00627185 | 0.02673178 | 0.016314386 |
| GO:0006534 | cysteine metabolic process | 3/1052 | 11/28832 | 0.272727273 | 7.474593847 | 4.179515617 | 0.006415278 | 0.026752677 | 0.016327139 |
| GO:0010248 | establishment or maintenance of transmembrane electrochemical gradient | 3/1052 | 11/28832 | 0.272727273 | 7.474593847 | 4.179515617 | 0.006415278 | 0.026752677 | 0.016327139 |
| GO:0021707 | cerebellar granule cell differentiation | 3/1052 | 11/28832 | 0.272727273 | 7.474593847 | 4.179515617 | 0.006415278 | 0.026752677 | 0.016327139 |
| GO:0031642 | negative regulation of myelination | 3/1052 | 11/28832 | 0.272727273 | 7.474593847 | 4.179515617 | 0.006415278 | 0.026752677 | 0.016327139 |

| | | | | | | | | | |
|---|---|---|---|---|---|---|---|---|---|
| GO:0046940 | nucleoside monophosphate phosphorylation | 3/1052 | 11/28832 | 0.272727273 | 7.474593847 | 4.179515617 | 0.006415278 | 0.026752677 | 0.016327139 |
| GO:0060717 | chorion development | 3/1052 | 11/28832 | 0.272727273 | 7.474593847 | 4.179515617 | 0.006415278 | 0.026752677 | 0.016327139 |
| GO:1902966 | positive regulation of protein localization to early endosome | 3/1052 | 11/28832 | 0.272727273 | 7.474593847 | 4.179515617 | 0.006415278 | 0.026752677 | 0.016327139 |
| GO:0006071 | glycerol metabolic process | 4/1052 | 21/28832 | 0.19047619 | 5.220351258 | 3.764876045 | 0.006420447 | 0.026752677 | 0.016327139 |
| GO:0015693 | magnesium ion transport | 4/1052 | 21/28832 | 0.19047619 | 5.220351258 | 3.764876045 | 0.006420447 | 0.026752677 | 0.016327139 |
| GO:0035457 | cellular response to interferon-alpha | 4/1052 | 21/28832 | 0.19047619 | 5.220351258 | 3.764876045 | 0.006420447 | 0.026752677 | 0.016327139 |
| GO:0043117 | positive regulation of vascular permeability | 4/1052 | 21/28832 | 0.19047619 | 5.220351258 | 3.764876045 | 0.006420447 | 0.026752677 | 0.016327139 |
| GO:0045836 | positive regulation of meiotic nuclear division | 4/1052 | 21/28832 | 0.19047619 | 5.220351258 | 3.764876045 | 0.006420447 | 0.026752677 | 0.016327139 |
| GO:0034405 | response to fluid shear stress | 5/1052 | 33/28832 | 0.151515152 | 4.152552137 | 3.526161067 | 0.006501666 | 0.026967864 | 0.016458468 |
| GO:0032963 | collagen metabolic process | 11/1052 | 127/28832 | 0.086614173 | 2.373821143 | 3.019429425 | 0.006887 | 0.028431664 | 0.017351824 |
| GO:0009311 | oligosaccharide metabolic process | 6/1052 | 47/28832 | 0.127659574 | 3.498746056 | 3.336257292 | 0.00696776 | 0.028612926 | 0.017462448 |
| GO:0043045 | epigenetic programming of gene expression | 6/1052 | 47/28832 | 0.127659574 | 3.498746056 | 3.336257292 | 0.00696776 | 0.028612926 | 0.017462448 |
| GO:0050892 | intestinal absorption | 6/1052 | 47/28832 | 0.127659574 | 3.498746056 | 3.336257292 | 0.00696776 | 0.028612926 | 0.017462448 |
| GO:0090559 | regulation of membrane permeability | 8/1052 | 77/28832 | 0.103896104 | 2.847464323 | 3.158903355 | 0.006971492 | 0.028612926 | 0.017462448 |
| GO:2000352 | negative regulation of endothelial cell apoptotic process | 5/1052 | 34/28832 | 0.147058824 | 4.030418251 | 3.440586413 | 0.007398015 | 0.030092595 | 0.018365489 |
| GO:0051048 | negative regulation of secretion | 16/1052 | 221/28832 | 0.07239819 | 1.984205908 | 2.858164286 | 0.007417013 | 0.030147459 | 0.018398973 |
| GO:0035461 | vitamin transmembrane transport | 4/1052 | 22/28832 | 0.181818182 | 4.983062565 | 3.636875563 | 0.007623981 | 0.030782877 | 0.018786768 |
| GO:0090050 | positive regulation of cell migration involved in sprouting angiogenesis | 4/1052 | 22/28832 | 0.181818182 | 4.983062565 | 3.636875563 | 0.007623981 | 0.030782877 | 0.018786768 |
| GO:0010712 | regulation of collagen metabolic process | 7/1052 | 63/28832 | 0.111111111 | 3.045204901 | 3.162396485 | 0.007898785 | 0.031696285 | 0.01934422 |
| GO:0050708 | regulation of protein secretion | 22/1052 | 343/28832 | 0.064139942 | 1.757873383 | 2.747746822 | 0.007949328 | 0.0317915 | 0.01940233 |
| GO:0031623 | receptor internalization | 13/1052 | 166/28832 | 0.078313253 | 2.146319117 | 2.88235609 | 0.008040061 | 0.032130878 | 0.019609452 |
| GO:0032414 | positive regulation of ion transmembrane transporter activity | 8/1052 | 79/28832 | 0.101265823 | 2.775376618 | 3.074921702 | 0.008113652 | 0.032354073 | 0.019745667 |
| GO:0002444 | myeloid leukocyte mediated immunity | 11/1052 | 130/28832 | 0.084615385 | 2.319040655 | 2.933225013 | 0.008161867 | 0.032498962 | 0.019834093 |
| GO:0044782 | cilium organization | 27/1052 | 449/28832 | 0.06013363 | 1.648073031 | 2.693330028 | 0.008195351 | 0.032584858 | 0.019886515 |
| GO:0002176 | male germ cell proliferation | 3/1052 | 12/28832 | 0.25 | 6.851711027 | 3.945464212 | 0.008323789 | 0.032644768 | 0.019923079 |
| GO:0060700 | regulation of ribonuclease activity | 3/1052 | 12/28832 | 0.25 | 6.851711027 | 3.945464212 | 0.008323789 | 0.032644768 | 0.019923079 |
| GO:0070244 | negative regulation of thymocyte apoptotic process | 3/1052 | 12/28832 | 0.25 | 6.851711027 | 3.945464212 | 0.008323789 | 0.032644768 | 0.019923079 |
| GO:0035116 | embryonic hindlimb morphogenesis | 5/1052 | 35/28832 | 0.142857143 | 3.915263444 | 3.358225137 | 0.008376944 | 0.032712534 | 0.019964436 |
| GO:0098815 | modulation of excitatory postsynaptic potential | 7/1052 | 64/28832 | 0.109375 | 2.997623574 | 3.113295965 | 0.008596628 | 0.033261777 | 0.020299639 |
| GO:0016051 | carbohydrate biosynthetic process | 16/1052 | 225/28832 | 0.071111111 | 1.948931136 | 2.780746507 | 0.008737668 | 0.033735908 | 0.020589001 |
| GO:0007416 | synapse assembly | 20/1052 | 305/28832 | 0.06557377 | 1.797170105 | 2.723607207 | 0.008780443 | 0.033877155 | 0.020675203 |
| GO:0010921 | regulation of phosphatase activity | 4/1052 | 23/28832 | 0.173913043 | 4.766407671 | 3.516403909 | 0.008966853 | 0.0343541 | 0.020966283 |
| GO:0030903 | notochord development | 4/1052 | 23/28832 | 0.173913043 | 4.766407671 | 3.516403909 | 0.008966853 | 0.0343541 | 0.020966283 |
| GO:0042953 | lipoprotein transport | 4/1052 | 23/28832 | 0.173913043 | 4.766407671 | 3.516403909 | 0.008966853 | 0.0343541 | 0.020966283 |
| GO:0045104 | intermediate filament cytoskeleton organization | 9/1052 | 97/28832 | 0.092783505 | 2.542903061 | 2.962037206 | 0.009016576 | 0.034496284 | 0.021053057 |
| GO:0051648 | vesicle localization | 16/1052 | 226/28832 | 0.07079646 | 1.940307547 | 2.761640713 | 0.009095473 | 0.034725284 | 0.021192816 |
| GO:0008206 | bile acid metabolic process | 6/1052 | 50/28832 | 0.12 | 3.288821293 | 3.152158195 | 0.009406796 | 0.035664992 | 0.021766319 |
| GO:1902459 | positive regulation of stem cell population maintenance | 6/1052 | 50/28832 | 0.12 | 3.288821293 | 3.152158195 | 0.009406796 | 0.035664992 | 0.021766319 |
| GO:0033280 | response to vitamin D | 5/1052 | 36/28832 | 0.138888889 | 3.806506126 | 3.278859177 | 0.009442002 | 0.035674858 | 0.02177234 |
| GO:0045103 | intermediate filament-based process | 9/1052 | 98/28832 | 0.091836735 | 2.516955071 | 2.927246629 | 0.009616367 | 0.036283547 | 0.022143823 |
| GO:0051453 | regulation of intracellular pH | 8/1052 | 82/28832 | 0.097560976 | 2.673838449 | 2.953745981 | 0.010082515 | 0.037937718 | 0.023153362 |
| GO:0032964 | collagen biosynthetic process | 7/1052 | 66/28832 | 0.106060606 | 2.906786496 | 3.017907328 | 0.010126519 | 0.038024834 | 0.023206529 |
| GO:0060271 | cilium assembly | 25/1052 | 414/28832 | 0.060386473 | 1.655002663 | 2.612270126 | 0.010150197 | 0.038061499 | 0.023228906 |
| GO:0002456 | T cell mediated immunity | 13/1052 | 171/28832 | 0.076023392 | 2.083561248 | 2.765524133 | 0.010179261 | 0.038135105 | 0.023273827 |
| GO:0006766 | vitamin metabolic process | 9/1052 | 99/28832 | 0.090909091 | 2.491531282 | 2.8928844 | 0.010245766 | 0.03834103 | 0.023399503 |
| GO:0098884 | postsynaptic neurotransmitter receptor internalization | 6/1052 | 51/28832 | 0.117647059 | 3.224334601 | 3.093882872 | 0.01034014 | 0.03843439 | 0.02345648 |
| GO:0001768 | establishment of T cell polarity | 4/1052 | 24/28832 | 0.166666667 | 4.567807351 | 3.402687494 | 0.010454979 | 0.03843439 | 0.02345648 |
| GO:0030220 | platelet formation | 4/1052 | 24/28832 | 0.166666667 | 4.567807351 | 3.402687494 | 0.010454979 | 0.03843439 | 0.02345648 |
| GO:0044872 | lipoprotein localization | 4/1052 | 24/28832 | 0.166666667 | 4.567807351 | 3.402687494 | 0.010454979 | 0.03843439 | 0.02345648 |
| GO:2000291 | regulation of myoblast proliferation | 4/1052 | 24/28832 | 0.166666667 | 4.567807351 | 3.402687494 | 0.010454979 | 0.03843439 | 0.02345648 |
| GO:0042270 | protection from natural killer cell mediated cytotoxicity | 3/1052 | 13/28832 | 0.230769231 | 6.324656332 | 3.736761738 | 0.010530643 | 0.03843439 | 0.02345648 |
| GO:0046618 | xenobiotic export from cell | 3/1052 | 13/28832 | 0.230769231 | 6.324656332 | 3.736761738 | 0.010530643 | 0.03843439 | 0.02345648 |
| GO:0072182 | regulation of nephron tubule epithelial cell differentiation | 3/1052 | 13/28832 | 0.230769231 | 6.324656332 | 3.736761738 | 0.010530643 | 0.03843439 | 0.02345648 |
| GO:1902946 | protein localization to early endosome | 3/1052 | 13/28832 | 0.230769231 | 6.324656332 | 3.736761738 | 0.010530643 | 0.03843439 | 0.02345648 |
| GO:1904383 | response to sodium phosphate | 3/1052 | 13/28832 | 0.230769231 | 6.324656332 | 3.736761738 | 0.010530643 | 0.03843439 | 0.02345648 |
| GO:2000678 | negative regulation of transcription regulatory region DNA binding | 3/1052 | 13/28832 | 0.230769231 | 6.324656332 | 3.736761738 | 0.010530643 | 0.03843439 | 0.02345648 |
| GO:0051450 | myoblast proliferation | 5/1052 | 37/28832 | 0.135135135 | 3.703627582 | 3.202290974 | 0.010596627 | 0.038623683 | 0.023572006 |
| GO:0051402 | neuron apoptotic process | 24/1052 | 396/28832 | 0.060606061 | 1.661020855 | 2.577508487 | 0.011138708 | 0.040357532 | 0.024630173 |
| GO:1904951 | positive regulation of establishment of protein localization | 23/1052 | 375/28832 | 0.061333333 | 1.680953105 | 2.582913629 | 0.011199974 | 0.040525834 | 0.024732887 |
| GO:0016202 | regulation of striated muscle tissue development | 6/1052 | 52/28832 | 0.115384615 | 3.162328166 | 3.037032816 | 0.011337327 | 0.040968637 | 0.02500313 |
| GO:0007632 | visual behavior | 8/1052 | 84/28832 | 0.095238095 | 2.610175629 | 2.875945662 | 0.011578731 | 0.041686216 | 0.025441068 |
| GO:0032411 | positive regulation of transporter activity | 8/1052 | 84/28832 | 0.095238095 | 2.610175629 | 2.875945662 | 0.011578731 | 0.041686216 | 0.025441068 |
| GO:0051279 | regulation of release of sequestered calcium ion into cytosol | 8/1052 | 84/28832 | 0.095238095 | 2.610175629 | 2.875945662 | 0.011578731 | 0.041686216 | 0.025441068 |
| GO:0090398 | cellular senescence | 9/1052 | 101/28832 | 0.089108911 | 2.442194029 | 2.825404362 | 0.01159686 | 0.041686216 | 0.025441068 |
| GO:0036075 | replacement ossification | 5/1052 | 38/28832 | 0.131578947 | 3.606163698 | 3.128341036 | 0.011844136 | 0.042333642 | 0.025836191 |
| GO:0007127 | meiosis I | 11/1052 | 137/28832 | 0.080291971 | 2.200549527 | 2.740998843 | 0.01184661 | 0.042333642 | 0.025836191 |
| GO:0050764 | regulation of phagocytosis | 10/1052 | 119/28832 | 0.084033613 | 2.303096143 | 2.771929988 | 0.011891384 | 0.042465903 | 0.02591691 |
| GO:0044321 | response to leptin | 4/1052 | 25/28832 | 0.16 | 4.385095057 | 3.295061 | 0.012093824 | 0.042908764 | 0.026187188 |
| GO:1901570 | fatty acid derivative biosynthetic process | 4/1052 | 25/28832 | 0.16 | 4.385095057 | 3.295061 | 0.012093824 | 0.042908764 | 0.026187188 |
| GO:2000345 | regulation of hepatocyte proliferation | 4/1052 | 25/28832 | 0.16 | 4.385095057 | 3.295061 | 0.012093824 | 0.042908764 | 0.026187188 |
| GO:0051602 | response to electrical stimulus | 6/1052 | 53/28832 | 0.113207547 | 3.102661597 | 2.981542953 | 0.012400499 | 0.043883009 | 0.026781769 |
| GO:0072538 | T-helper 17 type immune response | 6/1052 | 53/28832 | 0.113207547 | 3.102661597 | 2.981542953 | 0.012400499 | 0.043883009 | 0.026781769 |
| GO:0032526 | response to retinoic acid | 10/1052 | 120/28832 | 0.083333333 | 2.283903676 | 2.742602988 | 0.012562014 | 0.044425844 | 0.027113061 |

| | | | | | | | | | |
|---|---|---|---|---|---|---|---|---|---|
| GO:0051785 | positive regulation of nuclear division | 7/1052 | 69/28832 | 0.101449275 | 2.780404475 | 2.881361401 | 0.012779868 | 0.045050683 | 0.0274944 |
| GO:0000280 | nuclear division | 27/1052 | 466/28832 | 0.057939914 | 1.587950195 | 2.490038431 | 0.012899376 | 0.045413442 | 0.027715791 |
| GO:0072172 | mesonephric tubule formation | 3/1052 | 14/28832 | 0.214285714 | 5.872895166 | 3.548875548 | 0.01304381 | 0.045425005 | 0.027722848 |
| GO:2000303 | regulation of ceramide biosynthetic process | 3/1052 | 14/28832 | 0.214285714 | 5.872895166 | 3.548875548 | 0.01304381 | 0.045425005 | 0.027722848 |
| GO:2000644 | regulation of receptor catabolic process | 3/1052 | 14/28832 | 0.214285714 | 5.872895166 | 3.548875548 | 0.01304381 | 0.045425005 | 0.027722848 |
| GO:0031342 | negative regulation of cell killing | 5/1052 | 39/28832 | 0.128205128 | 3.513697962 | 3.05684586 | 0.013187711 | 0.045693457 | 0.027886684 |
| GO:0046189 | phenol-containing compound biosynthetic process | 6/1052 | 54/28832 | 0.111111111 | 3.045204901 | 2.927352373 | 0.013531751 | 0.046561139 | 0.02841623 |
| GO:1901861 | regulation of muscle tissue development | 6/1052 | 54/28832 | 0.111111111 | 3.045204901 | 2.927352373 | 0.013531751 | 0.046561139 | 0.02841623 |
| GO:0000768 | syncytium formation by plasma membrane fusion | 7/1052 | 70/28832 | 0.1 | 2.740684411 | 2.837468972 | 0.013765942 | 0.047277757 | 0.028853581 |
| GO:0140253 | cell-cell fusion | 7/1052 | 70/28832 | 0.1 | 2.740684411 | 2.837468972 | 0.013765942 | 0.047277757 | 0.028853581 |
| GO:0070555 | response to interleukin-1 | 9/1052 | 104/28832 | 0.086538462 | 2.371746125 | 2.727151833 | 0.013867209 | 0.047341695 | 0.028892602 |
| GO:0007130 | synaptonemal complex assembly | 4/1052 | 26/28832 | 0.153846154 | 4.216437555 | 3.192948563 | 0.013888397 | 0.047341695 | 0.028892602 |
| GO:0036344 | platelet morphogenesis | 4/1052 | 26/28832 | 0.153846154 | 4.216437555 | 3.192948563 | 0.013888397 | 0.047341695 | 0.028892602 |
| GO:0051043 | regulation of membrane protein ectodomain proteolysis | 4/1052 | 26/28832 | 0.153846154 | 4.216437555 | 3.192948563 | 0.013888397 | 0.047341695 | 0.028892602 |
| GO:0001964 | startle response | 5/1052 | 40/28832 | 0.125 | 3.425855513 | 2.987656139 | 0.014630398 | 0.049685232 | 0.030322862 |
| GO:0071312 | cellular response to alkaloid | 5/1052 | 40/28832 | 0.125 | 3.425855513 | 2.987656139 | 0.014630398 | 0.049685232 | 0.030322862 |
| GO:0019233 | sensory perception of pain | 9/1052 | 105/28832 | 0.085714286 | 2.349158066 | 2.695156254 | 0.014692415 | 0.049833974 | 0.030413638 |
| GO:1904659 | D-glucose transmembrane transport | 10/1052 | 123/28832 | 0.081300813 | 2.228198708 | 2.656340608 | 0.01474503 | 0.04991959 | 0.03046589 |

**Supplementary Table 11:** Summary of GO process terms associated with DEGs exhibiting >2 fold down- or upregulation in marked *Tead4* KD versus unmarked outer control cell clones at the E3.5 stage (RNA-Seq data). Relates to data in Fig. 4.

**Supplementary Table 12:**

| ID | Description | GeneRatio | BgRatio | RichFactor | FoldEnrichment | zScore | pvalue | p.adjust | qvalue |
|---|---|---|---|---|---|---|---|---|---|
| GO:0051015 | actin filament binding | 25/523 | 221/28366 | 0.113122172 | 6.135417838 | 10.50401952 | 5.61E-13 | 4.55E-10 | 3.47E-10 |
| GO:0030695 | GTPase regulator activity | 29/523 | 430/28366 | 0.06744186 | 3.657850505 | 7.611441102 | 2.55E-09 | 5.16E-07 | 3.94E-07 |
| GO:0050839 | cell adhesion molecule binding | 22/523 | 313/28366 | 0.07028754 | 3.812191889 | 6.856645383 | 1.06E-07 | 1.71E-05 | 1.31E-05 |
| GO:0005516 | calmodulin binding | 17/523 | 198/28366 | 0.085858586 | 4.656720165 | 7.076696015 | 1.81E-07 | 2.44E-05 | 1.86E-05 |
| GO:0051020 | GTPase binding | 20/523 | 337/28366 | 0.059347181 | 3.218818617 | 5.615871662 | 5.40E-06 | 0.000546647 | 0.00041771 |
| GO:0005544 | calcium-dependent phospholipid binding | 7/523 | 53/28366 | 0.132075472 | 7.163389733 | 6.155294627 | 5.15E-05 | 0.004171081 | 0.003187259 |
| GO:0000146 | microfilament motor activity | 6/523 | 38/28366 | 0.157894737 | 8.563751635 | 6.394481363 | 6.38E-05 | 0.004701655 | 0.003592688 |
| GO:0017017 | MAP kinase tyrosine/serine/threonine phosphatase activity | 4/523 | 13/28366 | 0.307692308 | 16.68833652 | 7.754133761 | 7.16E-05 | 0.004830786 | 0.00369136 |
| GO:0022804 | active transmembrane transporter activity | 21/523 | 439/28366 | 0.047835991 | 2.59448512 | 4.614501727 | 7.78E-05 | 0.004845367 | 0.003702502 |
| GO:0019838 | growth factor binding | 11/523 | 148/28366 | 0.074324324 | 4.031135342 | 5.067074575 | 9.95E-05 | 0.00575444 | 0.004397155 |
| GO:0044548 | S100 protein binding | 4/523 | 15/28366 | 0.266666667 | 14.46322498 | 7.148168257 | 0.00013268 | 0.006321797 | 0.004830691 |
| GO:0005178 | integrin binding | 11/523 | 155/28366 | 0.070967742 | 3.849084068 | 4.874679842 | 0.000149999 | 0.006749936 | 0.005157846 |
| GO:0140793 | histone H2AXY142 phosphatase activity | 8/523 | 86/28366 | 0.093023256 | 5.045311041 | 5.149270036 | 0.000190635 | 0.007353062 | 0.005618714 |
| GO:0072341 | modified amino acid binding | 9/523 | 115/28366 | 0.07826087 | 4.244642115 | 4.778405848 | 0.000286957 | 0.010565222 | 0.00807323 |
| GO:0048306 | calcium-dependent protein binding | 8/523 | 93/28366 | 0.086021505 | 4.665556446 | 4.852657769 | 0.000327101 | 0.011519633 | 0.008802527 |
| GO:0001223 | transcription coactivator binding | 6/523 | 54/28366 | 0.111111111 | 6.026343743 | 5.066966299 | 0.00046578 | 0.013973391 | 0.010677523 |
| GO:0004620 | phospholipase activity | 8/523 | 99/28366 | 0.080808081 | 4.38279545 | 4.621018426 | 0.000499579 | 0.014452109 | 0.011043327 |
| GO:0001227 | DNA-binding transcription repressor activity, RNA polymerase II-specific | 15/523 | 302/28366 | 0.049668874 | 2.693895382 | 4.056014457 | 0.000540155 | 0.015087094 | 0.01152854 |
| GO:0035091 | phosphatidylinositol binding | 15/523 | 303/28366 | 0.04950495 | 2.685004638 | 4.04147216 | 0.0005589 | 0.015090295 | 0.011530985 |
| GO:0003712 | transcription coregulator activity | 20/523 | 478/28366 | 0.041841004 | 2.269334464 | 3.835884836 | 0.00064272 | 0.016793641 | 0.012832568 |
| GO:0015103 | inorganic anion transmembrane transporter activity | 10/523 | 158/28366 | 0.063291139 | 3.432727449 | 4.202619546 | 0.000732339 | 0.017975598 | 0.013735739 |
| GO:0008375 | acetylglucosaminyltransferase activity | 5/523 | 44/28366 | 0.113636364 | 6.163306101 | 4.697604784 | 0.001254684 | 0.029036981 | 0.022188102 |
| GO:0001618 | virus receptor activity | 5/523 | 45/28366 | 0.111111111 | 6.026343743 | 4.624751219 | 0.001390404 | 0.029249338 | 0.022350371 |
| GO:0043178 | alcohol binding | 7/523 | 90/28366 | 0.077777778 | 4.21844062 | 4.191234823 | 0.001391669 | 0.029249338 | 0.022350371 |
| GO:0015605 | organophosphate ester transmembrane transporter activity | 5/523 | 48/28366 | 0.104166667 | 5.649697259 | 4.418737796 | 0.001862591 | 0.029249338 | 0.022350371 |
| GO:0005347 | ATP transmembrane transporter activity | 3/523 | 14/28366 | 0.214285714 | 11.62223436 | 5.448441241 | 0.001949493 | 0.029249338 | 0.022350371 |
| GO:0005536 | D-glucose binding | 3/523 | 14/28366 | 0.214285714 | 11.62223436 | 5.448441241 | 0.001949493 | 0.029249338 | 0.022350371 |
| GO:0140333 | glycerophospholipid flippase activity | 3/523 | 14/28366 | 0.214285714 | 11.62223436 | 5.448441241 | 0.001949493 | 0.029249338 | 0.022350371 |
| GO:0001786 | phosphatidylserine binding | 6/523 | 71/28366 | 0.084507042 | 4.583416368 | 4.14339009 | 0.001986066 | 0.029249338 | 0.022350371 |
| GO:0140791 | histone H2AXS140 phosphatase activity | 6/523 | 71/28366 | 0.084507042 | 4.583416368 | 4.14339009 | 0.001986066 | 0.029249338 | 0.022350371 |
| GO:0180004 | RNA polymerase II CTD heptapeptide repeat Y1 phosphatase activity | 6/523 | 71/28366 | 0.084507042 | 4.583416368 | 4.14339009 | 0.001986066 | 0.029249338 | 0.022350371 |
| GO:0180005 | RNA polymerase II CTD heptapeptide repeat T4 phosphatase activity | 6/523 | 71/28366 | 0.084507042 | 4.583416368 | 4.14339009 | 0.001986066 | 0.029249338 | 0.022350371 |
| GO:0180006 | RNA polymerase II CTD heptapeptide repeat S2 phosphatase activity | 6/523 | 71/28366 | 0.084507042 | 4.583416368 | 4.14339009 | 0.001986066 | 0.029249338 | 0.022350371 |
| GO:0180007 | RNA polymerase II CTD heptapeptide repeat S5 phosphatase activity | 6/523 | 71/28366 | 0.084507042 | 4.583416368 | 4.14339009 | 0.001986066 | 0.029249338 | 0.022350371 |
| GO:0180008 | RNA polymerase II CTD heptapeptide repeat S7 phosphatase activity | 6/523 | 71/28366 | 0.084507042 | 4.583416368 | 4.14339009 | 0.001986066 | 0.029249338 | 0.022350371 |
| GO:0008514 | organic anion transmembrane transporter activity | 13/523 | 279/28366 | 0.046594982 | 2.527176409 | 3.513361644 | 0.002181214 | 0.030461786 | 0.023276843 |
| GO:0140789 | histone phosphatase activity | 8/523 | 126/28366 | 0.063492063 | 3.443624996 | 3.767660048 | 0.002384227 | 0.032732604 | 0.025012048 |
| GO:0005006 | epidermal growth factor receptor activity | 5/523 | 51/28366 | 0.098039216 | 5.317362126 | 4.229407267 | 0.002442534 | 0.03297421 | 0.025196667 |
| GO:0046982 | protein heterodimerization activity | 14/523 | 319/28366 | 0.043887147 | 2.380311322 | 3.39792203 | 0.002600237 | 0.033278096 | 0.025428877 |
| GO:0017124 | SH3 domain binding | 8/523 | 128/28366 | 0.0625 | 3.389818356 | 3.713959257 | 0.00262938 | 0.033278096 | 0.025428877 |
| GO:0046875 | ephrin receptor binding | 4/523 | 32/28366 | 0.125 | 6.779636711 | 4.483382683 | 0.002728231 | 0.033482836 | 0.025585325 |
| GO:0090482 | vitamin transmembrane transporter activity | 4/523 | 32/28366 | 0.125 | 6.779636711 | 4.483382683 | 0.002728231 | 0.033482836 | 0.025585325 |
| GO:0141003 | histone H2AX kinase activity | 15/523 | 359/28366 | 0.04178273 | 2.266173831 | 3.308959767 | 0.002966586 | 0.035864701 | 0.027405386 |
| GO:0140996 | histone H3 kinase activity | 15/523 | 362/28366 | 0.041436464 | 2.247393385 | 3.273647488 | 0.003206049 | 0.038189705 | 0.029181997 |
| GO:0015662 | P-type ion transporter activity | 4/523 | 34/28366 | 0.117647059 | 6.380834552 | 4.302636978 | 0.003418005 | 0.039551198 | 0.030222358 |
| GO:0140358 | P-type transmembrane transporter activity | 4/523 | 34/28366 | 0.117647059 | 6.380834552 | 4.302636978 | 0.003418005 | 0.039551198 | 0.030222358 |
| GO:0035173 | histone kinase activity | 15/523 | 366/28366 | 0.040983607 | 2.222831709 | 3.227100149 | 0.003549845 | 0.039935753 | 0.030516209 |

**Supplementary Table 12:** Summary of GO function terms associated with DEGs exhibiting >2 fold downregulation in marked *Tead4* KD versus unmarked outer control cell clones at the E3.5 stage (RNA-Seq data). Relates to data in Fig. S8.

**Supplementary Table 13:**

| ID | Description | GeneRatio | BgRatio | RichFactor | FoldEnrichment | zScore | pvalue | p.adjust | qvalue |
|---|---|---|---|---|---|---|---|---|---|
| GO:0007015 | actin filament organization | 40/519 | 474/28832 | 0.084388186 | 4.688015739 | 10.96136716 | 6.89E-16 | 3.01E-12 | 2.25E-12 |
| GO:0022604 | regulation of cell morphogenesis | 26/519 | 283/28832 | 0.091872792 | 5.103807948 | 9.393022945 | 1.37E-11 | 2.99E-08 | 2.23E-08 |
| GO:0001890 | placenta development | 20/519 | 199/28832 | 0.100502513 | 5.583214725 | 8.783829629 | 6.57E-10 | 9.58E-07 | 7.16E-07 |
| GO:0007264 | small GTPase-mediated signal transduction | 29/519 | 440/28832 | 0.065909091 | 3.661446838 | 7.616697859 | 2.51E-09 | 2.18E-06 | 1.63E-06 |
| GO:0032970 | regulation of actin filament-based process | 28/519 | 416/28832 | 0.067307692 | 3.739143323 | 7.619044983 | 3.00E-09 | 2.18E-06 | 1.63E-06 |
| GO:0110053 | regulation of actin filament organization | 22/519 | 281/28832 | 0.078291815 | 4.34934414 | 7.638759878 | 1.02E-08 | 4.94E-06 | 3.69E-06 |
| GO:0045216 | cell-cell junction organization | 20/519 | 239/28832 | 0.083682008 | 4.648785482 | 7.668987098 | 1.56E-08 | 6.84E-06 | 5.11E-06 |
| GO:1902106 | negative regulation of leukocyte differentiation | 14/519 | 126/28832 | 0.111111111 | 6.172554057 | 7.878155666 | 6.89E-08 | 2.43E-05 | 1.82E-05 |
| GO:0002064 | epithelial cell development | 20/519 | 262/28832 | 0.076335878 | 4.240685993 | 7.134327884 | 7.22E-08 | 2.43E-05 | 1.82E-05 |
| GO:0031589 | cell-substrate adhesion | 24/519 | 376/28832 | 0.063829787 | 3.545935309 | 6.727829056 | 1.09E-07 | 3.27E-05 | 2.44E-05 |
| GO:0002573 | myeloid leukocyte differentiation | 21/519 | 295/28832 | 0.071186441 | 3.954619379 | 6.906038257 | 1.12E-07 | 3.27E-05 | 2.44E-05 |
| GO:0030100 | regulation of endocytosis | 23/519 | 352/28832 | 0.065340909 | 3.629882641 | 6.721377615 | 1.33E-07 | 3.41E-05 | 2.55E-05 |
| GO:0051336 | regulation of hydrolase activity | 25/519 | 424/28832 | 0.058962264 | 3.275529865 | 6.390954504 | 2.68E-07 | 5.58E-05 | 4.18E-05 |
| GO:0042060 | wound healing | 24/519 | 403/28832 | 0.05955335 | 3.308366442 | 6.318253855 | 3.89E-07 | 7.72E-05 | 5.77E-05 |
| GO:0031346 | positive regulation of cell projection organization | 26/519 | 465/28832 | 0.055913978 | 3.106188493 | 6.199227799 | 4.29E-07 | 8.12E-05 | 6.07E-05 |
| GO:0001822 | kidney development | 23/519 | 377/28832 | 0.061007958 | 3.389174243 | 6.322077157 | 4.46E-07 | 8.12E-05 | 6.07E-05 |
| GO:0007162 | negative regulation of cell adhesion | 21/519 | 322/28832 | 0.065217391 | 3.62302086 | 6.408430209 | 4.80E-07 | 8.17E-05 | 6.11E-05 |
| GO:0150105 | protein localization to cell-cell junction | 7/519 | 28/28832 | 0.25 | 13.88824663 | 9.237767768 | 5.02E-07 | 8.17E-05 | 6.11E-05 |
| GO:0007163 | establishment or maintenance of cell polarity | 18/519 | 243/28832 | 0.074074074 | 4.115036038 | 6.602179342 | 5.05E-07 | 8.17E-05 | 6.11E-05 |
| GO:0030048 | actin filament-based movement | 13/519 | 128/28832 | 0.1015625 | 5.642100193 | 7.126378276 | 5.78E-07 | 9.02E-05 | 6.74E-05 |
| GO:0007009 | plasma membrane organization | 15/519 | 178/28832 | 0.084269663 | 4.681431448 | 6.670443013 | 8.96E-07 | 0.000126256 | 9.44E-05 |
| GO:0001778 | plasma membrane repair | 7/519 | 31/28832 | 0.225806452 | 12.54422276 | 8.706875796 | 1.06E-06 | 0.000140841 | 0.000105304 |
| GO:1905244 | regulation of modification of synaptic structure | 7/519 | 31/28832 | 0.225806452 | 12.54422276 | 8.706875796 | 1.06E-06 | 0.000140841 | 0.000105304 |
| GO:0010721 | negative regulation of cell development | 20/519 | 315/28832 | 0.063492063 | 3.527173747 | 6.106031605 | 1.36E-06 | 0.000170565 | 0.000127528 |
| GO:1902903 | regulation of supramolecular fiber organization | 23/519 | 403/28832 | 0.05707196 | 3.170517841 | 5.94094701 | 1.41E-06 | 0.000170565 | 0.000127528 |
| GO:0034446 | substrate adhesion-dependent cell spreading | 11/519 | 101/28832 | 0.108910891 | 6.050325264 | 6.88375259 | 2.14E-06 | 0.000233496 | 0.000174581 |
| GO:0060485 | mesenchyme development | 20/519 | 329/28832 | 0.060790274 | 3.377081247 | 5.871070236 | 2.64E-06 | 0.000274795 | 0.000205459 |
| GO:0001894 | tissue homeostasis | 19/519 | 303/28832 | 0.062706271 | 3.483520606 | 5.883915065 | 2.98E-06 | 0.00029642 | 0.000221628 |
| GO:0060249 | anatomical structure homeostasis | 19/519 | 303/28832 | 0.062706271 | 3.483520606 | 5.883915065 | 2.98E-06 | 0.00029642 | 0.000221628 |
| GO:0051345 | positive regulation of hydrolase activity | 16/519 | 223/28832 | 0.071748879 | 3.985864503 | 6.060267244 | 3.27E-06 | 0.000315734 | 0.000236069 |
| GO:0038127 | ERBB signaling pathway | 14/519 | 173/28832 | 0.080924855 | 4.495617406 | 6.243627948 | 3.33E-06 | 0.000315734 | 0.000236069 |
| GO:0050678 | regulation of epithelial cell proliferation | 23/519 | 434/28832 | 0.052995392 | 2.944052281 | 5.524964793 | 4.84E-06 | 0.000429692 | 0.000321273 |
| GO:0030879 | mammary gland development | 14/519 | 179/28832 | 0.078212291 | 4.344926319 | 6.077830845 | 4.95E-06 | 0.000429692 | 0.000321273 |
| GO:0006942 | regulation of striated muscle contraction | 10/519 | 91/28832 | 0.10989011 | 6.104723793 | 6.60332381 | 5.71E-06 | 0.000480228 | 0.000359058 |
| GO:0099563 | modification of synaptic structure | 8/519 | 55/28832 | 0.145454545 | 8.080434402 | 7.116051416 | 6.05E-06 | 0.000498918 | 0.000373032 |
| GO:0008544 | epidermis development | 22/519 | 410/28832 | 0.053658537 | 2.980891959 | 5.469479355 | 6.38E-06 | 0.000516091 | 0.000385872 |
| GO:0006936 | muscle contraction | 19/519 | 330/28832 | 0.057575758 | 3.198505284 | 5.438353118 | 1.01E-05 | 0.000764684 | 0.00057174 |
| GO:0042692 | muscle cell differentiation | 24/519 | 488/28832 | 0.049180328 | 2.732114091 | 5.22488124 | 1.06E-05 | 0.000771273 | 0.000576666 |
| GO:0045444 | fat cell differentiation | 17/519 | 275/28832 | 0.061818182 | 3.434184621 | 5.491413637 | 1.18E-05 | 0.000817765 | 0.000611428 |
| GO:0032570 | response to progesterone | 7/519 | 44/28832 | 0.159090909 | 8.837975127 | 7.044417052 | 1.27E-05 | 0.000837581 | 0.000626244 |
| GO:0045862 | positive regulation of proteolysis | 15/519 | 222/28832 | 0.067567568 | 3.75358017 | 5.576168275 | 1.35E-05 | 0.000866442 | 0.000647823 |
| GO:2000241 | regulation of reproductive process | 15/519 | 225/28832 | 0.066666667 | 3.703532434 | 5.511975426 | 1.59E-05 | 0.000962604 | 0.000719721 |
| GO:0060348 | bone development | 16/519 | 255/28832 | 0.062745098 | 3.485677585 | 5.397928856 | 1.77E-05 | 0.001031862 | 0.000771504 |
| GO:0097529 | myeloid leukocyte migration | 16/519 | 255/28832 | 0.062745098 | 3.485677585 | 5.397928856 | 1.77E-05 | 0.001031862 | 0.000771504 |
| GO:0072659 | protein localization to plasma membrane | 18/519 | 314/28832 | 0.057324841 | 3.184566106 | 5.269762836 | 1.83E-05 | 0.001053085 | 0.000787373 |
| GO:0016055 | Wnt signaling pathway | 23/519 | 474/28832 | 0.048523207 | 2.69560905 | 5.039618774 | 1.99E-05 | 0.001116215 | 0.000834573 |
| GO:0006935 | chemotaxis | 23/519 | 475/28832 | 0.048421053 | 2.689934084 | 5.028136 | 2.06E-05 | 0.001139125 | 0.000851703 |
| GO:0042063 | gliogenesis | 21/519 | 412/28832 | 0.050970874 | 2.831584264 | 5.069721032 | 2.22E-05 | 0.001184942 | 0.000885959 |
| GO:0098742 | cell-cell adhesion via plasma-membrane adhesion molecules | 15/519 | 232/28832 | 0.064655172 | 3.591787921 | 5.366375089 | 2.27E-05 | 0.001184942 | 0.000885959 |
| GO:0007589 | body fluid secretion | 11/519 | 129/28832 | 0.085271318 | 4.737076369 | 5.759486573 | 2.28E-05 | 0.001184942 | 0.000885959 |
| GO:0050878 | regulation of body fluid levels | 20/519 | 384/28832 | 0.052083333 | 2.893384714 | 5.057072695 | 2.55E-05 | 0.001296714 | 0.000969529 |
| GO:0046434 | organophosphate catabolic process | 16/519 | 264/28832 | 0.060606061 | 3.366847667 | 5.230617224 | 2.71E-05 | 0.001306151 | 0.000976585 |
| GO:0003007 | heart morphogenesis | 17/519 | 301/28832 | 0.056478405 | 3.137544089 | 5.047315116 | 3.75E-05 | 0.001761178 | 0.0013168 |
| GO:0070372 | regulation of ERK1 and ERK2 cascade | 18/519 | 333/28832 | 0.054054054 | 3.002864136 | 4.977134574 | 3.97E-05 | 0.001825941 | 0.001365223 |
| GO:0044409 | symbiont entry into host | 9/519 | 92/28832 | 0.097826087 | 5.434531289 | 5.767914882 | 4.28E-05 | 0.001893767 | 0.001415935 |
| GO:0061448 | connective tissue development | 18/519 | 335/28832 | 0.053731343 | 2.984936589 | 4.947548464 | 4.29E-05 | 0.001893767 | 0.001415935 |
| GO:0034109 | homotypic cell-cell adhesion | 9/519 | 93/28832 | 0.096774194 | 5.376095469 | 5.722858706 | 4.67E-05 | 0.002006871 | 0.001500501 |
| GO:0050803 | regulation of synapse structure or activity | 20/519 | 401/28832 | 0.049875312 | 2.770722519 | 4.8344516 | 4.68E-05 | 0.002006871 | 0.001500501 |
| GO:0032535 | regulation of cellular component size | 20/519 | 404/28832 | 0.04950495 | 2.750147847 | 4.796371953 | 5.19E-05 | 0.002203416 | 0.001647454 |
| GO:0055117 | regulation of cardiac muscle contraction | 8/519 | 74/28832 | 0.108108108 | 6.005728272 | 5.837466195 | 5.56E-05 | 0.00230892 | 0.001726337 |
| GO:0001666 | response to hypoxia | 18/519 | 342/28832 | 0.052631579 | 2.923841395 | 4.845701762 | 5.60E-05 | 0.00230892 | 0.001726337 |
| GO:0030162 | regulation of proteolysis | 22/519 | 473/28832 | 0.046511628 | 2.583859838 | 4.702431176 | 5.64E-05 | 0.00230892 | 0.001726337 |
| GO:0045214 | sarcomere organization | 7/519 | 55/28832 | 0.127272727 | 7.070380102 | 6.100916196 | 5.65E-05 | 0.00230892 | 0.001726337 |
| GO:0045859 | regulation of protein kinase activity | 22/519 | 477/28832 | 0.046121593 | 2.56219225 | 4.657999421 | 6.38E-05 | 0.002509895 | 0.001876602 |
| GO:0051495 | positive regulation of cytoskeleton organization | 13/519 | 201/28832 | 0.064676617 | 3.592979227 | 4.994584848 | 7.86E-05 | 0.002986673 | 0.00223308 |
| GO:0043588 | skin development | 18/519 | 352/28832 | 0.051136364 | 2.840777719 | 4.704606191 | 8.09E-05 | 0.003047519 | 0.002278574 |
| GO:0120192 | tight junction assembly | 9/519 | 100/28832 | 0.09 | 4.999768786 | 5.424660278 | 8.26E-05 | 0.003084591 | 0.002306292 |
| GO:0009913 | epidermal cell differentiation | 15/519 | 261/28832 | 0.057471264 | 3.192700374 | 4.817895893 | 8.72E-05 | 0.003231167 | 0.002415884 |
| GO:1900024 | regulation of substrate adhesion-dependent cell spreading | 7/519 | 59/28832 | 0.118644068 | 6.591032298 | 5.820308727 | 8.93E-05 | 0.00325093 | 0.00243066 |
| GO:0007343 | egg activation | 4/519 | 14/28832 | 0.285714286 | 15.87228186 | 7.535816149 | 9.00E-05 | 0.00325093 | 0.00243066 |

| | | | | | | | | | |
|---|---|---|---|---|---|---|---|---|---|
| GO:0007605 | sensory perception of sound | 16/519 | 293/28832 | 0.054607509 | 3.033610185 | 4.736992861 | 9.35E-05 | 0.003347393 | 0.002502784 |
| GO:0008016 | regulation of heart contraction | 13/519 | 205/28832 | 0.063414634 | 3.522872315 | 4.908003572 | 9.59E-05 | 0.003408005 | 0.002548102 |
| GO:0051216 | cartilage development | 14/519 | 235/28832 | 0.059574468 | 3.309539622 | 4.813042519 | 0.000101959 | 0.003564473 | 0.002665091 |
| GO:0006909 | phagocytosis | 14/519 | 236/28832 | 0.059322034 | 3.295516149 | 4.794069185 | 0.00010664 | 0.003669413 | 0.002743552 |
| GO:0032878 | regulation of establishment or maintenance of cell polarity | 5/519 | 27/28832 | 0.185185185 | 10.28759009 | 6.536895909 | 0.000107856 | 0.003682273 | 0.002753167 |
| GO:0043254 | regulation of protein-containing complex assembly | 21/519 | 461/28832 | 0.045553145 | 2.530613268 | 4.485379586 | 0.000110492 | 0.003743016 | 0.002798584 |
| GO:0048144 | fibroblast proliferation | 11/519 | 154/28832 | 0.071428571 | 3.968070465 | 5.000123428 | 0.000115052 | 0.003808546 | 0.002847579 |
| GO:0031274 | positive regulation of pseudopodium assembly | 4/519 | 15/28832 | 0.266666667 | 14.81412974 | 7.245450007 | 0.000121003 | 0.003859189 | 0.002885444 |
| GO:0010927 | cellular component assembly involved in morphogenesis | 11/519 | 155/28832 | 0.070967742 | 3.942470011 | 4.973150745 | 0.000121869 | 0.003859189 | 0.002885444 |
| GO:0006672 | ceramide metabolic process | 9/519 | 106/28832 | 0.08490566 | 4.716763006 | 5.190399805 | 0.000129657 | 0.003934744 | 0.002941936 |
| GO:0038066 | p38MAPK cascade | 7/519 | 63/28832 | 0.111111111 | 6.172554057 | 5.564594436 | 0.000136194 | 0.00410462 | 0.003068948 |
| GO:1900744 | regulation of p38MAPK cascade | 6/519 | 45/28832 | 0.133333333 | 7.407064868 | 5.823549369 | 0.000148403 | 0.00435915 | 0.003259256 |
| GO:0055057 | neuroblast division | 4/519 | 16/28832 | 0.25 | 13.88824663 | 6.981641897 | 0.000159046 | 0.004513191 | 0.00337443 |
| GO:0060047 | heart contraction | 14/519 | 247/28832 | 0.056680162 | 3.148752272 | 4.591835431 | 0.000171788 | 0.00475134 | 0.003552489 |
| GO:0015711 | organic anion transport | 21/519 | 477/28832 | 0.044025157 | 2.445728966 | 4.310740042 | 0.000176454 | 0.004819402 | 0.003603378 |
| GO:0052372 | modulation by symbiont of entry into host | 5/519 | 30/28832 | 0.166666667 | 9.258831085 | 6.127572742 | 0.000182107 | 0.004872942 | 0.003643409 |
| GO:0061351 | neural precursor cell proliferation | 13/519 | 219/28832 | 0.059360731 | 3.297665866 | 4.621126104 | 0.000185105 | 0.004872942 | 0.003643409 |
| GO:0051014 | actin filament severing | 4/519 | 17/28832 | 0.235294118 | 13.07129094 | 6.740458726 | 0.000205032 | 0.005323002 | 0.00397991 |
| GO:0003014 | renal system process | 10/519 | 141/28832 | 0.070921986 | 3.939928121 | 4.737988623 | 0.000247024 | 0.00613337 | 0.004585808 |
| GO:0051701 | biological process involved in interaction with host | 10/519 | 141/28832 | 0.070921986 | 3.939928121 | 4.737988623 | 0.000247024 | 0.00613337 | 0.004585808 |
| GO:0051446 | positive regulation of meiotic cell cycle | 5/519 | 32/28832 | 0.15625 | 8.680154143 | 5.885308993 | 0.000249826 | 0.00613337 | 0.004585808 |
| GO:0031269 | pseudopodium assembly | 4/519 | 18/28832 | 0.222222222 | 12.34510811 | 6.518740652 | 0.000259874 | 0.006274298 | 0.004691177 |
| GO:0048738 | cardiac muscle tissue development | 15/519 | 289/28832 | 0.051903114 | 2.883373002 | 4.3566838 | 0.000265397 | 0.006372445 | 0.004764559 |
| GO:1900076 | regulation of cellular response to insulin stimulus | 8/519 | 94/28832 | 0.085106383 | 4.727913746 | 4.901422041 | 0.000300069 | 0.006993308 | 0.005228767 |
| GO:0007411 | axon guidance | 14/519 | 261/28832 | 0.053639847 | 2.979853683 | 4.350219956 | 0.00030225 | 0.006993308 | 0.005228767 |
| GO:1901292 | nucleoside phosphate catabolic process | 12/519 | 201/28832 | 0.059701493 | 3.31659621 | 4.462217131 | 0.000307595 | 0.007043331 | 0.005266169 |
| GO:0046632 | alpha-beta T cell differentiation | 10/519 | 145/28832 | 0.068965517 | 3.831240449 | 4.627418476 | 0.000309455 | 0.007043331 | 0.005266169 |
| GO:0032102 | negative regulation of response to external stimulus | 19/519 | 429/28832 | 0.044289044 | 2.46038868 | 4.126060433 | 0.000328713 | 0.007328968 | 0.005479734 |
| GO:0040013 | negative regulation of locomotion | 17/519 | 362/28832 | 0.046961326 | 2.608841908 | 4.170561307 | 0.000343615 | 0.007473062 | 0.005587471 |
| GO:0048286 | lung alveolus development | 7/519 | 73/28832 | 0.095890411 | 5.326998707 | 5.011663376 | 0.000343727 | 0.007473062 | 0.005587471 |
| GO:1903035 | negative regulation of response to wounding | 8/519 | 96/28832 | 0.083333333 | 4.629415543 | 4.822583337 | 0.000346458 | 0.007495162 | 0.005603994 |
| GO:0050920 | regulation of chemotaxis | 13/519 | 236/28832 | 0.055084746 | 3.060122138 | 4.302460719 | 0.000381662 | 0.008038231 | 0.006010037 |
| GO:0090025 | regulation of monocyte chemotaxis | 5/519 | 35/28832 | 0.142857143 | 7.93614093 | 5.55902843 | 0.000385255 | 0.008038231 | 0.006010037 |
| GO:0048678 | response to axon injury | 8/519 | 98/28832 | 0.081632653 | 4.534937674 | 4.745886445 | 0.000398528 | 0.008242986 | 0.006163128 |
| GO:0031268 | pseudopodium organization | 4/519 | 20/28832 | 0.2 | 11.1105973 | 6.12386623 | 0.000399889 | 0.008242986 | 0.006163128 |
| GO:0006644 | phospholipid metabolic process | 17/519 | 368/28832 | 0.046195652 | 2.566306442 | 4.094239705 | 0.000414524 | 0.008504544 | 0.006358691 |
| GO:0050767 | regulation of neurogenesis | 20/519 | 473/28832 | 0.042283298 | 2.348963489 | 4.005031136 | 0.00041871 | 0.008550286 | 0.006392891 |
| GO:0060562 | epithelial tube morphogenesis | 18/519 | 403/28832 | 0.044665012 | 2.481274832 | 4.054412785 | 0.000423319 | 0.008604204 | 0.006433204 |
| GO:0045598 | regulation of fat cell differentiation | 10/519 | 151/28832 | 0.066225166 | 3.679005729 | 4.468744852 | 0.000427433 | 0.00864761 | 0.006465659 |
| GO:0010976 | positive regulation of neuron projection development | 13/519 | 239/28832 | 0.054393305 | 3.021710563 | 4.249214513 | 0.000430346 | 0.008666405 | 0.006479711 |
| GO:0051260 | protein homooligomerization | 12/519 | 211/28832 | 0.056872038 | 3.159411555 | 4.262406195 | 0.000476642 | 0.009340466 | 0.006983694 |
| GO:0006066 | alcohol metabolic process | 17/519 | 373/28832 | 0.045576408 | 2.531905552 | 4.031783244 | 0.000482936 | 0.009416516 | 0.007040555 |
| GO:0015693 | magnesium ion transport | 4/519 | 21/28832 | 0.19047619 | 10.58152124 | 5.94683016 | 0.000486987 | 0.009416516 | 0.007040555 |
| GO:2001185 | regulation of CD8-positive, alpha-beta T cell activation | 4/519 | 21/28832 | 0.19047619 | 10.58152124 | 5.94683016 | 0.000486987 | 0.009416516 | 0.007040555 |
| GO:0046466 | membrane lipid catabolic process | 5/519 | 37/28832 | 0.135135135 | 7.50716034 | 5.362340854 | 0.000502234 | 0.009544527 | 0.007136267 |
| GO:0048145 | regulation of fibroblast proliferation | 9/519 | 127/28832 | 0.070866142 | 3.936825816 | 4.490780845 | 0.000502344 | 0.009544527 | 0.007136267 |
| GO:1905954 | positive regulation of lipid localization | 9/519 | 127/28832 | 0.070866142 | 3.936825816 | 4.490780845 | 0.000502344 | 0.009544527 | 0.007136267 |
| GO:0045766 | positive regulation of angiogenesis | 11/519 | 184/28832 | 0.059782609 | 3.321102455 | 4.276383605 | 0.000535708 | 0.010047405 | 0.00751226 |
| GO:2000146 | negative regulation of cell motility | 16/519 | 344/28832 | 0.046511628 | 2.583859838 | 4.001156835 | 0.000563745 | 0.010438845 | 0.007804932 |
| GO:0010876 | lipid localization | 20/519 | 485/28832 | 0.041237113 | 2.290844805 | 3.88161034 | 0.000573112 | 0.010567504 | 0.007901128 |
| GO:0036376 | sodium ion export across plasma membrane | 4/519 | 22/28832 | 0.181818182 | 10.100543 | 5.781327939 | 0.000586785 | 0.010729084 | 0.008021938 |
| GO:0032370 | positive regulation of lipid transport | 8/519 | 104/28832 | 0.076923077 | 4.273306655 | 4.527633395 | 0.000594044 | 0.010816546 | 0.008087332 |
| GO:0150172 | regulation of phosphatidylcholine metabolic process | 3/519 | 10/28832 | 0.3 | 16.66589595 | 6.708319958 | 0.000633306 | 0.011389078 | 0.008515403 |
| GO:0006869 | lipid transport | 19/519 | 456/28832 | 0.041666667 | 2.314707771 | 3.831392557 | 0.000689377 | 0.012143665 | 0.009079594 |
| GO:0031345 | negative regulation of cell projection organization | 12/519 | 220/28832 | 0.054545455 | 3.030162901 | 4.092500177 | 0.000690004 | 0.012143665 | 0.009079594 |
| GO:0016197 | endosomal transport | 14/519 | 284/28832 | 0.049295775 | 2.738527504 | 3.986332123 | 0.000699227 | 0.012143665 | 0.009079594 |
| GO:0070254 | mucus secretion | 4/519 | 23/28832 | 0.173913043 | 9.661388959 | 5.626106588 | 0.000700275 | 0.012143665 | 0.009079594 |
| GO:0010038 | response to metal ion | 17/519 | 386/28832 | 0.044041451 | 2.446634121 | 3.874024307 | 0.000708006 | 0.012229198 | 0.009143546 |
| GO:0035924 | cellular response to vascular endothelial growth factor stimulus | 8/519 | 107/28832 | 0.074766355 | 4.153494319 | 4.424605049 | 0.000717413 | 0.012342887 | 0.009228549 |
| GO:1903037 | regulation of leukocyte cell-cell adhesion | 17/519 | 387/28832 | 0.043927649 | 2.440312069 | 3.862155013 | 0.000728537 | 0.012436349 | 0.009298429 |
| GO:0034113 | heterotypic cell-cell adhesion | 6/519 | 61/28832 | 0.098360656 | 5.464228182 | 4.72557229 | 0.000793974 | 0.013448325 | 0.010055064 |
| GO:0032868 | response to insulin | 16/519 | 356/28832 | 0.04494382 | 2.496763439 | 3.847328419 | 0.000812231 | 0.013629466 | 0.010190501 |
| GO:0015748 | organophosphate ester transport | 10/519 | 164/28832 | 0.06097561 | 3.387377226 | 4.151115912 | 0.000814025 | 0.013629466 | 0.010190501 |
| GO:0051258 | protein polymerization | 14/519 | 290/28832 | 0.048275862 | 2.681868314 | 3.897349687 | 0.000856102 | 0.014062209 | 0.010514054 |
| GO:0048840 | otolith development | 3/519 | 11/28832 | 0.272727273 | 15.1508145 | 6.355413987 | 0.000859178 | 0.014062209 | 0.010514054 |
| GO:0060717 | chorion development | 3/519 | 11/28832 | 0.272727273 | 15.1508145 | 6.355413987 | 0.000859178 | 0.014062209 | 0.010514054 |
| GO:1904338 | regulation of dopaminergic neuron differentiation | 3/519 | 11/28832 | 0.272727273 | 15.1508145 | 6.355413987 | 0.000859178 | 0.014062209 | 0.010514054 |
| GO:0060541 | respiratory system development | 14/519 | 293/28832 | 0.04778157 | 2.654408912 | 3.853699821 | 0.000945061 | 0.015183521 | 0.011352438 |
| GO:0015867 | ATP transport | 4/519 | 25/28832 | 0.16 | 8.888477842 | 5.342380119 | 0.00097233 | 0.01545077 | 0.011552256 |
| GO:1905475 | regulation of protein localization to membrane | 12/519 | 230/28832 | 0.052173913 | 2.898416688 | 3.913612722 | 0.001015552 | 0.015785771 | 0.01180273 |
| GO:0046850 | regulation of bone remodeling | 6/519 | 64/28832 | 0.09375 | 5.208092486 | 4.56290068 | 0.00102462 | 0.015785771 | 0.01180273 |
| GO:1903532 | positive regulation of secretion by cell | 17/519 | 400/28832 | 0.0425 | 2.361001927 | 3.711125301 | 0.001045448 | 0.016030204 | 0.011985488 |
| GO:0038084 | vascular endothelial growth factor signaling pathway | 7/519 | 88/28832 | 0.079545455 | 4.418987563 | 4.348964299 | 0.001063348 | 0.016191048 | 0.012105748 |

| | | | | | | | | | |
|---|---|---|---|---|---|---|---|---|---|
| GO:0045022 | early endosome to late endosome transport | 5/519 | 44/28832 | 0.113636364 | 6.312839376 | 4.774939406 | 0.001128566 | 0.016951253 | 0.01267414 |
| GO:0098911 | regulation of ventricular cardiac muscle cell action potential | 3/519 | 12/28832 | 0.25 | 13.88824663 | 6.04585964 | 0.001130303 | 0.016951253 | 0.01267414 |
| GO:0046890 | regulation of lipid biosynthetic process | 12/519 | 233/28832 | 0.051502146 | 2.861098018 | 3.861823053 | 0.001135046 | 0.016951253 | 0.01267414 |
| GO:0045682 | regulation of epidermis development | 6/519 | 66/28832 | 0.090909091 | 5.050271501 | 4.460021511 | 0.00120481 | 0.017667848 | 0.013209924 |
| GO:0045576 | mast cell activation | 7/519 | 90/28832 | 0.077777778 | 4.32078784 | 4.271933463 | 0.001213643 | 0.017678727 | 0.013218058 |
| GO:0043603 | amide metabolic process | 19/519 | 479/28832 | 0.039665971 | 2.203563139 | 3.596314911 | 0.00122673 | 0.017809996 | 0.013316206 |
| GO:0090287 | regulation of cellular response to growth factor stimulus | 15/519 | 336/28832 | 0.044642857 | 2.480044041 | 3.694646686 | 0.001248914 | 0.018043724 | 0.01349096 |
| GO:0009132 | nucleoside diphosphate metabolic process | 9/519 | 146/28832 | 0.061643836 | 3.424499169 | 3.976344998 | 0.001358146 | 0.018901591 | 0.014132371 |
| GO:1901136 | carbohydrate derivative catabolic process | 13/519 | 271/28832 | 0.04797048 | 2.664903412 | 3.728271849 | 0.0013693 | 0.018996322 | 0.014203199 |
| GO:0050764 | regulation of phagocytosis | 8/519 | 119/28832 | 0.067226891 | 3.734654555 | 4.047224135 | 0.0014323 | 0.019554757 | 0.014620731 |
| GO:0035335 | peptidyl-tyrosine dephosphorylation | 4/519 | 28/28832 | 0.142857143 | 7.93614093 | 4.971541977 | 0.001508134 | 0.020119968 | 0.015043329 |
| GO:0098901 | regulation of cardiac muscle cell action potential | 4/519 | 28/28832 | 0.142857143 | 7.93614093 | 4.971541977 | 0.001508134 | 0.020119968 | 0.015043329 |
| GO:0007595 | lactation | 6/519 | 69/28832 | 0.086956522 | 4.830694479 | 4.313259207 | 0.001519357 | 0.020119968 | 0.015043329 |
| GO:0150116 | regulation of cell-substrate junction organization | 6/519 | 69/28832 | 0.086956522 | 4.830694479 | 4.313259207 | 0.001519357 | 0.020119968 | 0.015043329 |
| GO:0003002 | regionalization | 18/519 | 452/28832 | 0.039823009 | 2.212287074 | 3.517135367 | 0.001562241 | 0.020563235 | 0.015374752 |
| GO:0045936 | negative regulation of phosphate metabolic process | 16/519 | 381/28832 | 0.041994751 | 2.332933817 | 3.546033987 | 0.001638678 | 0.021376185 | 0.015982579 |
| GO:0051250 | negative regulation of lymphocyte activation | 10/519 | 180/28832 | 0.055555556 | 3.086277028 | 3.801468052 | 0.001644119 | 0.021383339 | 0.015987928 |
| GO:0048644 | muscle organ morphogenesis | 7/519 | 95/28832 | 0.073684211 | 4.093377954 | 4.088788648 | 0.00166256 | 0.021511181 | 0.016083513 |
| GO:0002548 | monocyte chemotaxis | 5/519 | 48/28832 | 0.104166667 | 5.786769428 | 4.493742075 | 0.001677738 | 0.021511181 | 0.016083513 |
| GO:0071364 | cellular response to epidermal growth factor stimulus | 5/519 | 48/28832 | 0.104166667 | 5.786769428 | 4.493742075 | 0.001677738 | 0.021511181 | 0.016083513 |
| GO:0014909 | smooth muscle cell migration | 8/519 | 122/28832 | 0.06557377 | 3.642818788 | 3.960511623 | 0.001678561 | 0.021511181 | 0.016083513 |
| GO:0051348 | negative regulation of transferase activity | 11/519 | 212/28832 | 0.051886792 | 2.882466281 | 3.724610893 | 0.001696287 | 0.021611587 | 0.016158585 |
| GO:0098739 | import across plasma membrane | 11/519 | 212/28832 | 0.051886792 | 2.882466281 | 3.724610893 | 0.001696287 | 0.021611587 | 0.016158585 |
| GO:0016042 | lipid catabolic process | 15/519 | 347/28832 | 0.043227666 | 2.40142593 | 3.555882173 | 0.00171148 | 0.021697719 | 0.016222984 |
| GO:0035148 | tube formation | 10/519 | 181/28832 | 0.055248619 | 3.069225774 | 3.780923253 | 0.001712978 | 0.021697719 | 0.016222984 |
| GO:0031579 | membrane raft organization | 4/519 | 29/28832 | 0.137931034 | 7.662480898 | 4.860004833 | 0.001724792 | 0.021721441 | 0.016240721 |
| GO:0072523 | purine-containing compound catabolic process | 10/519 | 182/28832 | 0.054945055 | 3.052361896 | 3.760520075 | 0.001784152 | 0.022276412 | 0.016655662 |
| GO:0001765 | membrane raft assembly | 3/519 | 14/28832 | 0.214285714 | 11.9042114 | 5.525186592 | 0.001820684 | 0.02234941 | 0.016710242 |
| GO:0030195 | negative regulation of blood coagulation | 5/519 | 49/28832 | 0.102040816 | 5.668672093 | 4.428370722 | 0.001841143 | 0.022537235 | 0.016850675 |
| GO:0006509 | membrane protein ectodomain proteolysis | 5/519 | 50/28832 | 0.1 | 5.555298651 | 4.364775997 | 0.002015894 | 0.024268482 | 0.018145096 |
| GO:1900047 | negative regulation of hemostasis | 5/519 | 50/28832 | 0.1 | 5.555298651 | 4.364775997 | 0.002015894 | 0.024268482 | 0.018145096 |
| GO:0140895 | cell surface toll-like receptor signaling pathway | 6/519 | 73/28832 | 0.082191781 | 4.565998891 | 4.130249916 | 0.002031518 | 0.024322558 | 0.018185528 |
| GO:2000177 | regulation of neural precursor cell proliferation | 8/519 | 126/28832 | 0.063492063 | 3.527173747 | 3.849059469 | 0.002057698 | 0.024568695 | 0.01836956 |
| GO:0007159 | leukocyte cell-cell adhesion | 17/519 | 427/28832 | 0.039812646 | 2.211711407 | 3.415358716 | 0.0020892 | 0.024876848 | 0.01859996 |
| GO:0016050 | vesicle organization | 16/519 | 391/28832 | 0.040920716 | 2.27326799 | 3.432071731 | 0.002125966 | 0.025177426 | 0.018824697 |
| GO:0019827 | stem cell population maintenance | 11/519 | 219/28832 | 0.050228311 | 2.790332656 | 3.60076419 | 0.002188098 | 0.025635357 | 0.019167084 |
| GO:0046636 | negative regulation of alpha-beta T cell activation | 5/519 | 51/28832 | 0.098039216 | 5.446371227 | 4.302872275 | 0.00220245 | 0.025665885 | 0.019189909 |
| GO:0050819 | negative regulation of coagulation | 5/519 | 51/28832 | 0.098039216 | 5.446371227 | 4.302872275 | 0.00220245 | 0.025665885 | 0.019189909 |
| GO:0086069 | bundle of His cell to Purkinje myocyte communication | 3/519 | 15/28832 | 0.2 | 11.1105973 | 5.30296361 | 0.002245605 | 0.02596109 | 0.019410628 |
| GO:0033673 | negative regulation of kinase activity | 10/519 | 188/28832 | 0.053191489 | 2.954946091 | 3.64097306 | 0.002263054 | 0.025987894 | 0.019430669 |
| GO:0043393 | regulation of protein binding | 10/519 | 188/28832 | 0.053191489 | 2.954946091 | 3.64097306 | 0.002263054 | 0.025987894 | 0.019430669 |
| GO:0044843 | cell cycle G1/S phase transition | 12/519 | 253/28832 | 0.04743083 | 2.634924261 | 3.536347809 | 0.002265764 | 0.025987894 | 0.019430669 |
| GO:0042472 | inner ear morphogenesis | 8/519 | 129/28832 | 0.062015504 | 3.44514645 | 3.768397206 | 0.002383883 | 0.027058625 | 0.020231235 |
| GO:0051180 | vitamin transport | 5/519 | 52/28832 | 0.096153846 | 5.341633319 | 4.242579664 | 0.002401267 | 0.027185334 | 0.020325973 |
| GO:0048771 | tissue remodeling | 11/519 | 223/28832 | 0.049327354 | 2.740281846 | 3.532167335 | 0.002517408 | 0.028207884 | 0.021090515 |
| GO:0070849 | response to epidermal growth factor | 5/519 | 53/28832 | 0.094339623 | 5.240847784 | 4.183823524 | 0.002612806 | 0.028979602 | 0.021667514 |
| GO:0016052 | carbohydrate catabolic process | 10/519 | 192/28832 | 0.052083333 | 2.893384714 | 3.563884015 | 0.002636094 | 0.029090235 | 0.021750232 |
| GO:0051782 | negative regulation of cell division | 3/519 | 16/28832 | 0.1875 | 10.41618497 | 5.100804896 | 0.002727118 | 0.029868436 | 0.022332079 |
| GO:1902036 | regulation of hematopoietic stem cell differentiation | 3/519 | 16/28832 | 0.1875 | 10.41618497 | 5.100804896 | 0.002727118 | 0.029868436 | 0.022332079 |
| GO:0150146 | cell junction disassembly | 4/519 | 33/28832 | 0.121212121 | 6.733695335 | 4.46193733 | 0.002807922 | 0.030297823 | 0.022653124 |
| GO:0038145 | macrophage colony-stimulating factor signaling pathway | 5/519 | 54/28832 | 0.092592593 | 5.143795047 | 4.126534024 | 0.002837527 | 0.030346787 | 0.022689733 |
| GO:0045124 | regulation of bone resorption | 5/519 | 54/28832 | 0.092592593 | 5.143795047 | 4.126534024 | 0.002837527 | 0.030346787 | 0.022689733 |
| GO:0098727 | maintenance of cell number | 11/519 | 227/28832 | 0.04845815 | 2.691994941 | 3.465066711 | 0.002885685 | 0.03075718 | 0.022996576 |
| GO:0007178 | cell surface receptor protein serine/threonine kinase signaling pathway | 16/519 | 404/28832 | 0.03960396 | 2.200118278 | 3.288987075 | 0.00293495 | 0.031172755 | 0.023307294 |
| GO:0150115 | cell-substrate junction organization | 7/519 | 105/28832 | 0.066666667 | 3.703532434 | 3.757523315 | 0.002938942 | 0.031172755 | 0.023307294 |
| GO:0001824 | blastocyst development | 9/519 | 164/28832 | 0.054878049 | 3.048639504 | 3.56212665 | 0.003000184 | 0.031745288 | 0.023735366 |
| GO:0002752 | cell surface pattern recognition receptor signaling pathway | 6/519 | 79/28832 | 0.075949367 | 4.219214166 | 3.879202232 | 0.003031343 | 0.031843675 | 0.023808929 |
| GO:0016485 | protein processing | 12/519 | 263/28832 | 0.045627376 | 2.534737027 | 3.385204529 | 0.003106816 | 0.032402828 | 0.024226997 |
| GO:0141091 | transforming growth factor beta receptor superfamily signaling pathway | 15/519 | 370/28832 | 0.040540541 | 2.252148102 | 3.282044324 | 0.00314921 | 0.032688944 | 0.024440921 |
| GO:0034612 | response to tumor necrosis factor | 11/519 | 230/28832 | 0.047826087 | 2.656881964 | 3.415685489 | 0.003189413 | 0.032949724 | 0.024635901 |
| GO:0141176 | gene silencing by piRNA-directed DNA methylation | 3/519 | 17/28832 | 0.176470588 | 9.803468208 | 4.915747154 | 0.003267565 | 0.033362756 | 0.024944717 |
| GO:0018212 | peptidyl-tyrosine modification | 13/519 | 300/28832 | 0.043333333 | 2.407296082 | 3.317420972 | 0.003316013 | 0.03362176 | 0.02513837 |
| GO:0033674 | positive regulation of kinase activity | 13/519 | 300/28832 | 0.043333333 | 2.407296082 | 3.317420972 | 0.003316013 | 0.03362176 | 0.02513837 |
| GO:0044772 | mitotic cell cycle phase transition | 17/519 | 447/28832 | 0.03803132 | 2.112753402 | 3.210175904 | 0.003333854 | 0.033646521 | 0.025156883 |
| GO:0030198 | extracellular matrix organization | 14/519 | 337/28832 | 0.041543027 | 2.307839202 | 3.269682316 | 0.003434531 | 0.034550767 | 0.025832972 |
| GO:1902932 | positive regulation of alcohol biosynthetic process | 4/519 | 35/28832 | 0.114285714 | 6.348912744 | 4.286931064 | 0.003492771 | 0.034927712 | 0.026114806 |
| GO:0044706 | multi-multicellular organism process | 12/519 | 267/28832 | 0.04494382 | 2.496763439 | 3.326689521 | 0.003507088 | 0.034990811 | 0.026161984 |
| GO:0002696 | positive regulation of leukocyte activation | 15/519 | 377/28832 | 0.039787798 | 2.210331028 | 3.202699078 | 0.00374562 | 0.036865671 | 0.027563783 |
| GO:0061430 | bone trabecula morphogenesis | 3/519 | 18/28832 | 0.166666667 | 9.258831085 | 4.745408981 | 0.003869103 | 0.037487678 | 0.028028846 |
| GO:0140966 | piRNA-mediated heterochromatin formation | 3/519 | 18/28832 | 0.166666667 | 9.258831085 | 4.745408981 | 0.003869103 | 0.037487678 | 0.028028846 |
| GO:1903830 | magnesium ion transmembrane transport | 3/519 | 18/28832 | 0.166666667 | 9.258831085 | 4.745408981 | 0.003869103 | 0.037487678 | 0.028028846 |
| GO:0036037 | CD8-positive, alpha-beta T cell activation | 4/519 | 37/28832 | 0.108108108 | 6.005728272 | 4.125059137 | 0.004282995 | 0.04007856 | 0.029966 |
| GO:0043304 | regulation of mast cell degranulation | 4/519 | 37/28832 | 0.108108108 | 6.005728272 | 4.125059137 | 0.004282995 | 0.04007856 | 0.029966 |

| | | | | | | | | | |
|---|---|---|---|---|---|---|---|---|---|
| GO:0010639 | negative regulation of organelle organization | 15/519 | 384/28832 | 0.0390625 | 2.170038536 | 3.125075367 | 0.004431489 | 0.040934577 | 0.030606028 |
| GO:0030865 | cortical cytoskeleton organization | 5/519 | 60/28832 | 0.083333333 | 4.629415543 | 3.810200956 | 0.004488453 | 0.040934577 | 0.030606028 |
| GO:0031529 | ruffle organization | 5/519 | 60/28832 | 0.083333333 | 4.629415543 | 3.810200956 | 0.004488453 | 0.040934577 | 0.030606028 |
| GO:0001821 | histamine secretion | 3/519 | 19/28832 | 0.157894737 | 8.771524186 | 4.587851491 | 0.004533715 | 0.040934577 | 0.030606028 |
| GO:0016137 | glycoside metabolic process | 3/519 | 19/28832 | 0.157894737 | 8.771524186 | 4.587851491 | 0.004533715 | 0.040934577 | 0.030606028 |
| GO:0080154 | regulation of fertilization | 3/519 | 19/28832 | 0.157894737 | 8.771524186 | 4.587851491 | 0.004533715 | 0.040934577 | 0.030606028 |
| GO:2000047 | regulation of cell-cell adhesion mediated by cadherin | 3/519 | 19/28832 | 0.157894737 | 8.771524186 | 4.587851491 | 0.004533715 | 0.040934577 | 0.030606028 |
| GO:2000725 | regulation of cardiac muscle cell differentiation | 3/519 | 19/28832 | 0.157894737 | 8.771524186 | 4.587851491 | 0.004533715 | 0.040934577 | 0.030606028 |
| GO:0021700 | developmental maturation | 16/519 | 423/28832 | 0.037825059 | 2.101294998 | 3.089345385 | 0.004562109 | 0.041021432 | 0.030670968 |
| GO:0015698 | inorganic anion transport | 9/519 | 175/28832 | 0.051428571 | 2.857010735 | 3.336096959 | 0.004597485 | 0.041090899 | 0.030722908 |
| GO:0007422 | peripheral nervous system development | 6/519 | 86/28832 | 0.069767442 | 3.875789757 | 3.616081707 | 0.004616849 | 0.041090899 | 0.030722908 |
| GO:1904888 | cranial skeletal system development | 6/519 | 86/28832 | 0.069767442 | 3.875789757 | 3.616081707 | 0.004616849 | 0.041090899 | 0.030722908 |
| GO:0048041 | focal adhesion assembly | 6/519 | 87/28832 | 0.068965517 | 3.831240449 | 3.580764932 | 0.004885216 | 0.042696786 | 0.0319236 |
| GO:1901264 | carbohydrate derivative transport | 6/519 | 87/28832 | 0.068965517 | 3.831240449 | 3.580764932 | 0.004885216 | 0.042696786 | 0.0319236 |
| GO:0006790 | sulfur compound metabolic process | 14/519 | 351/28832 | 0.03988604 | 2.215788636 | 3.102805899 | 0.004901749 | 0.042755777 | 0.031967706 |
| GO:0021846 | cell proliferation in forebrain | 4/519 | 39/28832 | 0.102564103 | 5.697742206 | 3.974647452 | 0.005185636 | 0.044433782 | 0.033222319 |
| GO:0072337 | modified amino acid transport | 4/519 | 39/28832 | 0.102564103 | 5.697742206 | 3.974647452 | 0.005185636 | 0.044433782 | 0.033222319 |
| GO:0006911 | phagocytosis, engulfment | 5/519 | 63/28832 | 0.079365079 | 4.408967184 | 3.667341049 | 0.005529019 | 0.046644426 | 0.034875177 |
| GO:0036005 | response to macrophage colony-stimulating factor | 5/519 | 63/28832 | 0.079365079 | 4.408967184 | 3.667341049 | 0.005529019 | 0.046644426 | 0.034875177 |
| GO:0072089 | stem cell proliferation | 8/519 | 149/28832 | 0.053691275 | 2.982710685 | 3.285192195 | 0.005714937 | 0.047570043 | 0.035567244 |
| GO:0034142 | toll-like receptor 4 signaling pathway | 5/519 | 64/28832 | 0.078125 | 4.340077071 | 3.621697953 | 0.005910748 | 0.04855257 | 0.036301861 |
| GO:0051893 | regulation of focal adhesion assembly | 5/519 | 64/28832 | 0.078125 | 4.340077071 | 3.621697953 | 0.005910748 | 0.04855257 | 0.036301861 |
| GO:0006887 | exocytosis | 15/519 | 397/28832 | 0.037783375 | 2.098979339 | 2.985236564 | 0.005975435 | 0.048808695 | 0.036493361 |
| GO:0045836 | positive regulation of meiotic nuclear division | 3/519 | 21/28832 | 0.142857143 | 7.93614093 | 4.30495858 | 0.006059252 | 0.04912603 | 0.036730627 |
| GO:1901569 | fatty acid derivative catabolic process | 3/519 | 21/28832 | 0.142857143 | 7.93614093 | 4.30495858 | 0.006059252 | 0.04912603 | 0.036730627 |

**Supplementary Table 13:** Summary of GO process terms associated with DEGs exhibiting >2 fold downregulation in marked *Tead4* KD versus unmarked outer control cell clones at the E3.5 stage (RNA-Seq data). Relates to data in Fig. S9.

**Supplementary Table 14:**

| ID | Description | GeneRatio | BgRatio | RichFactor | FoldEnrichment | zScore | pvalue | p.adjust | qvalue |
|---|---|---|---|---|---|---|---|---|---|
| GO:0046943 | carboxylic acid transmembrane transporter activity | 18/539 | 199/28366 | 0.090452261 | 4.760239043 | 7.40839753 | 5.58E-08 | 2.56E-05 | 1.83E-05 |
| GO:0008514 | organic anion transmembrane transporter activity | 20/539 | 279/28366 | 0.071684588 | 3.772551054 | 6.47711427 | 4.69E-07 | 0.000132974 | 9.52E-05 |
| GO:0061629 | RNA polymerase II-specific DNA-binding transcription factor binding | 23/539 | 387/28366 | 0.059431525 | 3.127708025 | 5.865485999 | 1.75E-06 | 0.000372658 | 0.000266745 |
| GO:0050839 | cell adhesion molecule binding | 19/539 | 313/28366 | 0.060702875 | 3.194615517 | 5.433673526 | 1.02E-05 | 0.001740224 | 0.001245634 |
| GO:0042578 | phosphoric ester hydrolase activity | 20/539 | 356/28366 | 0.056179775 | 2.956577933 | 5.170328987 | 1.86E-05 | 0.00263916 | 0.001889083 |
| GO:1901681 | sulfur compound binding | 18/539 | 303/28366 | 0.059405941 | 3.126361616 | 5.178976884 | 2.32E-05 | 0.002738493 | 0.001960185 |
| GO:0015293 | symporter activity | 12/539 | 147/28366 | 0.081632653 | 4.296088751 | 5.576212777 | 2.58E-05 | 0.002738493 | 0.001960185 |
| GO:0005319 | lipid transporter activity | 13/539 | 176/28366 | 0.073863636 | 3.887228032 | 5.347397443 | 3.46E-05 | 0.003268199 | 0.002339342 |
| GO:0030246 | carbohydrate binding | 17/539 | 293/28366 | 0.058020478 | 3.053448745 | 4.917290286 | 5.22E-05 | 0.004435925 | 0.003175189 |
| GO:0043425 | bHLH transcription factor binding | 6/539 | 37/28366 | 0.162162162 | 8.534122248 | 6.38220111 | 6.45E-05 | 0.004966166 | 0.003554729 |
| GO:0016757 | glycosyltransferase activity | 16/539 | 271/28366 | 0.05904059 | 3.1071343 | 4.850817395 | 7.01E-05 | 0.004966166 | 0.003554729 |
| GO:0051015 | actin filament binding | 14/539 | 221/28366 | 0.063348416 | 3.333842628 | 4.847520573 | 9.36E-05 | 0.005758132 | 0.00412161 |
| GO:0015081 | sodium ion transmembrane transporter activity | 12/539 | 168/28366 | 0.071428571 | 3.759077657 | 4.991855377 | 9.48E-05 | 0.005758132 | 0.00412161 |
| GO:0005504 | fatty acid binding | 7/539 | 63/28366 | 0.111111111 | 5.847454133 | 5.360685515 | 0.000189696 | 0.008270081 | 0.005919637 |
| GO:0170055 | lipid transmembrane transporter activity | 7/539 | 63/28366 | 0.111111111 | 5.847454133 | 5.360685515 | 0.000189696 | 0.008270081 | 0.005919637 |
| GO:0005178 | integrin binding | 11/539 | 155/28366 | 0.070967742 | 3.734825543 | 4.751588458 | 0.00019459 | 0.008270081 | 0.005919637 |
| GO:0005539 | glycosaminoglycan binding | 14/539 | 246/28366 | 0.056910569 | 2.995037483 | 4.373846501 | 0.00028528 | 0.01063894 | 0.007615241 |
| GO:0042277 | peptide binding | 16/539 | 307/28366 | 0.052117264 | 2.742779789 | 4.272953734 | 0.000292189 | 0.01063894 | 0.007615241 |
| GO:0004857 | enzyme inhibitor activity | 20/539 | 439/28366 | 0.045558087 | 2.397589394 | 4.107260952 | 0.000321594 | 0.01063894 | 0.007615241 |
| GO:0001217 | DNA-binding transcription repressor activity | 16/539 | 310/28366 | 0.051612903 | 2.716236759 | 4.228611169 | 0.000325426 | 0.01063894 | 0.007615241 |
| GO:0005006 | epidermal growth factor receptor activity | 6/539 | 51/28366 | 0.117647059 | 6.191422023 | 5.164350654 | 0.000399259 | 0.012120363 | 0.008675629 |
| GO:0005030 | neurotrophin receptor activity | 6/539 | 51/28366 | 0.117647059 | 6.191422023 | 5.164350654 | 0.000399259 | 0.012120363 | 0.008675629 |
| GO:0016829 | lyase activity | 12/539 | 199/28366 | 0.060301508 | 3.173492695 | 4.282200608 | 0.0004551 | 0.013339129 | 0.009548008 |
| GO:0019838 | growth factor binding | 10/539 | 148/28366 | 0.067567568 | 3.55588427 | 4.338718872 | 0.000554693 | 0.014626203 | 0.010469282 |
| GO:0005518 | collagen binding | 7/539 | 76/28366 | 0.092105263 | 4.847231716 | 4.674027349 | 0.000606571 | 0.015164264 | 0.010854421 |
| GO:0043177 | organic acid binding | 12/539 | 210/28366 | 0.057142857 | 3.007262126 | 4.063318278 | 0.000733243 | 0.016867814 | 0.012073804 |
| GO:0015370 | solute:sodium symporter activity | 7/539 | 82/28366 | 0.085365854 | 4.492556224 | 4.407907789 | 0.00095691 | 0.020334332 | 0.014555101 |
| GO:0001730 | 2'-5'-oligoadenylate synthetase activity | 3/539 | 11/28366 | 0.272727273 | 14.35284196 | 6.164641254 | 0.001004729 | 0.020829748 | 0.014909714 |
| GO:0005109 | frizzled binding | 5/539 | 42/28366 | 0.119047619 | 6.265129428 | 4.752356221 | 0.001157983 | 0.022352729 | 0.015999848 |
| GO:0035251 | UDP-glucosyltransferase activity | 3/539 | 12/28366 | 0.25 | 13.1567718 | 5.862114575 | 0.001320794 | 0.022846073 | 0.016352979 |
| GO:0051087 | protein-folding chaperone binding | 10/539 | 166/28366 | 0.060240964 | 3.170306458 | 3.903041194 | 0.001338262 | 0.022846073 | 0.016352979 |
| GO:0005104 | fibroblast growth factor receptor binding | 4/539 | 27/28366 | 0.148148148 | 7.796605511 | 4.917385625 | 0.001599015 | 0.024602274 | 0.017610049 |
| GO:0005201 | extracellular matrix structural constituent | 9/539 | 142/28366 | 0.063380282 | 3.335519611 | 3.883032195 | 0.001620856 | 0.024602274 | 0.017610049 |
| GO:0019842 | vitamin binding | 9/539 | 142/28366 | 0.063380282 | 3.335519611 | 3.883032195 | 0.001620856 | 0.024602274 | 0.017610049 |
| GO:0052650 | all-trans-retinol dehydrogenase (NADP+) activity | 3/539 | 13/28366 | 0.230769231 | 12.14471243 | 5.593628463 | 0.001692903 | 0.024809789 | 0.017758585 |
| GO:0140326 | ATPase-coupled intramembrane lipid transporter activity | 4/539 | 28/28366 | 0.142857143 | 7.518155314 | 4.802547896 | 0.001837775 | 0.026020232 | 0.018625008 |
| GO:0004602 | glutathione peroxidase activity | 4/539 | 29/28366 | 0.137931034 | 7.258908579 | 4.693245306 | 0.002100139 | 0.026031935 | 0.018633385 |
| GO:0140359 | ABC-type transporter activity | 5/539 | 48/28366 | 0.104166667 | 5.48198825 | 4.325264056 | 0.002123538 | 0.026031935 | 0.018633385 |
| GO:0001221 | transcription coregulator binding | 9/539 | 150/28366 | 0.06 | 3.157625232 | 3.687453063 | 0.002358187 | 0.026031935 | 0.018633385 |
| GO:0016500 | protein-hormone receptor activity | 6/539 | 72/28366 | 0.083333333 | 4.3855906 | 4.003189016 | 0.002478755 | 0.026181983 | 0.018740788 |
| GO:0004467 | long-chain fatty acid-CoA ligase activity | 3/539 | 15/28366 | 0.2 | 10.52541744 | 5.13568098 | 0.002618198 | 0.026181983 | 0.018740788 |
| GO:0008970 | phospholipase A1 activity | 3/539 | 15/28366 | 0.2 | 10.52541744 | 5.13568098 | 0.002618198 | 0.026181983 | 0.018740788 |
| GO:0016717 | oxidoreductase activity, acting on paired donors, with oxidation of a pair of donors resulting in the reduction of molecular oxygen to two molecules of water | 3/539 | 15/28366 | 0.2 | 10.52541744 | 5.13568098 | 0.002618198 | 0.026181983 | 0.018740788 |
| GO:0140789 | histone phosphatase activity | 8/539 | 126/28366 | 0.063492063 | 3.341402362 | 3.665907994 | 0.002866328 | 0.027686118 | 0.019817432 |
| GO:0022834 | ligand-gated channel activity | 10/539 | 185/28366 | 0.054054054 | 2.844707416 | 3.503383986 | 0.00296507 | 0.028318087 | 0.020269789 |
| GO:0038064 | collagen receptor activity | 5/539 | 53/28366 | 0.094339623 | 4.964819547 | 4.02087895 | 0.003293731 | 0.030431213 | 0.021782342 |
| GO:0004364 | glutathione transferase activity | 4/539 | 33/28366 | 0.121212121 | 6.379040873 | 4.302967996 | 0.003408289 | 0.031151028 | 0.022297578 |
| GO:0004303 | estradiol 17-beta-dehydrogenase [NAD(P)+] activity | 3/539 | 17/28366 | 0.176470588 | 9.287133035 | 4.756773605 | 0.003804076 | 0.033632754 | 0.024073971 |
| GO:0030695 | GTPase regulator activity | 17/539 | 430/28366 | 0.039534884 | 2.080605773 | 3.142476095 | 0.003877659 | 0.033632754 | 0.024073971 |
| GO:0016887 | ATP hydrolysis activity | 16/539 | 396/28366 | 0.04040404 | 2.126346958 | 3.141420927 | 0.00406793 | 0.034926676 | 0.025000147 |
| GO:0046978 | TAP1 binding | 3/539 | 18/28366 | 0.166666667 | 8.7711812 | 4.590020852 | 0.004501053 | 0.036572922 | 0.026178513 |
| GO:0046979 | TAP2 binding | 3/539 | 18/28366 | 0.166666667 | 8.7711812 | 4.590020852 | 0.004501053 | 0.036572922 | 0.026178513 |
| GO:0008374 | O-acyltransferase activity | 5/539 | 57/28366 | 0.087719298 | 4.616411158 | 3.803693783 | 0.004517832 | 0.036572922 | 0.026178513 |
| GO:1990841 | promoter-specific chromatin binding | 6/539 | 83/28366 | 0.072289157 | 3.804367749 | 3.560933439 | 0.005031136 | 0.039966971 | 0.028607937 |
| GO:0005261 | monoatomic cation channel activity | 14/539 | 334/28366 | 0.041916168 | 2.205925811 | 3.085456048 | 0.005080745 | 0.039987347 | 0.028622522 |
| GO:0140793 | histone H2AXY142 phosphatase activity | 6/539 | 86/28366 | 0.069767442 | 3.671657246 | 3.453367888 | 0.005976996 | 0.044908404 | 0.032144963 |
| GO:0043295 | glutathione binding | 3/539 | 20/28366 | 0.15 | 7.89406308 | 4.292367934 | 0.006113845 | 0.045189286 | 0.032346015 |
| GO:0019200 | carbohydrate kinase activity | 3/539 | 21/28366 | 0.142857143 | 7.518155314 | 4.158614886 | 0.007033365 | 0.049002954 | 0.035075799 |
| GO:0016616 | oxidoreductase activity, acting on the CH-OH group of donors, NAD or NADP as acceptor | 8/539 | 147/28366 | 0.054421769 | 2.864059167 | 3.153552993 | 0.007220391 | 0.049897014 | 0.035715757 |

**Supplementary Table 14:** Summary of GO function terms associated with DEGs exhibiting >2 fold upregulation in marked *Tead4* KD versus unmarked outer control cell clones at the E3.5 stage (RNA-Seq data). Relates to data in Fig. S10.

**Supplementary Table 15:**

| ID | Description | GeneRatio | BgRatio | RichFactor | FoldEnrichment | zScore | pvalue | p.adjust | qvalue |
|---|---|---|---|---|---|---|---|---|---|
| GO:0060070 | canonical Wnt signaling pathway | 29/533 | 319/28832 | 0.090909091 | 4.917618966 | 9.656149722 | 2.23E-12 | 1.04E-08 | 6.63E-09 |
| GO:0050767 | regulation of neurogenesis | 34/533 | 473/28832 | 0.071881607 | 3.88834988 | 8.692462533 | 2.06E-11 | 2.44E-08 | 1.56E-08 |
| GO:0006631 | fatty acid metabolic process | 34/533 | 477/28832 | 0.071278826 | 3.855743173 | 8.631204846 | 2.58E-11 | 2.44E-08 | 1.56E-08 |
| GO:0010717 | regulation of epithelial to mesenchymal transition | 16/533 | 108/28832 | 0.148148148 | 8.013897575 | 10.02205429 | 1.58E-10 | 1.05E-07 | 6.72E-08 |
| GO:0090287 | regulation of cellular response to growth factor stimulus | 26/533 | 336/28832 | 0.077380952 | 4.18583043 | 8.061358899 | 1.04E-09 | 5.31E-07 | 3.40E-07 |
| GO:0030198 | extracellular matrix organization | 26/533 | 337/28832 | 0.077151335 | 4.173409568 | 8.04201094 | 1.11E-09 | 5.31E-07 | 3.40E-07 |
| GO:0045859 | regulation of protein kinase activity | 31/533 | 477/28832 | 0.064989518 | 3.51553054 | 7.602945388 | 1.87E-09 | 7.21E-07 | 4.62E-07 |
| GO:0048562 | embryonic organ morphogenesis | 26/533 | 348/28832 | 0.074712644 | 4.041491449 | 7.834001426 | 2.19E-09 | 7.81E-07 | 5.00E-07 |
| GO:0032102 | negative regulation of response to external stimulus | 29/533 | 429/28832 | 0.067599068 | 3.656691026 | 7.608437913 | 2.58E-09 | 8.30E-07 | 5.32E-07 |
| GO:0043086 | negative regulation of catalytic activity | 27/533 | 377/28832 | 0.071618037 | 3.874092395 | 7.709035022 | 2.69E-09 | 8.30E-07 | 5.32E-07 |
| GO:0072001 | renal system development | 27/533 | 392/28832 | 0.068877551 | 3.725849064 | 7.457408867 | 6.18E-09 | 1.60E-06 | 1.03E-06 |
| GO:0010721 | negative regulation of cell development | 24/533 | 315/28832 | 0.076190476 | 4.121433039 | 7.644785334 | 6.23E-09 | 1.60E-06 | 1.03E-06 |
| GO:0042060 | wound healing | 27/533 | 403/28832 | 0.066997519 | 3.62415095 | 7.280621568 | 1.11E-08 | 2.42E-06 | 1.55E-06 |
| GO:0060562 | epithelial tube morphogenesis | 27/533 | 403/28832 | 0.066997519 | 3.62415095 | 7.280621568 | 1.11E-08 | 2.42E-06 | 1.55E-06 |
| GO:0018212 | peptidyl-tyrosine modification | 23/533 | 300/28832 | 0.076666667 | 4.147191995 | 7.520135581 | 1.15E-08 | 2.42E-06 | 1.55E-06 |
| GO:0050678 | regulation of epithelial cell proliferation | 28/533 | 434/28832 | 0.064516129 | 3.489923137 | 7.172901073 | 1.34E-08 | 2.55E-06 | 1.63E-06 |
| GO:0015849 | organic acid transport | 26/533 | 381/28832 | 0.06824147 | 3.691441009 | 7.257825101 | 1.43E-08 | 2.55E-06 | 1.63E-06 |
| GO:0045936 | negative regulation of phosphate metabolic process | 26/533 | 381/28832 | 0.06824147 | 3.691441009 | 7.257825101 | 1.43E-08 | 2.55E-06 | 1.63E-06 |
| GO:0070372 | regulation of ERK1 and ERK2 cascade | 24/533 | 333/28832 | 0.072072072 | 3.898652874 | 7.301488894 | 1.83E-08 | 3.13E-06 | 2.01E-06 |
| GO:0045165 | cell fate commitment | 23/533 | 310/28832 | 0.074193548 | 4.013411608 | 7.320777862 | 2.12E-08 | 3.48E-06 | 2.23E-06 |
| GO:0048608 | reproductive structure development | 26/533 | 389/28832 | 0.066838046 | 3.615524484 | 7.12777206 | 2.18E-08 | 3.48E-06 | 2.23E-06 |
| GO:0070371 | ERK1 and ERK2 cascade | 25/533 | 364/28832 | 0.068681319 | 3.715234109 | 7.154633751 | 2.39E-08 | 3.58E-06 | 2.29E-06 |
| GO:0015711 | organic anion transport | 29/533 | 477/28832 | 0.060796646 | 3.288722118 | 6.917439083 | 2.72E-08 | 3.94E-06 | 2.52E-06 |
| GO:0051962 | positive regulation of nervous system development | 25/533 | 384/28832 | 0.065104167 | 3.521732333 | 6.827263146 | 6.77E-08 | 8.39E-06 | 5.37E-06 |
| GO:0042063 | gliogenesis | 26/533 | 412/28832 | 0.063106796 | 3.413686952 | 6.772132193 | 6.87E-08 | 8.39E-06 | 5.37E-06 |
| GO:0043603 | amide metabolic process | 28/533 | 479/28832 | 0.058455115 | 3.162059795 | 6.548530605 | 1.07E-07 | 1.21E-05 | 7.75E-06 |
| GO:0007517 | muscle organ development | 25/533 | 395/28832 | 0.063291139 | 3.423658774 | 6.656348433 | 1.16E-07 | 1.24E-05 | 7.91E-06 |
| GO:0003002 | regionalization | 27/533 | 452/28832 | 0.059734513 | 3.23126733 | 6.561793234 | 1.17E-07 | 1.24E-05 | 7.91E-06 |
| GO:0007626 | locomotory behavior | 21/533 | 292/28832 | 0.071917808 | 3.890308155 | 6.812656666 | 1.47E-07 | 1.45E-05 | 9.28E-06 |
| GO:0007548 | sex differentiation | 24/533 | 375/28832 | 0.064 | 3.462003752 | 6.585938947 | 1.70E-07 | 1.64E-05 | 1.05E-05 |
| GO:0048705 | skeletal system morphogenesis | 20/533 | 270/28832 | 0.074074074 | 4.006948787 | 6.812735361 | 1.81E-07 | 1.71E-05 | 1.10E-05 |
| GO:0031346 | positive regulation of cell projection organization | 27/533 | 465/28832 | 0.058064516 | 3.140930824 | 6.387491276 | 2.06E-07 | 1.91E-05 | 1.23E-05 |
| GO:0030336 | negative regulation of cell migration | 22/533 | 328/28832 | 0.067073171 | 3.628243262 | 6.569880797 | 2.49E-07 | 2.27E-05 | 1.45E-05 |
| GO:0019216 | regulation of lipid metabolic process | 24/533 | 389/28832 | 0.061696658 | 3.337407216 | 6.369852757 | 3.31E-07 | 2.80E-05 | 1.79E-05 |
| GO:0048814 | regulation of dendrite morphogenesis | 12/533 | 100/28832 | 0.12 | 6.491257036 | 7.549127438 | 3.38E-07 | 2.80E-05 | 1.79E-05 |
| GO:0007369 | gastrulation | 17/533 | 209/28832 | 0.081339713 | 4.399974865 | 6.770163969 | 4.06E-07 | 3.31E-05 | 2.12E-05 |
| GO:0023019 | signal transduction involved in regulation of gene expression | 7/533 | 27/28832 | 0.259259259 | 14.02432076 | 9.292037173 | 4.57E-07 | 3.65E-05 | 2.34E-05 |
| GO:0048638 | regulation of developmental growth | 24/533 | 398/28832 | 0.060301508 | 3.261938209 | 6.236073227 | 5.00E-07 | 3.86E-05 | 2.48E-05 |
| GO:0007159 | leukocyte cell-cell adhesion | 25/533 | 427/28832 | 0.058548009 | 3.167084814 | 6.191563945 | 5.00E-07 | 3.86E-05 | 2.48E-05 |
| GO:0015908 | fatty acid transport | 13/533 | 124/28832 | 0.10483871 | 5.671125098 | 7.153832793 | 5.39E-07 | 4.10E-05 | 2.63E-05 |
| GO:0050890 | cognition | 24/533 | 403/28832 | 0.05955335 | 3.221467511 | 6.163389447 | 6.25E-07 | 4.52E-05 | 2.90E-05 |
| GO:0031589 | cell-substrate adhesion | 23/533 | 376/28832 | 0.061170213 | 3.308929783 | 6.184799643 | 6.74E-07 | 4.81E-05 | 3.08E-05 |
| GO:0007162 | negative regulation of cell adhesion | 21/533 | 322/28832 | 0.065217391 | 3.527857085 | 6.260222327 | 7.37E-07 | 5.10E-05 | 3.27E-05 |
| GO:0044248 | cellular catabolic process | 21/533 | 323/28832 | 0.06501548 | 3.516934926 | 6.242954497 | 7.75E-07 | 5.28E-05 | 3.38E-05 |
| GO:0031644 | regulation of nervous system process | 15/533 | 173/28832 | 0.086705202 | 4.690214621 | 6.681163096 | 8.68E-07 | 5.67E-05 | 3.63E-05 |
| GO:0051961 | negative regulation of nervous system development | 15/533 | 173/28832 | 0.086705202 | 4.690214621 | 6.681163096 | 8.68E-07 | 5.67E-05 | 3.63E-05 |
| GO:0040013 | negative regulation of locomotion | 22/533 | 362/28832 | 0.060773481 | 3.287469033 | 6.010681117 | 1.31E-06 | 8.09E-05 | 5.19E-05 |
| GO:0022409 | positive regulation of cell-cell adhesion | 21/533 | 334/28832 | 0.062874251 | 3.401107728 | 6.057390902 | 1.33E-06 | 8.09E-05 | 5.19E-05 |
| GO:0061053 | somite development | 11/533 | 94/28832 | 0.117021277 | 6.330126542 | 7.103642375 | 1.35E-06 | 8.10E-05 | 5.19E-05 |
| GO:0016053 | organic acid biosynthetic process | 21/533 | 335/28832 | 0.062686567 | 3.390955168 | 6.040907395 | 1.39E-06 | 8.27E-05 | 5.30E-05 |
| GO:0044282 | small molecule catabolic process | 22/533 | 369/28832 | 0.059620596 | 3.225105122 | 5.903795328 | 1.79E-06 | 0.000100962 | 6.47E-05 |
| GO:0017145 | stem cell division | 8/533 | 46/28832 | 0.173913043 | 9.407618892 | 7.831937006 | 1.82E-06 | 0.000100962 | 6.47E-05 |
| GO:0048010 | vascular endothelial growth factor receptor signaling pathway | 11/533 | 97/28832 | 0.113402062 | 6.134349432 | 6.951421165 | 1.85E-06 | 0.000100962 | 6.47E-05 |
| GO:0030324 | lung development | 18/533 | 259/28832 | 0.069498069 | 3.759415272 | 6.122050926 | 1.85E-06 | 0.000100962 | 6.47E-05 |
| GO:0062012 | regulation of small molecule metabolic process | 23/533 | 401/28832 | 0.057356608 | 3.102637403 | 5.819000804 | 2.02E-06 | 0.000108917 | 6.98E-05 |
| GO:0031099 | regeneration | 17/533 | 236/28832 | 0.072033898 | 3.89658791 | 6.131941231 | 2.19E-06 | 0.000115601 | 7.41E-05 |
| GO:0016042 | lipid catabolic process | 21/533 | 347/28832 | 0.060518732 | 3.273688707 | 5.847840829 | 2.43E-06 | 0.000123777 | 7.93E-05 |
| GO:0061384 | heart trabecula morphogenesis | 7/533 | 36/28832 | 0.194444444 | 10.51824057 | 7.842413875 | 3.72E-06 | 0.000174193 | 0.000111599 |
| GO:0010876 | lipid localization | 25/533 | 485/28832 | 0.051546392 | 2.788340651 | 5.450991177 | 4.89E-06 | 0.000211909 | 0.000135762 |
| GO:0001763 | morphogenesis of a branching structure | 17/533 | 252/28832 | 0.067460317 | 3.649185503 | 5.796815006 | 5.31E-06 | 0.000227691 | 0.000145873 |
| GO:0050866 | negative regulation of cell activation | 16/533 | 226/28832 | 0.07079646 | 3.829650169 | 5.860927 | 5.43E-06 | 0.000229375 | 0.000146952 |
| GO:0042692 | muscle cell differentiation | 25/533 | 488/28832 | 0.051229508 | 2.771199213 | 5.415700823 | 5.44E-06 | 0.000229375 | 0.000146952 |
| GO:0045926 | negative regulation of growth | 17/533 | 253/28832 | 0.067193676 | 3.634761845 | 5.776782614 | 5.59E-06 | 0.000230846 | 0.000147895 |
| GO:0008543 | fibroblast growth factor receptor signaling pathway | 12/533 | 130/28832 | 0.092307692 | 4.993274643 | 6.262571388 | 5.63E-06 | 0.000230846 | 0.000147895 |
| GO:0042632 | cholesterol homeostasis | 11/533 | 109/28832 | 0.100917431 | 5.45900821 | 6.400951475 | 5.83E-06 | 0.00023715 | 0.000151933 |
| GO:0003229 | ventricular cardiac muscle tissue development | 9/533 | 71/28832 | 0.126760563 | 6.856961657 | 6.781214426 | 6.35E-06 | 0.000252523 | 0.000161782 |
| GO:1902003 | regulation of amyloid-beta formation | 7/533 | 39/28832 | 0.179487179 | 9.709145139 | 7.469169 | 6.53E-06 | 0.000256075 | 0.000164058 |
| GO:0141091 | transforming growth factor beta receptor superfamily signaling pathway | 21/533 | 370/28832 | 0.056756757 | 3.070189138 | 5.500296402 | 6.57E-06 | 0.000256075 | 0.000164058 |
| GO:0045668 | negative regulation of osteoblast differentiation | 9/533 | 72/28832 | 0.125 | 6.761726079 | 6.717881329 | 7.14E-06 | 0.000271979 | 0.000174247 |
| GO:0032535 | regulation of cellular component size | 22/533 | 404/28832 | 0.054455446 | 2.94570245 | 5.405076785 | 7.68E-06 | 0.000282269 | 0.000180839 |
| GO:0006979 | response to oxidative stress | 23/533 | 437/28832 | 0.052631579 | 2.847042559 | 5.339549261 | 8.34E-06 | 0.000298367 | 0.000191153 |
| GO:0048546 | digestive tract morphogenesis | 8/533 | 56/28832 | 0.142857143 | 7.727686947 | 6.915960254 | 8.43E-06 | 0.000298367 | 0.000191153 |
| GO:0072524 | pyridine-containing compound metabolic process | 15/533 | 209/28832 | 0.071770335 | 3.882330763 | 5.73941063 | 9.01E-06 | 0.000313834 | 0.000201062 |
| GO:0046434 | organophosphate catabolic process | 17/533 | 264/28832 | 0.064393939 | 3.483313435 | 5.562903111 | 9.80E-06 | 0.000333934 | 0.000213939 |
| GO:0007015 | actin filament organization | 24/533 | 474/28832 | 0.050632911 | 2.738927019 | 5.238906412 | 1.02E-05 | 0.00034364 | 0.000220157 |
| GO:0008344 | adult locomotory behavior | 11/533 | 116/28832 | 0.094827586 | 5.129585301 | 6.11619575 | 1.06E-05 | 0.000356416 | 0.000228343 |
| GO:0043588 | skin development | 20/533 | 352/28832 | 0.056818182 | 3.073511854 | 5.37174978 | 1.07E-05 | 0.000356416 | 0.000228343 |
| GO:0001756 | somitogenesis | 9/533 | 76/28832 | 0.118421053 | 6.405845759 | 6.476108582 | 1.12E-05 | 0.000367769 | 0.000235616 |
| GO:0071774 | response to fibroblast growth factor | 13/533 | 163/28832 | 0.079754601 | 4.314230136 | 5.823419882 | 1.15E-05 | 0.000374425 | 0.00023988 |
| GO:0010038 | response to metal ion | 21/533 | 386/28832 | 0.054404145 | 2.942927413 | 5.27409002 | 1.25E-05 | 0.000392746 | 0.000251618 |
| GO:0006641 | triglyceride metabolic process | 11/533 | 119/28832 | 0.092436975 | 5.000268025 | 6.001104632 | 1.36E-05 | 0.000416834 | 0.00026705 |
| GO:0001570 | vasculogenesis | 10/533 | 99/28832 | 0.101010101 | 5.464021074 | 6.106062707 | 1.53E-05 | 0.000455241 | 0.000291656 |

| | | | | | | | | | |
|---|---|---|---|---|---|---|---|---|---|
| GO:0019827 | stem cell population maintenance | 15/533 | 219/28832 | 0.068493151 | 3.705055386 | 5.514732289 | 1.57E-05 | 0.00046396 | 0.000297242 |
| GO:0061448 | connective tissue development | 19/533 | 335/28832 | 0.056716418 | 3.068007057 | 5.224957493 | 1.81E-05 | 0.000516743 | 0.000331058 |
| GO:0046463 | acylglycerol biosynthetic process | 8/533 | 62/28832 | 0.129032258 | 6.979846275 | 6.468801824 | 1.82E-05 | 0.000516743 | 0.000331058 |
| GO:0034446 | substrate adhesion-dependent cell spreading | 10/533 | 101/28832 | 0.099009901 | 5.355822637 | 6.018155657 | 1.83E-05 | 0.000516743 | 0.000331058 |
| GO:0050777 | negative regulation of immune response | 15/533 | 222/28832 | 0.067567568 | 3.65498707 | 5.44989182 | 1.84E-05 | 0.000518275 | 0.00033204 |
| GO:0043583 | ear development | 17/533 | 279/28832 | 0.0609319 | 3.296038519 | 5.288875106 | 2.00E-05 | 0.000559135 | 0.000358217 |
| GO:0072089 | stem cell proliferation | 12/533 | 149/28832 | 0.080536913 | 4.356548346 | 5.63743656 | 2.26E-05 | 0.000608431 | 0.000389799 |
| GO:0098727 | maintenance of cell number | 15/533 | 227/28832 | 0.066079295 | 3.574480747 | 5.344283814 | 2.40E-05 | 0.000632059 | 0.000404937 |
| GO:0001666 | response to hypoxia | 19/533 | 342/28832 | 0.055555556 | 3.005211591 | 5.119587236 | 2.41E-05 | 0.000632059 | 0.000404937 |
| GO:1990845 | adaptive thermogenesis | 13/533 | 175/28832 | 0.074285714 | 4.018397213 | 5.49652195 | 2.46E-05 | 0.000640153 | 0.000410122 |
| GO:0001947 | heart looping | 8/533 | 65/28832 | 0.123076923 | 6.657699524 | 6.26696376 | 2.59E-05 | 0.000667133 | 0.000427407 |
| GO:0030856 | regulation of epithelial cell differentiation | 13/533 | 177/28832 | 0.073446328 | 3.972991594 | 5.444876343 | 2.77E-05 | 0.000709541 | 0.000454577 |
| GO:0008544 | epidermis development | 21/533 | 410/28832 | 0.051219512 | 2.770658491 | 4.955726499 | 3.04E-05 | 0.000758685 | 0.000486061 |
| GO:0034205 | amyloid-beta formation | 7/533 | 49/28832 | 0.142857143 | 7.727686947 | 6.468501733 | 3.11E-05 | 0.000763852 | 0.000489372 |
| GO:0010770 | positive regulation of cell morphogenesis | 8/533 | 67/28832 | 0.119402985 | 6.458962225 | 6.13936161 | 3.24E-05 | 0.000786827 | 0.000504091 |
| GO:0006790 | sulfur compound metabolic process | 19/533 | 351/28832 | 0.054131054 | 2.928154883 | 4.98799226 | 3.43E-05 | 0.00080876 | 0.000518142 |
| GO:0035701 | hematopoietic stem cell migration | 4/533 | 11/28832 | 0.363636364 | 19.67047587 | 8.499729801 | 3.44E-05 | 0.00080876 | 0.000518142 |
| GO:0045923 | positive regulation of fatty acid metabolic process | 7/533 | 50/28832 | 0.14 | 7.573133208 | 6.384176193 | 3.56E-05 | 0.000825174 | 0.000528658 |
| GO:0050878 | regulation of body fluid levels | 20/533 | 384/28832 | 0.052083333 | 2.817385866 | 4.920336463 | 3.71E-05 | 0.000850406 | 0.000544824 |
| GO:0032970 | regulation of actin filament-based process | 21/533 | 416/28832 | 0.050480769 | 2.73069707 | 4.879711708 | 3.76E-05 | 0.000858236 | 0.00054984 |
| GO:0031348 | negative regulation of defense response | 17/533 | 295/28832 | 0.057627119 | 3.117270328 | 5.016387472 | 4.05E-05 | 0.000903331 | 0.000578731 |
| GO:0048545 | response to steroid hormone | 19/533 | 356/28832 | 0.053370787 | 2.887029112 | 4.91668087 | 4.16E-05 | 0.000912968 | 0.000584905 |
| GO:0009261 | ribonucleotide catabolic process | 12/533 | 160/28832 | 0.075 | 4.057035647 | 5.321565516 | 4.56E-05 | 0.000979541 | 0.000627556 |
| GO:0050863 | regulation of T cell activation | 20/533 | 391/28832 | 0.051150895 | 2.766946733 | 4.827778294 | 4.77E-05 | 0.00100887 | 0.000646346 |
| GO:0061371 | determination of heart left/right asymmetry | 8/533 | 71/28832 | 0.112676056 | 6.095077029 | 5.899101252 | 4.96E-05 | 0.001034973 | 0.000663069 |
| GO:0007492 | endoderm development | 9/533 | 92/28832 | 0.097826087 | 5.291785627 | 5.658436751 | 5.26E-05 | 0.001077858 | 0.000690544 |
| GO:0150104 | transport across blood-brain barrier | 5/533 | 23/28832 | 0.217391304 | 11.75952362 | 7.084362611 | 5.41E-05 | 0.001100331 | 0.000704941 |
| GO:0001894 | tissue homeostasis | 17/533 | 303/28832 | 0.056105611 | 3.034966161 | 4.887009064 | 5.65E-05 | 0.001114918 | 0.000714287 |
| GO:0060249 | anatomical structure homeostasis | 17/533 | 303/28832 | 0.056105611 | 3.034966161 | 4.887009064 | 5.65E-05 | 0.001114918 | 0.000714287 |
| GO:0043254 | regulation of protein-containing complex assembly | 22/533 | 461/28832 | 0.047722343 | 2.581483275 | 4.697704093 | 5.71E-05 | 0.001121184 | 0.000718301 |
| GO:0008202 | steroid metabolic process | 19/533 | 366/28832 | 0.051912568 | 2.808148535 | 4.777705071 | 6.02E-05 | 0.001177364 | 0.000754294 |
| GO:0045664 | regulation of neuron differentiation | 15/533 | 247/28832 | 0.060728745 | 3.285049107 | 4.94974504 | 6.32E-05 | 0.001224835 | 0.000784706 |
| GO:0035264 | multicellular organism growth | 15/533 | 250/28832 | 0.06 | 3.245628518 | 4.894062652 | 7.24E-05 | 0.001363403 | 0.000873482 |
| GO:0035050 | embryonic heart tube development | 9/533 | 96/28832 | 0.09375 | 5.071294559 | 5.483563751 | 7.35E-05 | 0.001366408 | 0.000875407 |
| GO:1903035 | negative regulation of response to wounding | 9/533 | 96/28832 | 0.09375 | 5.071294559 | 5.483563751 | 7.35E-05 | 0.001366408 | 0.000875407 |
| GO:0034248 | regulation of amide metabolic process | 8/533 | 75/28832 | 0.106666667 | 5.770006254 | 5.676566167 | 7.37E-05 | 0.001366408 | 0.000875407 |
| GO:1905330 | regulation of morphogenesis of an epithelium | 8/533 | 75/28832 | 0.106666667 | 5.770006254 | 5.676566167 | 7.37E-05 | 0.001366408 | 0.000875407 |
| GO:0050868 | negative regulation of T cell activation | 11/533 | 143/28832 | 0.076923077 | 4.161062202 | 5.200573855 | 7.48E-05 | 0.001381229 | 0.000884902 |
| GO:0042180 | ketone metabolic process | 15/533 | 251/28832 | 0.059760956 | 3.232697727 | 4.875688978 | 7.57E-05 | 0.001386476 | 0.000888264 |
| GO:0030098 | lymphocyte differentiation | 22/533 | 473/28832 | 0.046511628 | 2.515991099 | 4.562361339 | 8.30E-05 | 0.001497065 | 0.000959114 |
| GO:0022604 | regulation of cell morphogenesis | 16/533 | 283/28832 | 0.056537102 | 3.058307202 | 4.775467283 | 8.48E-05 | 0.00150616 | 0.000964941 |
| GO:0070661 | leukocyte proliferation | 19/533 | 379/28832 | 0.050131926 | 2.711826818 | 4.603872756 | 9.53E-05 | 0.001661081 | 0.001064194 |
| GO:0006090 | pyruvate metabolic process | 11/533 | 147/28832 | 0.074829932 | 4.04783602 | 5.08429511 | 9.59E-05 | 0.001664696 | 0.001066509 |
| GO:0045671 | negative regulation of osteoclast differentiation | 6/533 | 41/28832 | 0.146341463 | 7.916167117 | 6.081867373 | 0.000100727 | 0.001735578 | 0.001111921 |
| GO:0001818 | negative regulation of cytokine production | 17/533 | 318/28832 | 0.053459119 | 2.89180738 | 4.655531994 | 0.000102014 | 0.00175124 | 0.001121955 |
| GO:0003158 | endothelium development | 11/533 | 149/28832 | 0.073825503 | 3.993502651 | 5.027689026 | 0.000108223 | 0.001825985 | 0.001169841 |
| GO:0030900 | forebrain development | 22/533 | 482/28832 | 0.045643154 | 2.469012012 | 4.463548392 | 0.000108863 | 0.001825985 | 0.001169841 |
| GO:0120162 | positive regulation of cold-induced thermogenesis | 9/533 | 101/28832 | 0.089108911 | 4.820240373 | 5.278176622 | 0.000109126 | 0.001825985 | 0.001169841 |
| GO:2000027 | regulation of animal organ morphogenesis | 9/533 | 101/28832 | 0.089108911 | 4.820240373 | 5.278176622 | 0.000109126 | 0.001825985 | 0.001169841 |
| GO:0002832 | negative regulation of response to biotic stimulus | 11/533 | 150/28832 | 0.073333333 | 3.9668793 | 4.999754791 | 0.000114877 | 0.001915304 | 0.001227064 |
| GO:0034599 | cellular response to oxidative stress | 15/533 | 263/28832 | 0.057034221 | 3.085198211 | 4.662143225 | 0.00012718 | 0.002061109 | 0.001320476 |
| GO:0034204 | lipid translocation | 7/533 | 61/28832 | 0.114754098 | 6.207486236 | 5.587576508 | 0.000130558 | 0.002101168 | 0.001346141 |
| GO:0003183 | mitral valve morphogenesis | 4/533 | 15/28832 | 0.266666667 | 14.42501563 | 7.137464471 | 0.000134059 | 0.002150044 | 0.001377454 |
| GO:0032370 | positive regulation of lipid transport | 9/533 | 104/28832 | 0.086538462 | 4.681194978 | 5.161319066 | 0.000136738 | 0.002177942 | 0.001395327 |
| GO:0045123 | cellular extravasation | 8/533 | 82/28832 | 0.097560976 | 5.277444744 | 5.32329241 | 0.000139106 | 0.00220054 | 0.001409805 |
| GO:0070664 | negative regulation of leukocyte proliferation | 9/533 | 105/28832 | 0.085714286 | 4.636612168 | 5.123354569 | 0.000147147 | 0.002291364 | 0.001467993 |
| GO:0050807 | regulation of synapse organization | 19/533 | 393/28832 | 0.048346056 | 2.6152223 | 4.424654145 | 0.000152361 | 0.002327002 | 0.001490825 |
| GO:0009636 | response to toxic substance | 14/533 | 238/28832 | 0.058823529 | 3.181988743 | 4.638861869 | 0.000153598 | 0.002327002 | 0.001490825 |
| GO:1903825 | organic acid transmembrane transport | 11/533 | 156/28832 | 0.070512821 | 3.814307019 | 4.837070309 | 0.000162581 | 0.002438711 | 0.001562393 |
| GO:0035924 | cellular response to vascular endothelial growth factor stimulus | 9/533 | 107/28832 | 0.08411215 | 4.54994652 | 5.048839834 | 0.000169956 | 0.002530363 | 0.00162111 |
| GO:0043542 | endothelial cell migration | 14/533 | 241/28832 | 0.058091286 | 3.142378925 | 4.583508582 | 0.000174959 | 0.002551536 | 0.001634675 |
| GO:1901136 | carbohydrate derivative catabolic process | 15/533 | 271/28832 | 0.055350554 | 2.994122249 | 4.526449014 | 0.000176484 | 0.002551536 | 0.001634675 |
| GO:0022602 | ovulation cycle process | 7/533 | 64/28832 | 0.109375 | 5.916510319 | 5.403809369 | 0.000177259 | 0.002551536 | 0.001634675 |
| GO:0098815 | modulation of excitatory postsynaptic potential | 7/533 | 64/28832 | 0.109375 | 5.916510319 | 5.403809369 | 0.000177259 | 0.002551536 | 0.001634675 |
| GO:0055074 | calcium ion homeostasis | 18/533 | 366/28832 | 0.049180328 | 2.660351244 | 4.387177493 | 0.000182947 | 0.002622529 | 0.001680158 |
| GO:0042593 | glucose homeostasis | 17/533 | 334/28832 | 0.050898204 | 2.753277685 | 4.423078636 | 0.000183322 | 0.002622529 | 0.001680158 |
| GO:0005996 | monosaccharide metabolic process | 16/533 | 303/28832 | 0.052805281 | 2.85643874 | 4.458272143 | 0.000186012 | 0.00265282 | 0.001699564 |
| GO:0046395 | carboxylic acid catabolic process | 14/533 | 243/28832 | 0.057613169 | 3.116515724 | 4.547084937 | 0.000190582 | 0.002693139 | 0.001725395 |
| GO:0010565 | regulation of ketone metabolic process | 11/533 | 159/28832 | 0.06918239 | 3.742338962 | 4.75872975 | 0.000192145 | 0.002706968 | 0.001734254 |
| GO:0050803 | regulation of synapse structure or activity | 19/533 | 401/28832 | 0.047381546 | 2.563048289 | 4.325700304 | 0.000196924 | 0.002757527 | 0.001766646 |
| GO:0060688 | regulation of morphogenesis of a branching structure | 7/533 | 66/28832 | 0.106060606 | 5.737222127 | 5.287664504 | 0.000215281 | 0.002952157 | 0.001891338 |
| GO:0097035 | regulation of membrane lipid distribution | 7/533 | 66/28832 | 0.106060606 | 5.737222127 | 5.287664504 | 0.000215281 | 0.002952157 | 0.001891338 |
| GO:0006874 | intracellular calcium ion homeostasis | 17/533 | 339/28832 | 0.050147493 | 2.712668869 | 4.353234645 | 0.0002183 | 0.002984723 | 0.001912202 |
| GO:0042445 | hormone metabolic process | 15/533 | 277/28832 | 0.054151625 | 2.929267615 | 4.427914125 | 0.000223612 | 0.003048355 | 0.001952969 |
| GO:0038084 | vascular endothelial growth factor signaling pathway | 8/533 | 88/28832 | 0.090909091 | 4.917618966 | 5.051238103 | 0.000227804 | 0.00308735 | 0.001977951 |
| GO:0008064 | regulation of actin polymerization or depolymerization | 11/533 | 163/28832 | 0.067484663 | 3.650502423 | 4.657186699 | 0.000238549 | 0.003214172 | 0.002059201 |
| GO:0006096 | glycolytic process | 9/533 | 113/28832 | 0.079646018 | 4.30835644 | 4.835871719 | 0.000256735 | 0.00338815 | 0.002170663 |
| GO:0002244 | hematopoietic progenitor cell differentiation | 12/533 | 192/28832 | 0.0625 | 3.380863039 | 4.542620261 | 0.000257309 | 0.00338815 | 0.002170663 |
| GO:0016052 | carbohydrate catabolic process | 12/533 | 192/28832 | 0.0625 | 3.380863039 | 4.542620261 | 0.000257309 | 0.00338815 | 0.002170663 |
| GO:0050905 | neuromuscular process | 13/533 | 222/28832 | 0.058558559 | 3.16765546 | 4.449546251 | 0.000273225 | 0.00357739 | 0.002291902 |
| GO:0071825 | protein-lipid complex organization | 6/533 | 49/28832 | 0.12244898 | 6.623731669 | 5.407076508 | 0.000276514 | 0.003610263 | 0.002312963 |
| GO:0042178 | xenobiotic catabolic process | 5/533 | 32/28832 | 0.15625 | 8.452157598 | 5.78853534 | 0.00028242 | 0.003663209 | 0.002346883 |
| GO:0034375 | high-density lipoprotein particle remodeling | 4/533 | 18/28832 | 0.222222222 | 12.02084636 | 6.418851637 | 0.000287579 | 0.003698445 | 0.002369458 |
| GO:0050870 | positive regulation of T cell activation | 14/533 | 253/28832 | 0.055335968 | 2.993333284 | 4.370434118 | 0.000288056 | 0.003698445 | 0.002369458 |
| GO:0098773 | skin epidermis development | 11/533 | 167/28832 | 0.065868263 | 3.563065239 | 4.558792616 | 0.000294087 | 0.003765453 | 0.002412387 |

| | | | | | | | | | |
|---|---|---|---|---|---|---|---|---|---|
| GO:0002697 | regulation of immune effector process | 21/533 | 485/28832 | 0.043298969 | 2.342206147 | 4.091141059 | 0.000314966 | 0.003956282 | 0.002534644 |
| GO:0007596 | blood coagulation | 12/533 | 197/28832 | 0.060913706 | 3.295054333 | 4.435937349 | 0.000325435 | 0.004030497 | 0.002582191 |
| GO:0034109 | homotypic cell-cell adhesion | 8/533 | 93/28832 | 0.086021505 | 4.65323085 | 4.842734698 | 0.000333098 | 0.004052258 | 0.002596133 |
| GO:0042246 | tissue regeneration | 8/533 | 93/28832 | 0.086021505 | 4.65323085 | 4.842734698 | 0.000333098 | 0.004052258 | 0.002596133 |
| GO:0050817 | coagulation | 12/533 | 200/28832 | 0.06 | 3.245628518 | 4.373558934 | 0.000373216 | 0.004446927 | 0.002848983 |
| GO:0042633 | hair cycle | 10/533 | 145/28832 | 0.068965517 | 3.730607492 | 4.523854463 | 0.000381423 | 0.00448704 | 0.002874681 |
| GO:0051099 | positive regulation of binding | 10/533 | 146/28832 | 0.068493151 | 3.705055386 | 4.497027108 | 0.000402836 | 0.004624926 | 0.00296302 |
| GO:0097006 | regulation of plasma lipoprotein particle levels | 7/533 | 73/28832 | 0.095890411 | 5.18707754 | 4.915791522 | 0.000403122 | 0.004624926 | 0.00296302 |
| GO:0006575 | modified amino acid metabolic process | 12/533 | 202/28832 | 0.059405941 | 3.213493582 | 4.332625892 | 0.00040826 | 0.0046608 | 0.002986003 |
| GO:0042552 | myelination | 12/533 | 202/28832 | 0.059405941 | 3.213493582 | 4.332625892 | 0.00040826 | 0.0046608 | 0.002986003 |
| GO:0002064 | epithelial cell development | 14/533 | 262/28832 | 0.053435115 | 2.890508858 | 4.218734117 | 0.000409637 | 0.004665029 | 0.002988712 |
| GO:0014855 | striated muscle cell proliferation | 8/533 | 96/28832 | 0.083333333 | 4.507817386 | 4.72462505 | 0.000413332 | 0.004695569 | 0.003008278 |
| GO:0043279 | response to alkaloid | 9/533 | 121/28832 | 0.074380165 | 4.023506427 | 4.57390705 | 0.000426734 | 0.004755319 | 0.003046558 |
| GO:0009743 | response to carbohydrate | 15/533 | 296/28832 | 0.050675676 | 2.741240302 | 4.132529201 | 0.000451387 | 0.004946046 | 0.003168749 |
| GO:0006091 | generation of precursor metabolites and energy | 21/533 | 500/28832 | 0.042 | 2.271939962 | 3.937503121 | 0.000469353 | 0.005082833 | 0.003256384 |
| GO:0032640 | tumor necrosis factor production | 12/533 | 206/28832 | 0.058252427 | 3.151095648 | 4.252270773 | 0.000486733 | 0.005210183 | 0.003337973 |
| GO:0045446 | endothelial cell differentiation | 9/533 | 124/28832 | 0.072580645 | 3.92616353 | 4.481422393 | 0.000510588 | 0.005427921 | 0.003477469 |
| GO:0033674 | positive regulation of kinase activity | 15/533 | 300/28832 | 0.05 | 2.704690432 | 4.073314621 | 0.000518894 | 0.005478528 | 0.003509891 |
| GO:0072526 | pyridine-containing compound catabolic process | 9/533 | 125/28832 | 0.072 | 3.894754221 | 4.451236923 | 0.00054137 | 0.005613538 | 0.003596387 |
| GO:0050867 | positive regulation of cell activation | 18/533 | 401/28832 | 0.044887781 | 2.428151011 | 3.952375179 | 0.000544945 | 0.005637991 | 0.003612053 |
| GO:0048872 | homeostasis of number of cells | 19/533 | 436/28832 | 0.043577982 | 2.357299 | 3.919206105 | 0.00055384 | 0.00567931 | 0.003638524 |
| GO:0032355 | response to estradiol | 9/533 | 126/28832 | 0.071428571 | 3.863843474 | 4.421362493 | 0.000573658 | 0.005792816 | 0.003711244 |
| GO:0046651 | lymphocyte proliferation | 16/533 | 337/28832 | 0.047477745 | 2.568252042 | 3.974242784 | 0.000600813 | 0.006014623 | 0.003853347 |
| GO:0098869 | cellular oxidant detoxification | 7/533 | 78/28832 | 0.08974359 | 4.854572569 | 4.678237939 | 0.000603942 | 0.006032917 | 0.003865067 |
| GO:1905954 | positive regulation of lipid localization | 9/533 | 127/28832 | 0.070866142 | 3.833419509 | 4.391793157 | 0.000607504 | 0.006042452 | 0.003871176 |
| GO:0050910 | detection of mechanical stimulus involved in sensory perception of sound | 4/533 | 22/28832 | 0.181818182 | 9.835237933 | 5.68939221 | 0.000648339 | 0.006314458 | 0.00404544 |
| GO:0006066 | alcohol metabolic process | 17/533 | 373/28832 | 0.045576408 | 2.465401466 | 3.909391215 | 0.000649837 | 0.006314458 | 0.00404544 |
| GO:0070252 | actin-mediated cell contraction | 8/533 | 104/28832 | 0.076923077 | 4.161062202 | 4.432052829 | 0.000706328 | 0.006792183 | 0.004351501 |
| GO:0060021 | roof of mouth development | 8/533 | 105/28832 | 0.076190476 | 4.121433039 | 4.397556704 | 0.000752446 | 0.007088596 | 0.004541402 |
| GO:1902806 | regulation of cell cycle G1/S phase transition | 11/533 | 189/28832 | 0.058201058 | 3.148316904 | 4.066566788 | 0.000831935 | 0.007693026 | 0.004928638 |
| GO:0051347 | positive regulation of transferase activity | 16/533 | 348/28832 | 0.045977011 | 2.487071661 | 3.830266074 | 0.000845212 | 0.007772929 | 0.004979829 |
| GO:0072091 | regulation of stem cell proliferation | 8/533 | 107/28832 | 0.074766355 | 4.044396907 | 4.329832094 | 0.000851945 | 0.00781934 | 0.005009563 |
| GO:0008354 | germ cell migration | 3/533 | 11/28832 | 0.272727273 | 14.7528569 | 6.260984909 | 0.000928014 | 0.008352126 | 0.005350899 |
| GO:0031642 | negative regulation of myelination | 3/533 | 11/28832 | 0.272727273 | 14.7528569 | 6.260984909 | 0.000928014 | 0.008352126 | 0.005350899 |
| GO:0048318 | axial mesoderm development | 3/533 | 11/28832 | 0.272727273 | 14.7528569 | 6.260984909 | 0.000928014 | 0.008352126 | 0.005350899 |
| GO:0048771 | tissue remodeling | 12/533 | 223/28832 | 0.053811659 | 2.910877594 | 3.931351485 | 0.000976657 | 0.008739006 | 0.005598759 |
| GO:0045913 | positive regulation of carbohydrate metabolic process | 8/533 | 110/28832 | 0.072727273 | 3.934095173 | 4.231273651 | 0.001020723 | 0.00902872 | 0.005784368 |
| GO:0060325 | face morphogenesis | 5/533 | 42/28832 | 0.119047619 | 6.439739123 | 4.841609942 | 0.001025115 | 0.009033099 | 0.005787174 |
| GO:0030316 | osteoclast differentiation | 9/533 | 137/28832 | 0.065693431 | 3.553607866 | 4.11167983 | 0.001044773 | 0.009171448 | 0.005875809 |
| GO:0032868 | response to insulin | 16/533 | 356/28832 | 0.04494382 | 2.43118241 | 3.728965882 | 0.001072079 | 0.00934421 | 0.005986491 |
| GO:0010712 | regulation of collagen metabolic process | 6/533 | 63/28832 | 0.095238095 | 5.151791298 | 4.527425566 | 0.001080912 | 0.00934421 | 0.005986491 |
| GO:0036005 | response to macrophage colony-stimulating factor | 6/533 | 63/28832 | 0.095238095 | 5.151791298 | 4.527425566 | 0.001080912 | 0.00934421 | 0.005986491 |
| GO:0034103 | regulation of tissue remodeling | 7/533 | 86/28832 | 0.081395349 | 4.402984423 | 4.337389154 | 0.001082598 | 0.00934421 | 0.005986491 |
| GO:1904888 | cranial skeletal system development | 7/533 | 86/28832 | 0.081395349 | 4.402984423 | 4.337389154 | 0.001082598 | 0.00934421 | 0.005986491 |
| GO:0051258 | protein polymerization | 14/533 | 290/28832 | 0.048275862 | 2.611425244 | 3.785078352 | 0.001102661 | 0.009499691 | 0.006086102 |
| GO:0006814 | sodium ion transport | 13/533 | 258/28832 | 0.050387597 | 2.725657024 | 3.821085269 | 0.001116247 | 0.009598898 | 0.00614966 |
| GO:0060078 | regulation of postsynaptic membrane potential | 10/533 | 167/28832 | 0.05988024 | 3.239150217 | 3.982661589 | 0.001142018 | 0.009789031 | 0.006271472 |
| GO:0048663 | neuron fate commitment | 7/533 | 88/28832 | 0.079545455 | 4.302916596 | 4.2586628 | 0.001239377 | 0.010418881 | 0.006674993 |
| GO:0019693 | ribose phosphate metabolic process | 17/533 | 396/28832 | 0.042929293 | 2.322208956 | 3.635978846 | 0.001248619 | 0.010418881 | 0.006674993 |
| GO:0008037 | cell recognition | 10/533 | 169/28832 | 0.059171598 | 3.200817079 | 3.937988024 | 0.001249359 | 0.010418881 | 0.006674993 |
| GO:0051928 | positive regulation of calcium ion transport | 9/533 | 141/28832 | 0.063829787 | 3.452796296 | 4.006877897 | 0.001278962 | 0.01060463 | 0.006793996 |
| GO:0070227 | lymphocyte apoptotic process | 8/533 | 114/28832 | 0.070175439 | 3.796056746 | 4.10515183 | 0.001286244 | 0.010645963 | 0.006820476 |
| GO:0006937 | regulation of muscle contraction | 10/533 | 170/28832 | 0.058823529 | 3.181988743 | 3.915900368 | 0.001306034 | 0.010790495 | 0.006913072 |
| GO:0048511 | rhythmic process | 16/533 | 364/28832 | 0.043956044 | 2.37774983 | 3.630366678 | 0.001348541 | 0.011102108 | 0.007112712 |
| GO:0032964 | collagen biosynthetic process | 6/533 | 66/28832 | 0.090909091 | 4.917618966 | 4.372827406 | 0.001379805 | 0.011339351 | 0.007264704 |
| GO:0006734 | NADH metabolic process | 5/533 | 45/28832 | 0.111111111 | 6.010423181 | 4.616261196 | 0.001407008 | 0.011481481 | 0.007355762 |
| GO:0032528 | microvillus organization | 4/533 | 27/28832 | 0.148148148 | 8.013897575 | 5.003976654 | 0.001445928 | 0.011696119 | 0.007493273 |
| GO:0050860 | negative regulation of T cell receptor signaling pathway | 4/533 | 27/28832 | 0.148148148 | 8.013897575 | 5.003976654 | 0.001445928 | 0.011696119 | 0.007493273 |
| GO:0035051 | cardiocyte differentiation | 11/533 | 203/28832 | 0.054187192 | 2.931191601 | 3.789467673 | 0.001482251 | 0.011969046 | 0.007668126 |
| GO:0045824 | negative regulation of innate immune response | 8/533 | 117/28832 | 0.068376068 | 3.698721958 | 4.01425148 | 0.001519294 | 0.012183269 | 0.007805371 |
| GO:0015833 | peptide transport | 16/533 | 369/28832 | 0.043360434 | 2.345530998 | 3.570051383 | 0.001550138 | 0.012387739 | 0.007936368 |
| GO:0048845 | venous blood vessel morphogenesis | 3/533 | 13/28832 | 0.230769231 | 12.48318661 | 5.683325866 | 0.001564846 | 0.012440926 | 0.007970443 |
| GO:0050920 | regulation of chemotaxis | 12/533 | 236/28832 | 0.050847458 | 2.750532642 | 3.705795791 | 0.001582705 | 0.012561364 | 0.008047603 |
| GO:0140374 | antiviral innate immune response | 6/533 | 68/28832 | 0.088235294 | 4.772983114 | 4.274866837 | 0.001611536 | 0.012768324 | 0.008180195 |
| GO:0033622 | integrin activation | 4/533 | 28/28832 | 0.142857143 | 7.727686947 | 4.887944906 | 0.001662502 | 0.013103969 | 0.00839523 |
| GO:0045995 | regulation of embryonic development | 7/533 | 93/28832 | 0.075268817 | 4.071576994 | 4.071692454 | 0.0017095 | 0.013339284 | 0.008545987 |
| GO:0009266 | response to temperature stimulus | 10/533 | 177/28832 | 0.056497175 | 3.05614738 | 3.765724817 | 0.001764094 | 0.013673203 | 0.008759917 |
| GO:0030278 | regulation of ossification | 9/533 | 148/28832 | 0.060810811 | 3.289488363 | 3.832281072 | 0.001789414 | 0.013846298 | 0.008870813 |
| GO:0007009 | plasma membrane organization | 10/533 | 178/28832 | 0.056179775 | 3.038978013 | 3.744879333 | 0.001838997 | 0.014182618 | 0.00908628 |
| GO:0009306 | protein secretion | 18/533 | 447/28832 | 0.040268456 | 2.178274173 | 3.445589883 | 0.001849016 | 0.014236197 | 0.009120607 |
| GO:0070193 | synaptonemal complex organization | 4/533 | 29/28832 | 0.137931034 | 7.461214984 | 4.777517002 | 0.00190061 | 0.014560874 | 0.009328615 |
| GO:0051235 | maintenance of location | 12/533 | 242/28832 | 0.049586777 | 2.682337618 | 3.606796789 | 0.001952124 | 0.014906251 | 0.009549886 |
| GO:0035592 | establishment of protein localization to extracellular region | 18/533 | 450/28832 | 0.04 | 2.163752345 | 3.414705399 | 0.001987841 | 0.015030411 | 0.009629431 |
| GO:0007584 | response to nutrient | 10/533 | 180/28832 | 0.055555556 | 3.005211591 | 3.703624092 | 0.001996514 | 0.015071407 | 0.009655695 |
| GO:0071805 | potassium ion transmembrane transport | 11/533 | 211/28832 | 0.052132701 | 2.820056374 | 3.641594452 | 0.002010873 | 0.015147474 | 0.009704429 |
| GO:0010543 | regulation of platelet activation | 5/533 | 49/28832 | 0.102040816 | 5.519776391 | 4.345651283 | 0.002067334 | 0.015454992 | 0.009901444 |
| GO:1903115 | regulation of actin filament-based movement | 5/533 | 49/28832 | 0.102040816 | 5.519776391 | 4.345651283 | 0.002067334 | 0.015454992 | 0.009901444 |
| GO:0006941 | striated muscle contraction | 10/533 | 181/28832 | 0.055248619 | 2.988608212 | 3.68321047 | 0.002079254 | 0.015519072 | 0.009942498 |
| GO:0090382 | phagosome maturation | 4/533 | 30/28832 | 0.133333333 | 7.212507817 | 4.672229482 | 0.002161315 | 0.015979389 | 0.010237406 |
| GO:2000179 | positive regulation of neural precursor cell proliferation | 6/533 | 72/28832 | 0.083333333 | 4.507817386 | 4.089937737 | 0.002162477 | 0.015979389 | 0.010237406 |
| GO:0106027 | neuron projection organization | 8/533 | 124/28832 | 0.064516129 | 3.489923137 | 3.813319793 | 0.002193844 | 0.016089349 | 0.010307853 |
| GO:0071692 | protein localization to extracellular region | 18/533 | 456/28832 | 0.039473684 | 2.13528192 | 3.353655653 | 0.002291923 | 0.01675562 | 0.010734708 |
| GO:1904427 | positive regulation of calcium ion transmembrane transport | 7/533 | 98/28832 | 0.071428571 | 3.863843474 | 3.897374906 | 0.002307684 | 0.016844279 | 0.010791509 |
| GO:0042594 | response to starvation | 12/533 | 247/28832 | 0.048582996 | 2.628039286 | 3.526567265 | 0.002311462 | 0.016845327 | 0.01079218 |
| GO:0006813 | potassium ion transport | 12/533 | 248/28832 | 0.048387097 | 2.617442353 | 3.510759364 | 0.002389432 | 0.017324039 | 0.011098873 |

| | | | | | | | | | |
|---|---|---|---|---|---|---|---|---|---|
| GO:0071383 | cellular response to steroid hormone stimulus | 11/533 | 216/28832 | 0.050925926 | 2.754777291 | 3.552649237 | 0.002412754 | 0.017324039 | 0.011098873 |
| GO:0032530 | regulation of microvillus organization | 3/533 | 15/28832 | 0.2 | 10.81876173 | 5.220184831 | 0.002422002 | 0.017324039 | 0.011098873 |
| GO:0043518 | negative regulation of DNA damage response, signal transduction by p53 class mediator | 3/533 | 15/28832 | 0.2 | 10.81876173 | 5.220184831 | 0.002422002 | 0.017324039 | 0.011098873 |
| GO:0070106 | interleukin-27-mediated signaling pathway | 3/533 | 15/28832 | 0.2 | 10.81876173 | 5.220184831 | 0.002422002 | 0.017324039 | 0.011098873 |
| GO:2000035 | regulation of stem cell division | 3/533 | 15/28832 | 0.2 | 10.81876173 | 5.220184831 | 0.002422002 | 0.017324039 | 0.011098873 |
| GO:0050819 | negative regulation of coagulation | 5/533 | 51/28832 | 0.098039216 | 5.303314572 | 4.221270234 | 0.002471085 | 0.0174065 | 0.011151703 |
| GO:0034394 | protein localization to cell surface | 6/533 | 74/28832 | 0.081081081 | 4.385984484 | 4.002482086 | 0.00248607 | 0.017485488 | 0.011202308 |
| GO:0032963 | collagen metabolic process | 8/533 | 127/28832 | 0.062992126 | 3.407484008 | 3.731594195 | 0.002546478 | 0.01782919 | 0.011422505 |
| GO:0097191 | extrinsic apoptotic signaling pathway | 12/533 | 250/28832 | 0.048 | 2.596502814 | 3.479375478 | 0.002551808 | 0.017832185 | 0.011424424 |
| GO:0051336 | regulation of hydrolase activity | 17/533 | 424/28832 | 0.04009434 | 2.168855535 | 3.32760179 | 0.0025546 | 0.017832185 | 0.011424424 |
| GO:1901879 | regulation of protein depolymerization | 7/533 | 100/28832 | 0.07 | 3.786566604 | 3.830843516 | 0.002587622 | 0.018035531 | 0.0115547 |
| GO:0034764 | positive regulation of transmembrane transport | 12/533 | 251/28832 | 0.047808765 | 2.586158182 | 3.463798018 | 0.002636302 | 0.018347234 | 0.011754397 |
| GO:0033628 | regulation of cell adhesion mediated by integrin | 5/533 | 52/28832 | 0.096153846 | 5.201327753 | 4.161508183 | 0.002693094 | 0.018547533 | 0.011882721 |
| GO:0036215 | response to stem cell factor | 5/533 | 52/28832 | 0.096153846 | 5.201327753 | 4.161508183 | 0.002693094 | 0.018547533 | 0.011882721 |
| GO:0036216 | cellular response to stem cell factor stimulus | 5/533 | 52/28832 | 0.096153846 | 5.201327753 | 4.161508183 | 0.002693094 | 0.018547533 | 0.011882721 |
| GO:0035767 | endothelial cell chemotaxis | 4/533 | 32/28832 | 0.125 | 6.761726079 | 4.475476342 | 0.002754701 | 0.018721466 | 0.011994153 |
| GO:0042537 | benzene-containing compound metabolic process | 4/533 | 32/28832 | 0.125 | 6.761726079 | 4.475476342 | 0.002754701 | 0.018721466 | 0.011994153 |
| GO:0070841 | inclusion body assembly | 4/533 | 32/28832 | 0.125 | 6.761726079 | 4.475476342 | 0.002754701 | 0.018721466 | 0.011994153 |
| GO:0009914 | hormone transport | 17/533 | 428/28832 | 0.039719626 | 2.148585857 | 3.28551542 | 0.00281147 | 0.019055802 | 0.01220835 |
| GO:0044843 | cell cycle G1/S phase transition | 12/533 | 253/28832 | 0.04743083 | 2.565714243 | 3.432868454 | 0.002812118 | 0.019055802 | 0.01220835 |
| GO:0008038 | neuron recognition | 5/533 | 53/28832 | 0.094339623 | 5.103189493 | 4.103265061 | 0.002929191 | 0.019563853 | 0.01253384 |
| GO:0051602 | response to electrical stimulus | 5/533 | 53/28832 | 0.094339623 | 5.103189493 | 4.103265061 | 0.002929191 | 0.019563853 | 0.01253384 |
| GO:0090181 | regulation of cholesterol metabolic process | 5/533 | 53/28832 | 0.094339623 | 5.103189493 | 4.103265061 | 0.002929191 | 0.019563853 | 0.01253384 |
| GO:0003413 | chondrocyte differentiation involved in endochondral bone morphogenesis | 3/533 | 16/28832 | 0.1875 | 10.14258912 | 5.02019121 | 0.002940278 | 0.019563853 | 0.01253384 |
| GO:0141196 | transposable element silencing by piRNA-mediated DNA methylation | 3/533 | 16/28832 | 0.1875 | 10.14258912 | 5.02019121 | 0.002940278 | 0.019563853 | 0.01253384 |
| GO:1902950 | regulation of dendritic spine maintenance | 3/533 | 16/28832 | 0.1875 | 10.14258912 | 5.02019121 | 0.002940278 | 0.019563853 | 0.01253384 |
| GO:0032869 | cellular response to insulin stimulus | 13/533 | 290/28832 | 0.044827586 | 2.42489487 | 3.346936971 | 0.003113824 | 0.020500817 | 0.013134118 |
| GO:2000045 | regulation of G1/S transition of mitotic cell cycle | 9/533 | 161/28832 | 0.055900621 | 3.023877501 | 3.534139216 | 0.003161281 | 0.020754305 | 0.013296518 |
| GO:0120254 | olefinic compound metabolic process | 10/533 | 192/28832 | 0.052083333 | 2.817385866 | 3.467521533 | 0.003184988 | 0.020821466 | 0.013339546 |
| GO:0042113 | B cell activation | 14/533 | 326/28832 | 0.042944785 | 2.323046996 | 3.297037174 | 0.003237007 | 0.021067328 | 0.013497061 |
| GO:0008542 | visual learning | 6/533 | 78/28832 | 0.076923077 | 4.161062202 | 3.836534624 | 0.003240778 | 0.021067328 | 0.013497061 |
| GO:2000241 | regulation of reproductive process | 11/533 | 225/28832 | 0.048888889 | 2.6445862 | 3.398753405 | 0.003298645 | 0.021383526 | 0.013699637 |
| GO:0019233 | sensory perception of pain | 7/533 | 105/28832 | 0.066666667 | 3.606253909 | 3.671758839 | 0.003401728 | 0.022020964 | 0.01410802 |
| GO:0000302 | response to reactive oxygen species | 11/533 | 226/28832 | 0.048672566 | 2.632884491 | 3.382120132 | 0.003411355 | 0.022048374 | 0.014125581 |
| GO:0035790 | platelet-derived growth factor receptor-alpha signaling pathway | 5/533 | 55/28832 | 0.090909091 | 4.917618966 | 3.991064007 | 0.00344565 | 0.022048374 | 0.014125581 |
| GO:0001845 | phagolysosome assembly | 3/533 | 17/28832 | 0.176470588 | 9.545966229 | 4.837090735 | 0.003521699 | 0.022190434 | 0.014216593 |
| GO:0071285 | cellular response to lithium ion | 3/533 | 17/28832 | 0.176470588 | 9.545966229 | 4.837090735 | 0.003521699 | 0.022190434 | 0.014216593 |
| GO:0141176 | gene silencing by piRNA-directed DNA methylation | 3/533 | 17/28832 | 0.176470588 | 9.545966229 | 4.837090735 | 0.003521699 | 0.022190434 | 0.014216593 |
| GO:0006935 | chemotaxis | 18/533 | 475/28832 | 0.037894737 | 2.049870643 | 3.166360565 | 0.003523659 | 0.022190434 | 0.014216593 |
| GO:0030308 | negative regulation of cell growth | 10/533 | 195/28832 | 0.051282051 | 2.774041468 | 3.411341724 | 0.003556085 | 0.022333948 | 0.014308537 |
| GO:0015748 | organophosphate ester transport | 9/533 | 164/28832 | 0.054878049 | 2.968562669 | 3.469607789 | 0.003572349 | 0.022405732 | 0.014354527 |
| GO:1902903 | regulation of supramolecular fiber organization | 16/533 | 403/28832 | 0.039702233 | 2.147645008 | 3.184103793 | 0.003704683 | 0.023023846 | 0.01475053 |
| GO:0035116 | embryonic hindlimb morphogenesis | 4/533 | 35/28832 | 0.114285714 | 6.182149558 | 4.209962764 | 0.003840041 | 0.023668338 | 0.015163432 |
| GO:0019882 | antigen processing and presentation | 8/533 | 136/28832 | 0.058823529 | 3.181988743 | 3.500412218 | 0.003875275 | 0.023853784 | 0.01528224 |
| GO:0043113 | receptor clustering | 6/533 | 81/28832 | 0.074074074 | 4.006948787 | 3.719203798 | 0.00390997 | 0.024003589 | 0.015378215 |
| GO:0051147 | regulation of muscle cell differentiation | 9/533 | 167/28832 | 0.053892216 | 2.915235196 | 3.406530561 | 0.004024237 | 0.024608618 | 0.015765835 |
| GO:0032941 | secretion by tissue | 5/533 | 57/28832 | 0.087719298 | 4.745070932 | 3.884165464 | 0.004024451 | 0.024608618 | 0.015765835 |
| GO:0006116 | NADH oxidation | 3/533 | 18/28832 | 0.166666667 | 9.015634772 | 4.66853168 | 0.004168523 | 0.025190488 | 0.016138617 |
| GO:0070365 | hepatocyte differentiation | 3/533 | 18/28832 | 0.166666667 | 9.015634772 | 4.66853168 | 0.004168523 | 0.025190488 | 0.016138617 |
| GO:0140966 | piRNA-mediated heterochromatin formation | 3/533 | 18/28832 | 0.166666667 | 9.015634772 | 4.66853168 | 0.004168523 | 0.025190488 | 0.016138617 |
| GO:1902093 | positive regulation of flagellated sperm motility | 3/533 | 18/28832 | 0.166666667 | 9.015634772 | 4.66853168 | 0.004168523 | 0.025190488 | 0.016138617 |
| GO:2000155 | positive regulation of cilium-dependent cell motility | 3/533 | 18/28832 | 0.166666667 | 9.015634772 | 4.66853168 | 0.004168523 | 0.025190488 | 0.016138617 |
| GO:0003401 | axis elongation | 4/533 | 36/28832 | 0.111111111 | 6.010423181 | 4.128264252 | 0.004257853 | 0.025497608 | 0.016335378 |
| GO:0010526 | transposable element silencing | 4/533 | 36/28832 | 0.111111111 | 6.010423181 | 4.128264252 | 0.004257853 | 0.025497608 | 0.016335378 |
| GO:0042886 | amide transport | 16/533 | 410/28832 | 0.03902439 | 2.110977898 | 3.109409655 | 0.004365819 | 0.026043206 | 0.016684923 |
| GO:0051495 | positive regulation of cytoskeleton organization | 10/533 | 201/28832 | 0.049751244 | 2.69123426 | 3.302109351 | 0.004400865 | 0.026151292 | 0.016754169 |
| GO:0060759 | regulation of response to cytokine stimulus | 9/533 | 170/28832 | 0.052941176 | 2.863789869 | 3.344845594 | 0.004519626 | 0.026719984 | 0.017118509 |
| GO:0009612 | response to mechanical stimulus | 12/533 | 270/28832 | 0.044444444 | 2.404169272 | 3.18137565 | 0.00472407 | 0.027822193 | 0.017824654 |
| GO:0051851 | host-mediated perturbation of symbiont process | 7/533 | 112/28832 | 0.0625 | 3.380863039 | 3.464648014 | 0.00485246 | 0.028360714 | 0.018169664 |
| GO:0097061 | dendritic spine organization | 7/533 | 112/28832 | 0.0625 | 3.380863039 | 3.464648014 | 0.00485246 | 0.028360714 | 0.018169664 |
| GO:0003353 | positive regulation of cilium movement | 3/533 | 19/28832 | 0.157894737 | 8.541127678 | 4.512599259 | 0.004882815 | 0.028360714 | 0.018169664 |
| GO:0030033 | microvillus assembly | 3/533 | 19/28832 | 0.157894737 | 8.541127678 | 4.512599259 | 0.004882815 | 0.028360714 | 0.018169664 |
| GO:0001825 | blastocyst formation | 5/533 | 60/28832 | 0.083333333 | 4.507817386 | 3.732806594 | 0.00501822 | 0.028880963 | 0.018502969 |
| GO:0031529 | ruffle organization | 5/533 | 60/28832 | 0.083333333 | 4.507817386 | 3.732806594 | 0.00501822 | 0.028880963 | 0.018502969 |
| GO:1902930 | regulation of alcohol biosynthetic process | 5/533 | 60/28832 | 0.083333333 | 4.507817386 | 3.732806594 | 0.00501822 | 0.028880963 | 0.018502969 |
| GO:0002040 | sprouting angiogenesis | 8/533 | 142/28832 | 0.056338028 | 3.047538514 | 3.356749174 | 0.005022234 | 0.028880963 | 0.018502969 |
| GO:0032197 | retrotransposition | 4/533 | 38/28832 | 0.105263158 | 5.694085119 | 3.973742046 | 0.00518323 | 0.029696254 | 0.019025295 |
| GO:0046879 | hormone secretion | 16/533 | 418/28832 | 0.038277512 | 2.070576407 | 3.025850969 | 0.005236512 | 0.029830102 | 0.019111047 |
| GO:0002237 | response to molecule of bacterial origin | 18/533 | 495/28832 | 0.036363636 | 1.967047587 | 2.978389024 | 0.005368433 | 0.030400879 | 0.019476723 |
| GO:0009267 | cellular response to starvation | 10/533 | 207/28832 | 0.048309179 | 2.61322747 | 3.196803534 | 0.005396367 | 0.030428422 | 0.019494369 |
| GO:0070228 | regulation of lymphocyte apoptotic process | 6/533 | 87/28832 | 0.068965517 | 3.730607492 | 3.500625566 | 0.005547895 | 0.031206911 | 0.019993118 |
| GO:1990830 | cellular response to leukemia inhibitory factor | 13/533 | 311/28832 | 0.041800643 | 2.261155988 | 3.068842071 | 0.005572129 | 0.031305235 | 0.02005611 |
| GO:0034587 | piRNA processing | 3/533 | 20/28832 | 0.15 | 8.114071295 | 4.367716295 | 0.005666457 | 0.031336446 | 0.020076106 |
| GO:0050746 | regulation of lipoprotein metabolic process | 3/533 | 20/28832 | 0.15 | 8.114071295 | 4.367716295 | 0.005666457 | 0.031336446 | 0.020076106 |
| GO:0060973 | cell migration involved in heart development | 3/533 | 20/28832 | 0.15 | 8.114071295 | 4.367716295 | 0.005666457 | 0.031336446 | 0.020076106 |
| GO:1900424 | regulation of defense response to bacterium | 3/533 | 20/28832 | 0.15 | 8.114071295 | 4.367716295 | 0.005666457 | 0.031336446 | 0.020076106 |
| GO:0031342 | negative regulation of cell killing | 4/533 | 39/28832 | 0.102564103 | 5.548082936 | 3.900543571 | 0.005692619 | 0.031336446 | 0.020076106 |
| GO:0061744 | motor behavior | 4/533 | 39/28832 | 0.102564103 | 5.548082936 | 3.900543571 | 0.005692619 | 0.031336446 | 0.020076106 |
| GO:0051963 | regulation of synapse assembly | 10/533 | 209/28832 | 0.04784689 | 2.588220509 | 3.162527281 | 0.005764677 | 0.031650517 | 0.02027732 |
| GO:0007140 | male meiotic nuclear division | 5/533 | 62/28832 | 0.080645161 | 4.362403922 | 3.637338423 | 0.005770159 | 0.031650517 | 0.02027732 |
| GO:0007033 | vacuole organization | 11/533 | 243/28832 | 0.04526749 | 2.448690926 | 3.112341956 | 0.005846266 | 0.031992258 | 0.020496261 |
| GO:1990823 | response to leukemia inhibitory factor | 13/533 | 313/28832 | 0.041533546 | 2.246707707 | 3.043529946 | 0.00587024 | 0.032010072 | 0.020507673 |
| GO:0070527 | platelet aggregation | 5/533 | 63/28832 | 0.079365079 | 4.293159415 | 3.591108767 | 0.006174404 | 0.033390072 | 0.021391788 |
| GO:0015850 | organic hydroxy compound transport | 13/533 | 315/28832 | 0.041269841 | 2.232442896 | 3.018408935 | 0.006180945 | 0.033390072 | 0.021391788 |
| GO:0001964 | startle response | 4/533 | 40/28832 | 0.1 | 5.409380863 | 3.829830952 | 0.006234318 | 0.033444519 | 0.02142667 |

| | | | | | | | | | |
|---|---|---|---|---|---|---|---|---|---|
| GO:0086004 | regulation of cardiac muscle cell contraction | 4/533 | 40/28832 | 0.1 | 5.409380863 | 3.829830952 | 0.006234318 | 0.033444519 | 0.02142667 |
| GO:1903522 | regulation of blood circulation | 12/533 | 280/28832 | 0.042857143 | 2.318306084 | 3.042180075 | 0.006262255 | 0.033555549 | 0.021497804 |
| GO:0002687 | positive regulation of leukocyte migration | 9/533 | 180/28832 | 0.05 | 2.704690432 | 3.148561769 | 0.006520684 | 0.034583006 | 0.022156058 |
| GO:0000729 | DNA double-strand break processing | 3/533 | 21/28832 | 0.142857143 | 7.727686947 | 4.232570189 | 0.006521154 | 0.034583006 | 0.022156058 |
| GO:0003228 | atrial cardiac muscle tissue development | 3/533 | 21/28832 | 0.142857143 | 7.727686947 | 4.232570189 | 0.006521154 | 0.034583006 | 0.022156058 |
| GO:0006071 | glycerol metabolic process | 3/533 | 21/28832 | 0.142857143 | 7.727686947 | 4.232570189 | 0.006521154 | 0.034583006 | 0.022156058 |
| GO:2000736 | regulation of stem cell differentiation | 6/533 | 90/28832 | 0.066666667 | 3.606253909 | 3.398500975 | 0.006532551 | 0.034603855 | 0.022169415 |
| GO:0046034 | ATP metabolic process | 11/533 | 247/28832 | 0.044534413 | 2.409036012 | 3.052174673 | 0.006581099 | 0.034713226 | 0.022239484 |
| GO:0060324 | face development | 5/533 | 64/28832 | 0.078125 | 4.226078799 | 3.545831029 | 0.006598134 | 0.034713226 | 0.022239484 |
| GO:0099003 | vesicle-mediated transport in synapse | 13/533 | 318/28832 | 0.040880503 | 2.211382114 | 2.981079495 | 0.006671424 | 0.035059015 | 0.022461019 |
| GO:0016311 | dephosphorylation | 9/533 | 181/28832 | 0.049723757 | 2.68974739 | 3.129673932 | 0.006753084 | 0.035327929 | 0.022633302 |
| GO:0031128 | developmental induction | 4/533 | 41/28832 | 0.097560976 | 5.277444744 | 3.761455036 | 0.006809179 | 0.035501177 | 0.022744296 |
| GO:0052652 | cyclic purine nucleotide metabolic process | 4/533 | 41/28832 | 0.097560976 | 5.277444744 | 3.761455036 | 0.006809179 | 0.035501177 | 0.022744296 |
| GO:0006816 | calcium ion transport | 17/533 | 470/28832 | 0.036170213 | 1.956584567 | 2.869536715 | 0.007051927 | 0.036561167 | 0.023423393 |
| GO:0070588 | calcium ion transmembrane transport | 14/533 | 359/28832 | 0.038997214 | 2.109507857 | 2.903146189 | 0.007433338 | 0.037854738 | 0.024252135 |
| GO:0009074 | aromatic amino acid family catabolic process | 3/533 | 22/28832 | 0.136363636 | 7.37642845 | 4.106058334 | 0.00744844 | 0.037854738 | 0.024252135 |
| GO:0034349 | glial cell apoptotic process | 3/533 | 22/28832 | 0.136363636 | 7.37642845 | 4.106058334 | 0.00744844 | 0.037854738 | 0.024252135 |
| GO:0051341 | regulation of oxidoreductase activity | 5/533 | 66/28832 | 0.075757576 | 4.098015805 | 3.457990309 | 0.007505917 | 0.038063373 | 0.0243858 |
| GO:0099171 | presynaptic modulation of chemical synaptic transmission | 5/533 | 66/28832 | 0.075757576 | 4.098015805 | 3.457990309 | 0.007505917 | 0.038063373 | 0.0243858 |
| GO:0120034 | positive regulation of plasma membrane bounded cell projection assembly | 7/533 | 122/28832 | 0.057377049 | 3.103743118 | 3.195684825 | 0.007675145 | 0.038794217 | 0.024854025 |
| GO:0035725 | sodium ion transmembrane transport | 9/533 | 185/28832 | 0.048648649 | 2.63159069 | 3.055381338 | 0.00774677 | 0.03907103 | 0.025031369 |
| GO:0022411 | cellular component disassembly | 16/533 | 437/28832 | 0.036613272 | 1.980551346 | 2.834640808 | 0.007877849 | 0.039602854 | 0.025372089 |
| GO:0022600 | digestive system process | 7/533 | 123/28832 | 0.056910569 | 3.078509434 | 3.170322451 | 0.008012466 | 0.040135136 | 0.025713102 |
| GO:0033627 | cell adhesion mediated by integrin | 6/533 | 94/28832 | 0.063829787 | 3.452796296 | 3.268925712 | 0.008035066 | 0.040135136 | 0.025713102 |
| GO:0030866 | cortical actin cytoskeleton organization | 4/533 | 43/28832 | 0.093023256 | 5.031982198 | 3.631177318 | 0.008061664 | 0.040135136 | 0.025713102 |
| GO:0061081 | positive regulation of myeloid leukocyte cytokine production involved in immune response | 4/533 | 43/28832 | 0.093023256 | 5.031982198 | 3.631177318 | 0.008061664 | 0.040135136 | 0.025713102 |
| GO:0071398 | cellular response to fatty acid | 4/533 | 43/28832 | 0.093023256 | 5.031982198 | 3.631177318 | 0.008061664 | 0.040135136 | 0.025713102 |
| GO:0031345 | negative regulation of cell projection organization | 10/533 | 220/28832 | 0.045454545 | 2.458809483 | 2.980874714 | 0.008153873 | 0.040463812 | 0.025923673 |
| GO:0043524 | negative regulation of neuron apoptotic process | 10/533 | 221/28832 | 0.045248869 | 2.447683648 | 2.964907852 | 0.008403947 | 0.041443722 | 0.026551466 |
| GO:0010921 | regulation of phosphatase activity | 3/533 | 23/28832 | 0.130434783 | 7.055714169 | 3.987246701 | 0.008449691 | 0.041443722 | 0.026551466 |
| GO:0030903 | notochord development | 3/533 | 23/28832 | 0.130434783 | 7.055714169 | 3.987246701 | 0.008449691 | 0.041443722 | 0.026551466 |
| GO:0044851 | hair cycle phase | 3/533 | 23/28832 | 0.130434783 | 7.055714169 | 3.987246701 | 0.008449691 | 0.041443722 | 0.026551466 |
| GO:0055062 | phosphate ion homeostasis | 3/533 | 23/28832 | 0.130434783 | 7.055714169 | 3.987246701 | 0.008449691 | 0.041443722 | 0.026551466 |
| GO:0033555 | multicellular organismal response to stress | 7/533 | 125/28832 | 0.056 | 3.029253283 | 3.120364096 | 0.00871971 | 0.042467376 | 0.027207283 |
| GO:0019674 | NAD metabolic process | 4/533 | 44/28832 | 0.090909091 | 4.917618966 | 3.569034103 | 0.008740858 | 0.042467376 | 0.027207283 |
| GO:0000082 | G1/S transition of mitotic cell cycle | 10/533 | 223/28832 | 0.044843049 | 2.425731329 | 2.933233919 | 0.008921735 | 0.043255485 | 0.027712195 |
| GO:0010907 | positive regulation of glucose metabolic process | 5/533 | 69/28832 | 0.072463768 | 3.919841205 | 3.332534091 | 0.009025255 | 0.043439309 | 0.027829964 |
| GO:0030193 | regulation of blood coagulation | 5/533 | 69/28832 | 0.072463768 | 3.919841205 | 3.332534091 | 0.009025255 | 0.043439309 | 0.027829964 |
| GO:0061082 | myeloid leukocyte cytokine production | 5/533 | 69/28832 | 0.072463768 | 3.919841205 | 3.332534091 | 0.009025255 | 0.043439309 | 0.027829964 |
| GO:0006644 | phospholipid metabolic process | 14/533 | 368/28832 | 0.038043478 | 2.057916633 | 2.80307877 | 0.00912262 | 0.043816939 | 0.028071898 |
| GO:0003018 | vascular process in circulatory system | 11/533 | 259/28832 | 0.042471042 | 2.297420444 | 2.878462735 | 0.00922603 | 0.044221974 | 0.028331389 |
| GO:0006470 | protein dephosphorylation | 6/533 | 97/28832 | 0.06185567 | 3.346008781 | 3.176272926 | 0.00931431 | 0.044552971 | 0.028543446 |
| GO:0046677 | response to antibiotic | 4/533 | 45/28832 | 0.088888889 | 4.808338545 | 3.508742637 | 0.009456356 | 0.044735144 | 0.028660158 |
| GO:1900744 | regulation of p38MAPK cascade | 4/533 | 45/28832 | 0.088888889 | 4.808338545 | 3.508742637 | 0.009456356 | 0.044735144 | 0.028660158 |
| GO:0017157 | regulation of exocytosis | 10/533 | 225/28832 | 0.044444444 | 2.404169272 | 2.901900245 | 0.009463622 | 0.044735144 | 0.028660158 |
| GO:0002709 | regulation of T cell mediated immunity | 7/533 | 127/28832 | 0.05511811 | 2.981548507 | 3.071395233 | 0.009471733 | 0.044735144 | 0.028660158 |
| GO:0048145 | regulation of fibroblast proliferation | 7/533 | 127/28832 | 0.05511811 | 2.981548507 | 3.071395233 | 0.009471733 | 0.044735144 | 0.028660158 |
| GO:0002819 | regulation of adaptive immune response | 11/533 | 260/28832 | 0.042307692 | 2.288584211 | 2.864422471 | 0.009478813 | 0.044735144 | 0.028660158 |
| GO:0046579 | positive regulation of Ras protein signal transduction | 3/533 | 24/28832 | 0.125 | 6.761726079 | 3.875338002 | 0.009526124 | 0.044735144 | 0.028660158 |
| GO:0097398 | cellular response to interleukin-17 | 3/533 | 24/28832 | 0.125 | 6.761726079 | 3.875338002 | 0.009526124 | 0.044735144 | 0.028660158 |
| GO:0141005 | transposable element silencing by heterochromatin formation | 3/533 | 24/28832 | 0.125 | 6.761726079 | 3.875338002 | 0.009526124 | 0.044735144 | 0.028660158 |
| GO:0046503 | glycerolipid catabolic process | 5/533 | 70/28832 | 0.071428571 | 3.863843474 | 3.292279285 | 0.009575528 | 0.044876209 | 0.028750533 |
| GO:0002460 | adaptive immune response based on somatic recombination of immune receptors built from immunoglobulin superfamily domains | 17/533 | 487/28832 | 0.034907598 | 1.888284901 | 2.713228954 | 0.009823731 | 0.045895401 | 0.029403492 |
| GO:0044782 | cilium organization | 16/533 | 449/28832 | 0.035634744 | 1.927619016 | 2.71876442 | 0.010034272 | 0.046789589 | 0.029976365 |
| GO:0010935 | regulation of macrophage cytokine production | 4/533 | 46/28832 | 0.086956522 | 4.703809446 | 3.45020423 | 0.010208873 | 0.047129607 | 0.030194202 |
| GO:0048730 | epidermis morphogenesis | 4/533 | 46/28832 | 0.086956522 | 4.703809446 | 3.45020423 | 0.010208873 | 0.047129607 | 0.030194202 |
| GO:0006766 | vitamin metabolic process | 6/533 | 99/28832 | 0.060606061 | 3.278412644 | 3.116501972 | 0.010243954 | 0.047197542 | 0.030237726 |
| GO:0045069 | regulation of viral genome replication | 6/533 | 99/28832 | 0.060606061 | 3.278412644 | 3.116501972 | 0.010243954 | 0.047197542 | 0.030237726 |
| GO:0071695 | anatomical structure maturation | 13/533 | 336/28832 | 0.038690476 | 2.092915215 | 2.765489874 | 0.010298442 | 0.047354441 | 0.030338245 |
| GO:0043467 | regulation of generation of precursor metabolites and energy | 8/533 | 161/28832 | 0.049689441 | 2.687891112 | 2.947432425 | 0.010405468 | 0.047751828 | 0.030592836 |
| GO:0055064 | chloride ion homeostasis | 3/533 | 25/28832 | 0.12 | 6.491257036 | 3.769646914 | 0.010678811 | 0.048585671 | 0.031127049 |
| GO:1901570 | fatty acid derivative biosynthetic process | 3/533 | 25/28832 | 0.12 | 6.491257036 | 3.769646914 | 0.010678811 | 0.048585671 | 0.031127049 |
| GO:2000345 | regulation of hepatocyte proliferation | 3/533 | 25/28832 | 0.12 | 6.491257036 | 3.769646914 | 0.010678811 | 0.048585671 | 0.031127049 |
| GO:0035821 | modulation of process of another organism | 7/533 | 130/28832 | 0.053846154 | 2.912743542 | 2.999716786 | 0.010687149 | 0.048585671 | 0.031127049 |
| GO:0050688 | regulation of defense response to virus | 5/533 | 72/28832 | 0.069444444 | 3.756514488 | 3.21395654 | 0.010744404 | 0.048585671 | 0.031127049 |
| GO:0070542 | response to fatty acid | 5/533 | 72/28832 | 0.069444444 | 3.756514488 | 3.21395654 | 0.010744404 | 0.048585671 | 0.031127049 |
| GO:0009615 | response to virus | 15/533 | 414/28832 | 0.036231884 | 1.959920603 | 2.699893551 | 0.010807477 | 0.048728264 | 0.031218403 |
| GO:0045055 | regulated exocytosis | 11/533 | 265/28832 | 0.041509434 | 2.245403377 | 2.795172221 | 0.010823775 | 0.048754323 | 0.031235098 |
| GO:0050892 | intestinal absorption | 4/533 | 47/28832 | 0.085106383 | 4.603728394 | 3.3933275 | 0.010999097 | 0.049352194 | 0.031618132 |
| GO:0016032 | viral process | 13/533 | 339/28832 | 0.038348083 | 2.074393841 | 2.730877273 | 0.011029246 | 0.049439609 | 0.031674136 |
| GO:0021915 | neural tube development | 9/533 | 196/28832 | 0.045918367 | 2.483899376 | 2.860778439 | 0.011056185 | 0.04951248 | 0.031720821 |
| GO:0055007 | cardiac muscle cell differentiation | 8/533 | 163/28832 | 0.049079755 | 2.654910853 | 2.907836924 | 0.011153943 | 0.049902052 | 0.031970405 |

**Supplementary Table 15:** Summary of GO process terms associated with DEGs exhibiting >2 fold upregulation in marked *Tead4* KD versus unmarked outer control cell clones at the E3.5 stage (RNA-Seq data). Relates to data in Fig. S11.

**Supplementary Tables 16: RT-qPCR (*Krt8* & *Krt18* KD, *Tead4* KD, Rnd1 & Rnd3 KD and *Tead4* , *Rnd1* & *Rnd3* KD)**

si*K8* & si*K18*

| | | | H2afz | *Krt8* | | | *Krt18* | | |
|---|---|---|---|---|---|---|---|---|---|
| | | | Ct | Ct | 2^-ΔCt | 2^-ΔΔCt | Ct | 2^-ΔCt | 2^-ΔΔCt |
| BioRep1 | Tech1 | si*K8* & si*K18* | 29.34 | 32.01 | 0.152 | 0.320 | 35.13 | 0.018 | 0.120 |
| | | | 29.25 | 32.12 | 0.141 | 0.296 | 34.02 | 0.038 | 0.259 |
| | | | | 32.07 | 0.146 | 0.307 | 34.8 | 0.022 | 0.151 |
| | | siNTC | 22.38 | 23.47 | 0.459 | 0.964 | 25.22 | 0.136 | 0.933 |
| | | | 22.31 | 23.4 | 0.482 | 1.012 | 25.05 | 0.154 | 1.050 |
| | | | 22.35 | 23.38 | 0.489 | 1.026 | 25.09 | 0.149 | 1.021 |
| | Tech2 | si*K8* & si*K18* | 29.19 | 32.34 | 0.115 | 0.214 | 34.15 | 0.033 | 0.185 |
| | | | 29.3 | 32.52 | 0.102 | 0.189 | 34.29 | 0.030 | 0.168 |
| | | | 29.18 | | | | 34.35 | 0.029 | 0.161 |
| | | siNTC | 22.71 | 23.64 | 0.545 | 1.012 | 25.32 | 0.170 | 0.955 |
| | | | 22.57 | 23.63 | 0.548 | 1.019 | 25.19 | 0.186 | 1.045 |
| | | | 23.01 | 23.7 | 0.522 | 0.970 | 25.25 | 0.178 | 1.002 |

si*Tead4*

| | | | H2afz | *Tead4* | | | *Rnd1* | | | *Rnd3* | | |
|---|---|---|---|---|---|---|---|---|---|---|---|---|
| | | | Ct | Ct | 2^-ΔCt | 2^-ΔΔCt | Ct | 2^-ΔCt | 2^-ΔΔCt | Ct | 2^-ΔCt | 2^-ΔΔCt |
| BioRep1 | Tech1 | si*Tead4* | 24.12 | 33.85 | 0.001 | 0.057 | 30.65 | 0.011 | 1.437 | 27 | 0.136 | 2.056 |
| | | | 24.11 | 33.94 | 0.001 | 0.053 | 30.59 | 0.011 | 1.498 | 28.66 | 0.043 | 1.117 |
| | | | 24.12 | 33.86 | 0.001 | 0.056 | 30.67 | 0.011 | 1.417 | 28.46 | 0.049 | 1.283 |
| | | siNTC | 22.51 | 28.18 | 0.019 | 0.942 | 29.67 | 0.007 | 0.922 | N/A | | |
| | | | 22.43 | 28.03 | 0.022 | 1.045 | 29.58 | 0.007 | 0.982 | 27.08 | 0.042 | 1.087 |
| | | | 22.55 | 28.07 | 0.021 | 1.016 | 29.41 | 0.008 | 1.104 | 27.32 | 0.035 | 0.920 |

ds*Rnd1* + ds*Rnd3*

| | | | H2afz | *Tead4* | | | *Rnd1* | | | *Rnd3* | | |
|---|---|---|---|---|---|---|---|---|---|---|---|---|
| | | | Ct | Ct | 2^-ΔCt | 2^-ΔΔCt | Ct | 2^-ΔCt | 2^-ΔΔCt | Ct | 2^-ΔCt | 2^-ΔΔCt |
| BioRep2 | Tech3 | ds*Rnd1* + ds*Rnd3* | 23.63 | 29.38 | 0.019 | 0.810 | 31.27 | 0.005 | 0.260 | 30.44 | 0.009 | 0.217 |
| | | | 23.45 | 29.23 | 0.021 | 0.899 | 31.32 | 0.005 | 0.251 | 30.69 | 0.008 | 0.182 |
| | | | 23.91 | 29.1 | 0.023 | 0.984 | 31.89 | 0.003 | 0.169 | 30.95 | 0.006 | 0.152 |
| | | ds*EGFP* | 23.93 | 28.89 | 0.022 | 0.935 | 29.36 | 0.016 | 0.803 | 28.03 | 0.040 | 0.946 |
| | | | 23.13 | 28.74 | 0.024 | 1.038 | 28.77 | 0.024 | 1.209 | 27.82 | 0.046 | 1.094 |
| | | | 23.08 | 28.75 | 0.024 | 1.030 | 29.00 | 0.020 | 1.030 | 28.00 | 0.041 | 0.966 |

si*Tead4* + ds*Rnd1* + ds*Rnd3*

| | | | H2afz | *Tead4* | | | *Rnd1* | | | *Rnd3* | | |
|---|---|---|---|---|---|---|---|---|---|---|---|---|
| | | | Ct | Ct | 2^-ΔCt | 2^-ΔΔCt | Ct | 2^-ΔCt | 2^-ΔΔCt | Ct | 2^-ΔCt | 2^-ΔΔCt |
| BioRep3 | Tech5 | si*Tead4* + ds*Rnd1* + ds*Rnd3* | 23 | 32.45 | 0.001 | 0.083 | 30.260 | 0.007 | 0.652 | 27.470 | 0.046 | 1.014 |
| | | | 23.02 | 32.32 | 0.002 | 0.090 | 29.750 | 0.010 | 0.929 | 27.360 | 0.050 | 1.094 |
| | | | 23.1 | 31.7 | 0.002 | 0.139 | 30.150 | 0.007 | 0.704 | 27.300 | 0.052 | 1.141 |
| | | siNTC + ds*EGFP* | 21.89 | 27.4 | 0.021 | 1.159 | 28.500 | 0.010 | 0.935 | 26.290 | 0.045 | 0.973 |
| | | | 21.71 | 27.37 | 0.021 | 1.184 | 28.240 | 0.012 | 1.120 | 26.280 | 0.045 | 0.979 |
| | | | 21.8 | 28.07 | 0.013 | 0.729 | 28.470 | 0.010 | 0.955 | 26.180 | 0.048 | 1.050 |
| | Tech6 | si*Tead4* + ds*Rnd1* + ds*Rnd3* | 23.02 | 32.48 | 0.001 | 0.082 | 30.280 | 0.007 | 0.654 | 27.490 | 0.046 | 1.009 |
| | | | 23.04 | 32.34 | 0.002 | 0.090 | 29.780 | 0.009 | 0.924 | 27.380 | 0.050 | 1.089 |
| | | | 23.12 | 31.73 | 0.002 | 0.138 | 30.170 | 0.007 | 0.705 | 27.320 | 0.052 | 1.136 |
| | | siNTC + ds*EGFP* | 21.92 | 27.42 | 0.021 | 1.162 | 28.530 | 0.010 | 0.935 | 26.310 | 0.045 | 0.973 |
| | | | 21.73 | 27.4 | 0.021 | 1.178 | 28.270 | 0.011 | 1.120 | 26.300 | 0.045 | 0.979 |
| | | | 21.83 | 28.09 | 0.013 | 0.730 | 28.500 | 0.010 | 0.955 | 26.200 | 0.048 | 1.050 |

**Supplementary Tables S16:** Summary of confirmatory RT-qPCR data showing the effects of global gene knockdown (KD), incorporating controls, by combined si*K8* and si*K18* (assaying *Krt8* and *Krt18* expression), si*Tead4* (assaying *Tead4*, *Rnd1* and *Rnd3* expression), combined ds*Rnd1* and ds*Rnd3* (assaying *Tead4*, *Rnd1* and *Rnd3* expression) and combined si*Tead4,* ds*Rnd1* and ds*Rnd3* (assaying *Tead4*, *Rnd1* and *Rnd3* expression); after microinjection into both 2-cell blastomeres and culture to the E3.5 early-blastocyst stage and normalisation of all data to *H2afz* expression levels. Relates to data in Figs. 5 and S12 & S14.

Supplementary Tables 17: KRT8 normalised fluorescence intensity (Relative Fluorescence Units - RFU)

| | Whole embryo | | Individual outer cells | | | |
|---|---|---|---|---|---|---|
| | | | siNTC | | si*Tead4* | |
| | siNTC | si*Tead4* | Unmarked | Marked | Unmarked | Marked |
| | 1316758.10 | 1019456.38 | 20419.80 | 52008.59 | 39066.88 | 2026.77 |
| | 638946.23 | 195489.58 | 55604.10 | 71944.95 | 64419.58 | 6665.44 |
| | 190440.22 | 495666.76 | 60534.20 | 30273.70 | 2981.22 | 1847.77 |
| | 102821.37 | 137470.92 | 22682.00 | 24672.65 | 1001.32 | 615.71 |
| | 79119.46 | 388294.49 | 4269.35 | 4775.72 | 2729.53 | 1193.04 |
| | 940986.35 | 125032.56 | 46762.20 | 53828.34 | 12572.43 | 338.59 |
| | 465366.87 | 73528.28 | 24066.00 | 20001.23 | 3252.63 | 687.78 |
| | 312887.32 | 717261.64 | 21761.80 | 37743.86 | 56701.87 | 480.99 |
| | 399902.07 | 135194.03 | 19729.00 | 26833.33 | 16574.65 | 873.02 |
| | 444963.77 | 377826.11 | 54704.90 | 6623.58 | 7360.06 | 911.79 |
| | 208507.56 | 304265.62 | 27048.70 | 30679.63 | 6611.97 | 564.88 |
| | 1144408.87 | 842820.85 | 2140.17 | 3420.83 | 4221.92 | 2169.24 |
| | 1601371.54 | 169504.59 | 1188.88 | 10458.27 | 2592.09 | 269.48 |
| | 621753.18 | 466300.13 | 35692.60 | 2578.48 | 4508.02 | 1100.91 |
| | 216057.94 | 32604.21 | 1994.76 | 2463.29 | 5131.38 | 2063.07 |
| | 674647.73 | 538396.36 | 13849.90 | 871.13 | 201.89 | 2098.82 |
| | 648432.83 | 144466.36 | 25352.40 | 15295.36 | 8332.07 | 1360.68 |
| | 255672.89 | 1157121.81 | 1335.68 | 890.39 | 1074.47 | 1587.69 |
| | 241462.76 | 567600.24 | 15542.10 | 4641.29 | 47.93 | 2031.79 |
| | 891058.81 | 30532.67 | 2794.25 | 3638.42 | 48325.41 | 2239.87 |
| | 1260942.28 | 76739.31 | 6241.18 | 15238.25 | 10838.95 | 557.65 |
| | 1074310.56 | 42353.05 | 5269.97 | 11367.56 | 22638.38 | 716.29 |
| | 1140504.55 | 12528.24 | 5244.91 | 888.53 | 10317.39 | 2821.18 |
| | | 1208342.59 | 13509.90 | 25218.02 | 3243.05 | 4234.77 |
| | | 287358.68 | 5050.84 | 8765.46 | 6012.67 | 248.82 |
| | | | 4320.07 | 16836.10 | 5352.84 | 2012.25 |
| | | | 12263.60 | 8638.29 | 2408.59 | 541.04 |
| | | | 18222.90 | 1115.69 | 4422.15 | 1765.40 |
| | | | 30653.90 | 22663.35 | 3442.96 | 423.20 |
| | | | 8060.78 | 29114.84 | 1893.49 | 158.75 |
| | | | 7612.06 | 7854.98 | 1000.51 | 374.59 |
| | | | 34743.30 | 6400.19 | 1021.95 | 117.24 |
| | | | 23575.40 | 6297.74 | 9205.57 | 491.17 |
| | | | 1238.47 | 11292.10 | 1393.59 | 381.43 |
| | | | 5515.43 | 3114.83 | 24524.75 | 826.82 |
| | | | 16186.60 | 1943.16 | 3521.64 | 1297.42 |
| | | | 3293.41 | 3213.53 | 23096.60 | 179.82 |
| | | | 5710.81 | 4483.12 | 7783.94 | 796.16 |
| | | | 2332.27 | 9323.92 | 17176.11 | 2.77 |
| | | | 16916.30 | 20090.66 | 2510.28 | 420.16 |
| | | | 1558.98 | 3719.25 | 12886.11 | 3.68 |
| | | | 994.29 | 2660.83 | 22297.33 | 165.71 |
| | | | 37332.90 | 7052.59 | 4923.01 | 2853.80 |
| | | | 35940.30 | 21303.92 | 6685.62 | 1439.60 |
| | | | 15961.80 | 21077.67 | 7527.98 | 469.35 |
| | | | 9259.79 | 18455.32 | 13509.11 | 1064.56 |
| | | | 6750.59 | 48294.84 | 1620.41 | 404.02 |
| | | | 10509.40 | 10923.05 | 10092.46 | 780.17 |
| | | | 6835.67 | 43044.61 | 11587.75 | 889.15 |
| | | | 29304.80 | 11217.86 | 10965.35 | 1129.48 |
| | | | 46487.00 | 2747.14 | 20611.92 | 353.95 |
| | | | 54873.60 | 23563.31 | 19624.02 | 5948.09 |
| | | | 60379.60 | 13103.22 | 5620.81 | 478.10 |
| | | | 49293.70 | 40277.37 | 14444.63 | 221.96 |
| | | | 75291.20 | 18004.20 | 7072.63 | 726.09 |
| | | | 24794.20 | 8012.52 | 2420.44 | 154.55 |
| | | | 14161.20 | 7546.92 | 2763.34 | 370.53 |
| | | | 22722.50 | 6309.24 | 29752.41 | 277.02 |
| | | | 14834.30 | 5126.84 | 12064.98 | 3.67 |
| | | | 3001.94 | 5825.49 | 13242.74 | 843.58 |
| | | | 9552.40 | 17746.57 | 27691.98 | 155.06 |
| | | | 10360.30 | 6164.58 | 23411.28 | 0.46 |
| | | | 1758.75 | 10581.29 | 19227.24 | 409.98 |
| | | | 19601.60 | 9124.97 | 13867.56 | 1716.90 |
| | | | 9961.80 | 318.89 | 15096.55 | 816.09 |
| | | | 11851.20 | 36817.10 | 17892.20 | 612.02 |
| | | | 15299.90 | 14732.98 | 6754.42 | 306.82 |
| | | | 16405.60 | 14717.62 | 3727.50 | 927.12 |
| | | | 14501.80 | 6785.68 | 22448.44 | 174.63 |
| | | | 47769.20 | 35430.46 | 18903.18 | 83.78 |
| | | | 30654.40 | 33727.03 | 20893.27 | 1519.62 |
| | | | 76074.80 | 41891.09 | 58931.63 | 16.86 |
| | | | 18632.80 | 6446.03 | 27474.65 | 499.88 |
| | | | 13789.20 | 20561.65 | 16439.96 | 840.95 |
| | | | 12223.00 | 6170.86 | 21015.51 | 894.66 |
| | | | | | 9833.89 | 480.77 |
| | | | | | 11136.96 | 923.17 |
| | | | | | 451.93 | 269.75 |
| | | | | | 6002.11 | 139.62 |
| | | | | | 366.62 | 365.03 |
| | | | | | 614.02 | 5417.13 |
| | | | | | 5916.82 | 3500.56 |
| | | | | | 6957.27 | 855.05 |
| | | | | | 106088.60 | 2823.98 |
| | | | | | 32822.41 | 971.02 |
| | | | | | 31450.18 | 3824.00 |
| | | | | | 77235.99 | 4421.96 |
| | | | | | 19970.80 | 830.84 |
| | | | | | 8408.79 | 1206.95 |
| | | | | | 18288.27 | 1425.20 |
| | | | | | 12971.70 | 2056.01 |
| | | | | | 19444.32 | 1575.57 |
| | | | | | 20214.31 | 20.03 |
| | | | | | 4914.36 | 365.64 |
| | | | | | 4038.36 | 1151.21 |
| | | | | | 21631.15 | 62.40 |
| | | | | | 5663.76 | |
| n | 23 | 25 | 75 | 75 | 97 | 96 |
| Mean | 646579.27 | 381846.22 | 20162.13 | 15959.82 | 14612.62 | 1143.04 |
| Median | 621753.18 | 287358.68 | 14834.30 | 10581.29 | 9833.89 | 788.17 |
| Sd | 443461.97 | 360420.49 | 18380.69 | 14901.49 | 17067.35 | 1287.33 |
| SEM | 92468.21 | 72084.10 | 2122.42 | 1720.68 | 1732.93 | 131.39 |

**Supplementary Tables S17:** Normalised fluorescence intensity (Relative Fluorescence Units; RFU) quantification of whole-embryo KRT8 levels in control siNTC and si*Tead4*–microinjected blastocysts, including expression in outer-cells, for unmarked/marked cell clones. Relates to data in Fig. 5.

### Supplementary Tables 18a: +48h (Clonal *Krt8* KD - E3.5)

#### siNTC

| Embryo Number | Total | Outer | Inner | Marked | Unmarked | Marked | | Unmarked | |
|---|---|---|---|---|---|---|---|---|---|
| | | | | | | Outer | Inner | Outer | Inner |
| _001 | 32 | 24 | 8 | 15 | 17 | 12 | 3 | 12 | 5 |
| _002 | 36 | 20 | 16 | 16 | 20 | 8 | 8 | 12 | 8 |
| _003 | 32 | 21 | 11 | 16 | 16 | 11 | 5 | 10 | 6 |
| _004 | 31 | 16 | 15 | 15 | 16 | 8 | 7 | 8 | 8 |
| _005 | 33 | 20 | 13 | 16 | 17 | 9 | 7 | 11 | 6 |
| _006 | 28 | 17 | 11 | 16 | 12 | 10 | 6 | 7 | 5 |
| _007 | 30 | 20 | 10 | 15 | 15 | 10 | 5 | 10 | 5 |
| _008 | 32 | 17 | 15 | 16 | 16 | 9 | 7 | 8 | 8 |
| _009 | 34 | 20 | 14 | 17 | 17 | 9 | 8 | 11 | 6 |
| _010 | 31 | 17 | 14 | 15 | 16 | 8 | 7 | 9 | 7 |
| _011 | 34 | 19 | 15 | 16 | 18 | 11 | 5 | 8 | 10 |
| _012 | 32 | 20 | 12 | 17 | 15 | 10 | 7 | 10 | 5 |
| _013 | 32 | 17 | 15 | 16 | 16 | 9 | 7 | 8 | 8 |
| _014 | 29 | 19 | 10 | 13 | 16 | 11 | 2 | 8 | 8 |
| _015 | 28 | 15 | 13 | 14 | 14 | 8 | 6 | 7 | 7 |
| _016 | 45 | 22 | 23 | 16 | 29 | 11 | 5 | 11 | 18 |
| _017 | 55 | 29 | 26 | 21 | 34 | 13 | 8 | 16 | 18 |
| _018 | 30 | 19 | 11 | 13 | 17 | 9 | 4 | 10 | 7 |
| Mean | 33.56 | 19.56 | 14.00 | 15.72 | 17.83 | 9.78 | 5.94 | 9.78 | 8.06 |
| Median | 32.00 | 19.50 | 13.50 | 16.00 | 16.00 | 9.50 | 6.50 | 10.00 | 7.00 |
| Sd | 6.55 | 3.24 | 4.43 | 1.74 | 5.31 | 1.48 | 1.73 | 2.24 | 3.87 |
| SEM | 1.55 | 0.76 | 1.04 | 0.41 | 1.25 | 0.35 | 0.41 | 0.53 | 0.91 |

### si*K8*

| Embryo Number | Total | Outer | Inner | Marked | Unmarked | Marked | | Unmarked | |
|---|---|---|---|---|---|---|---|---|---|
| | | | | | | Outer | Inner | Outer | Inner |
| _001 | 32 | 18 | 14 | 15 | 17 | 9 | 6 | 9 | 8 |
| _002 | 34 | 17 | 17 | 16 | 18 | 9 | 7 | 8 | 10 |
| _003 | 31 | 17 | 14 | 15 | 16 | 9 | 6 | 8 | 8 |
| _004 | 30 | 14 | 16 | 16 | 14 | 8 | 8 | 6 | 8 |
| _005 | 29 | 14 | 15 | 14 | 15 | 6 | 8 | 8 | 7 |
| _006 | 31 | 18 | 13 | 12 | 19 | 7 | 5 | 11 | 8 |
| _007 | 29 | 19 | 10 | 12 | 17 | 7 | 5 | 12 | 5 |
| _008 | 28 | 17 | 11 | 13 | 15 | 7 | 6 | 10 | 5 |
| _009 | 35 | 22 | 13 | 15 | 20 | 8 | 7 | 14 | 6 |
| _010 | 34 | 20 | 14 | 15 | 19 | 9 | 6 | 11 | 8 |
| _011 | 35 | 23 | 12 | 16 | 19 | 9 | 7 | 14 | 5 |
| _012 | 33 | 17 | 16 | 16 | 17 | 10 | 6 | 7 | 10 |
| _013 | 30 | 21 | 9 | 16 | 14 | 10 | 6 | 11 | 3 |
| _014 | 31 | 20 | 11 | 16 | 15 | 12 | 4 | 8 | 7 |
| _015 | 32 | 19 | 13 | 16 | 16 | 9 | 7 | 10 | 6 |
| _016 | 32 | 18 | 14 | 16 | 16 | 11 | 5 | 7 | 9 |
| _017 | 33 | 16 | 17 | 17 | 16 | 8 | 9 | 8 | 8 |
| _018 | 45 | 27 | 18 | 22 | 23 | 13 | 9 | 14 | 9 |
| _019 | 32 | 17 | 15 | 15 | 17 | 8 | 7 | 9 | 8 |
| _020 | 39 | 24 | 15 | 21 | 18 | 14 | 7 | 10 | 8 |
| _021 | 41 | 21 | 20 | 18 | 23 | 9 | 9 | 12 | 11 |
| Mean | 33.14 | 19.00 | 14.14 | 15.81 | 17.33 | 9.14 | 6.67 | 9.86 | 7.48 |
| Median | 32.00 | 18.00 | 14.00 | 16.00 | 17.00 | 9.00 | 7.00 | 10.00 | 8.00 |
| Sd | 4.15 | 3.21 | 2.71 | 2.40 | 2.52 | 2.01 | 1.39 | 2.39 | 1.94 |
| SEM | 0.91 | 0.70 | 0.59 | 0.52 | 0.55 | 0.44 | 0.30 | 0.52 | 0.42 |

**Supplementary Tables S18a:** Blastocyst lineage/positional contributions from injected/marked control siNTC and si*K8* (targeting *Krt8* mRNA) cell clones, versus equivalent non-injected/unmarked clones, specifically at +48 h (E3.5) post-single 2-cell stage microinjection. Data include raw cell counts. Relates to data in Fig. S12.

### Supplementary Tables 18b: +48h (Clonal *Krt8* & *Krt18* KD - E3.5)

#### siNTC

| Embryo Number | Total | Outer | Inner | Marked | Unmarked | Marked Outer | Marked Inner | Unmarked Outer | Unmarked Inner |
|---|---|---|---|---|---|---|---|---|---|
| _001 | 31 | 20 | 11 | 16 | 15 | 12 | 4 | 8 | 7 |
| _002 | 31 | 21 | 10 | 16 | 15 | 10 | 6 | 11 | 4 |
| _003 | 32 | 18 | 14 | 16 | 16 | 8 | 8 | 10 | 6 |
| _004 | 31 | 18 | 13 | 14 | 17 | 9 | 5 | 9 | 8 |
| _005 | 32 | 19 | 13 | 16 | 16 | 8 | 8 | 11 | 5 |
| _006 | 32 | 19 | 13 | 16 | 16 | 11 | 5 | 8 | 8 |
| _007 | 25 | 16 | 9 | 16 | 9 | 10 | 6 | 6 | 3 |
| _008 | 37 | 24 | 13 | 12 | 25 | 12 | 0 | 12 | 13 |
| _009 | 36 | 20 | 16 | 23 | 13 | 11 | 12 | 9 | 4 |
| _010 | 37 | 22 | 15 | 19 | 18 | 11 | 8 | 11 | 7 |
| _011 | 38 | 24 | 14 | 15 | 23 | 13 | 2 | 11 | 12 |
| _012 | 32 | 19 | 13 | 16 | 16 | 10 | 6 | 9 | 7 |
| _013 | 30 | 19 | 11 | 13 | 17 | 8 | 5 | 11 | 6 |
| _014 | 31 | 15 | 16 | 15 | 16 | 7 | 8 | 8 | 8 |
| _015 | 36 | 24 | 12 | 16 | 20 | 12 | 4 | 12 | 8 |
| _016 | 33 | 21 | 12 | 16 | 17 | 10 | 6 | 11 | 6 |
| Mean | 32.75 | 19.94 | 12.81 | 15.94 | 16.81 | 10.13 | 5.81 | 9.81 | 7.00 |
| Median | 32.00 | 19.50 | 13.00 | 16.00 | 16.00 | 10.00 | 6.00 | 10.50 | 7.00 |
| Sd | 3.34 | 2.67 | 1.97 | 2.43 | 3.67 | 1.75 | 2.76 | 1.72 | 2.66 |
| SEM | 0.83 | 0.67 | 0.49 | 0.61 | 0.92 | 0.44 | 0.69 | 0.43 | 0.66 |

### si*K8* & si*K18*

| Embryo Number | Total | Outer | Inner | Marked | Unmarked | Marked Outer | Marked Inner | Unmarked Outer | Unmarked Inner |
|---|---|---|---|---|---|---|---|---|---|
| _001 | 33 | 17 | 16 | 16 | 17 | 9 | 7 | 8 | 9 |
| _002 | 31 | 18 | 13 | 15 | 16 | 9 | 6 | 9 | 7 |
| _003 | 36 | 22 | 14 | 18 | 18 | 11 | 7 | 11 | 7 |
| _004 | 42 | 27 | 15 | 20 | 22 | 13 | 7 | 14 | 8 |
| _005 | 28 | 19 | 9 | 14 | 14 | 9 | 5 | 10 | 4 |
| _006 | 33 | 23 | 10 | 17 | 16 | 13 | 4 | 10 | 6 |
| _007 | 31 | 19 | 12 | 16 | 15 | 10 | 6 | 9 | 6 |
| _008 | 31 | 20 | 11 | 16 | 15 | 9 | 7 | 11 | 4 |
| _009 | 30 | 18 | 12 | 15 | 15 | 9 | 6 | 9 | 6 |
| _010 | 32 | 21 | 11 | 16 | 16 | 11 | 5 | 10 | 6 |
| _011 | 32 | 21 | 11 | 16 | 16 | 10 | 6 | 11 | 5 |
| _012 | 38 | 19 | 19 | 17 | 21 | 8 | 9 | 11 | 10 |
| _013 | 41 | 34 | 7 | 22 | 19 | 19 | 3 | 15 | 4 |
| _014 | 30 | 25 | 5 | 14 | 16 | 13 | 1 | 12 | 4 |
| _015 | 49 | 32 | 17 | 26 | 23 | 17 | 9 | 15 | 8 |
| _016 | 30 | 21 | 9 | 15 | 15 | 12 | 3 | 9 | 6 |
| _017 | 29 | 18 | 11 | 16 | 13 | 9 | 7 | 9 | 4 |
| _018 | 28 | 19 | 9 | 16 | 12 | 11 | 5 | 8 | 4 |
| _019 | 31 | 19 | 12 | 16 | 15 | 9 | 7 | 10 | 5 |
| _020 | 39 | 24 | 15 | 17 | 22 | 12 | 5 | 12 | 10 |
| _021 | 32 | 20 | 12 | 16 | 16 | 11 | 5 | 9 | 7 |
| Mean | 33.62 | 21.71 | 11.90 | 16.86 | 16.76 | 11.14 | 5.71 | 10.57 | 6.19 |
| Median | 32.00 | 20.00 | 12.00 | 16.00 | 16.00 | 11.00 | 6.00 | 10.00 | 6.00 |
| Sd | 5.36 | 4.53 | 3.33 | 2.78 | 3.02 | 2.76 | 1.93 | 2.06 | 1.97 |
| SEM | 1.17 | 0.99 | 0.73 | 0.61 | 0.66 | 0.60 | 0.42 | 0.45 | 0.43 |

**Supplementary Tables S18b:** Blastocyst lineage/positional contributions from injected/marked control siNTC and si*K8* & *siK18* (co-targeting *Krt8* & *Krt18* mRNA) cell clones, versus equivalent non-injected/unmarked clones, specifically at +48 h (E3.5) post-single 2-cell stage microinjection. Data include raw cell counts. Relates to data in Fig. S12.

**Supplementary Tables 19: +48h (Clonal *Tead4* KD and *Krt8* /*Krt8* -HA & *Krt18* OE - E3.5)**

***K8/K8-HA* & *K18* mRNA +**

**siNTC**

| Embryo Number | Total | Outer | Inner | Marked | Unmarked | Marked Outer | Marked Inner | Unmarked Outer | Unmarked Inner |
|---|---|---|---|---|---|---|---|---|---|
| _001 | 32 | 20 | 12 | 16 | 16 | 9 | 7 | 11 | 5 |
| _002 | 33 | 18 | 15 | 17 | 16 | 9 | 8 | 9 | 7 |
| _003 | 35 | 21 | 14 | 19 | 16 | 12 | 7 | 9 | 7 |
| _004 | 34 | 21 | 13 | 16 | 18 | 10 | 6 | 11 | 7 |
| _005 | 30 | 17 | 13 | 16 | 14 | 9 | 7 | 8 | 6 |
| _006 | 47 | 33 | 14 | 26 | 21 | 19 | 7 | 14 | 7 |
| _007 | 32 | 22 | 10 | 16 | 16 | 12 | 4 | 10 | 6 |
| _008 | 32 | 18 | 14 | 16 | 16 | 10 | 6 | 8 | 8 |
| _009 | 33 | 17 | 16 | 16 | 17 | 8 | 8 | 9 | 8 |
| _010 | 31 | 17 | 14 | 16 | 15 | 8 | 8 | 9 | 6 |
| _011 | 33 | 18 | 15 | 15 | 18 | 7 | 8 | 11 | 7 |
| _012 | 31 | 17 | 14 | 15 | 16 | 8 | 7 | 9 | 7 |
| _013 | 32 | 17 | 15 | 17 | 15 | 9 | 8 | 8 | 7 |
| Mean | 33.46 | 19.69 | 13.77 | 17.00 | 16.46 | 10.00 | 7.00 | 9.69 | 6.77 |
| Median | 32.00 | 18.00 | 14.00 | 16.00 | 16.00 | 9.00 | 7.00 | 9.00 | 7.00 |
| Sd | 4.27 | 4.39 | 1.54 | 2.89 | 1.76 | 3.08 | 1.15 | 1.70 | 0.83 |
| SEM | 1.19 | 1.22 | 0.43 | 0.80 | 0.49 | 0.85 | 0.32 | 0.47 | 0.23 |

**si*Tead4***

| Embryo Number | Total | Outer | Inner | Marked | Unmarked | Marked Outer | Marked Inner | Unmarked Outer | Unmarked Inner |
|---|---|---|---|---|---|---|---|---|---|
| _001 | 31 | 17 | 14 | 15 | 16 | 7 | 8 | 10 | 6 |
| _002 | 31 | 21 | 10 | 13 | 18 | 7 | 6 | 14 | 4 |
| _003 | 35 | 20 | 15 | 16 | 19 | 8 | 8 | 12 | 7 |
| _004 | 34 | 17 | 17 | 15 | 19 | 4 | 11 | 13 | 6 |
| _005 | 34 | 18 | 16 | 16 | 18 | 9 | 7 | 9 | 9 |
| _006 | 31 | 15 | 16 | 17 | 14 | 7 | 10 | 8 | 6 |
| _007 | 31 | 13 | 18 | 15 | 16 | 5 | 10 | 8 | 8 |
| _008 | 35 | 25 | 10 | 21 | 14 | 13 | 8 | 12 | 2 |
| _009 | 31 | 17 | 14 | 15 | 16 | 7 | 8 | 10 | 6 |
| _010 | 31 | 16 | 15 | 15 | 16 | 5 | 10 | 11 | 5 |
| _011 | 35 | 19 | 16 | 16 | 19 | 7 | 9 | 12 | 7 |
| _012 | 33 | 22 | 11 | 14 | 19 | 9 | 5 | 13 | 6 |
| Mean | 32.67 | 18.33 | 14.33 | 15.67 | 17.00 | 7.33 | 8.33 | 11.00 | 6.00 |
| Median | 32.00 | 17.50 | 15.00 | 15.00 | 17.00 | 7.00 | 8.00 | 11.50 | 6.00 |
| Sd | 1.83 | 3.28 | 2.67 | 1.97 | 1.91 | 2.35 | 1.78 | 2.00 | 1.81 |
| SEM | 0.53 | 0.95 | 0.77 | 0.57 | 0.55 | 0.68 | 0.51 | 0.58 | 0.52 |

Legend (cell shading of Embryo Number):
- *K8* mRNA
- *K8* -HA mRNA

**Supplementary Tables S19:** Summary of blastocyst lineage/positional contributions from injected/marked control siNTC and si*Tead4* cell clones, in which *Krt8* or *Krt8*-HA (as indicated) plus *Krt18* recombinant mRNA was co-injected, compared with equivalent non-injected/unmarked clones, at +48 h (E3.5) post-single 2-cell stage blastomere microinjection. Relates to data in Figs. 5 and S13E&F.

**Supplementary Tables 20: +48h (Clonal *Tead4* , *Rnd1* & *Rnd3* KD - E3.5)**

**siNTC + ds*EGFP***

| Embryo Number | Total | Outer | Inner | Marked | Unmarked | Marked | | Unmarked | |
|---|---|---|---|---|---|---|---|---|---|
| | | | | | | Outer | Inner | Outer | Inner |
| _001 | 26 | 18 | 8 | 11 | 15 | 6 | 5 | 12 | 3 |
| _002 | 27 | 18 | 9 | 12 | 15 | 8 | 4 | 10 | 5 |
| _003 | 31 | 20 | 11 | 15 | 16 | 11 | 4 | 9 | 7 |
| _004 | 29 | 20 | 9 | 15 | 14 | 12 | 3 | 8 | 6 |
| _005 | 33 | 20 | 13 | 14 | 19 | 8 | 6 | 12 | 7 |
| _006 | 30 | 20 | 10 | 14 | 16 | 9 | 5 | 11 | 5 |
| _007 | 47 | 30 | 17 | 25 | 22 | 15 | 10 | 15 | 7 |
| _008 | 32 | 20 | 12 | 16 | 16 | 12 | 4 | 8 | 8 |
| _009 | 30 | 17 | 13 | 15 | 15 | 10 | 5 | 7 | 8 |
| _010 | 28 | 17 | 11 | 14 | 14 | 9 | 5 | 8 | 6 |
| _011 | 32 | 18 | 14 | 16 | 16 | 9 | 7 | 9 | 7 |
| _012 | 35 | 20 | 15 | 18 | 17 | 11 | 7 | 9 | 8 |
| _013 | 32 | 21 | 11 | 17 | 15 | 13 | 4 | 8 | 7 |
| _014 | 36 | 19 | 17 | 16 | 20 | 9 | 7 | 10 | 10 |
| _015 | 32 | 18 | 14 | 18 | 14 | 11 | 7 | 7 | 7 |
| _016 | 32 | 18 | 14 | 16 | 16 | 11 | 5 | 7 | 9 |
| _017 | 29 | 18 | 11 | 14 | 15 | 7 | 7 | 11 | 4 |
| _018 | 25 | 15 | 10 | 10 | 15 | 6 | 4 | 9 | 6 |
| _019 | 45 | 30 | 15 | 27 | 18 | 18 | 9 | 12 | 6 |
| _020 | 32 | 19 | 13 | 16 | 16 | 8 | 8 | 11 | 5 |
| _021 | 22 | 16 | 6 | 10 | 12 | 7 | 3 | 9 | 3 |
| _022 | 25 | 14 | 11 | 15 | 10 | 8 | 7 | 6 | 4 |
| _023 | 30 | 15 | 15 | 15 | 15 | 7 | 8 | 8 | 7 |
| _024 | 30 | 16 | 14 | 16 | 14 | 9 | 7 | 7 | 7 |
| _025 | 30 | 16 | 14 | 16 | 14 | 8 | 8 | 8 | 6 |
| _026 | 31 | 19 | 12 | 15 | 16 | 11 | 4 | 8 | 8 |
| _027 | 31 | 17 | 14 | 14 | 17 | 7 | 7 | 10 | 7 |
| _028 | 32 | 20 | 12 | 11 | 21 | 8 | 3 | 12 | 9 |
| _029 | 16 | 10 | 6 | 8 | 8 | 6 | 2 | 4 | 4 |
| _030 | 31 | 17 | 14 | 12 | 19 | 7 | 5 | 10 | 9 |
| _031 | 34 | 18 | 16 | 18 | 16 | 11 | 7 | 7 | 9 |
| _032 | 41 | 23 | 18 | 19 | 22 | 10 | 9 | 13 | 9 |
| _033 | 29 | 17 | 12 | 14 | 15 | 9 | 5 | 8 | 7 |
| _034 | 32 | 17 | 15 | 16 | 16 | 9 | 7 | 8 | 8 |
| _035 | 34 | 24 | 10 | 15 | 19 | 10 | 5 | 14 | 5 |
| _036 | 31 | 21 | 10 | 16 | 15 | 11 | 5 | 10 | 5 |
| _037 | 47 | 27 | 20 | 18 | 29 | 9 | 9 | 18 | 11 |
| _038 | 34 | 19 | 15 | 17 | 17 | 8 | 9 | 11 | 6 |
| Mean | 31.66 | 19.00 | 12.66 | 15.37 | 16.29 | 9.42 | 5.95 | 9.58 | 6.71 |
| Median | 31.00 | 18.00 | 13.00 | 15.00 | 16.00 | 9.00 | 5.50 | 9.00 | 7.00 |
| Sd | 5.97 | 3.88 | 3.09 | 3.54 | 3.52 | 2.49 | 2.01 | 2.68 | 1.89 |
| SEM | 0.97 | 0.63 | 0.50 | 0.57 | 0.57 | 0.40 | 0.33 | 0.43 | 0.31 |

**si*Tead4* + ds*Rnd1* + ds*Rnd3***

| Embryo Number | Total | Outer | Inner | Marked | Unmarked | Marked | | Unmarked | |
|---|---|---|---|---|---|---|---|---|---|
| | | | | | | Outer | Inner | Outer | Inner |
| _001 | 33 | 17 | 16 | 18 | 15 | 8 | 10 | 9 | 6 |
| _002 | 30 | 18 | 12 | 15 | 15 | 9 | 6 | 9 | 6 |
| _003 | 25 | 17 | 8 | 11 | 14 | 8 | 3 | 9 | 5 |
| _004 | 36 | 23 | 13 | 15 | 21 | 7 | 8 | 16 | 5 |
| _005 | 33 | 24 | 9 | 15 | 18 | 11 | 4 | 13 | 5 |
| _006 | 32 | 20 | 12 | 16 | 16 | 10 | 6 | 10 | 6 |
| _007 | 40 | 24 | 16 | 14 | 26 | 11 | 3 | 13 | 13 |
| _008 | 42 | 27 | 15 | 19 | 23 | 9 | 10 | 18 | 5 |
| _009 | 29 | 15 | 14 | 14 | 15 | 7 | 7 | 8 | 7 |
| _010 | 30 | 16 | 14 | 14 | 16 | 6 | 8 | 10 | 6 |
| _011 | 23 | 15 | 8 | 10 | 13 | 7 | 3 | 8 | 5 |
| _012 | 30 | 15 | 15 | 14 | 16 | 6 | 8 | 9 | 7 |
| _013 | 19 | 13 | 6 | 14 | 5 | 9 | 5 | 4 | 1 |
| _014 | 30 | 23 | 7 | 16 | 14 | 13 | 3 | 10 | 4 |
| _015 | 25 | 15 | 10 | 16 | 9 | 8 | 8 | 7 | 2 |
| _016 | 31 | 15 | 16 | 16 | 15 | 7 | 9 | 8 | 7 |
| _017 | 28 | 19 | 9 | 17 | 11 | 11 | 6 | 8 | 3 |
| _018 | 33 | 22 | 11 | 16 | 17 | 9 | 7 | 13 | 4 |
| _019 | 33 | 19 | 14 | 15 | 18 | 8 | 7 | 11 | 7 |
| _020 | 32 | 15 | 17 | 16 | 16 | 5 | 11 | 10 | 6 |
| _021 | 40 | 21 | 19 | 17 | 23 | 7 | 10 | 14 | 9 |
| _022 | 33 | 22 | 11 | 16 | 17 | 12 | 4 | 10 | 7 |
| _023 | 35 | 17 | 18 | 16 | 19 | 6 | 10 | 11 | 8 |
| _024 | 32 | 16 | 16 | 16 | 16 | 8 | 8 | 8 | 8 |
| _025 | 28 | 14 | 14 | 14 | 14 | 8 | 6 | 6 | 8 |
| _026 | 27 | 16 | 11 | 14 | 13 | 7 | 7 | 9 | 4 |
| _027 | 30 | 18 | 12 | 14 | 16 | 10 | 4 | 8 | 8 |
| _028 | 30 | 14 | 16 | 14 | 16 | 6 | 8 | 8 | 8 |
| _029 | 31 | 16 | 15 | 13 | 18 | 7 | 6 | 9 | 9 |
| _030 | 29 | 16 | 13 | 14 | 15 | 8 | 6 | 8 | 7 |
| _031 | 33 | 21 | 12 | 16 | 17 | 9 | 7 | 12 | 5 |
| _032 | 47 | 22 | 25 | 19 | 28 | 3 | 16 | 19 | 9 |
| _033 | 36 | 18 | 18 | 14 | 22 | 8 | 6 | 10 | 12 |
| _034 | 30 | 19 | 11 | 15 | 15 | 10 | 5 | 9 | 6 |
| _035 | 29 | 16 | 13 | 12 | 17 | 7 | 5 | 9 | 8 |
| _036 | 28 | 15 | 13 | 14 | 14 | 9 | 5 | 6 | 8 |
| _037 | 34 | 22 | 12 | 15 | 19 | 10 | 5 | 12 | 7 |
| _038 | 32 | 18 | 14 | 16 | 16 | 8 | 8 | 10 | 6 |
| _039 | 32 | 18 | 14 | 16 | 16 | 9 | 7 | 9 | 7 |
| _040 | 33 | 18 | 15 | 13 | 20 | 5 | 8 | 13 | 7 |
| _041 | 35 | 21 | 14 | 14 | 21 | 8 | 6 | 13 | 8 |
| _042 | 32 | 16 | 16 | 13 | 19 | 7 | 6 | 9 | 10 |
| _043 | 35 | 22 | 13 | 16 | 19 | 9 | 7 | 13 | 6 |
| _044 | 30 | 15 | 15 | 16 | 14 | 10 | 6 | 5 | 9 |
| Mean | 31.70 | 18.25 | 13.45 | 14.95 | 16.75 | 8.18 | 6.77 | 10.07 | 6.68 |
| Median | 32.00 | 18.00 | 14.00 | 15.00 | 16.00 | 8.00 | 6.50 | 9.00 | 7.00 |
| Sd | 4.83 | 3.32 | 3.46 | 1.79 | 4.06 | 1.94 | 2.48 | 3.07 | 2.29 |
| SEM | 0.73 | 0.50 | 0.52 | 0.27 | 0.61 | 0.29 | 0.37 | 0.46 | 0.35 |

**Supplementary Tables S20:** Summary of blastocyst lineage/positional contributions from injected/marked control siNTC/dsEGFP and si*Tead4* plus ds*Rnd1* & ds*Rnd3* cell clones, compared with equivalent non-injected/unmarked clones, at +48 h (E3.5) post-single 2-cell stage blastomere microinjection. Relates to data in Figs. 5 and S14A.

**Supplementary Tables 21a: E3.75 (+/- Jasplakinolide; PARD6B & TEAD4 IF)**

| DMSO | | | | Jasplakinolide | | | |
|---|---|---|---|---|---|---|---|
| Embryo Number | Total | Outer | Inner | Embryo Number | Total | Outer | Inner |
| _001 | 58 | 37 | 21 | _001 | 44 | 27 | 17 |
| _002 | 31 | 19 | 12 | _002 | 32 | 19 | 13 |
| _003 | 61 | 30 | 31 | _003 | 39 | 27 | 12 |
| _004 | 38 | 27 | 11 | _004 | 29 | 18 | 11 |
| _005 | 32 | 21 | 11 | _005 | 29 | 19 | 10 |
| _006 | 30 | 17 | 13 | _006 | 43 | 25 | 18 |
| _007 | 42 | 27 | 15 | _007 | 43 | 28 | 15 |
| _008 | 60 | 32 | 28 | _008 | 45 | 33 | 12 |
| _009 | 57 | 35 | 22 | _009 | 64 | 30 | 34 |
| _010 | 35 | 20 | 15 | _010 | 34 | 20 | 14 |
| _011 | 43 | 28 | 15 | _011 | 47 | 25 | 22 |
| _012 | 30 | 15 | 15 | _012 | 32 | 16 | 16 |
| _013 | 47 | 31 | 16 | _013 | 37 | 24 | 13 |
| _014 | 52 | 29 | 23 | _014 | 42 | 26 | 16 |
| _015 | 59 | 33 | 26 | _015 | 32 | 16 | 16 |
| _016 | 31 | 22 | 9 | _016 | 62 | 38 | 24 |
| _017 | 32 | 18 | 14 | _017 | 45 | 30 | 15 |
| _018 | 38 | 23 | 15 | _018 | 29 | 18 | 11 |
| _019 | 28 | 15 | 13 | _019 | 35 | 21 | 14 |
| _020 | 27 | 17 | 10 | | | | |
| Mean | 41.55 | 24.80 | 16.75 | Mean | 40.16 | 24.21 | 15.95 |
| Median | 38.00 | 25.00 | 15.00 | Median | 39.00 | 25.00 | 15.00 |
| Sd | 12.16 | 6.95 | 6.26 | Sd | 10.06 | 6.07 | 5.65 |
| SEM | 2.72 | 1.55 | 1.40 | SEM | 2.31 | 1.39 | 1.30 |

**Supplementary Tables S21a:** Total outer and inner cell numbers in blastocysts treated with DMSO vehicle control or Jasplakinolide for 12 hours from the E3.25 stage (IF-stained for TEAD4 and PARD6B). Relates to data in Figs. S15.

**Supplementary Tables 21b: E3.75 (+/- Jasplakinolide; YAP1 IF & phalloidin stain)**

**DMSO**

| Embryo Number | Total | Outer | Inner |
|---|---|---|---|
| _001 | 39 | 22 | 17 |
| _002 | 45 | 30 | 15 |
| _003 | 59 | 37 | 22 |
| _004 | 47 | 24 | 23 |
| _005 | 39 | 22 | 17 |
| _006 | 41 | 29 | 12 |
| _007 | 34 | 20 | 14 |
| _008 | 52 | 25 | 27 |
| _009 | 62 | 37 | 25 |
| _010 | 55 | 41 | 14 |
| _011 | 40 | 24 | 16 |
| _012 | 48 | 30 | 18 |
| _013 | 48 | 25 | 23 |
| _014 | 35 | 21 | 14 |
| _015 | 32 | 18 | 14 |
| _016 | 42 | 28 | 14 |
| _017 | 34 | 21 | 13 |
| _018 | 35 | 22 | 13 |
| _019 | 57 | 39 | 18 |
| _020 | 40 | 27 | 13 |
| _021 | 57 | 32 | 25 |
| _022 | 55 | 35 | 20 |
| _023 | 55 | 34 | 21 |
| _024 | 38 | 23 | 15 |
| Mean | 45.38 | 27.75 | 17.63 |
| Median | 43.50 | 26.00 | 16.50 |
| Sd | 9.20 | 6.62 | 4.52 |
| SEM | 1.88 | 1.35 | 0.92 |

**Jasplakinolide**

| Embryo Number | Total | Outer | Inner |
|---|---|---|---|
| _001 | 53 | 29 | 24 |
| _002 | 30 | 18 | 12 |
| _003 | 38 | 25 | 13 |
| _004 | 57 | 33 | 24 |
| _005 | 40 | 27 | 13 |
| _006 | 54 | 34 | 20 |
| _007 | 56 | 27 | 29 |
| _008 | 34 | 17 | 17 |
| _009 | 32 | 25 | 7 |
| _010 | 36 | 21 | 15 |
| _011 | 36 | 18 | 18 |
| _012 | 34 | 20 | 14 |
| _013 | 53 | 35 | 18 |
| _014 | 27 | 16 | 11 |
| _015 | 30 | 15 | 15 |
| _016 | 51 | 29 | 22 |
| _017 | 43 | 27 | 16 |
| _018 | 33 | 19 | 14 |
| _019 | 54 | 30 | 24 |
| _020 | 54 | 29 | 25 |
| _021 | 61 | 31 | 30 |
| _022 | 42 | 30 | 12 |
| _023 | 38 | 24 | 14 |
| _024 | 46 | 27 | 19 |
| _025 | 43 | 23 | 20 |
| _026 | 36 | 24 | 12 |
| Mean | 42.73 | 25.12 | 17.62 |
| Median | 41.00 | 26.00 | 16.50 |
| Sd | 10.01 | 5.73 | 5.80 |
| SEM | 1.96 | 1.12 | 1.14 |

**Supplementary Tables S21b:** Total outer and inner cell numbers in blastocysts treated with DMSO vehicle control or Jasplakinolide for 12 hours from the E3.25 stage (IF-stained for YAP1 and counterstained with phalloidin for actin). Relates to data in Figs. S15.

| Supplementary Tables 21c: E3.75 (+/- Jasplakinolide; PARD6B & TEAD4 normalised fluorescence intensity - Relative Fluorescence Unitis/RFU) | | | | | |
|---|---|---|---|---|---|
| | Whole embryo PARD6B (RFU) | | | Whole embryo TEAD4 (RFU) | |
| | DMSO | Jasp | | DMSO | Jasp |
| | 12519657.15 | 6101019.14 | | 7154414.11 | 3195004.69 |
| | 5848487.15 | 8617862.77 | | 4632287.14 | 5012680.56 |
| | 10365011.81 | 5470472.43 | | 6699858.51 | 3793977.57 |
| | 6058491.26 | 10812122.18 | | 4232055.89 | 4212980.12 |
| | 4904928.83 | 9881065.73 | | 3470849.97 | 5439256.08 |
| | 8267776.08 | 7020163.94 | | 5459878.29 | 3756307.47 |
| | 7648952.60 | 8171306.24 | | 4960424.77 | 3871986.25 |
| | 8749428.09 | 9135125.76 | | 4573072.87 | 4463946.86 |
| | 9737528.84 | 7166130.12 | | 5368310.48 | 4251616.39 |
| | 7829795.42 | 5787942.28 | | 4610411.31 | 3345598.82 |
| | 6703241.03 | 7608093.39 | | 3946226.86 | 4566882.66 |
| | 5250088.95 | 6350683.45 | | 3997116.93 | 3202990.76 |
| | 9646454.37 | 5586883.79 | | 4800300.12 | 3655286.87 |
| | 5932248.93 | 5798080.46 | | 4137053.15 | 3441752.64 |
| | 6283668.04 | 6962855.90 | | 3750542.55 | 4233863.98 |
| | 8022061.81 | 9043190.48 | | 4772644.90 | 5114425.59 |
| | 9236430.43 | 12471594.82 | | 5452658.19 | 5153314.19 |
| | 5110707.26 | 6940010.78 | | 2492217.51 | 4028478.30 |
| | 4647218.64 | 6012146.34 | | 2928697.71 | 3412748.82 |
| | 3306234.25 | | | 2596350.24 | |
| n | 20 | 19 | | 20 | 19 |
| Mean | 7303420.55 | 7628250.00 | | 4501768.57 | 4113320.98 |
| Median | 7176096.82 | 7020163.94 | | 4591742.09 | 4028478.30 |
| Sd | 2299018.61 | 1941551.09 | | 1200605.84 | 696454.36 |
| SEM | 514076.19 | 445422.37 | | 268463.63 | 159777.59 |

**Supplementary Tables S21c:** Normalised fluorescence intensity (Relative Fluorescence Units; RFU) quantification of whole embryo PARD6B and TEAD4 levels in Jasplakinolide- or vehicle DMSO-treated control E3.75 blastocysts (i.e. 12 hours from the E3.25 stage). Relates to data in Figs. S15. *Statistical details are provided in supplementary statistics Excel workbook.*

**Supplementary Tables 21d: E3.75 (+/- Jasplakinolide; YAP1 normalised fluorescence intensity - Relative Fluorescence Unitis/RFU)**

| | Whole embryo | | Individual outer cells* – Nuclear YAP1 | | | | | |
|---|---|---|---|---|---|---|---|---|
| | | | Mural | | Polar | | Boundary | |
| | DMSO | Jasp | DMSO | Jasp | DMSO | Jasp | DMSO | Jasp |
| | 12519657.15 | 6101019.14 | 179533.72 | 103123.95 | 120233.38 | 89188.39 | 224874.51 | 43046.00 |
| | 5848487.15 | 8617862.77 | 181671.95 | 116168.51 | 125473.93 | 116074.83 | 137336.40 | 80766.50 |
| | 10365011.81 | 5470472.43 | 117519.68 | 122897.18 | 113060.93 | 141364.71 | 154610.85 | 144755.34 |
| | 6058491.26 | 10812122.18 | 135184.72 | 111535.54 | 99505.18 | 103904.49 | 171530.81 | 150548.62 |
| | 4904928.83 | 9881065.73 | 28502.21 | 180176.47 | 139284.08 | 68431.62 | 175118.84 | 129715.69 |
| | 8267776.08 | 7020163.94 | 252093.24 | 156553.89 | 173330.13 | 109175.76 | 239640.08 | 142183.62 |
| | 7648952.60 | 8171306.24 | 252761.25 | 102030.87 | 310412.34 | 61787.58 | 237465.94 | 72150.99 |
| | 8749428.09 | 9135125.76 | 123990.84 | 115308.01 | 231367.04 | 123742.88 | 250884.21 | 115456.25 |
| | 9737528.84 | 7166130.12 | 191849.48 | 77335.07 | 215657.52 | 137585.69 | 135677.71 | 72873.14 |
| | 7829795.42 | 5787942.28 | 119319.02 | 93326.32 | 125521.34 | 123431.82 | 113440.43 | 100505.14 |
| | 6703241.03 | 7608093.39 | 171436.21 | 100171.19 | 115498.12 | 142382.14 | 154740.29 | 92208.58 |
| | 5250088.95 | 6350683.45 | 32113.11 | 182350.91 | | 85638.68 | 161510.20 | 105005.56 |
| | 9646454.37 | 5586883.79 | 42452.14 | 101145.44 | | 48803.65 | 144637.53 | 167565.68 |
| | 5932248.93 | 5798080.46 | | 90018.21 | | 123359.18 | | 73924.09 |
| | 6283668.04 | 6962855.90 | | 80678.71 | | 147458.88 | | 99318.69 |
| | 8022061.81 | 9043190.48 | | 111217.21 | | 62083.56 | | 79767.45 |
| | 9236430.43 | 12471594.82 | | 167383.91 | | 33675.30 | | 170246.92 |
| | 5110707.26 | 6940010.78 | | 184640.63 | | 92326.93 | | 89863.87 |
| | 4647218.64 | 6012146.34 | | 116585.79 | | 82270.01 | | 132960.78 |
| | 3306234.25 | | | | | | | |
| n | 20 | 19 | 13 | 19 | 11 | 19 | 13 | 19 |
| Mean | 7303420.55 | 7628250.00 | 140648.27 | 121718.31 | 160849.45 | 99615.06 | 177035.98 | 108571.73 |
| Median | 7176096.82 | 7020163.94 | 135184.72 | 111535.54 | 125521.34 | 103904.49 | 161510.20 | 100505.14 |
| Sd | 2299018.61 | 1941551.09 | 74749.62 | 34762.15 | 65741.40 | 33990.66 | 45615.03 | 35787.47 |
| SEM | 514076.19 | 445422.37 | 20731.81 | 7974.98 | 19821.78 | 7797.99 | 12651.33 | 8210.21 |

| | Individual outer cells* – Cytoplasmic YAP1 | | | | | |
|---|---|---|---|---|---|---|
| | Mural | | Polar | | Boundary | |
| | DMSO | Jasp | DMSO | Jasp | DMSO | Jasp |
| | 83146.54 | 66835.96 | 56658.23 | 48372.08 | 150289.00 | 180557.89 |
| | 132869.29 | 83056.17 | 84543.03 | 182466.62 | 112775.92 | 122276.55 |
| | 54824.27 | 42965.88 | 108506.96 | 211595.50 | 40680.99 | 113741.09 |
| | 43127.99 | 101387.55 | 102161.04 | 96528.14 | 94492.06 | 123292.04 |
| | 56466.48 | 77786.81 | 178470.59 | 100707.39 | 171732.20 | 183299.04 |
| | 152486.37 | 55500.98 | 94029.46 | 116166.84 | 179616.56 | 100762.79 |
| | 143855.29 | 75921.18 | 350162.25 | 61789.04 | 110600.51 | 36778.56 |
| | 62643.54 | 60103.86 | 135714.12 | 92003.83 | 171709.96 | 120620.37 |
| | 124630.93 | 32678.56 | 191138.96 | 109084.68 | 146008.69 | 82858.35 |
| | 56905.49 | 30385.09 | 122999.15 | 109954.65 | 88696.73 | 71089.25 |
| | 153286.80 | 44647.82 | 89956.01 | 113326.42 | 158760.19 | 36562.45 |
| | 45389.22 | 79893.42 | | 80036.38 | 255217.59 | 74488.77 |
| | 76611.25 | 79307.13 | | 59344.49 | 166582.99 | 134290.23 |
| | | 52982.16 | | 86356.00 | | 74161.45 |
| | | 48233.50 | | 78318.75 | | 67128.04 |
| | | 95431.08 | | 126733.25 | | 136275.42 |
| | | 90624.48 | | 60371.56 | | 98225.41 |
| | | 148683.70 | | 59061.89 | | 201503.05 |
| | | 72953.35 | | 86186.70 | | 153218.11 |
| n | 13 | 19 | 11 | 19 | 13 | 19 |
| Mean | 91249.50 | 70493.61 | 137667.25 | 98863.38 | 142089.49 | 111112.05 |
| Median | 76611.25 | 72953.35 | 108506.96 | 92003.83 | 150289.00 | 113741.09 |
| Sd | 43249.01 | 28008.46 | 80977.47 | 41440.60 | 53364.76 | 47090.61 |
| SEM | 11995.12 | 6425.58 | 24415.63 | 9507.13 | 14800.72 | 10803.33 |

| | Individual inner cells* – Nuclear YAP | | Individual inner cells* – Cytoplasmic YAP1 | |
|---|---|---|---|---|
| | DMSO | Jasp | DMSO | Jasp |
| | 67340.60 | 40493.63 | 125096.00 | 96500.84 |
| | 61542.91 | 49056.37 | 143607.17 | 100177.41 |
| | 67942.21 | 45285.83 | 127446.32 | 111364.59 |
| | 69554.54 | 55985.56 | 119962.96 | 218730.78 |
| | 66142.48 | 52346.04 | 121680.38 | 177927.68 |
| | 51935.13 | 44549.24 | 110087.19 | 158303.31 |
| | 43940.95 | 36724.94 | 145553.40 | 110984.32 |
| | 81697.98 | 62087.99 | 168974.56 | 187147.37 |
| | 66799.08 | 50791.72 | 143250.51 | 93318.99 |
| | 68760.77 | 39425.14 | 134163.91 | 80539.62 |
| | 68672.04 | 30047.42 | 187422.47 | 61554.30 |
| | 114265.39 | 35594.75 | 311279.25 | 124351.39 |
| | 81083.42 | 51636.58 | 424779.23 | 118099.76 |
| | 81235.24 | 57935.07 | 204696.42 | 139429.22 |
| | 68232.88 | 41752.51 | 202271.62 | 84985.93 |
| | 42874.26 | 44596.15 | 72687.37 | 59306.57 |
| | 58576.36 | 53676.92 | 149375.86 | 111295.75 |
| | 43236.31 | 30040.06 | 127434.43 | 62606.97 |
| | 63603.73 | 46434.91 | 200418.54 | 109627.83 |
| | 58640.04 | 56844.81 | 122763.16 | 131860.02 |
| | 46588.15 | 41925.35 | 133835.21 | 104952.77 |
| | | 40606.31 | | 89686.76 |
| | | 45906.20 | | 133502.59 |
| | | 48804.16 | | 105413.59 |
| | | 53822.51 | | 68868.06 |
| | | 26701.76 | | 58738.51 |
| | | 46363.24 | | 68852.97 |
| | | 36128.65 | | 96373.59 |
| | | 46240.59 | | 150964.28 |
| | | 36137.61 | | 108221.58 |
| n | 21 | 30 | 21 | 30 |
| Mean | 65364.97 | 44931.40 | 165561.24 | 110789.58 |
| Median | 66799.08 | 45596.01 | 143250.51 | 106817.58 |
| Sd | 16301.62 | 8768.20 | 77048.38 | 39140.55 |
| SEM | 3557.30 | 1600.85 | 16813.33 | 7146.05 |

| | Nuclear:Cytoplasmic YAP1** – Outer cells | | | | | | Nuclear:Cytoplasmic YAP1** – Inner cells | | Jasp treatment Nuc:Cyto (RFU) – Outer cells | |
|---|---|---|---|---|---|---|---|---|---|---|
| | Mural | | Polar | | Boundary | | | | | |
| | DMSO | Jasp | DMSO | Jasp | DMSO | Jasp | DMSO | Jasp | Typical | Atypical |
| | 2.29 | 1.55 | 2.16 | 1.88 | 1.50 | 0.24 | 0.54 | 0.42 | 1.88 | 0.24 |
| | 1.41 | 1.38 | 1.55 | 0.64 | 1.27 | 0.66 | 0.44 | 0.50 | 0.64 | 0.66 |
| | 2.20 | 2.89 | 1.07 | 0.67 | 3.83 | 1.33 | 0.54 | 0.41 | 0.67 | 0.68 |
| | 3.25 | 1.12 | 1.04 | 1.10 | 1.85 | 1.23 | 0.59 | 0.26 | 1.10 | 0.72 |
| | 0.52 | 2.32 | 0.79 | 0.68 | 1.04 | 0.72 | 0.56 | 0.30 | 1.33 | 0.97 |
| | 1.66 | 2.86 | 2.70 | 0.97 | 1.33 | 1.48 | 0.49 | 0.29 | 1.23 | 0.83 |
| | 1.76 | 1.37 | 1.26 | 1.04 | 2.15 | 1.99 | 0.30 | 0.34 | 1.48 | 1.04 |
| | 1.98 | 1.95 | 1.71 | 1.41 | 1.50 | 0.97 | 0.51 | 0.34 | 1.04 | 0.52 |
| | 1.54 | 2.66 | 1.13 | 1.26 | 0.94 | 0.88 | 0.51 | 0.57 | 1.41 | 1.50 |
| | 2.10 | 3.08 | 1.02 | 1.13 | 1.28 | 1.43 | 0.56 | 0.48 | 1.99 | 0.59 |
| | 1.12 | 2.24 | 1.28 | 1.27 | 0.99 | 2.58 | 0.49 | 0.51 | 0.97 | |
| | 0.75 | 2.31 | | 1.11 | 0.66 | 1.49 | 0.37 | 0.29 | 1.26 | |
| | 0.53 | 1.27 | | 0.83 | 0.88 | 1.27 | 0.19 | 0.44 | 1.13 | |
| | | 1.74 | | 1.43 | | 1.04 | 0.40 | 0.42 | 0.88 | |
| | | 1.67 | | 1.90 | | 1.50 | 0.35 | 0.49 | 1.43 | |
| | | 1.21 | | 0.52 | | 0.59 | 0.59 | 0.86 | 1.27 | |
| | | 1.86 | | 0.57 | | 1.79 | 0.39 | 0.50 | 1.11 | |
| | | 1.24 | | 1.80 | | 0.44 | 0.34 | 0.49 | 2.58 | |
| | | 1.61 | | 0.95 | | 0.87 | 0.32 | 0.43 | 1.49 | |
| | | | | | | | 0.48 | 0.43 | 1.43 | |
| | | | | | | | 0.36 | 0.41 | 1.27 | |
| | | | | | | | | 0.45 | 1.90 | |
| | | | | | | | | 0.36 | 0.57 | |
| | | | | | | | | 0.47 | 1.80 | |
| | | | | | | | | 0.79 | 1.79 | |
| | | | | | | | | 0.46 | 0.57 | |
| | | | | | | | | 0.68 | 0.44 | |
| | | | | | | | | 0.38 | 0.87 | |
| | | | | | | | | 0.31 | | |
| | | | | | | | | 0.34 | | |
| n | 13 | 19 | 11 | 19 | 13 | 19 | 21 | 30 | 28 | 10 |
| Mean | 1.62 | 1.91 | 1.43 | 1.11 | 1.48 | 1.18 | 0.44 | 0.45 | 1.27 | 0.77 |
| Median | 1.66 | 1.74 | 1.26 | 1.10 | 1.28 | 1.23 | 0.48 | 0.43 | 1.26 | 0.70 |
| Sd | 0.78 | 0.63 | 0.57 | 0.43 | 0.82 | 0.57 | 0.11 | 0.14 | 0.50 | 0.34 |
| SEM | 0.22 | 0.14 | 0.17 | 0.10 | 0.23 | 0.13 | 0.02 | 0.03 | 0.09 | 0.11 |

*All individual readings represent an average of three consecutive, central slices.

**Ratios represent an average of the three consecuative slice ratios (i.e. not the ratio of the average slices)

**Supplementary Tables S21d:** Normalised fluorescence intensity (Relative Fluorescence Units; RFU) quantification of single cell cytoplasmic and nuclear YAP1 levels in Jasplakinolide- or vehicle DMSO-treated control E3.75 blastocysts, in inner and outer positions (delineated by position in the mural or polar TE or at the boundary). Whole embryo YAP1 expression levels are also shown (left) for comparison. Also shown are the associated single cell Nuclear:Cytoplasmic (Nuc:Cyto) expression ratios. For the Jasplakinolide-treated group, the Nuc:Cyto YAP1 expression ratios are also given for outer cells with typical morphology or atypical outer cells with apical-blebs. Relates to data in Figs. S15. *Statistical details are provided in supplementary statistics Excel workbook.*

### Supplementary Table 22: Microinjected construct summary

**siRNAs**

| Targeted mouse gene/mRNA | Construct abbreviation | Company supplier | Cat. No. | Injected concentration | Resuspension media |
|---|---|---|---|---|---|
| *Tead4* | si*Tead4* | ThermoFisher - murine Silencer-Select gene siRNAs | 4390771 - s74939 | 10µM | HPLC water |
| *Krt8* | si*K8* | ThermoFisher - murine Silencer-Select gene siRNAs | 4390771 - s201576 | 5µM | HPLC water |
| *Krt18* | si*K8* | ThermoFisher - murine Silencer-Select gene siRNAs | 4390771 - s68965 | 5µM | HPLC water |
| non-targeting negatve control | siNTC | Qiagen - All Stars murine negative control | 1027292 | 10µM | HPLC water |

**dsRNAs**

| Targeted mouse gene/mRNA | Construct abbreviation | T7-linked sense template PCR primer | T7-linked anti-sense template PCR primer | Injected concentration | Resuspension media |
|---|---|---|---|---|---|
| *Tead4* | *dsTead4* | taatacgactcactatagggTGTTGGAGTTCTCGGCTTTC | taatacgactcactatagggTCGGTAGATGTGGTGCTGAG | 100ng/µl | HPLC water |
| *Tfap2c* | ds*Tfap2c* | taatacgactcactatagggGACCTGCTGCTGCCTCAC | taatacgactcactatagggTCATCGAAATGGGTCCTTTG | 150ng/µl | HPLC water |
| *Rnd1* | dsRnd1 | taatacgactcactatagggGTCCTGGTGGGAGACGTG | taatacgactcactatagggAGAACCTGAGGTGTCCCAGA | 100ng/µl | HPLC water |
| *Rnd3* | dsRnd3 | taatacgactcactatagggACTATGACAACGTCCGTCCA | taatacgactcactatagggAGCCCACCAACAGCATCTT | 100ng/µl | HPLC water |
| *EFGP - negative control* | dsEGFP | taatacgactcactatagggAGAGTACAAATTTTCTGTCAGTGGAGAGG | taatacgactcactatagggAGATGTATAGTTCATCCATGCCATGTGTA | Accordingly | HPLC water |

**sgRNA**

| Targeted mouse gene/mRNA | Construct abbreviation | Company Source | Cat. No. | Injected concentration | Resuspension media |
|---|---|---|---|---|---|
| *Cdx2* | sg*Cdx2* | ThermoFisher - murine TrueGuide Synthetic sgRNA | A35533 - CRISPR52050_SGM | 100ng/µl | HPLC water |

**Recombinant mRNAs**

| Gene/mRNA (*mouse*) name | Construct abbreviation | Restriction site- & KOZAK-seq- linked sense cloning PCR primer (in-house cloning) | Restriction site-linked anti-sense cloning PCR primer (in-house cloning) | Plasmid vector | Plasmid name | IVT promoter utilised | Linearising restriction enzyme | Injected concentration | Resuspension media |
|---|---|---|---|---|---|---|---|---|---|
| *Krt8* | *Krt8* mRNA | gactatGAATTCgccaccATGTCCATCAGGGTGACTCAGAAATCC | GACTATgcggccgcTCACTTGGACACGACATCAGAAGAC | pRN3P | pRN3P:Krt8 | T3-RNApol | *SfiI* | 300ng/µl | HPLC water |
| *Krt8*-HA epitope tage (N-terminus) | *Krt8*-HA mRNA | gactatGAATTCgccaccATGgctacccatacgatgttcctgactatgctTCCATCAGGGTGACTCAGAAATC | GACTATgcggccgcTCACTTGGACACGACATCAGAAGAC | pRN3P | pRN3P:Krt8-HA_Nterm | T3-RNApol | *SfiI* | 300ng/µl | HPLC water |
| *Krt18* | *Krt18* mRNA | gactatCCCGGGgccaccATGAGCTTCACAACTCGCTCCACCA | GACTATgcggccgcTCAGTGCCTCAGAACTCTGGTG | pRN3P | pRN3P:Krt18 | T3-RNApol | *SfiI* | 300ng/µl | HPLC water |
| Cas9 (monomeric streptavidin/mSA fusion) | Cas9 mRNA | Cited in Gu et al., 2018 - PMID: 29889212 (acquired from Addgene, cat. #103882) | | pCS2+ | pCS2+Cas9-mSA | SP6-RNApol | *NotI* | 100ng/µl | HPLC water |
| Histone-H2B-RFP (C-terminus) | H2B-RFP mRNA | Acquired from authors: B. E. McGuinnes et al., 2009 - PMID: 19249208 | | pRN3 | pRN3:Hist1h2bb_RFP_Cterm | T3-RNApol | *SfiI* | 50ng/µl | HPLC water |
| Histone-H2B-YFP (C-terminus) | H2B-YFP mRNA | Acquired from authors: B. E. McGuinnes et al., 2009 - PMID: 19249208 | | pRN3 | pRN3:Hist1h2bb_Venus_Cterm | T3-RNApol | *SfiI* | 50ng/µl | HPLC water |

**Rhodamin-dextran conjugated beads/ RDB (70,000 MW tetramethylrhodamine-conjugated dextran beads)**

| Product description | Product abbreviation | Company Source | Cat. No. | Injected concentration | Resuspension media |
|---|---|---|---|---|---|
| 70,000 MW tetramethylrhodamine-conjugated dextran beads | RDBs | ThermoFisher | D1818 | 2µg/µl | HPLC water |

**Yellow-green fluorescent microspheres (for endocytic labelling of outer mebryo cells)**

| Product description | Product abbreviation | Company Source | Cat. No. | Media dilution | Resuspension media |
|---|---|---|---|---|---|
| 0.20µm Fluoresbrite YG microspheres | YGMs | Polysciences | 17151-10 (10ml) | 1:100 | BSA-free M2 |

**Supplementary Table S22:** Origin and concentrations of all microinjected constructs used in the study (siRNA, dsRNA, sgRNA, and recombinant mRNAs), including rhodamine-conjugated dextran beads/RDBs, and yellow-green fluorescent microspheres (for endocytic outer cell labelling). Provides supplier details or the primer sequences used to generate dsRNA IVT templates, or to amplify specific cDNA inserted cloned into the indicated plasmid IVT vectors (plus the restriction enzymes used to linearise plasmid DNA templates before IVT from the indicated RNA-polymerase promoter).

**Supplementary tables 23: antibodies used in IF (+fluorescently conjugated phalloidin)**

### Primary antibodies used

| # | ANTIGEN | cat. no. | supplier | species raised in and clonicity | dilution used | seconday antibody used (refer to below table) |
|---|---|---|---|---|---|---|
| 1 | TEAD4 | ab58310 | Abcam | mouse, polyclonal | 1:100 | B |
| 2 | PARD6B | sc-67393 | Santa Cruz | rabbit, polyclonal | 1:200 | C, D |
| 3 | TFAP2C | sc-12762 | Santa Cruz | mouse, monoclonal | 1:200 | B |
| 4 | CDX2 | MU392A-UC | BioGenex | mouse, monoclonal | 1:100 | B |
| 5 | YAP1 | sc101199 | Santa Cruz | mouse, monoclonal | 1:100 | A |
| 6 | CTNNB1 | 05-665 | Millipore | mouse, monoclonal | 1:200 | B |
| 7 | AMOT | 10061-1 | Gift from H. Sasaki group<br>Hirate et al., 2013; Curr. Biol. 23: 1181-94 | rabbit, polyclonal | 1:200 | C |
| 8 | KRT8 (TROMA-I antigen) | AB 531826 | Developmental Studies Hybridoma Bank (DHSB) | rat, monoclonal | 1:200 | E |
| 9 | HA-tag | ab9134 | Abcam | goat, polyclonal | 1:200 | F |
| 10 | Phallodin-Alexa$^{647nm}$ | A22287 | Invitrogen | N/A | 50X | N/A |

### Secondary fluorescent conjugated antibodies used

| # | SPECIES OF ANTIBODY TARGETED | cat. no. | supplier | species raised in and fluorophore | dilution used | used in combination with primary antibody (refer to above table) |
|---|---|---|---|---|---|---|
| A | mouse | A21202 | Invitrogen | donkey, Alexa488 | 1:500 | 5 |
| B | mouse | ab150107 | Abcam | donkey, Alexa647 | 1:500 | 1,3,4,6 |
| C | rabbit | ab150073 | Abcam | donkey, Alexa488 | 1:500 | 2,7 |
| D | rabbit | ab150075 | Abcam | donkey, Alexa647 | 1:500 | 2 |
| E | rat | A21208 | Invitrogen | donkey, Alexa488 | 1:500 | 8 |
| F | goat | ab150131 | Abcam | donkey, Alexa647 | 1:500 | 9 |

**Supplementary Table S23:** Details of the primary and secondary antibodies (and combinations) used in immuno-fluorescence (IF), including employed dilutions.

**Supplemenatrary Table 24: RT-qPCR oligonucleotide primer pair sequences**

| # | Gene | forward (5'-3') | reverse (5'-3') |
|---|---|---|---|
| 1 | *Tead4* | CGACAATGATGCAGAGGGTG | TCAGGATAATTTTGCGGCGG |
| 2 | *Gata3* | CCGAAACCGGAAGATGTCTA | AGATGTGGCTCAGGGATGAC |
| 3 | *Tfap2c* | CACGCGGAAGAGTATGTTGT | TAATTCTGCACTGCGGAGAC |
| 4 | *Cdx2* | TCAAGAAGAAGCAGCAGCAG | GCAAGGAGGTCACAGGACTC |
| 5 | *Krt8* | ACCGACGAGATCAACTTCC | CATAGACAGCACCACAGAC |
| 6 | *Krt18* | ACCTTCTCCACCAACTACC | CAGTCTCCTGTTCTCAGTTTC |
| 7 | *Rnd1* | CTTTGACATCAGCCGTCCAG | GTGTGCTGGGACAGTAGTCT |
| 8 | *Rnd3* | ATATGGCCAAGCAGATCGGA | GCAACTGCTGAGAGTTCTGG |
| 9 | *H2afz* | GCGCAGCCATCCTGGAGTA | CCGATCAGCGATTTGTGGA |

**Supplementary Tables S24:** Gene-specific RT-qPCR oligonucleotide primer sequences used (final concentration: 400nM).

### Supplementary Excel data and statistics workbook legends

**Supplementary data Excel workbook 1:** List of RNA-Seq identified significant DEGs (including genomic coordinates), unfiltered by minimum fold change expression level, between outer marked *Tead4* KD and unmarked outer cell clones within the same E3.5 embryo samples (a minimum RPMK >0.5 expression threshold in the unmarked clone was applied, and all sex-linked chromosomal genes were excluded). The RPMK expression value in unmarked control and marked *Tead4* KD outer clones, and accompanying fold expression changes, (plus, the false discovery rate/FDR statistic) are also given. Relates to data in Fig. 4.

**Supplementary data Excel workbook 2:** List of DEGs (RPMK >0.5 & fold change >2.0) identified in marked outer *Tead4* KD clones (E3.5) and their association with a TEAD4 chromatin immunoprecipitation (ChIP) peak identified in mouse TSCs (Trophoblastic Stem cells; as reported by Home et al., 2012[50]). DEG names and their chromosomal start and end position (and strand orientation), plus False Discovery Rate (FDR) statistic, are provided. The RPMK expression value in unmarked control and marked *Tead4* KD outer clones, and accompanying fold expression changes, are also given. Associations with a previously identified mouse TSC TEAD4 ChIP peak, at a given indicated genomic location, relative to the DEG are denoted by 0 (no association) or 1 (associated). Relates to data in Fig. 4 and S.Tabs 9.

**Supplementary data Excel workbook 3:** List of DEGs (RPMK >0.5 & fold change >10.0) identified in marked outer *Tead4* KD clones (E3.5) and their association with a TEAD4 chromatin immunoprecipitation (ChIP) peak identified in mouse TSCs (Trophoblastic Stem cells; as reported by Home et al., 2012[48]). DEG names and their chromosomal start and end position (and strand orientation), plus False Discovery Rate (FDR) statistic, are provided. The RPMK expression value in unmarked control and marked *Tead4* KD outer clones, and accompanying fold expression changes, are also given. Associations with a previously identified mouse TSC TEAD4 ChIP peak, at a given indicated genomic location, relative to the DEG, are denoted by 0 (no association) or 1 (associated). Relates to data in Fig. 4 and supplementary tables S.Tabs 9.

**Supplementary statistics workbook:** Statistical analyses were performed according to normality and experimental design. Parametric and non-parametric tests were applied to normally and non-normally distributed data, respectively. Pairwise t-tests were applied to within-group analyses, and unpaired t-tests were applied to between-group analyses, as indicated. Details of t-tests and corresponding p-values are provided.
